# Mapping AAV capsid sequences to functions through function-guided *in silico* evolution

**DOI:** 10.1101/2024.10.11.617764

**Authors:** Binjie Guo, Hanyu Zheng, Xurong Lin, Haohan Jiang, Aisheng Mo, Hongyan Wei, Ru Zhang, Yubin Yuan, Yile Wu, Hengguang Li, Yunshuo Zhang, Zhuoyuan Song, Xuebin Ni, Yan Huang, Bin Yu, Yumin Yang, Xiaosong Gu, Zuobing Chen, Ningtao Cheng, Xuhua Wang

## Abstract

Artificial intelligence (AI) offers significant potential to accelerate the functional engineering of adeno-associated virus (AAV) capsids in a time- and cost-efficient manner. However, existing approaches lack the ability to systematically map capsid sequences to multifunctional properties. To address this challenge, we developed ALICE, a knowledge-driven platform for *in silico* AAV capsid engineering that directly maps sequence to multiple functions. Our method incorporating a heuristic algorithm with contrastive learning principles, termed function-guided evolution (FE), to iteratively optimize high-performing capsid sequences generated by a naïve language model toward desired functions. We elucidate FE’s evolutionary mechanism, demonstrating its ability to navigate a function-guided landscape and generate capsids with tailored properties. After validating ALICE’s ability to design murine-specific AAV capsids targeting Ly6a/Ly6c1 with limited training data, we developed ALICE-X by integrating a curriculum learning strategy. This upgrade facilitated exploration of sequence space beyond existing wet-lab datasets, enabling successful targeting of the human transferrin receptor 1 (hTfR1). The resulting engineered AAV variant, AAV.ALICE-H3, demonstrated improved viability, specific receptor targeting, and ∼ 251-fold enhanced CNS tropism in human *TFRC* knock-in (*hTFRC* KI) mice compared to wildtype controls. This interpretable, knowledge-driven framework advances *in silico* AAV capsid design, offering broad applicability for translational gene therapy.

## Introduction

Recent advances in artificial intelligence (AI) have greatly expanded its applications in functional protein design^1–4^. However, the computational design of multifunctional, large natural protein complexes, such as adeno-associated virus (AAV) capsids, remains a formidable challenge^5^. Over the past decade, conventional methodologies like rational design^6–9^ and directed evolution^10–15^ have been developed to engineer AAV capsids to overcome limitations such as low viability and suboptimal tissue tropism. Nevertheless, these methods are constrained by their reliance on labor-intensive wet-lab experimentation, limiting their ability to comprehensively explore the vast sequence landscape of AAV variants^16,17^.

Recent progress in computational protein engineering, particularly through sequence-to-function mapping^18–23^, offers a promising alternative. As our understanding of AAV capsid mechanisms deepens^24–27^, we hypothesized that leveraging capsid-receptor binding data could enable the in silico prediction of functional outcomes, facilitating the design of optimized capsids. However, two key challenges persist: (i) the lack of a computational framework capable of predicting multiple functional properties from a single sequence, especially beyond the restricted sequence space defined by limited training data^28^, and (ii) the need for interpretable models that elucidate how capsid sequences evolve to achieve desired functions^16,29^.

To overcome these challenges, we developed AI-driven Ligand-Informed Capsid Engineering (ALICE), a generative AI architecture that map capsid sequences to multiple functions. ALICE begins with a naïve generative language model pretrained on 2.89 million peptide sequences to capture natural peptide linguistics. Through transfer learning^30^, the model is then fine-tuned on AAV capsid sequences to learn their biological principles. However, a naïve language model alone cannot effectively associate a single sequence with multiple functions. To overcome this limitation, we introduced function-guided evolution (FE), a heuristic algorithm incorporating contrastive learning principles, which steers sequence evolution toward desired functional properties, even when training data are scarce (∼ 100 entries). The FE module is trained on functional data, including capsid production fitness and binding capability to Ly6c1 and Ly6a receptors, which are enriched at the blood-brain barrier (BBB) in mice. By leveraging different algorithmic approaches within the FE module, we elucidated the evolutionary mechanisms guiding capsid sequences toward specific functions. Using these insights, we directed the FE module to optimize capsid sequences for enhanced viability and CNS tropism. Remarkably, ALICE-generated AAV.ALICE-N2 demonstrated an approximate two-fold increase in viability and a ∼372-fold enhancement in CNS cell transduction efficiency of mice compared to the original AAV9.

To enable translational applications, we upgraded ALICE to ALICE-X by incorporating a curriculum learning strategy^31^ into the FE module. This advancement enables the design of capsid sequences featuring novel motifs that target the human transferrin receptor (hTfR1), a critical mediator of blood-brain barrier (BBB) penetration. This curriculum learning framework decomposes the complex capsid design process into two phases: (1) a function-integration phase, where contrastive learning with positive and negative references is used to integrate different functional objectives; and (2) an exploration phase, where the search expands into new sequence space while the evaluation model ensures that multi-functional performance remains balanced and does not diverge. This strategy allows ALICE-X to explore sequence spaces beyond the limitations of wet-lab data. We further elucidated how ALICE-X’s curriculum learning balances novel sequence exploration with multi-functional integration. Using this approach, ALICE-X successfully designed novel AAV variants, including AAV.ALICE-H1-H3, which incorporates the previously unexplored ’YTK’ motif. These variants exhibit enhanced hTfR1-targeting capability and CNS tropism, surpassing the constraints of traditional wet-lab experimentation. Notably, AAV.ALICE-H3 demonstrated a ∼1.5-fold improvement in viability over the original AAV9 and exhibited ∼251-fold greater selectivity in transducing brain cells in human *TFRC* knock-in (*hTFRC* KI) mice compared to wild-type mice, underscoring its clinical potential. Collectively, ALICE-X represents a breakthrough in interpretable in silico AAV engineering, overcoming the empirical limitations of wet-lab methods and establishing a computational framework for designing multifunctional protein complexes.

## Results

### Workflow of ALICE for engineering AAV capsids for Ly6a/Ly6c1 targeting

It is believed that the amino acid sequences within proteins share an intrinsic resemblance with the architecture of human language, with each sequence akin to the alphabet of a verbal dialect.^1,2^ In our pilot experiment, we found that the peptide sequences used for the insertion modification of AAV capsids, which exhibit high production fitness and binding capability for Ly6a and Ly6c1 receptors, display similar architectures of protein and human language (Supplementary Fig. 1 and Supplementary Table1). This finding motivated us to pretrain a generative language model with a large peptide sequence dataset (Fig. 1a) and then semantically tune it with a limited AAV insertion peptide sequence dataset to generate functional capsid sequences (Fig. 1b). We then attempted to use a ranking filtration process module to rank the generated sequences on the basis of their multifunctional properties for further evolution (Fig. 1c). By incorporating a heuristic algorithm^32,33^ with contrastive learning principles,^34^ we subsequently sought to steer the evolution of the top-performing sequences generated by the naïve language model toward the desired functions (Fig. 1d). Finally, we sought to evaluate the comprehensive functions of the sequences through an hits-ranking module (Fig. 1e) and select the top candidates for wet experiment verification (Fig. 1F). Thus, the workflow of ALICE integrates 6 essential steps: (i) pretraining, (ii) semantic tuning, (iii) ranking filtration process (RankFlitPro), (iv) function-guided evolution (FE), (v) hits-ranking, and (vi) wet-lab validation (Fig. 1).

**Fig. 1.**
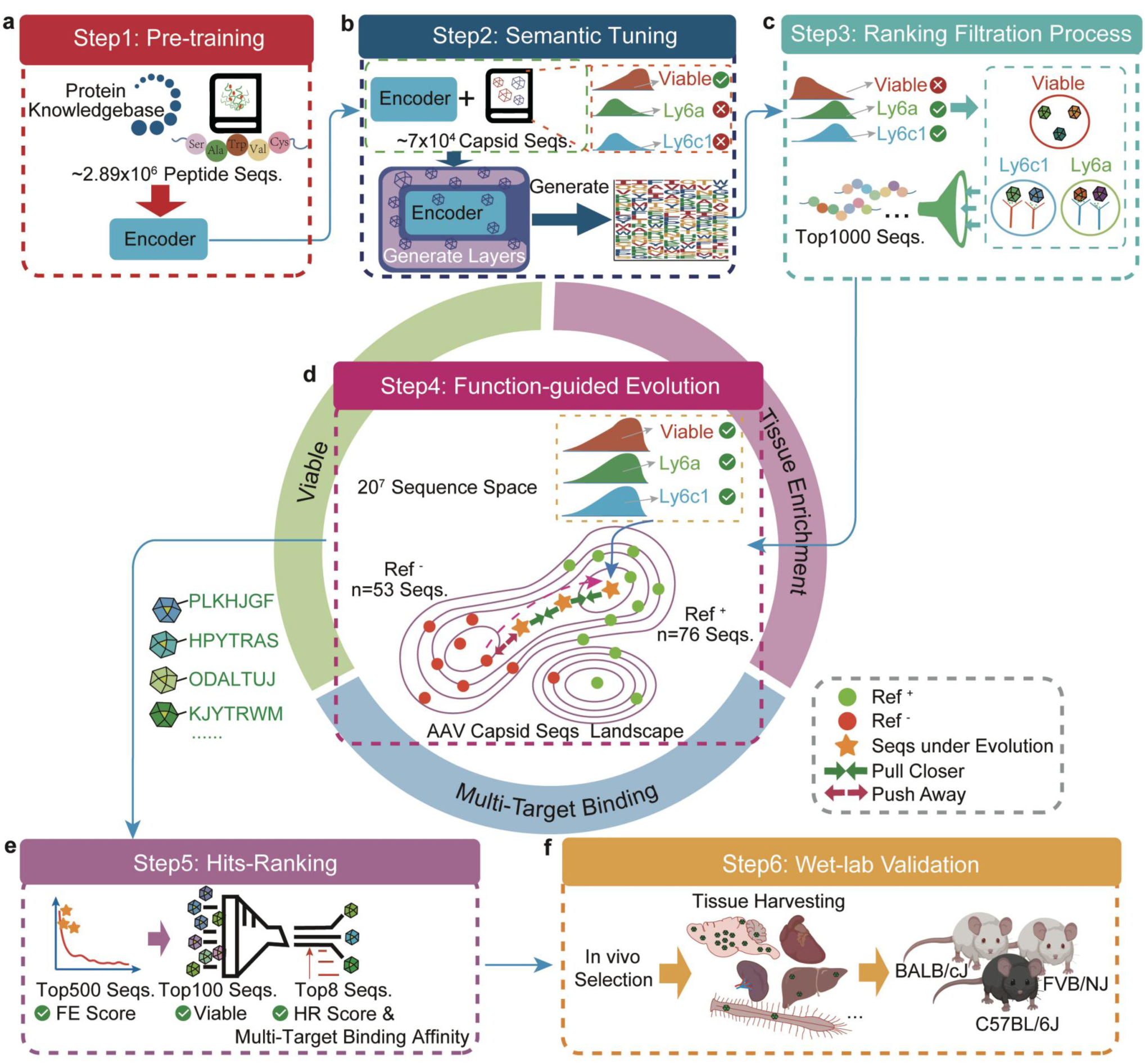
Workflow of the ALICE platform in mapping capsid sequences to functions. The workflow of the ALICE platform integrates 6 steps, including pretraining, semantic tuning, the ranking filtration process, function-guided evolution, hits-ranking, and wet-lab validation. **a,** A language model learns the foundational knowledge of peptide sequences in the pretraining step. **b,** During the semantic tuning step, the language model learns the distribution of AAV capsid sequences in the training dataset, which exhibit high viability but low binding affinity for Ly6a or Ly6c1. Subsequently, the generative model designs AAV capsid sequences with improved semantic representation and diverse functional properties. **c,** The Ranking Filtration Process (RankFiltPro) module selects the top 1,000 capsid sequences, prioritizing high production fitness and strong binding affinity for both Ly6a and Ly6c1. Although these sequences still exhibit relatively low viability, they provide an optimal initialization for the subsequent FE Module. **d,** The function-guided evolution (FE) module constructs a conformational AAV capsid sequence evolution landscape in which capsid sequences evolve and map to multiple functions of viability and multiple targeting from a large potential sequence space. **e,** In the hits-ranking (HR) module, the top 500 sequences were first selected, prioritizing the FE score. The top 100 sequences were then selected, prioritizing viability. Finally, the top candidate capsids were selected, prioritizing the HR score and the target binding affinity of both Ly6a and Ly6c1. **f,** In the wet-lab validation step, the potency of the AAV variants engineered by ALICE was evaluated through in vivo experiments using three distinct mouse strains: BLAB/cJ, C57B/6J and FVB/NJ.

### Generation of datasets and establishment of virtual evaluation methods

To train ALICE, we generated four main datasets: (i) 2.89 million peptide sequences from UniProt^35,36^ (***D***_***Pretrain***_) for pretraining (details in Supplementary Methods) and (ii) a publicly accessible dataset of 7-mer amino acid sequences, which were inserted between residues 588-589 in VP1 for AAV9 capsid modification: ***Lib***_***seq***_ These data include capsid sequence insertion-modified variant production fitness, binding capability to the target proteins Ly6a (widely expressed in C57BL/6J and FVB/NJ mice) and Ly6c1 (expressed in Ly6a-deficient BALB/cJ mice).^25^ The capsid sequences with high production fitness and target binding ability are selected for semantic tuning in the dataset of ***D***_***seq***_ (n = 72,753, details in Supplementary Methods). (iii) To promote capsid sequences evolution toward desired functions, we prioritized sequences in the FE reference dataset by focusing on two key factors: (1) high viability and strong binding capability to Ly6a and Ly6c1, and (2) AAV variants preformed desired functions in animals reported in previously studies.^8,12,37,38^ Sequences meeting these criteria were designated as ***Ref***\* (n = 76, Supplementary Table2), while those exhibiting weak binding to either Ly6a or Ly6c1 and low viability were classified as ***Ref***^+^ (n = 53, Supplementary Table2, detailed in the Supplementary Methods). (iv) To evaluate the production fitness, Ly6a binding, and Ly6c1 binding capabilities of the designed capsid sequences, three datasets (***D***_***Production Fitness***_, ***D***_***Ly6a***_ and ***D***_***Ly6c1***_) were constructed from ***Lib***_***seq***_(details in the Supplementary Methods).

To virtually evaluate the functionalities of the sequences designed by each module within ALICE and thereby optimize them, we established predictive methodologies to comprehensively examine the functionalities of the designed sequences. This includes evaluating the fitness of the sequence insertion-modified variants, as well as their ability to bind to the targets of Ly6a or Ly6c1. To identify the best prediction model, we selected a total of seven widely used candidate machine learning models, including support vector regression (SVR),^39^ eXtreme gradient boosting (XGBoost),^40^ K-nearest neighbors (KNN),^41^ random forest (RF),^42,43^ gradient boosting (GB),^44,45^ Bayesian Ridge,^46^ and AdaBoost.^43^ To evaluate the performance of each model, six evaluation indicators (Pearson, Spearman, RMSE, R^2^, MedAE, and C-index) were selected. We trained and tested these models on the ***D***_***Production Fitness***_ (Supplementary Fig. 2a), ***D***_***Ly6a***_(Supplementary Fig. 3a) and ***D***_***Ly6c1***_(Supplementary Fig. 4a) datasets to predict production fitness, Ly6a binding, and Ly6c1 binding capabilities, respectively (details in the Supplementary Methods). While XGBRegressor is generally robust for regression tasks, the negative R² values (Supplementary Fig. 3e and Supplementary Fig. 4e) indicate that its predictions performed worse than using a simple mean predictor, suggesting a significant model-data mismatch. Perhaps, the model has captured noise in the training data, leading to poor generalization on test data despite apparent training correlations. The R² penalized deviations in absolute prediction accuracy, which may disproportionately affect models like XGBRegressor that prioritize complex, nonlinear feature interactions over mean-squared error minimization. In contrast, the predictive performance of the GB model was the best among these candidate models (Supplementary Fig. 2-4), with Pearson correlations of 0.96, 0.81, and 0.81, respectively (Supplementary Table 4-6). Therefore, we use the GB model in the following evaluation tasks.

### The pretraining followed by a semantic-tuning strategy confronts a multifeature fusion conflict issue

To enable a language model to learn and generate functional capsid sequences, we selected BERT^47^ and RoBERTa^48^ for the pretraining module (Fig. 1a; Supplementary Methods) and incorporated a SeqGAN architecture^49,50^ for semantic tuning (Fig. 1b). To assess the necessity of pretraining, we trained SeqGAN directly on ***D***_***seq***_without pretraining (SeqGAN-only), and we further embedded BERT- and RoBERTa-based pretrained models trained on ***D***_***Pretrain***_into the SeqGAN framework (Extended Data Fig. 1a).

Through uniform manifold approximation and projection (UMAP)^51^ analysis of the properties of the generated sequences, we found that the features of the sequences generated by the RoBERTa-SeqGAN are more segregated than those generated by the SeqGAN-only or BERT-SeqGAN (Extended Data Fig. 1b-d). This suggests that RoBERTa-SeqGAN can generate sequences with features different from those in the original ***D***_***seq***_. Meanwhile, amino acid usage in RoBERTa-SeqGAN outputs showed the highest correlation with the training dataset (Pearson = 0.835 vs 0.643 or 0.158; Supplementary Fig. 5a-f), indicating effective learning of semantic and syntactic patterns from natural analogues.

Virtual evaluation with the GB model showed that SeqGAN-only failed to improve production fitness or Ly6a/Ly6c1 binding relative to training sequences (Extended Data Fig. 1e-h, j), underscoring the critical value of pretraining. BERT-SeqGAN modestly enhanced production fitness but at a substantial cost to Ly6a/Ly6c1 binding (Extended Data Fig. 1e-h, k). Conversely, RoBERTa-SeqGAN improved predicted binding to both Ly6a and Ly6c1 but exhibited markedly reduced production fitness (Extended Data Fig. 1e-h, l). These trends mirror the antagonistic relationship between binding capability and production fitness observed in natural capsids (Extended Data Fig. 1i), revealing a fundamental multifeature fusion conflict when mapping a single peptide to multiple functional requirements using a naïve language model.

To determine whether this conflict stems from limitations intrinsic to RoBERTa-SeqGAN, we evaluated alternative sequence-generation architectures (Supplementary Methods). Across all models tested, the same trade-off persisted (Supplementary Fig. 6), indicating that conventional generative models primarily capture natural sequence regularities and therefore reproduce inherent evolutionary constraints rather than introducing model-specific artifacts.

### Promotion of the evolution of capsid sequences via a function-directed evolution strategy

Inspired by biological evolution, where organisms achieve adaptive breakthroughs under environmental pressures^52^ and learn from limited samples—we developed an algorithmic paradigm that simulates natural evolutionary processes to address the challenge of fusing multiple functional features into a single sequence. We first applied a ranking filtration module to select the top 1,000 RoBERTa-SeqGAN-generated capsid sequences with high predicted binding capability and production fitness for downstream evolution (Fig. 1c; Supplementary Methods). We then introduced a function-guided evolution (FE) module that integrates a heuristic algorithm^32,33^ with contrastive learning principles^34^ to iteratively optimize sequences toward desired functions.

During optimization, the FE module computes MSA similarity between evolving sequences and predefined positive (***Ref***\*, n = 76) and negative (***Ref***^+^, n = 53) reference sets, functionally mimicking contrastive learning by encouraging proximity to functional sequences while discouraging similarity to non-functional ones. The ***Ref***\* and ***Ref***^+^ datasets are distinguished by production fitness, multitarget binding capacities and in vivo-validated capsid sequences (Supplementary Fig. 7a; Supplementary Table2). Evolutionary progress is quantified using multi-sequence alignment-based similarity to or , and the FE function is defined as:

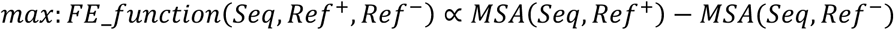

where denotes the Smith-Waterman algorithm^53^. This function enables adaptive search over sequence landscapes and provides the basis for the functional evolution score (FE score; Supplementary Methods), which reflects traits inferred from both ***Ref***\*and ***Ref***^+^reference sets.

To identify suitable models for the FE module, we evaluated four heuristic algorithms—simulated annealing (SA)^33^, genetic algorithm (GA)^32^, estimation of distribution algorithm (EDA)^54^, and a hybrid EDA-GA method (EDG)^54^. Only EDA and EDG converged rapidly (Supplementary Fig. 7b), yielding high FE scores and producing concentrated sequence distributions (Supplementary Fig. 7c-f), consistent with their superior optimization performance. Therefore, subsequent development of the FE module focused on the EDA and EDG frameworks.

### Refining the FE module by exploring the evolutionary mechanisms of capsid sequences within the model

To elucidate the mechanisms governing sequence evolution within the FE module, we examined the evolutionary trajectories of capsid sequences across training epochs (Extended Data Fig. 2a-d). By plotting the distances between the capsid sequences under evolution and the positive references (***Ref***\* ) against those with negative references (***Ref***^+^ ), we visualized the evolutionary trajectory across multiple well-established metrics (details in the Supplementary Methods, section “Metrics”): sequence evolutionary divergence (Extended Data Fig. 2a, Kullback‒Leibler (KL) divergence),^55^ functional similarity (Extended Data Fig. 2b, Blocks Substitution Matrix (BLOSUM) 62 similarity),^56^ and both local (Extended Data Fig. 2c, N-gram similarity)^57^ and global (Extended Data Fig. 2d, longest common subsequence (LCS)^58^ semantic similarity. We designated the evolutionary changes with reference to ***Ref***\*and ***Ref***^+^as ΔPE (positive evolutionary change) and ΔNE (negative evolutionary change), respectively.

Ideally, FE should guide sequences toward ***Ref***\* while diverging from ***Ref***^+^(Extended Data Fig. 2a, b, green arrows). However, the EDA-based approach initially drove evolution in a direction opposite to the ideal trajectory during the first five epochs (Extended Data Fig. 2a, b) and then reversed course by ∼180°, ultimately yielding sequences with reduced correlation to positive samples. Similar behavior was observed across semantic metrics (Extended Data Fig. 2c, d), indicating a form of partial or unstable evolution (Extended Data Fig. 2a, b). In contrast, the EDG-based approach consistently steered sequences toward ***Ref***\*and away from ***Ref***^+^ , particularly during early epochs (Extended Data Fig. 2a, b). The initial 5-epoch phase displayed an approximately balanced ΔPE:ΔNE ratio for KL divergence (Extended Data Fig. 2a). After epoch 5, EDG continued to promote similarity to ***Ref***\* while moderately increasing similarity to ***Ref***^+^, yielding final ΔPE:ΔNE ratios of ∼1.5:1 (KL divergence) and 3.7:1 (BLOSUM62) (Extended Data Fig. 2a, b). Notably, functional similarity to ***Ref***\*increased substantially (from −6.578 to 1.439), whereas similarity to ***Ref***^+^shifted only modestly (−8.002 to −5.853) (Extended Data Fig. 2b), indicating that FE preferentially incorporates beneficial features from ***Ref***\* while making minimal trade-offs with respect to ***Ref***^+^.

An additional and intriguing pattern emerged: EDG-guided evolution increased similarity to both ***Ref***\*and ***Ref***^+^under local and global semantic metrics (Extended Data Fig. 2c, d; Supplementary Fig. 8). We attribute this to intrinsic properties of capsid sequences. Although ***Ref***\*and ***Ref***^+^differ functionally, both are naturally occurring and likely follow the shared structural and semantic characteristic of capsid proteins ^59,60^. The EDG-based FE module appears to capture this shared “grammar”, while the functional distinctions between ***Ref***\*and ***Ref***^+^manifest as different “expressions” of that grammar, consistent with prior observations^61^. This enables the model to generate diverse sequences^1,18^ that conform to naïve capsid rules while exploring the functional spectrum between ***Ref***\*and ***Ref***^+^ . Comparable results obtained using single positive and negative sequences for trajectory tracking (Supplementary Fig. 9) further support these conclusions. Based on these findings, we adopted the EDG algorithm for the FE module in subsequent analyses.

#### Resolving the conflict in multifunction fusion through evolution across a function-guided landscape

To determine the biological implications of the capsid sequences generated through the selected EDG-based approach, we sought to analyze them from both functional and semantic perspectives. We found, from a functional perspective, that despite similar evolutionary distances to positive and negative samples, these sequences exhibited greater functional similarity to ***Ref***\* (Fig. 2a, top panel). Interestingly, we found that the capsid sequences under evolution exhibit a heightened sensitivity to local semantic information (N-gram similarity, Fig. 2a, bottom panel), as opposed to global information (LCS ratio, Fig. 2a, bottom panel), in agreement with previous observations that the functionality of proteins is dictated by specific regions or motifs within the sequence (local semantic information) rather than the overall sequence semantics^62,63^. This sensitivity to local motifs is crucial, as the AAV capsid protein often contains specific motif substructures.

**Fig. 2.**
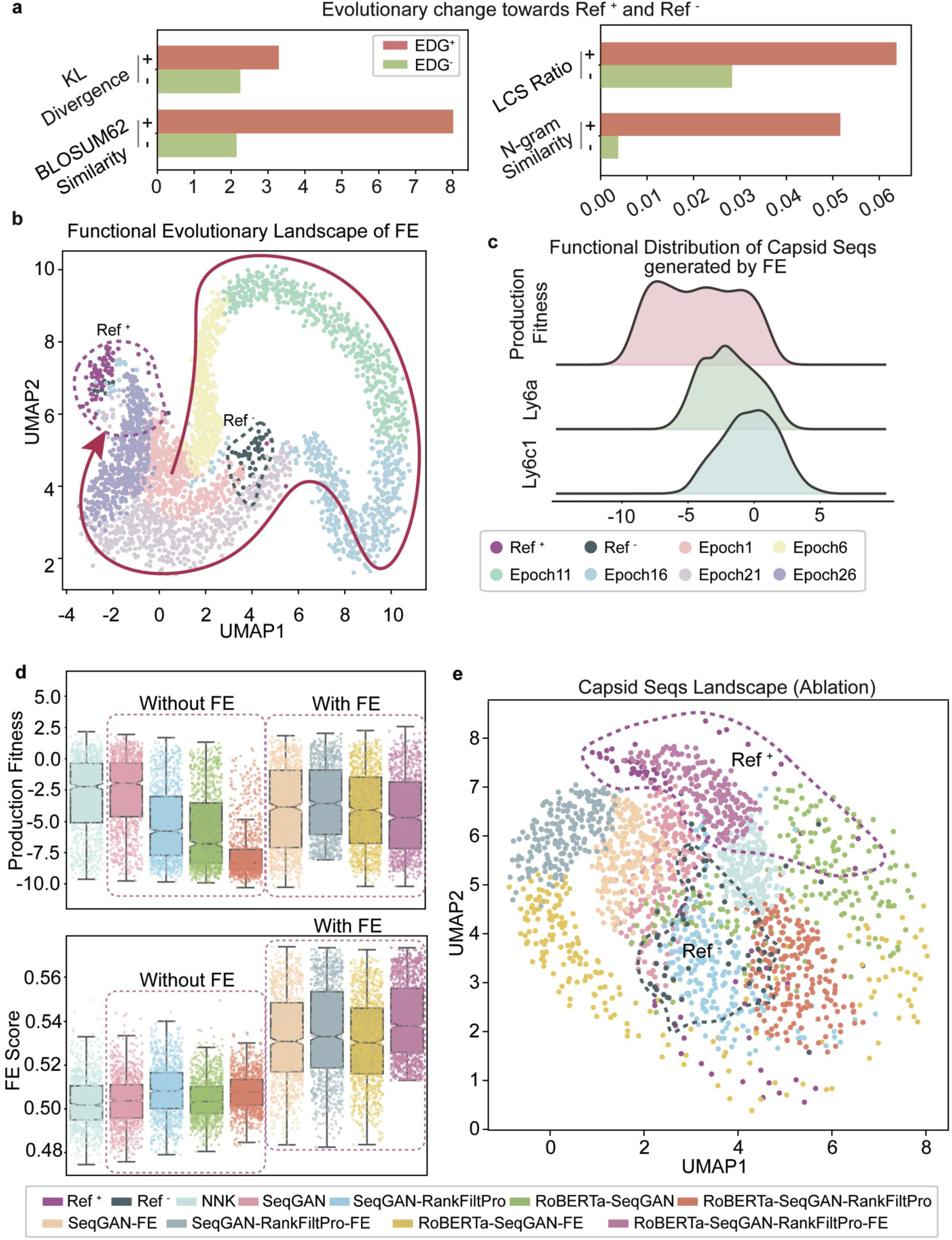
The FE module steers the evolutionary trajectory of sequences toward positive samples with desired functions and ablation study. **a,** Evolutionary changes in the sequences toward ***Ref***\* or ***Ref***^+^ engineered via the EDG model. **b,** Conformational evolution landscape of capsid sequences engineered with EDG-based FE at different epochs, showing progression toward ***Ref***\* and divergence from ***Ref***^+^. The colors of the dots represent the outcomes after different epochs, as indicated by the color bar provided at the bottom. **c,** The plot shows the functional distribution of FE-designed capsid sequences, including production fitness and Ly6a or Ly6c1 binding capacity. **d,** Performance of production fitness (top) and the FE score (bottom) across architectures with different ablations. The colors of the dots and histograms represent the outcomes from architectures with distinct ablations, as indicated by the color bar provided at the bottom. **e,** The diagram illustrates the functional distribution of the capsid sequence landscape engineered by architectures with different ablations. The colors of the dots represent the outcomes from architectures with distinct ablations, as indicated by the color bar provided at the bottom.

To visualize the evolutionary process of capsid sequences within the model, we depicted the dynamic landscape of capsid sequences as they evolved from epoch 0 (near ***Ref***^+^) to epoch 25 (close to ***Ref***\*) (Fig. 2b). We found that throughout the evolutionary process, both the FE score and production fitness of the sequences improved significantly (Supplementary Fig. 10a). We also found that FE ensures that evolved capsids adhere to specified functional goals without introducing unintended features (Supplementary Fig. 11a-d). As sequence diversity gradually increased, the models converged on variants containing the ’GYS’ motif, which is also among the most frequent and high-fitness patterns in the ***Ref***\*dataset (Supplementary Fig. 10b), indicating that ALICE successfully recovers and amplifies beneficial motifs present in the training data rather than generating patterns beyond those observed in the reference sequences. Thus, the FE adeptly recognizes local semantic structures in close proximity to positive samples, thus gradually exploring the functional motifs related to positive samples while diverging from negative ones (Supplementary Fig. 10b). This approach successfully integrated production fitness and multiple-target binding ability simultaneously into the capsid sequences (Fig. 2c). In conclusion, by evolving across a function-guided landscape informed by the ***Ref***^!^ dataset (experimentally validated high-fitness, high-affinity capsids), FE effectively resolved the intrinsic conflict of multifunctional integration (Extended Data Fig. 1i-l), thereby enabling a single sequence to be simultaneously mapped onto multiple functions.

#### An ablation study interprets the contribution of each module to the evolutionary landscape

To elucidate the contribution of each module within ALICE, we performed an ablation study across nine recombinant architectures for mapping capsid sequences to functions. Sequence performance was evaluated using production fitness prediction and FE scores. The RoBERTa+SeqGAN+RankFiltPro+FE (RSRF) architecture achieved the highest FE score while retaining strong production fitness (Fig. 2d). In UMAP space, sequences from different architectures formed distinct clusters (Fig. 2e). Architectures lacking the FE module clustered toward ***Ref***^+^ and away from ***Ref***\*, consistent with observations that naïve generative models cannot resolve multi-feature fusion conflicts (Extended Data Fig. 1i-l). Conversely, FE-containing combinations consistently shifted toward ***Ref***\*, with RSRF showing the closest proximity (Fig. 2e), underscoring the FE module’ s role in steering functional evolution (Fig. 2c). FE further identified sequences exhibiting low, stable single-site entropy at positions 4-7 (Supplementary Fig. 10a), and the RSRF configuration optimized this balance between diversity and specificity (Supplementary Fig. 10b). Motif analysis revealed recurrent GYSS signals in the terminal four positions (Supplementary Fig. 10b), consistent with prior pulldown evidence implicating this motif in Ly6c1 interaction^25^.

To examine how FE shapes ablation outcomes, we analyzed evolutionary trajectories relative to ***Ref***\*and ***Ref***^+^ , defining four trajectory types (Extended Data Fig. 2e) using KL divergence, BLOSUM62 similarity, N-gram, and LCS metrics. NNK-derived sequences served as controls. Combinations lacking FE displayed Type III trajectories—progressive divergence from both ***Ref***\* and ***Ref***^+^ (Supplementary Fig. 13a-d)—indicating reduced mapping performance (Fig. 2d). In contrast, architectures containing FE frequently showed Type I behavior: semantic approaches toward both ***Ref***\* and ***Ref***^+^, but more strongly toward ***Ref***\* (Supplementary Fig. 12d, e; Supplementary Fig. 13c, d). This pattern indicates that FE captures shared “capsid grammar” underlying both sets while retaining sensitivity to functional motifs distinguishing ***Ref***\*and ***Ref***^+^, such as GYSS. This balance enables FE to generate sequences functionally aligned with ***Ref***\*(supported by KL divergence and BLOSUM62) while exploring new semantic spaces (Supplementary Fig. 14).

Functionally, architectures incorporating FE but lacking pretraining exhibited Evo-Type I behavior—evolving toward both ***Ref***\*and ***Ref***^+^(Extended Data Fig. 2g, yellow arrow). When both FE and pretraining were present, architectures shifted toward Type II behavior—evolving toward ***Ref***\*while diverging from ***Ref***^+^(Extended Data Fig. 2f, blue arrow; Supplementary Fig. 13a). This demonstrates that pretraining reinforces FE-guided multifunctional fusion. Moreover, FE-containing architectures mapped sequences with motif segments satisfying multi-functional requirements while maintaining distributions distinct from the training data (Supplementary Fig. 14a, b), thus balancing novelty and functional enhancement. Collectively, these results demonstrate that the FE and pretraining modules function synergistically to guide capsid sequence evolution toward the desired multifunctional profiles.

#### In vivo experiments verified the potency of AAV variants engineered by ALICE

To validate ALICE-designed variants, we used the hits-ranking module to prioritize sequences with strong multifunctional potential for wet-lab testing (Fig. 1e; Supplementary Methods). From the top 500 candidates ranked by FE score, the top 100 were further selected based on predicted production fitness. Eight leading variants were then identified using HR scores and predicted Ly6c1/Ly6a binding (Fig. 3b). Notably, these eight sequences clustered closer to ***Ref***\* and away from ***Ref***^+^, suggesting potential CNS tropism (Fig. 3c). Variants carrying these sequences inserted at VP1 positions 588-589^11,12^ were packaged and designated AAV.ALICE-N1 through N8. All variants carried an AAV-EF1α-

**Fig. 3.**
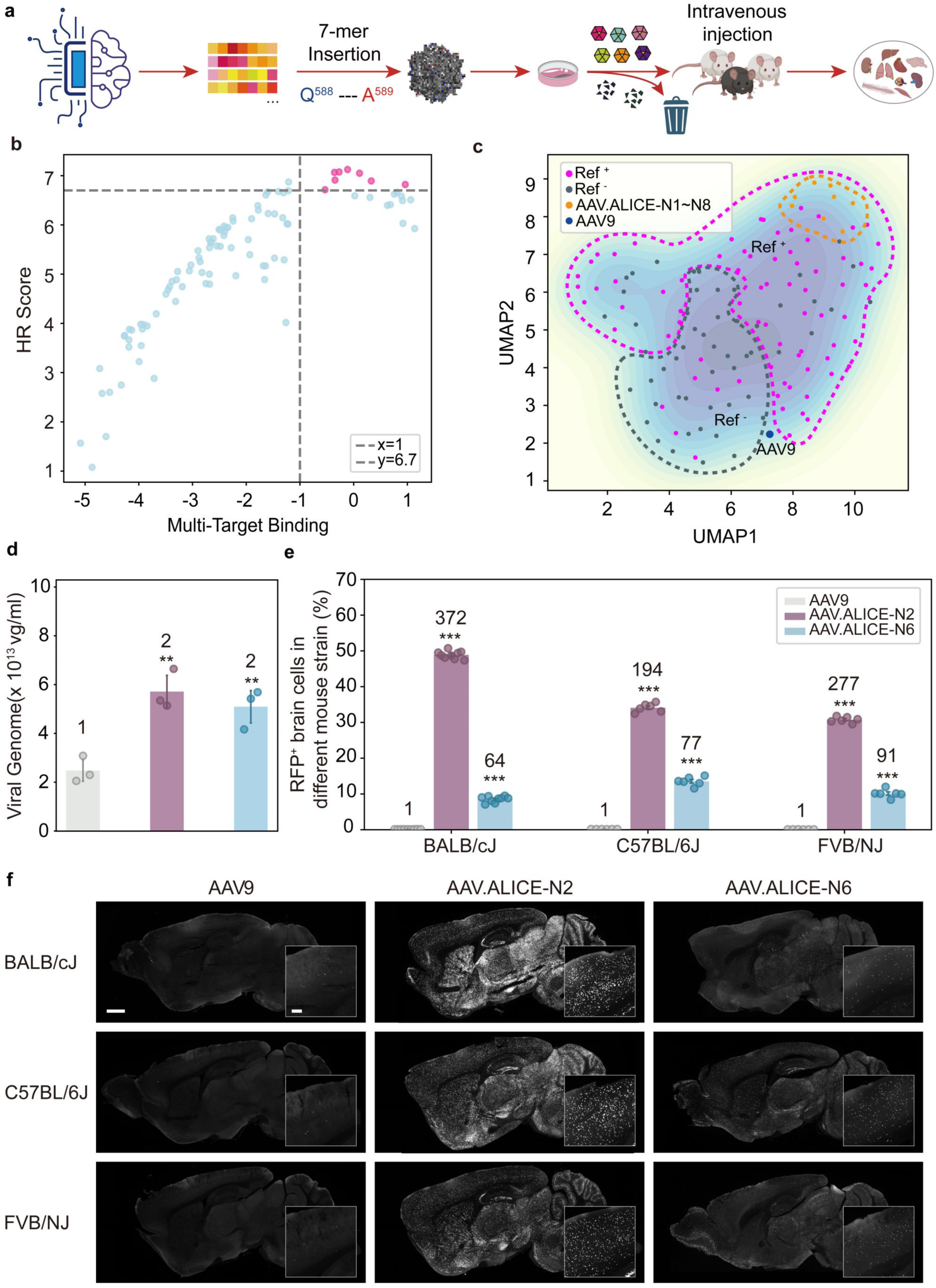
An *in vivo* study verified the ability of AAV variants engineered by ALICE to transduce CNS cells from different mouse strains. **a,** Workflow diagram of the wet laboratory validation process of the ALICE platform. **b,** The plot delineates the criteria employed for the selection of 8 candidates for in vivo experimental verification, focusing on those demonstrating high HR scores and multitarget binding capacities. **c,** The diagram illustrates the distribution of selected candidate sequences within the capsid sequence feature landscape in comparison to the sequences from the datasets of ***Ref***\*, ***Ref***^+^, and AAV9. **d,** Comparison of the titers of AAV9 and AAV.ALICE-N2 and AAV.ALICE-N6. The data are presented as the means ± s.e.m., n = 3 biologically independent replicates. The color of the bar indicates the results of different AAVs, as indicated by the color bar provided in (e). **e,** Quantification of the percentage of total RFP^+^ brain cells and the fold change relative to AAV9 in BALB/cJ, C57BL/6J, and FVB/NJ mice after intravenous injections of AAV variants (5 × 10^11^ vg per animal). The data are presented as the means ± s.e.m; n = 3-4 mice per group. Statistical significance was determined via one-way ANOVA with Tukey’s post hoc test (*P < 0.05, **P < 0.01, ***P < 0.001 versus AAV9). The color of the bar indicates the outcome of treatment with different AAVs, as indicated by the provided color bar. **f,** Representative images of brain regions in BALB/cJ, C57BL/6J, and FVB/NJ mice showing RFP-labeled cells (white dots) 21 days after intravenous administration of AAVs-EF1α-H2B-RFP (AAV9 on the left, AAV.ALICE-N2 in the middle, AAV.ALICE-N6 on the right; 5 × 10^11^ vg per animal). Scale bars: 1,000 μm for low-magnification images and 200 μm for enlarged images.

H2B-RFP reporter cassette to visualize transduced cells. Under identical production conditions, AAV.ALICE-N2, AAV.ALICE-N6 and AAV.ALICE-N8 exhibited ∼2-fold higher titers than AAV9 (Fig. 3e; Supplementary Table14), whereas AAV.ALICE-N3, N4 and N6 showed titers comparable to AAV9. AAV.ALICE-N1 and N5 generated insufficient titers (∼9×10¹⁰ vg ml⁻¹) and were excluded from in vivo studies.

To assess CNS tropism, BALB/cJ mice were intravenously injected with AAV9 or AAV.ALICE-N2, N3, N4, N6, N7 or N8 (5×10¹¹ vg per animal). Three weeks later, RFP signals were examined in brain tissue. Only sparse transduction was detected in mice treated with AAV9, N3, N4, N7 or N8 (Fig. 3f, g; Supplementary Fig. 15a-d). In contrast, AAV.ALICE-N2 and AAV.ALICE-N6 yielded robust brain transduction, exhibiting 372-fold and 64-fold increases over AAV9, respectively (Fig. 3f, g). Their enhanced tropism was further confirmed in C57BL/6J and FVB/NJ mice (Fig. 3f, g).

Cell-type analysis showed higher percentages of RFP-labeled cells in multiple brain regions following AAV.ALICE-N2 and AAV.ALICE-N6 treatment compared with AAV9 (Supplementary Fig. 16a-c). Approximately 30% of AAV.ALICE-N2-transduced cells were NeuN⁺ neurons, ∼40% were Olig2⁺ oligodendrocytes, and ∼3% were GFAP⁺ astrocytes (Supplementary Fig. 17a, b; Supplementary Fig. 18a). Across sampled regions in adult BALB/J mice, AAV.ALICE-N2 transduced 19-51% of neurons, 26-54% of oligodendrocytes and 1-7% of astrocytes (Supplementary Fig. 17b, Supplementary Fig. 18a).

Improved gene delivery was also observed in the spinal cord. AAV.ALICE-N2-EF1α-H2B-RFP labeled 8-12% of spinal cord cells versus 0-1% for AAV9, and 9-10% of ChAT⁺ motor neurons versus 0-1% for AAV9 (Supplementary Fig. 18b-d), demonstrating efficient targeting of spinal neurons. Systemic administration additionally revealed transduction in liver, skeletal muscle, heart and spleen for AAV.ALICE-N2 and AAV.ALICE-N6 (Supplementary Fig. 19a-c), with lower levels detected in kidney, lung and dorsal root ganglion (DRG) (Supplementary Fig. 19a-c).

### Validation of ALICE’s predictive capabilities and performance of ALICE-generated sequences

To evaluate ALICE’s predictive power across a broader spectrum of predicted performance levels, we conducted systematic experiments that included the top 100 predicted sequences from ALICE, 76 sequences from ***Ref***\*, and 53 sequences from ***Ref***^+^. Using rigorously validated protocols,^64^ we produced viral library and performed in vitro pull-down assays to quantify binding affinities for Ly6a and Ly6c1 receptors (see Supplementary Methods for detail). We further administered the viral library into mice to assess blood-brain barrier (BBB) penetration and CNS transduction efficiency, followed by RNA recovery and sequencing to map functional performance (see Supplementary Methods for details).

The results demonstrated a reasonable correlation between ALICE’s predictions and both in vitro (including production fitness and Ly6a/Ly6c1 log2 enrichment, Pearson r = 0.74; Supplementary Fig. 20a) and in vivo (brain log2 enrichment, Pearson r = 0.64; Supplementary Fig. 20b) experimental outcomes. Adjusting for production fitness had minimal impact on predicting in vivo outcomes (Supplementary Fig. 20c), as receptor engagement, specifically Ly6a and Ly6c1 log2 enrichment, accounted for the dominant effect. Notably, ALICE-generated variants exhibited dual affinity for both Ly6c1 and Ly6a receptors, a critical design objective to enable cross-strain compatibility, whereas existing controls showed preferential binding to only one receptor (Supplementary Fig20d). This dual targeting aligns with our observation that AAV.ALICE-N2 and N6 achieved robust BBB penetration and CNS transduction across all tested strains (BALB/cJ, C57BL/6J, and FVB/NJ, Fig. 3g).

Combining in vitro and in vivo results, we found that the top 30 ALICE-designed capsid variants demonstrated superior average performance in production fitness and Ly6c1/Ly6a binding capability compared to the ***Ref***\*samples and significantly outperformed ***Ref***^+^samples (Supplementary Fig20e-g). In CNS transduction, the top 30 variants performed comparably to those in ***Ref***\*(Supplementary Fig. 20h). Notably, their distribution in production fitness and CNS transduction closely matched that of the positive controls, which were independently validated in vivo in previous studies (Supplementary Fig. 20h; see Source Data for details). Among these top 30 variants, 37% exceeded the mean values for both production fitness and CNS transduction (Supplementary Fig. 20h). Notably, AAV.ALICE-N2 and N9 matched reported variants (AAV.CPP.21 and M-CREATE-generated PHP variants B, B2-B8, C1-C3, V1-V2) in brain enrichment while demonstrating superior production fitness (Supplementary Fig. 20h and Table 2). Both AAV.ALICE-N2 and N9 achieved near-maximal brain transduction efficiency (Fig. 3f, Supplementary Fig. 15e), suggesting these variants, along with others, represent a biological plateau for transvascular CNS targeting.

#### AAV.ALICE-N2 and N6 mediate their functions by targeting the Ly6a/Ly6c1 receptors

To further investigate the mechanism of action of AAV.ALICE-N2 and AAV.ALICE-N6, we assessed whether their CNS transduction is mediated through binding to the designed targets Ly6a and Ly6c1. Lentiviruses were used to stably express Ly6a, Ly6c1, Car4 and Slco1c1 on HEK293T cells (Car4 and Slco1c1 as controls; Supplementary Methods). Following an established protocol,^65^ we first evaluated AAV9 and PHP.eB infectivity across doses in HEK293T and Ly6a-transfected HEK293T cells. A dose of 1×10⁷ v.g. per well (low MOI) was selected because neither vector could infect unmodified HEK293T cells at this level, as these cells lack sufficient Ly6a/Ly6c1 for PHP.eB entry and lack endogenous AAV9 receptor expression. As expected, Ly6a expression selectively enhanced PHP.eB—but not AAV9—infectivity (Supplementary Fig. 21a, b). We then tested AAV.ALICE-N2 and AAV.ALICE-N6 against all four receptor proteins in triplicate at the same dose (1×10⁷ v.g. per well), using PHP.eB with Ly6a as a positive control and untransduced HEK293T cells as a blank control (Extended Data Fig. 3a-c).

We discovered that AAV.ALICE-N2 and AAV.ALICE-N6 displayed notable strong transduction in Ly6a- and Ly6c1-overexpressing cells (Extended Data Fig. 3b, c). In contrast, AAV9 demonstrated no significant transduction in cells overexpressing the target proteins. PHP.eB, employed as a positive control, successfully transduced Ly6a-overexpressing cells but failed to transduce Ly6c1-overexpressing cells (Extended Data Fig. 3b, c), which is consistent with a previous report.^26^ Furthermore, the lack of significant infection with AAV.ALICE-N2 and AAV.ALICE-N6 to cells overexpressing Car4 or Slco1c1 target proteins suggests that the sequence evolutionary path of ALICE is not only robust but also well adapted to the Ly6a and Ly6c1 target landscape, implying a controlled evolutionary process (Extended Data Fig. 3b, c).

#### ALICE’s versatility in engineering capsid sequences with novel motifs for translational hTfR1 targeting

Since Ly6a and Ly6c1 are not conserved in non-human primates (NHPs) or humans,^37^ capsids engineered using these targets lack translational potential for clinical applications. To design clinically relevant capsids, we extended ALICE to engineer AAV capsids targeting the human transferrin receptor (TFRC/TfR1), a well-validated receptor for transcytosis, with established safety and efficacy in both approved therapeutics (e.g., pabinafusp alfa/Lzcargo) and clinical-stage candidates (e.g., trontinemab, NCT07169578, NCT07170150). To do so, we generated a library of AAV variants containing insertion-modified capsid sequences, assessing their production fitness (Supplementary Fig. 22a) and binding capability to hTfR1 (Supplementary Fig. 22b), using established methods (see Supplementary Methods). Capsid sequences with consistently high production fitness and binding capability were selected for semantic tuning and ranking in in the dataset of ***D***_𝒔𝒆𝒒+𝒙_(n = 1,388, see Supplementary Methods). For the FE module, we prioritized sequences based on two criteria: High viability and strong hTfR1 binding capability. Sequences meeting these criteria were designated as ***Ref***\*(n = 27; Supplementary Table 12), while those with weak hTfR1 binding and low viability were classified as ***Ref***^+^ (n = 48; Supplementary Table 12; see Supplementary Methods).

Because ALICE tends to optimize known functional motifs (e.g., ’GYS’) from the training data, as observed in our study targeting Ly6a/Ly6c1, rather than exploring entirely novel ones, we sought to enhance its capability for discovering new sequence motifs beyond wet-lab-derived ones. To achieve this, we first modified the Kullback-Leibler (KL) divergence term in the FE module’s objective function (see Supplementary Methods). The objective function included a normalized KL divergence between generated sequences and a positive reference distribution (see Supplementary Methods). At a low novelty weight (0.2; Supplementary Fig. 23a, b), the sequences retained high fitness and hTfR1 binding capability but exhibiting minimal motif divergence from the reference, as the high-frequency AI-generated motifs (e.g., ’YHR’, ’HRL’, and ’YSR’) are conservative ones present in the ***Ref***\* dataset (Supplementary Fig. 23b). Increasing the weight (1.1) enhanced sequence exploration (Supplementary Fig. 24d) but disrupted dual-functionality integration and significantly reduced production fitness (Supplementary Fig. 23c). A further increased weight (2.0) exacerbated these effects, further diminishing fitness (Supplementary Fig. 23e). These results reveal a fundamental trade-off: excessive novelty regularization impedes multi-functional integration.

Since direct objective function modification proved insufficient, we hypothesized that advanced machine learning strategy is required to reconcile novel motif exploration with multi-functional integration. Inspired by curriculum learning^31^, we developed FE-Explore (FE-X), a two-phase evolutionary framework. The first phase (function integration) employs contrastive learning principles with positive and negative references to establish baseline functionality. The second phase (exploration) introduces an evaluation model to constrain divergence, ensuring that novel motif exploration remains balanced with multi-feature integration. To guide evolutionary trajectories, we compared two reward strategies: minimum reward and sum reward (see Supplementary Method for details). While both strategies preserved multifunctionality, the sum-reward strategy achieved comparable fitness distributions after 20 evolutionary rounds, while significantly outperforming in hTfR1 binding capability (Supplementary Fig. 25). Consequently, we adopted a composite objective function using the sum-reward strategy combining four components: (1) a quality term, (2) an exploration term, (3) a novelty term, and (4) a penalty discouraging convergence toward reference sequences (see Supplementary Methods). Under sum-reward guidance, FE-X generated sequences that diverged from both positive and negative references during the exploration stage (Fig. 4b), following multiple distinct evolutionary trajectories (Fig. 4c). These trajectories gave rise to diversified capsid sequences (Supplementary Fig. 25a, b) containing motifs rarely observed in ***Ref***\*samples (Supplementary Fig. 25c). Functional trajectory analysis further revealed an alternating pattern between novel motif exploration and multi-function fusion (Fig. 4d), underscoring the dynamic trade-offs imposed by reward-driven selection. At both local (Supplementary Fig. 25d) and global (Supplementary Fig. 25e) semantic levels, the evolutionary process progressively shifted away from the reference data (Supplementary Fig. 25d, e), reflecting a continuous balance between diversity expansion and multi-functional integration.

**Fig. 4.**
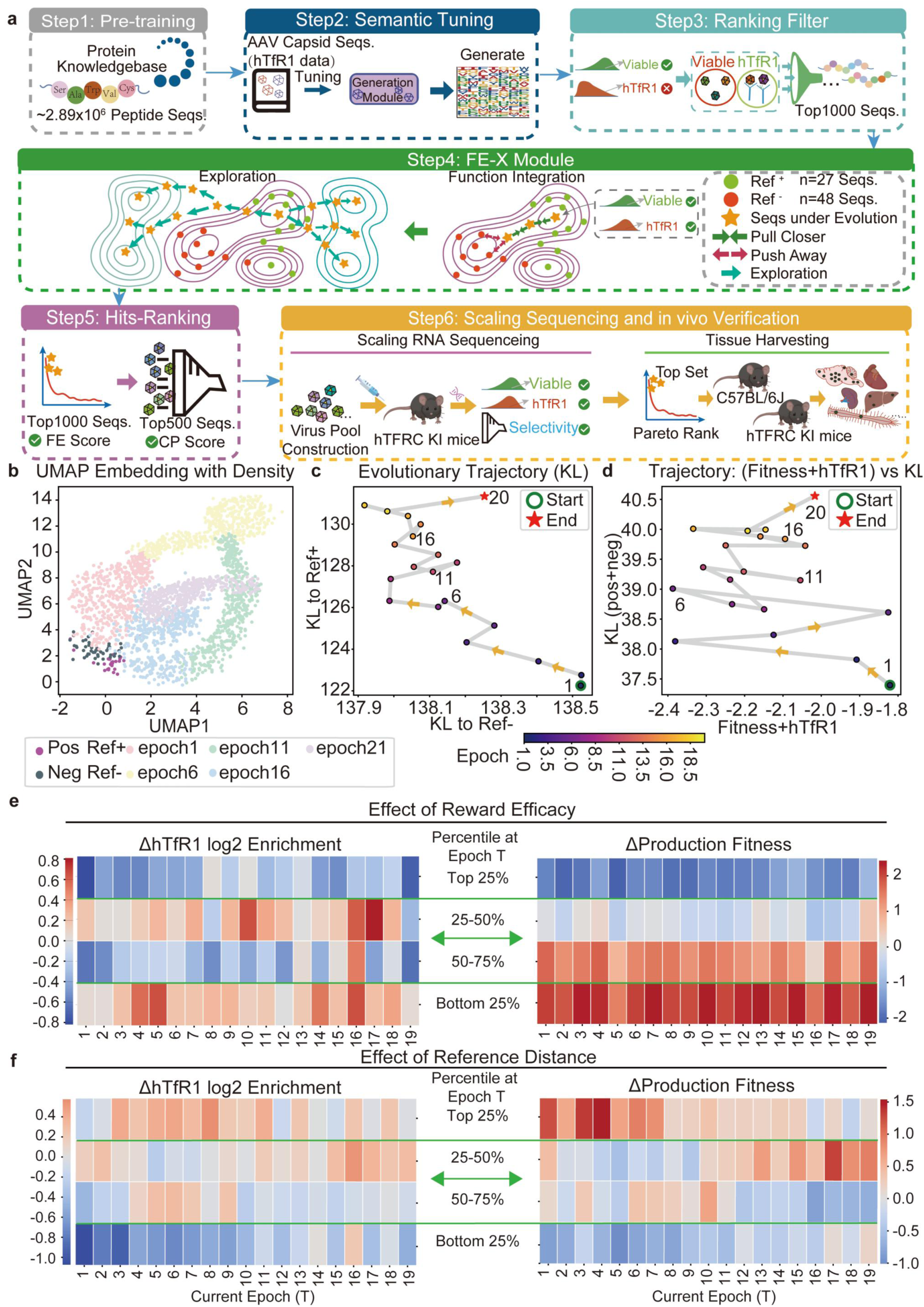
ALICE-X designs novel capsid sequences for translational hTfR1 targeting. **a,** Workflow diagram of the ALICE-X platform for expanded hTfR1-targeted sequence space exploration. **b,** UMAP visualization of evolutionary trajectories during exploration. ALICE-X generates diverse evolutionary distributions from the initial positive and negative reference samples (***Ref***^!^ and ***Ref***^−^). Unlike ALICE, which exhibits trajectories biased toward ***Ref***^!^, ALICE-X trajectories substantially diverge from both reference sets. **c,** Evolutionary trajectory of ALICE-X, quantified by its dynamic distance to ***Ref***^!^ and ***Ref***^−^. **d,** Joint trajectory of ALICE-X, integrating both reference distance and functional performance metrics during exploration. **e,** Effect of reward efficacy on the median change in hTfR1 log2 enrichment (left) and production fitness (right) in the subsequent epoch during evolution. **f,** Effect of reference distance on the median change in hTfR1 log2 enrichment (left) and production fitness (right) in the subsequent epoch during evolution.

To evaluate the individual contributions of each objective function term to functional outcomes during sequence exploration, we quantified evolutionary gain/loss by assessing hTfR1 enrichment and production fitness across training epochs using our evaluation model (see Supplementary Methods for details). Our analysis revealed that relatively low reward levels (bottom 25%) consistently yielded the most significant improvements in both hTfR1 binding capability and production fitness (Fig. 4e). In contrast, moderate rewards (25-50% or 50-75%) selectively enhanced either production fitness (Fig. 4e right) or binding capability (Fig. 4e, left), but not both simultaneously. But excessively strong rewards paradoxically diminished both production fitness and hTfR1 enrichment (Fig. 4e). Notably, sequences designed with the highest diversity metrics (top 25%), as defined by increased Hamming distance from reference sets (novelty term, see Supplementary Methods), exhibited significant improvements in both production fitness and hTfR1 binding capability (Fig. 4f). Furthermore, the diversity regularization (exploration term, see Supplementary Methods) in the FE-X module, which enforces dissimilarity across consecutive generations, also proved critical for functional optimization (Supplementary Fig. 26f, see Supplementary Methods for details). Sequences designed with the top 25% diversity metrics, exhibiting the greatest inter-generational distances, demonstrated optimal improvements in both hTfR1 enrichment and production fitness (Supplementary Fig. 25f).

#### Ablation study for functional validation of modules in ALICE-X

To systematically evaluate the functional contributions of each ALICE-X module in balancing production fitness with hTfR1 binding capability while enabling exploration of novel motifs, we compared five distinct design strategies: (1) ALICE-X, (2) ALICE, (3) Semantic Tuning with Hits-Ranking Module (ST Hits-Rank), (4) Semantic Tuning only (ST), and (5) Random Mutagenesis (RM) of ***Ref***\*capsid sequences. For each approach, 500 sequences were generated, assembled into an AAV variant library, and quantitatively assessed for both production fitness and hTfR1 binding capability using pulldown assays (see Supplementary Methods for details).

Comparative analyses revealed that ALICE-X uniquely produced sequences harboring novel motifs (e.g., ’YTK’ and ’TKS’) that were rarely observed in ***Ref***\*or the training dataset (Fig. 5a, b), while achieving an optimal balance between experimentally measured production fitness and hTfR1 binding capability (Fig. 5e). In contrast, ALICE generated preserved ***Ref***\*-derived motifs, like ’HRL’ and ’LHR’, yet retained reasonable experimentally measured production fitness and hTfR1 binding capacity (Supplementary Fig. 26a, b, and Fig. 5e). Ablation of the FE module severely restricted search capacity: both Semantic Tuning variants (with or without the Ranking Module) generated sequences clustered near ***Ref***\* and failed to reliably integrate production fitness with hTfR1 binding capability (Supplementary Fig. 26c-f, and Fig. 5e). Random mutagenesis of ***Ref***\*produced sequences with minimal divergence from the parental template, accompanied by markedly reduced dual functionality (Supplementary Fig. 26g, h, and Fig. 5e), underscoring the inherent limitations of stochastic mutational strategies.

**Fig. 5.**
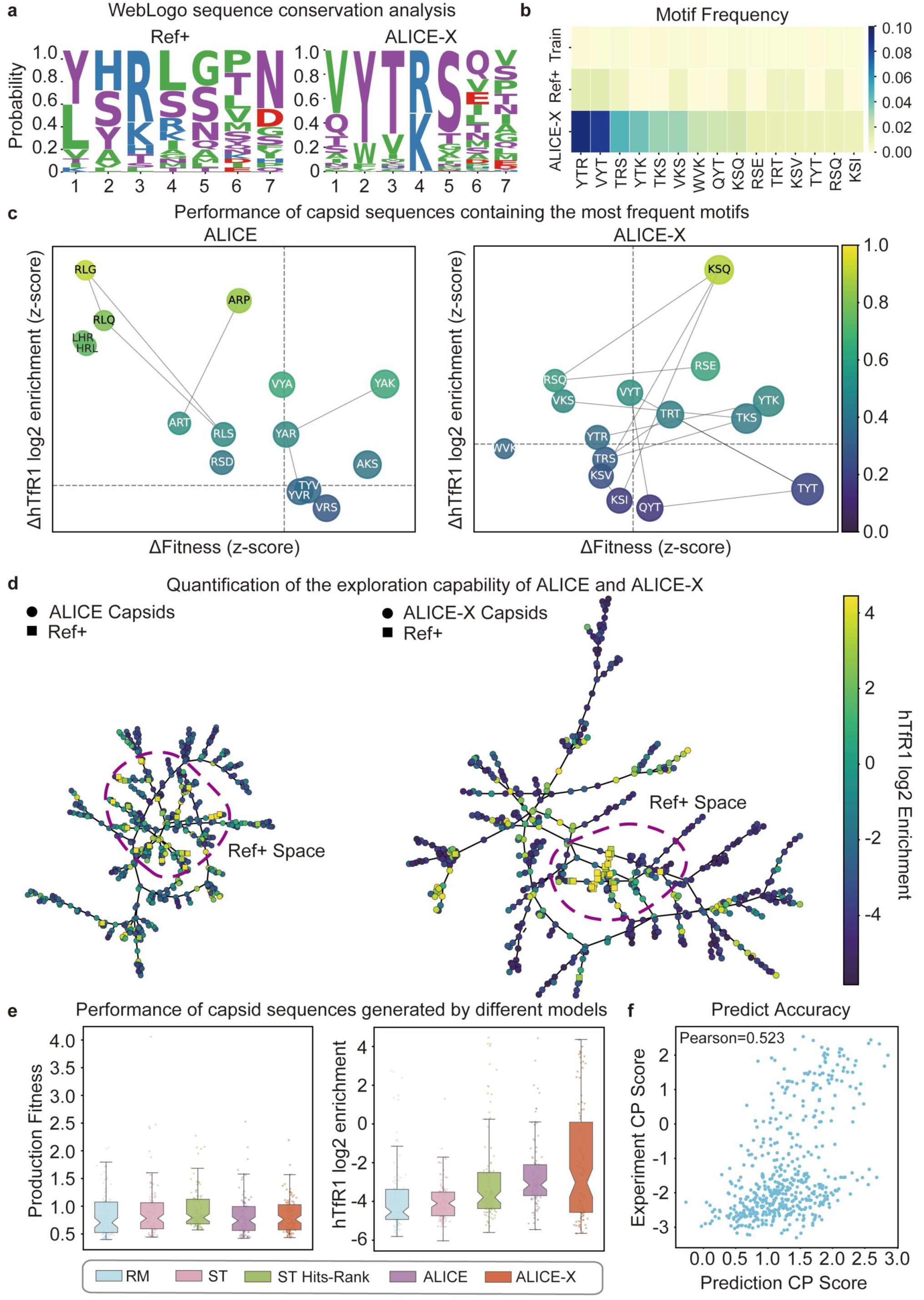
Generalization and novel motif exploration capacity of ALICE-X. **a,** Sequence WebLogo comparison of positive reference capsids (***Ref***^!^) and top 500 ALICE-X-generated variants. The frequency-based ’unit height’ normalization was applied, where the total stack height is fixed to 1 for all positions to display only relative amino acid frequencies. This approach was chosen to emphasize positional preferences and residue composition rather than global conservation differences across sites. Consequently, while all positions show stacks of equal height, their underlying Shannon entropy values remain distinct. **b,** Heatmap comparing the frequency of the top 15 ALICE-X motifs with their occurrence in ***Ref***^!^and the training dataset. **c,** Motif-level contributions to experimentally measured hTfR1 enrichment and production fitness. Scatter plots compare the effects of top 15 motifs from ALICE (left) and ALICE-X (right) models, showing normalized z-score differences in ΔFitness (x-axis) versus ΔhTfR1 log2 enrichment (y-axis). Node color indicates relative hTfR1 enrichment (normalized), while edges represent sequence similarity. **d,** Minimum spanning tree (MST) representation of sequence space for ALICE (left) and ALICE-X (right) capsids, together with ***Ref***^!^sequences, embedded in Hamming space. Node color indicates hTfR1 log2 enrichment. Purple dashed circles mark the ***Ref***^!^ sequence space. **e,** Boxplots comparing experimentally measured production fitness (left) and hTfR1 log2 enrichment (right) across capsid libraries generated by five models: random mutation (RM), semantic tuning only (ST), semantic tuning with hits-ranking module (ST hits-rank), ALICE, and ALICE-X. **f,** Correlation of predicted versus experimentally measured hTfR1 enrichment scores for ALICE-X variants (Pearson’s r = 0.523). CP scores (defined in Supplementary Methods, section 2.3) are shown on the x-axis, with experimental measurements on the y-axis.

Deeper analysis of capsid sequences containing these motifs, evaluated for experimentally measured production fitness and hTfR1 binding capacity (see Supplementary Methods), showed that ALICE-X preferentially enriched both a truly novel motif (’YTK’, absent from both ***Ref***\* and the training dataset) and several extremely low-frequency motifs (’KSQ’ and ’RSE’). Importantly, sequences containing these motifs consistently outperformed the global mean (dashed lines) across both metrics and surpassed all five alternative generation strategies (Fig. 5a-c, Extended Data Fig. 4). In contrast, ALICE generated only a limited number of dual-functional motifs, most of which (e.g., ’YAK’) were conservative motifs present in the ***Ref***\* dataset (Fig. 5c Left, Supplementary Fig. 26a, b). Architectures lacking FE or FE-X modules (ST hits-Rank) identified motifs (e.g., ’KSQ’, ’YTR’) that were similarly conservative in ***Ref***\*(Extended Data Fig. 4a and Supplementary Fig. 26c, d). Strategies using only Semantic Tuning (ST) (Extended Data Fig. 4b and Supplementary Fig. 26e, f) predominantly produced motifs in low-affinity regions, while random mutation (RM) of ***Ref***\*yielded sequences (e.g., ’KSQ’) that were either conservative or resided in low-fitness, low-capacity regions (Extended Data Fig. 4c and Supplementary Fig. 26g, h). These results align with our ablation studies and collectively demonstrate the critical importance of ALICE-X’s integrated design framework.

To analyze evolutionary relationships among designed capsid sequences, we constructed minimum spanning trees (MSTs) based on pairwise Hamming distances between designed capsid sequences (see details in the Supplementary Methods). Network analysis showed that ALICE-derived variants clustered near ***Ref***\*sequences (Fig. 5d, left), while ALICE-X successfully accessed novel sequence space with significantly greater motif diversity (Fig. 5d, right). Notably, ALICE-X achieved this expanded exploration while maintaining reasonable predictive accuracy (Pearson r = 0.523; Fig. 5f).

#### In vivo functional verification of variants engineered by ALICE-X

Given the well-documented nonlinear relationship between TfR1 binding affinity and blood-brain barrier (BBB) penetration^66^, we produced an AAV library incorporating ALICE-X generated capsid sequences and administered it via retro-orbital intravenous injection into humanized TFRC knock-in (*hTFRC* KI) and wild-type mice (see Supplementary Methods). Quantitative RNA-seq analysis of recovered capsid sequences in the brain tissues was performed to assess transduction efficiency.

Using Pareto front-based ranking, which integrated production fitness, enrichment gain in *hTFRC* KI mice relative to wild-type controls, and sequence logo analysis, we selected three candidates, AAV.ALICE-H1 (TYTKSEI), H2 (VYTKSDT), and H3 (QYTKSIT), from the top set (Fig. 6a, Supplementary Table 13), containing the conserved ‘YTK’ motif for individual validation (Fig. 6b). Each capsid was produced separately for in vivo characterization. AAV.ALICE-H3 demonstrated superior performance, exhibiting both ∼1.4× higher viability than AAV9 (Supplementary Table 14) and ∼251-fold greater CNS expression in *hTFRC* KI mice versus wild-type controls (Fig. 6c, g). In contrast, AAV.ALICE-H1 and H2 showed normal viability (Supplementary Table 14) and limited *hTFRC* KI mice selectivity (1.6- and 16.7-fold enrichment in KI mice, respectively), suggesting potential engagement of additional receptors for BBB penetration. To contextualize these findings, we also compared ALICE-X-designed variants against the current benchmark capsid BI-hTFR1 (YSRIGPN)^24^ within the same Pareto-front analysis. BI-hTFR1 received a Pareto rank of 9 (Fig. 6a and Supplementary Table 13), whereas the majority of ALICE-X sequences surpassed this benchmark, further supporting the ability of ALICE-X to identify multifunctional motifs with improved overall performance relative to established controls.

**Fig. 6.**
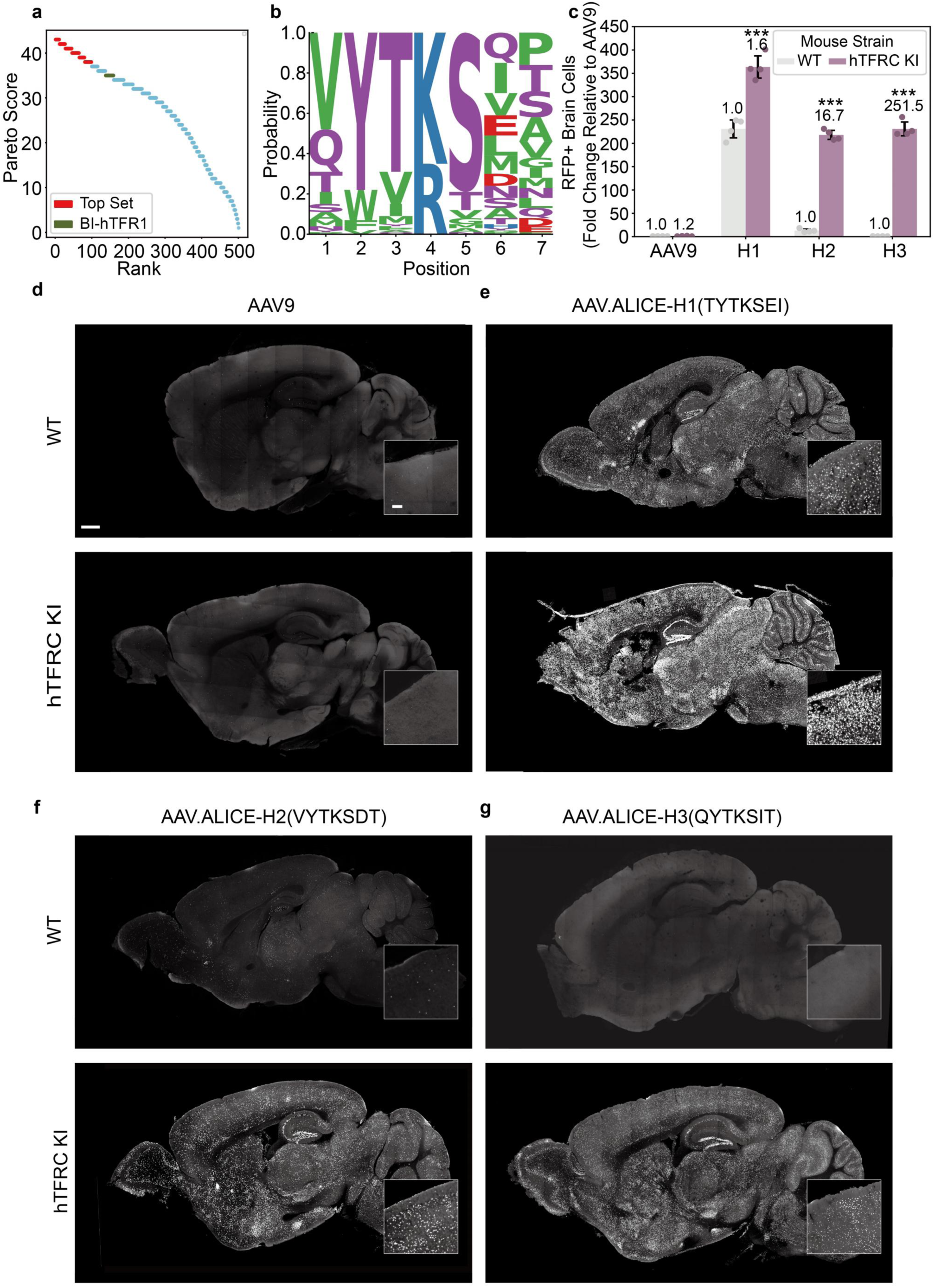
In vivo validation of capsids designed by ALICE-X. **a,** Top 3 candidate sequences were selected from the Pareto front-top set, ranked by combined brain enrichment and capsid production fitness (see Supplementary Methods), for individual in vivo validation **b,** Sequence WebLogo depicting the amino acid distribution at each position, generated from the 99 sequences in the top-6 Pareto set. **c,** Quantitative analysis of the percentage of total RFP-positive brain cells and the fold change relative to AAV9. Data are shown as mean ± s.e.m.; n = 4 mice per group. Statistical significance was determined by one-way ANOVA followed by Tukey’s post hoc test (*P < 0.05, **P < 0.01, ***P < 0.001 vs. AAV9). **d-g,** Whole-brain distribution of AAV9 (d), AAV.ALICE-H1 (e), AAV.ALICE-H2 (f), and AAV.ALICE-H3 (g) in wild-type (C57BL/6J; top) and hTfR1-KI (bottom) mice at 21 days post-IV injection (5×10¹¹ vg/mouse). Scale bars: 1 mm (overview) and 200 μm (insets).

To assess binding specificity, we performed pulldown assays of the AAV library against both human (hTfR1) and macaque (macTfR1) TfR1 (see Supplementary Methods for details). AAV.ALICE-H1-H3 demonstrated strong preferential binding to hTfR1, with minimal interaction with macTfR1, a profile similar to the positive control BI-hTFR1 (Supplementary Fig. 28a). When comparing the top ALICE-X-designed sequences with the ***Ref***\*, we found that their mean production fitness was comparable to that of ***Ref***\*, while their mean hTfR1 log2 enrichment was significantly higher. Together with the superior in vitro and in vivo performance described above (Fig. 6a), these results demonstrate the effectiveness of ALICE-X in exploring a broader and more functional sequence space.

We selected AAV.ALICE-H3 for further characterization based on its superior selectivity for *hTFRC* KI mice. Quantitative analysis demonstrated AAV.ALICE-H3-EF1α-H2B-RFP’s broad neural transduction, with 25-51% of cells transduced across various brain regions and spinal cord (Supplementary Fig. 29). Immunohistochemical profiling revealed predominant neuronal tropism (24-59% NeuN+ cells), with lower but regionally variable transduction of glia (4-11% SOX9+) and oligodendrocytes (1-11% Olig2+) (Supplementary Fig. 29). This cell-type preference may reflect the used EF1α promoter’s limited activity in non-neuronal lineages^67^. Importantly, peripheral transduction profiles mirrored AAV9 in both *hTFRC* KI and wild-type mice, with only marginally increased lung transduction (Supplementary Fig. 30).

To probe the binding mechanism, we pre-incubated hTfR1-conjugated Dynabeads with either: OKT9 antibody (targeting the hTfR1 apical domain), or holo-transferrin (sub-nanomolar affinity hTfR1 ligand), prior to AAV enrichment (Extended Data Fig. 5a). Both AAV.ALICE-H1-H3 and BI-hTFR1 showed significantly reduced hTfR1 binding after OKT9 treatment, but no inhibition with holo-Tf (Extended Data Fig. 5b-e). These results indicate that AAV.ALICE-H1-H3, like BI-hTFR1, specifically engages the apical domain of hTfR1 without occluding transferrin-binding sites^24^.

## Discussion

Traditional AAV capsid engineering relies heavily on in vivo selection, an approach that is inherently stochastic, labor-intensive, and costly. Although recent AI-based strategies have improved capsid diversity and fitness^64,68^, they still depend on extensive empirical screening. In contrast, ALICE establishes a fundamentally new, fully in silico framework that directly maps sequence to multifactorial function, integrating production fitness, receptor binding, and CNS tropism—three properties essential for therapeutic vector design.

A major advantage of ALICE is its ability to enable *in silico* knowledge-driven, multifunctional capsid design. Engineering sequences that simultaneously optimize multiple biophysical constraints is an NP-hard problem^69^, as no polynomial-time mapping exists between sequence and composite function. Our initial trials with naïve language-model pipelines (pretraining + semantic tuning; Fig. 1b) revealed this difficulty: while these models increased target-binding potential, they substantially impaired production fitness (Extended Data Fig. 1l), likely due to limited training data that bias models toward superficial sequence regularities rather than functionally critical motifs (Supplementary Fig. 13d). To resolve this, we developed an interpretable function-guided evolution (FE) strategy that explicitly balances beneficial sequence features while suppressing deleterious ones (Fig. 2c), enabling simultaneous optimization of otherwise antagonistic traits. To evaluate individual module contributions in ALICE, we employed predictive metrics as proxies in our ablation study (Fig. 2d). Although the metrics predicted by the chosen GradientBoostingRegressor model showed strong correlation with experimentally validated functional outcomes (Pearson and Spearman r = 0.7-0.9; Supplementary Figs. 2-4), these computational predictions inherently possess limitations. Through wet-lab experiments (Fig. 5e), we further validated the FE module’s contribution in ALICE-X, with results consistent with those obtained using predictive metrics in ALICE.

A second key innovation of ALICE-X is its capacity to generate genuinely novel motifs with integrated functionality, a longstanding challenge in machine learning-guided protein engineering^70^. Unlike methods constrained by limited training data or reward specifications (e.g., reinforcement-learning approaches such as AlphaZero^71^ or RLHF^72^), ALICE-X employs a curriculum learning framework^31^ that decomposes complex optimization tasks into progressive stages. This enables FE-X to explore beyond the manifold of the training distribution (Fig. 4b-f) while preserving multifunctional coherence, yielding motifs inaccessible to conventional experimental screens (Fig. 5). This framework establishes a generalizable paradigm for reconciling novelty with biological viability.

More importantly, ALICE-X is inherently designed with translational constraints in mind. While directed evolution or AI trained on animal datasets may overlook human-specific receptor biology, ALICE-X integrates human biophysical rules, such as hTfR1 epitope geometry, to avoid preclinical dead ends. This capability enabled the design of AAV.ALICE-H1-H3 carrying the novel ’YTK’ motif, which exhibited exceptional hTfR1 specificity and a well-defined mechanism of action (Fig. 6; Extended Data Fig. 5). AAV.ALICE-H3 exhibited a ∼251-fold reduction in off-target transduction in WT mice compared to hTFRC KI mice (Fig. 6c, g), further validating its human-specific precision. These results collectively demonstrate the translational potential of ALICE-X-generated capsid.

While exhaustive FE scoring of the entire 20⁷ (1.28 billion) 7-mer sequence space is theoretically feasible on modern hardware, this brute-force approach fails to address two key challenges in functional sequence design: uncertainty in predictive scoring and experimental throughput limitations. No predictive model, alignment-based or neural, achieves perfect calibration across such a vast combinatorial space, and large uncertainty regions persist^73,74^. By leveraging ***Ref***\* and ***Ref***^+^functional priors, FE-guided evolution efficiently enriches sequences in biologically meaningful regions while avoiding unproductive areas of sequence space. As shown in Fig. 2d and Fig. 5e, removing the FE module leads to capsid designs with poor production fitness or diminished target-binding affinity. By contrast, incorporating the FE module markedly enhances search efficiency compare with brute-force enumeration, which often over-prioritizes high-scoring yet biophysically implausible sequences and could divert substantial resources toward false-positive candidates during downstream in vivo validation. Notably, in vivo screening results lose reproducibility and reliability when AAV libraries exceed 10⁵ variants in mice^28^, a limitation that becomes even more pronounced in large animal models such as non-human primates (NHPs). Additionally, conventional generative models largely replicate the empirical distribution of training data (Extended Data Fig. 1; Supplementary Fig. 6) and rarely escape into sparsely populated functional basins. FE-X enables directed navigation toward multi-objective optima while retaining this capability for engineering longer capsid sequences, including insertions or replacements, where combinatorial expansion makes brute-force approaches computationally intractable. Therefore, FE-guided evolutionary modeling is not only computationally efficient but also conceptually essential for the discovery of multifunctional capsids.

Finally, ALICE is not intended to replace traditional vector engineering pipelines but to augment and accelerate them. By embedding mechanistic knowledge of AAV biology and capsid-receptor interactions into a scalable in silico framework, ALICE enables rapid, low-cost discovery of therapeutically optimized variants. Its modular structure also makes it readily adaptable to designing capsids for other tissues or cell types (e.g., retina, muscle, T cells) as additional receptor-capsid datasets become available. Given that most datasets in modern biomedicine are inherently sparse and multi-functional, the ability to leverage search algorithms that learn complex patterns from limited samples and efficiently explore broader design spaces represents one of the valuable pathways for accelerating AI-driven translation into practical therapeutic applications. By merging computational efficiency with mechanistic interpretability, ALICE presents a powerful blueprint for the next generation of mechanism-guided, multifunctional AAV vector design.

### Limitations of the study

While ALICE has demonstrated capabilities in designing multifunctional AAV vectors, several opportunities for improvement remain. First, ALICE currently lacks the capability to identify novel receptors, a limitation stemming from the absence of high-throughput capsid-receptor interaction data across diverse host receptors, which impedes systematic discovery. Although our prior studies confirmed ALICE-X’s ability to explore novel sequences for conventionally identified new receptors, the present work centers on establishing a generalizable sequence-to-function computational framework. Second, ALICE does not incorporate structural information into its coevolutionary strategy; integrating such data, for example, through AlphaFold^75^, could enhance its predictive performance. Additionally, this study was limited to 7-mer peptide insertions between VP1 residues 588-589 of the AAV9 capsid, restricting exploration to a sequence space of 20^7^ variants. Future work incorporating data from broader capsid modifications could significantly expand ALICE-X’s screening capacity. Another limitation is that we did not investigate other receptors that may influence the *hTFRC* KI mice selectivity of AAV.ALICE-H1 and H2, which warrants further study. Furthermore, while liver de-targeting remains an important translational goal, we will explore this optimization by introducing additional capsid mutations in future work, as it lies beyond the scope of the present study. Notably, although it is technically possible to engineer AAV capsids capable of engaging both human (hTfR1) and macaque (macTfR1) transferrin receptors, such dual targeting is not a prerequisite for translational development. The FDA has recognized the utility of humanized mouse models for preclinical biodistribution and safety assessments, and many human-specific biologics have advanced to clinical approval without requiring extensive large-animal evaluations^76^.

## Methods

### Plasmids

Plasmid libraries for functional AAV capsid selection were generated based on a previously described strategy with modifications.^11^ A pUC-Cap9XA vector, derived from pAAV2/9 by introducing the K449R mutation, was used as the template for all PCR amplifications. The vector was linearized with XbaI (NEB, R0145V) and AgeI (NEB, R3552L).

The pAAV-588-7NNK-lib, containing a seven-amino acid insertion between residues Q588 and A589, was created by PCR using the forward primer Assembly-XbaI-F (5′-ACTCATCGACCAATACTTGTACTATCTCTCTAGAAC-3′) and the reverse primer Assembly-AAV9-588i7NNK (5′-GTATTCCTTGGTTTTGAACCCAACCGGTCTGCGCCTGTGCMNNMNNMNNMNNMNNMNNMNNT TGGGCACTCTGGTGGTTTGTG-3′).

For the oligonucleotide libraries pAAV-588-OligoPool-lib1, −lib2, and −lib200, an oligo pool was used as the reverse primer with Assembly-XbaI-F as the forward primer for an initial 5 amplification cycles, followed by a second round of 25 cycles with Assembly-AgeI-R (5′-GTATTCCTTGGTTTTGAACCCAACCG-3′).

Purified PCR products were assembled into an XbaI/AgeI-digested pAAV-P41-Rep78-Cap9-588iSTOP-WPRE-hGH backbone (containing the AAV5 P41 promoter, Rep78, AAV9 Cap, WPRE, and hGH elements) using Gibson Assembly Master Mix (NEB, E2611). PCR and digested products were purified with the Gel DNA Recovery Kit (Genstone, TD407).

Additional AAV capsid plasmids were derived from pAAV2/9n (Addgene #112865). DNA fragments encoding peptide inserts were synthesized by GenScript and introduced between VP1 residues 588 and 589 using CloneEZ Seamless Cloning technology (GenScript).

For viral packaging, pAAV-EF1α-H2B-mCherry (containing the EF1α promoter, H2B, and mCherry cassette) and a pHelper plasmid (Agilent, 240071-12) were used for single-vector production. For library-scale packaging, a RepAAP plasmid was constructed by introducing a stop codon into the Cap sequence to block VP expression while maintaining Rep and AAP sequences required for packaging.

Coding sequences for Ly6a (NM_010738.3), Ly6c1 (NM_010741.4), Car4 (NM_000717.5), and Slco1c1 (NM_001145944.2) were synthesized and cloned into the pHAGE-P2A-eGFP, which is modified from pHAGE-EF1aL-eGFP-W (Addgene #126686), for lentivirus production.

The coding sequences for Ly6a(NM_001271416.1), Ly6c(NM_010741.3), human transferrin receptor 1 (hTfR1; NM_003234.4), or rhesus macaque transferrin receptor 1 (macTfR1; NM_001257303.1) were cloned into the pcDNA3.4 transfer plasmid for protein production.

### AAV production and purification

Recombinant AAVs were packaged via a standard three-plasmid co-transfection protocol. The AAV9, AAV.ALICE-N1 ∼ AAV.ALICE-N8 and AAV.ALICE-H1 ∼ AAV.ALICE-H3 (including pRC plasmids: pAAV2/9n, pAAV2/9n-N1∼N8, and pAAV2/9n-H1∼H3, a pAAV plasmid carrying a transgene expression cassette: pAAV-EF1α-H2B-RFP, and a pHelper plasmid (240071-12, Agilent Technologies) donor vectors were sent to the viral core for viral packaging and production. The purified AAVs were stored in a freezer at −80°C in small aliquots. Each virus was packaged three times or more. The viral core reported the average titers of AAV9 and AAV.ALICE-N1∼ AAV.ALICE-N8 values were 2.41 × 10^13^, 9.59 × 10^10^, 5.55 × 10^13^, 6.00 × 10^12^, 6.02 × 10^12^, 9.59 × 10^10^, 4.95 × 10^13^, 8.30 × 10^12^ and 3.86 × 10^13^ vector genomes (vg) per ml, respectively. AAV.ALICE-H1∼ AAV.ALICE-H3 values were 4.05 × 10^12^, 5.58 × 10^12^, 3.33 × 10^13^ vector genomes (vg) per ml, respectively.

The viral packaging process was performed using the ALICE-designed (pAAV-588-OligoPool-lib200), pAAV-588-7NNK-lib, pAAV-588-OligoPool-lib1, or ALICE-X-designed library (pAAV-588-OligoPool-lib2) oligo pool mutant plasmid libraries, and following a previously optimized protocol^77^ to reduce the production of mosaic capsids. The specific implementation steps included: (1) A triple-plasmid co-transfection system was employed, with each 150 mm culture dish containing the following amounts of plasmids: AAV-588-OligoPool mutant plasmid library: 50 ng; RepAAP plasmid: 11.4 μg; Helper plasmid: 22.8 μg; pUC19 plasmid: 5.7 μg. Polyethylenimine (PEI) was used as the transfection reagent at a dosage of 157 μl (10μg/μl) per 150 mm culture dish; (2) Transfected HEK293T cells were cultured at 37°C under 5% CO_2_ conditions. Six to twelve hours post-transfection, the medium was replaced with fresh DMEM medium to remove residual transfection reagents; (3) A timed collection strategy was implemented to maximize viral yield, with virus-containing medium was first collected at 72 hours post-transfection, and replaced with fresh DMEM medium, then cell pellets and the replaced medium were collected at 120 hours post-transfection; (4) The subsequent purification process utilized a multi-stage strategy: fractional precipitation with polyethylene glycol (PEG-8000) combined with iodixanol-based gradient centrifugation (using OptiSeal Polypropylene Tube #362183 and ultracentrifugation by Beckman VTi 50.1 rotor for 48,000 rpm and 1 hour in 4°C) for viral purification. Finally, real-time quantitative PCR (qPCR) with Primer ITR-F (5’-CGGCCTCAGTGAGCGAGC-3’) and ITR-R (5’-AGGAACCCCTAGTGATGG-3’) was used to determine AAV genome concentration: Viral genomic DNA was first extracted from the virus suspension; specific primers and probes were then designed targeting the sequence of interest to establish a qPCR detection system; quantitative analysis of viral genome copy numbers was performed based on a standard curve, with results standardized and expressed in units of genome copies per milliliter (vg/mL). This standardized workflow enabled precise quantitative assessment of viral production efficiency. The final outcome was a high-purity, high-titer AAV-588-OligoPool capsid mutant library suitable for subsequent in vivo screening and validation in animal models.

### Generation of HEK293T cell lines stably expressing receptor proteins

The lentiviral transfer plasmid containing the target genes was co-transfected with the envelope plasmid (pMD2.G) and the packaging plasmid (psPAX2) into HEK293FT cells. The crude lentiviral supernatant was collected at 24-hour intervals over 48 hours. For transduction, wild-type HEK293T cells were seeded in a 6-well plate at a density of 1 × 10⁵ cells per well and cultured at 37°C until reaching 30-40% confluence. The medium was then replaced with the crude lentivirus supernatant, and the cells were incubated for 12 hours. After replacing the supernatant with fresh medium, the cells were cultured for an additional 24 hours. The cell culture medium was supplemented with 2 µg/mL puromycin to select for positive cells. Cells expressing eGFP, indicating successful lentiviral transduction and stable receptor expression, were considered positive. Flow cytometry was performed once the percentage of eGFP-positive cells reached 10%.

### Flow cytometry

Once the proportion of EGFP-positive cells reached approximately 10%, the culture medium was carefully aspirated. The cells were subsequently treated with 0.25% trypsin, gently detached, and then rinsed and revitalized with culture medium. They were transferred into a 15 mL centrifuge tube and subjected to centrifugation at 800×g for 3 minutes to pellet the cells. After centrifugation, the cells were resuspended in FACS buffer composed of PBS supplemented with 2% FBS and 5 mM EDTA. The cell suspension was then adjusted to a concentration of approximately 1×10^7^ cells/mL and passed through a 40 µm filter to eliminate any cell aggregates. The cell samples were processed on a MoFlo Astrios EQ flow cytometer (Beckman Coulter), and the acquired data were meticulously analyzed via FlowJo V10 software.

### Determination of the infection titers of AAV

Wild-type HEK293T cells were seeded at 20% confluency in 96-well plates and cultured in FluoroBrite DMEM. The medium was supplemented with 0.5% FBS, 1% NEAA, penicillin‒streptomycin (100 U/mL), 1× GlutaMAX, and 15 mM HEPES, and the cells were maintained at 37°C in 5% CO_2_. The cells were then transduced with AAV9 and PHP.eB variants at six different titer gradients ranging from 1×10^10^ to 5×10^7^ viral genomes (v.g.) per well, with each condition tested in triplicate. After 24 hours of incubation, the cells were imaged via a 20× objective on an OLYMPUS IX73 microscope with autofocus for each well. The infection titer of 5×10^7^ v.g. per well, at which neither AAV9 nor PHP.eB could infect the cells, was selected as the selected titer for further experiments. HEK293T-Ly6a cells were similarly plated at 20% confluency in 96-well plates and cultured in the same medium. These cells were transduced with either AAV9 or PHP.eB at four titer gradients ranging from 5×10^7^ v.g. to 1×10^6^ v.g. per well and then cultured for an additional 24 hours. The same imaging process was applied. The infection titer of 1×10^7^ v.g. per well, where PHP.eB was able to infect the cells but AAV9 was not, was chosen for testing the AAVs in the receptor-characterization assay.

### In vitro characterization of both well-established and potential receptors for AAV

Wild-type 293T cells, as well as 293T cell lines generated to highly express various receptors, i.e., Ly6a (NM_010738.3), Ly6c1 (NM_010741.4), Car4 (NM_000717.5) and Slcoc1 (NM_001145944.2), were seeded in 96-well plates at an initial density of approximately 6,000 cells per well. Once the cells reached a confluence of approximately 20%, they were transduced by a technician blinded to the treatment groups, with the test rAAVs at a titer of 1×10^7^ v.g. per well. The cells were cultured at 37°C in 5% CO_2_ for an additional 24 hours. The cells were subsequently imaged by a technician blinded to the treatment groups via an OLYMPUS IX73 microscope with a 20x objective for autofocus on each well in triplicate.

### Synthetic oligo pool library design

The pAAV-588-OligoPool-lib200 library was designed using the ALICE model to prioritize variants with predicted binding affinity for Ly6a and Ly6c1. The top 100 amino acid sequences ranked by model evaluation were selected. To this set, 76 positive reference sequences (𝑅𝑒𝑓⁺) and 54 negative reference sequences (𝑅𝑒𝑓⁻) were added, each represented by two synonymous codon variants to provide biological replicates. In total, the library comprised 230 unique sequences (see Source Data and Supplementary Table2 for the full list of the sequences).

The pAAV-588-OligoPool-lib1 library was constructed by selecting up to 6402 variants (6,387 variants were recovered) from the pAAV-588-7NNK library based on performance in pulldown assays and the positive control BI-hTFR1 (see Source data for the full list of the sequences).0. Variants were prioritized by enrichment against human transferrin receptor 1 (hTfR1), requiring a substantially higher log₂ enrichment relative to the human IgG Fc-only control, while excluding sequences with low hTfR1 enrichment values.

The pAAV-588-OligoPool-lib2 library was generated to benchmark computational design strategies. A total of 2,501 variants (2,500 variants were recovered) were included, comprising 500 sequences each from five sources: ALICE, ALICE-X, semantic tuning only (ST), semantic tuning combined with Hits-Ranking (ST Hits-Rank), random mutagenesis (RM) of top-performing variants, and the positive control BI-hTFR1 (see Source data for the full list of the sequences).

### Pull-down assay for AAV receptors

Pull-down assays were performed to enrich adeno-associated virus (AAV) variants displaying affinity for Ly6a(NM_001271416.1), Ly6c(NM_010741.3), human transferrin receptor 1 (hTfR1; NM_003234.4), or rhesus macaque transferrin receptor 1 (macTfR1; NM_001257303.1), using Fc-tagged recombinant ectodomains, with Fc-only protein as control. Recombinant proteins were generated by Genscript Biotech Corporation, where coding sequences were synthesized and cloned into a pcDNA3.4 vector supplied by the company, transiently expressed in the proprietary HD293F system, and subsequently purified.^78^ All steps were carried out in low protein-binding microcentrifuge tubes to minimize nonspecific adsorption. Washing and incubation buffers consisted of DPBS containing 0.05% Tween-20 (DT) or DT supplemented with 1% BSA (BDT).

For receptor-specific pull-downs, Dynabeads Protein A (Thermo Fisher Scientific, 10001D) were pre-incubated with Fc-fusion proteins for >6 h at 4°C. After washing, beads were incubated with 1 × 10¹⁰ vector genomes of the AAV capsid library in BDT overnight at 4°C. Bead-protein-AAV complexes were washed three times with DT and subsequently treated with proteinase K (TIANGEN, #RT403) to release viral genomes for PCR recovery and next-generation sequencing (NGS). Pulldown assays for each target were performed in triplicate to minimize experimental errors.

To determine whether ALICE-X-generated variants interfere with the physiological function of human transferrin receptor 1 (hTfR1), a competitive binding assay was performed. Dynabeads Protein A were first incubated with hTfR1-Fc for 15 min at 4°C. Bead-receptor complexes were then incubated for 6 h with either the OKT9 antibody (Thermo Fisher, #14-0719-82) or human holo-transferrin (holo-Tf; Beyotime, #ST1135). The AAV-588-OligoPool-lib2 library was subsequently added for target enrichment. All downstream steps, including washing, Proteinase K digestion, and viral genome recovery, were performed as in the standard pull-down protocol.

### Viral Vector Administration and Tissue Collection

Adult mice (6-8 weeks old) were randomly assigned to receive viral vectors. Prior to injection, mice were briefly anesthetized with 2.5-3% isoflurane in an induction chamber. An eye drop of local anesthetic (oxybuprocaine 4 mg/mL) was applied to the right eye, and one minute later, AAV vectors were administered by a technician blinded to the labels of the viruses via retro-orbital sinus injection a BD MicroFine 0.3 ml insulin syringe fitted with a 31-gauge (0.3 mm × 8 mm) injection needle. Individual AAV capsid injections consisted of 5 × 10¹¹ viral genomes (vg) per mouse, while AAV-588-OligoPool lib200 or lib2 injections contained 1 × 10¹⁰ vg per mouse, each diluted in 10 - 50 μL of PBS supplemented with 0.001% F68. All mice recovered fully within minutes post-injection. Each individual viral vector was tested in at least three biological replicates, whereas AAV-588-OligoPool-lib200 or AAV-588-OligoPool-lib2 injections were evaluated in one or two animals. Three weeks post-injection, mice receiving AAV-588 OligoPool lib200 or lib2 were transcardially perfused with PBS, and tissues were collected for RNA recovery. In contrast, mice injected with individual AAV capsids were perfused with PBS followed by 4% paraformaldehyde (PFA), and tissues were harvested for histological analysis.

### Tissue RNA recovery

RNA extraction and analysis were performed to quantify AAV-588-OligoPool-lib200 and AAV-588-OligoPool-lib2 sequences in mouse brain and liver tissues. All procedures were conducted under RNase-free conditions to preserve RNA integrity. All tissue samples were cut to ∼50 mg each, flash-frozen in liquid nitrogen for 3 min, and homogenized using a tissue grinder (60 Hz, 60 s). Total RNA was subsequently extracted using the Ultrapure RNA Kit (Cowin, #CW0581), which incorporates TRIzol®-based lysis, following the manufacturer’s protocol. RNA was extracted from each organ in triplicate to reduce technical variability.

### Next-Generation Sequencing and data analysis

NGS libraries from RNA extraction and pull-down assay were generated by PCR amplification using primers NGS-1ST-F (5’-ACACTCTTTCCCTACACGACGCTCTTCCGATCTACTAACCCGGTAGCAACGGAGT-3’) and NGS-1ST-R (5’-GTGACTGGAGTTCAGACGTGTGCTCTTCCGATCTCTGCCAAACCATACCCGGAAGTATTCC-3’). For RNA samples, reverse transcription to cDNA was performed prior to amplification. Libraries were sequenced on an Illumina PE150 platform. Sequencing reads were aligned by matching to a defined left-flanking sequence (5’-GCCCAA-3’) and right-flanking sequence (5’-GCACAG-3’). Reads were excluded if the average quality score was <33 or if any nucleotide position between the flanking sequences had a quality score <20, in order to minimize sequencing uncertainty. Read counts were normalized to reads per million (RPM) prior to downstream analysis. Log₂ enrichment values were calculated as:

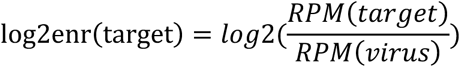

### Normalized Enrichment Analysis Approach for Pull-Down Assays

Raw sequencing values obtained from Fc-fusion pulldown assays are often influenced by experimental variability, including differences in yield, background binding, and competition effects, which can obscure true biological signals. To correct for these biases, we applied a correction strategy that models systematic deviations relative to a stable baseline condition and adjusts accordingly, thereby reducing assay-dependent inflation. This yields enrichment values that more faithfully capture relative capture efficiency rather than technical fluctuations, enabling more reliable comparisons across experimental groups.

### Histology and Immunostaining

For mice, PFA-fixed tissues were dehydrated in 15% and then 30% sucrose and embedded in the optimal cutting temperature compound for frozen sectioning. Typically, 60-µm-thick brain sections were cut for whole-section imaging of native fluorescence. Sections (10-20 µm thick) of the brain or other tissues (spinal cord, liver, muscle, heart, kidney, lung, spleen and DRG) were cut for standard immunohistochemistry with antibody staining and DAPI. The primary antibodies used included rabbit anti-NeuN (1:500, Ab177487, Abcam), chicken anti-GFAP (1:500, Ab4674, Abcam), rabbit anti-Olig2 (1:500, AB9610, Millipore), and goat anti-ChAT (1:200, AB144P, Millipore) antibodies. The secondary antibodies used were as follows: goat anti-chicken Alexa Fluor 647 (1:500, A32933, Thermo Fisher), donkey anti-rabbit Alexa Fluor 647 (1:500, A31573, Thermo Fisher), and donkey anti-goat Alexa Fluor 647 (1:500, A21447, Thermo Fisher). The tissue sections were blocked with PBS containing 5% donkey serum and 0.3% Triton X-100 surfactant and incubated with primary antibodies. The sections were subsequently washed three times with PBS and incubated with appropriate secondary antibodies. Finally, images were acquired by a technician blinded to the treatment groups via a virtual digital slide scanning system (VS120, Olympus) and a confocal laser scanning microscope (A1Ti, Nikon). Cell counting and analysis were performed via ImageJ.

### Statistical analysis

All experiments were randomized and blinded by an independent researcher before AAV treatment. Researchers remained blinded throughout the histological and biochemical assessments. The groups were unblinded at the end of each experiment during the statistical analysis. The statistical analyses were conducted via the SciPy Python library (version: 1.10.1). One-way ANOVA with Tukey’s post hoc test was used for data comparison. Statistical significance was determined when the p value was less than 0.05. Statistical analysis was conducted via the following Python libraries: Numpy v.1.26.4, SciPy v.1.10.1.1, seaborn v.0.12.2, scikit-learn v.1.2.2, statsmodels v.0.13.5, Matplotlib v.3.7.1, rdkit v.2023.9.5, pandas v.1.5.3, biopython v.1.80, and autogluon v.0.7.0.

### Construction and data preprocessing protocol for the ALICE system

In this work, the ALICE system uses the following 6 datasets for training and testing (Tables S1--2). (i) Pretraining dataset ***D***_***Pretrain***_ (Supplementary Table1): The pretraining dataset, ***D***_***Pretrain***_ , comprises 2,896,202 high-quality amino acid sequences meticulously curated from UniProt (specifically, the Swiss-Prot database, which has undergone manual review to ensure data integrity). To avoid the dominance of structural information from longer proteins, which could lead to insufficient learning of grammatical and semantic information, we limited the sequence length to 50 amino acids or less. After removing sequences containing nonnatural amino acids (X, Z, B, U, O), the dataset was split into training and test sets at an 8:2 ratio. (ii) Semantic-tuning dataset ***D***_***seq***_(Supplementary Table1): We used a publicly accessible dataset of 7-mer-modified AAV9 libraries ***Lib***_𝑨𝑨𝑽_ (random 7-mer amino acid sequences were inserted between residues 588-589 in VP1).^78^ These data include capsid variant production fitness^79^ (production fitness is an indicator used to evaluate the production capacity of a virus). The larger the value is, the greater the production capacity of the virus, and the smaller the value is, the weaker the production capacity of the virus.) Ly6a binds to Ly6a, which is widely expressed in C57BL/6J and FVB/NJ mice and is a known receptor for AAV-PHP. B family, facilitating blood‒brain barrier crossing^80^. Ly6a binding affinity^78^ represents the ability of AAV variants to bind to the Ly6a protein. and Ly6c1 binding affinity (Ly6c1 was identified as a novel receptor associated with BBB crossing in Ly6a-deficient mice^81^ such as BALB/cJ. Ly6c1 binding affinity^78^ represents the ability of AAV variants to bind to the Ly6c1 protein. Capsid sequences with high production fitness, high Ly6a binding affinity, and high Ly6c1 binding affinity were selected as training data for the semantic tuning model ***D***_***seq***_ (n = 72,753). (iii) Production fitness dataset ***D***_***Production Fitness***_ (Supplementary Table1): This dataset is derived from ***Lib***_𝑨𝑨𝑽_ and uses the production fitness metric to quantify the viral production capacity of different AAV sequences. A total of 119,851 data points are used to train a model to evaluate the viral production capacity of capsid sequences. (iv) Ly6a Target Binding Dataset ***D***_***Ly6a***_ (Supplementary Table1): This dataset is derived from ***Lib***_𝑨𝑨𝑽_, with a total of 89,169 data points, which can be used for model training and evaluation of the binding ability of capsid sequences to the Ly6a target. (v) Ly6c1 Target Binding Dataset ***D***_***Ly6c1***_ (Supplementary Table1): This dataset is derived from ***Lib***_𝑨𝑨𝑽_, with a total of 89,040 data points, which can be used for model training and evaluation of the binding ability of capsid sequences to the Ly6c1 target. (vi) Evolutionary Reference Datasets ***Ref***\*& ***Ref***^+^(Supplementary Table1-2): From ***Lib***_𝑨𝑨𝑽_, we selected sequences with high production fitness, strong binding affinity to the ly6a target, and strong binding affinity to the ly6c1 target, as well as sequences previously reported to be effective in vivo in mice ***Ref***\*. The sequences with poor performance in all three functions were chosen as ***Ref***^+^. These datasets are employed in the heuristic evolutionary approach within the function-guided evolution (FE) module for limited data scenarios.

To train ALICE, we generated four main datasets:

- **Pretraining datasets** ***D***_***Pretrain***_: The dataset comprises 2.89 million peptide sequences from UniProt. To train the BERT backbone, we implemented data preprocessing consistent with the original BERT methodology, including random masking and substitution of specific amino acids in these peptides. Additionally, we divided the sequences into pairs of sentences and randomly matched them to create the next sentence prediction task for BERT pretraining. For the RoBERTa-backbone, we employed a similar preprocessing approach tailored for RoBERTa tasks (details in the Supplementary Methods).
- ***Lib***_𝑨𝑨𝑽_: This dataset originates from a publicly available library of 7-mer-modified AAV9 capsid sequences, where amino acid sequences were inserted between residues 588-589 of the VP1 protein. The raw data were rigorously cleaned and preprocessed to remove low-quality or redundant sequences.
- **Semantic tuning dataset** ***D***_𝑺𝒆𝒒_ : This dataset comprises a publicly accessible AAV capsid sequence library, where 7-mer amino acid sequences are inserted between residues 588 and 589 in the VP1 of the AAV9 capsid. The data include information on capsid variant production fitness and binding affinity to the target proteins Ly6a and Ly6c1. Capsid sequences exhibiting high production fitness and target binding abilities were selected for the semantic-tuning model (***D***_𝑺𝒆𝒒_, n = 72,753; details in the Supplementary Methods).
- **Evolutionary reference datasets** ***Ref***\* **&** ***Ref***^+^: To promote the evolution of capsid sequences toward desired functions, we prioritized sequences in the FE reference dataset by focusing on two key factors:

▪ High viability and strong binding affinity to Ly6a or Ly6c1.

▪ AAV variants with desired functions as reported in previous studies.

Sequences meeting these criteria were designated as ***Ref***\* (n = 76, Supplementary Table2), while those exhibiting weak binding to Ly6a and Ly6c1 and low viability were classified as ***Ref***^+^ (n = 53, Supplementary Table2).

- **Capability evaluation dataset:** To evaluate the production fitness, Ly6a binding, and Ly6c1 binding capabilities of the designed capsid sequences during the module optimization process, three datasets ( ***D***_***Production Fitness***_, ***D***_***Ly6a***_ , and ***D***_***Ly6c1***_ ) were constructed (details in the Supplementary Methods).

Through rigorous examination and validation of data features and label distributions, we meticulously removed low-quality and anomalous data to ensure high-quality training data for ALICE.

### Training and inference regimens

For each component module of ALICE, we trained multiple candidate models via diverse architectures and hyperparameters. The optimal parameter configurations were determined through grid search, random search, and manual adjustment. The training data included sequences sourced from UniProt, the AAV capsid library, and other repositories (details in the Supplementary Methods). ALICE was trained using an RTX 3090 GPU. Architecture exploration was facilitated on a server equipped with four RTX 3090 GPUs. Training the entire ALICE system required only 7.25 GPU hours and involved 51 million parameters. Inference for ALICE can be efficiently conducted via a single RTX 3090 GPU.

### Metrics

All methods, including the RMSE, Pearson, Spearman, R², MedAE, and C-index, serve as traditional regression evaluation metrics, aiding in quantifying and selecting the best evaluation models among all candidate algorithms. The FE score, KL divergence, BLOSUM62 similarity, n-gram similarity, LCS ratio, mutual information and HR score are calculated via defined or well-established formulae. Where 𝑦:serves as the prediction value, 𝑦 is the mean of the true value, and 𝑦Y; are the true values in the dataset, i =1, 2…, n, where n is the total amount of the dataset. X denotes the ALICE designed capsid sequences, and Y denotes the reference sequences ***Ref***\* and ***Ref***^+^.

- **RMSE (**root mean squared error): The RMSE measures the average magnitude of the errors between the predicted and observed values, providing a straightforward measure of prediction accuracy by penalizing larger errors more heavily:

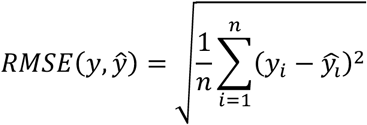

- **Pearson correlation coefficient**: This coefficient assesses the linear correlation between the predicted and observed values and ranges from −1--1. A value of 1 indicates a perfect positive linear relationship, −1 indicates a perfect negative linear relationship, and 0 indicates no linear relationship:

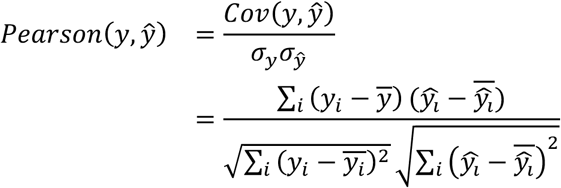

- **Spearman rank correlation coefficient**: Spearman’s coefficient evaluates the monotonic relationship between the predicted and observed values by considering their rank orders. It ranges from −1 to 1, with values closer to 1 or −1 indicating stronger correlations:

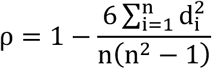

where 𝑑: is the difference between the ranks of the corresponding values 𝑦: and 𝑦Y;.

- **R^2^ (coefficient of determination)**: R^2^ quantifies the proportion of variance in the observed values that is predictable from the predicted values, ranging from 0--1. A value closer to 1 indicates a better fit of the model to the data:

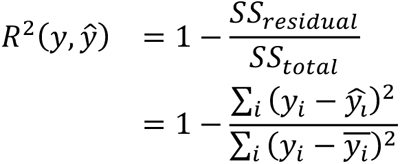

- **MedAE (**median absolute error): The MedAE calculates the median of the absolute errors between the predicted and observed values, providing a robust measure of prediction accuracy that is less sensitive to outliers than the RMSE is:

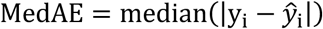

- **C-index (Concordance Index)**: The C-index evaluates the predictive accuracy of a model by measuring the agreement between the predicted and observed rankings. It ranges from 0.5 to 1, with values closer to 1 indicating better model performance in predicting the correct order of observations:

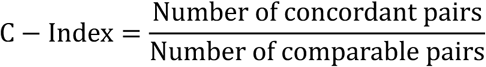

- **FE score:** We defined the FE score by the following FE function. The FE score measures the evolutionary changes toward 𝑅𝑒𝑓* or 𝑅𝑒𝑓^+^. It is correlated with the reference sequences, 𝑅𝑒𝑓*and 𝑅𝑒𝑓^+^, and is calculated via the following formula:

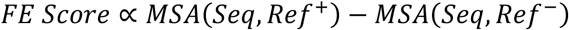

where 𝑆𝑒𝑞 denotes the capsid sequences designed by the FE module. The ablation study in Fig. 2d uses FE score and predicted production fitness as evaluation metrics because experimental measurements cannot feasibly be performed at the required scale for every ablated variant of the pipeline. FE is a biologically anchored scoring function derived from positive and negative reference sets, and in prior sections we show that it correlates with experimentally validated functional outcomes. While we acknowledge that predictive metrics have inherent limitations, they provide a consistent and biologically meaningful proxy for isolating each module’s contribution within the computational framework.

- **KL Divergence**^82^: KL divergence, or Kullback‒Leibler divergence, quantifies the difference between two probability distributions. It measures how one probability distribution diverges from a second, expected distribution, helping to understand the evolutionary divergence of sequences by quantifying deviations in amino acid distribution from a reference. A lower KL divergence indicates closer similarity to the reference distribution, suggesting less evolutionary change, whereas a higher KL divergence indicates more divergence:

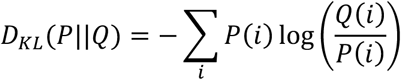

where 𝑃 is the probability distribution of ALICE-designed capsid sequences and where 𝑄 is the probability distribution of reference sequences 𝑅𝑒𝑓* or 𝑅𝑒𝑓^+^.

- **BLOSUM62 Similarity**^56^: BLOSUM62 (Blocks Substitution Matrix 62) is a scoring matrix for protein sequence alignment that is based on observed substitutions in blocks of local alignments from related proteins. High BLOSUM62 similarity scores indicate more conserved sequences, highlighting their potential functional or structural importance. While BLOSUM62 identifies conserved regions, KL divergence quantifies overall evolutionary change, making both measures complementary in evolutionary studies:

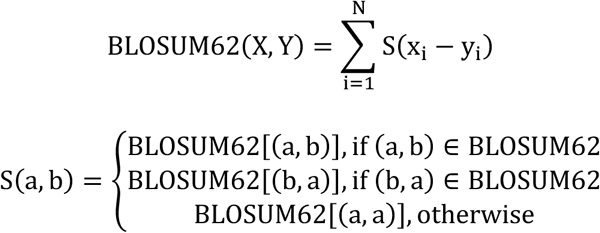

where N = minflen(X), len(Y)g and where S(a, b) is the BLOSUM62 score for amino acids a and

- b.
- **N-gram Similarity**^83^: This metric gauges local semantic similarity by measuring the similarity of short sequence fragments. An increase in n-gram similarity indicates that local sequence patterns are becoming more similar, reflecting evolutionary adaptations at the local sequence level:

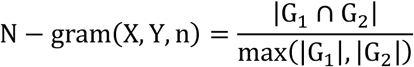

where G_?_ and G_<_ are the sets of n-grams from set X and set Y, respectively, and where |G| denotes the total count of n-grams in set G, including repetitions.

- **LCS ratio**^58^: The longest common subsequence (LCS) ratio measures the overall similarity between two capsid sequences. In this study, it assesses the global semantic similarity between capsid sequences designed via FE and the reference sequences 𝑅𝑒𝑓* and 𝑅𝑒𝑓^+^. The LCS ratio tracks the evolutionary trajectory of global semantic information, focusing on the overall structure rather than exact local matches:

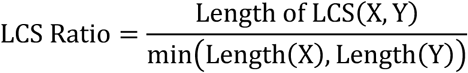

where Length of LCS(X, Y) is the length of the longest common subsequence between sequences X and Y . Length(X) and Length(Y) are the lengths of the capsid sequences in sets X and Y , respectively.

- **Mutual Information**^84^: This metric evaluates the correlation of capsid sequences designed by the FE module with 𝑅𝑒𝑓* and 𝑅𝑒𝑓^+^. According to the FE function, as evolutionary progress, sequences evolve in a direction guided by 𝑅𝑒𝑓* and 𝑅𝑒𝑓^+^, indicating increasing correlation with these reference sequences:

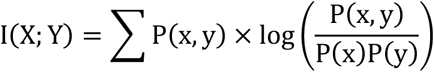

where P(x, y) is the joint probability distribution of X and Y , where P(x) , and where P(y) = represents the marginal probability distributions.

- **HR score**: We define the HR score to select ultimate candidates on the basis of multiple functions, including the binding capabilities to Ly6c1 and Ly6a and production fitness, via the following function:

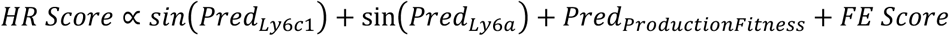

where 𝑃𝑟𝑒𝑑_N@PQ?_ and 𝑃𝑟𝑒𝑑_N@PI_ represent the predicted binding abilities of the capsid sequences to these receptors and where 𝑃𝑟𝑒𝑑_RDLGHQK:L=S:K=EFF_ represents the predicted production fitness, all of which are evaluated by the GB model. This score is used to rank and select the best candidates from a comprehensive perspective.

- **CP score**: To evaluate the multifunctional performance of ALICE and ALICE-X, we introduce a Comprehensive Prediction (CP) Score. The CP Score is defined separately for each task as follows:
- in Ly6c1/Ly6a task, the CP Score was defined as:

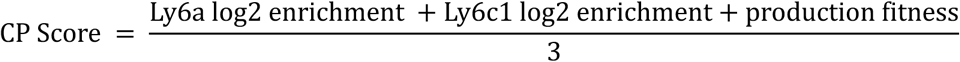

or:

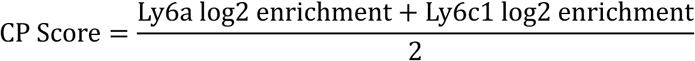

2 in hTfR1 task, the CP Score was defined as:

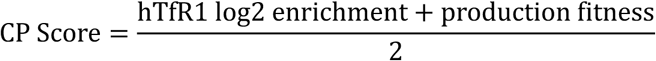

The Comprehensive Prediction (CP) Score provides an integrated metric that can be derived either from experimental measurements or from evaluation model predictions. For the hTfR1 task, for example, CP Scores are calculated using experimentally measured production fitness and hTfR1 log2 enrichment values to capture overall performance, and can likewise be computed from their predicted counterparts. This enables direct comparison of experimental outcomes with model-based predictions, thereby assessing the comprehensive predictive power of ALICE-designed capsid sequences. An analogous procedure was applied to the Ly6c1/Ly6a task.

### Definitions of the four evolution trajectory types

In the ablation experiment of this study, to facilitate the analysis of the evolutionary trajectories and types of capsid sequences generated by eight combination backbones compared with those of the NNK group in terms of semantics and functionality, we defined four evolution types:

- Evo-Type I: Evolution toward both ***Ref***\* and ***Ref***^+^.
- Evo-Type II: Evolution toward ***Ref***\* and away from ***Ref***^+^.
- Evo-Type III: Evolution away from both ***Ref***\* and ***Ref***^+^.
- Evo-Type IV: Evolution away from ***Ref***\* and toward ***Ref***^+^.

## Lead contact

Further information and requests for resources and reagents should be directed to and will be fulfilled by the lead contact, Xuhua Wang.

## Materials availability

All unique reagents generated in this study (plasmids generated in this work, stable cell lines etc.) are available upon reasonable request from the lead contact.

The data that support the findings of this study are available on reasonable request to the lead corresponding author (X.W.).

## Data and code availability

All the data generated and described in the article and its Supplementary Information are available upon publication. The pretraining dataset includes 2.89 million protein peptide sequences. The semantic tuning dataset contains 72,753 AAV capsid sequences to learn the semantic language of the AAV. The dataset for fusing multiple functionalities, comprising 129 sequences of positive and negative samples, is available in Supplementary Table2. A total of 119,851 capsid sequences were collected for training the model to predict production fitness. Additionally, 89,169 capsid sequences were used for training the model to predict Ly6a binding ability, and 89,040 capsid sequences were used to train the model to predict Ly6c1 binding ability (see Source data for full list of sequences).

- A total of 6,387 capsid sequences were compiled to train and test the evaluation model of ALICE-X, which predicts both production fitness and hTfR1 binding capability (see Source data for full list of sequences). For benchmarking multifunctional integration, a curated reference dataset comprising 75 sequences (positive ***Ref***\* , n = 27; negative ***Ref***^+^ , n = 48) was assembled (Supplementary Table12).
- From the full set of 6,387 sequences, a semantic tuning dataset was derived by selecting variants with comparatively favorable receptor engagement and packaging profiles. Variants were filtered using two relative log2 enrichment proxies—hTfR1 log2 enrichment > −2 and production fitness > −1, yielding 1,388 unique capsid sequences for fine-tuning.

To contribute to the scientific community, we offer the following models with associated code and sequencing data as open resources for noncommercial applications, accessible upon publication. The ALICE and ALICE-X system encompasses meticulously calibrated modules, each fine-tuned with optimized hyperparameters. The training and fine-tuning files for each module of the ALICE and ALICE-X model are deployed at the aforementioned link. Additionally, we provide scripts to regenerate our training datasets, as well as scripts for training and generating models that replicate our results.

Any additional information required to reanalyze the data reported in this paper is available from the lead contact upon request.

## Experimental model and study participant details

All animal protocols were approved by the Institutional Animal Care and Use Committee, used following the provisions of the Zhejiang University Animal Experimentation Committee (ZJU20210110), and were conducted in accordance with the guidelines for the care and use of laboratory animals. Both sexes of BALB/c, C57BL/6J, FVB/NJ mice, and *hTFRC* Knock in (KI) mice, aged 6-8 weeks, were obtained from the Zhejiang Center of Laboratory Animal, Shanghai SLAC Laboratory Animal, or Biocytogen. The mice were housed in groups of 2-4 per cage under a 12 h light‒dark cycle with standard rodent chow and water ad libitum.

## Supplemental Information

Further information on the research design is available in the Supplementary Information linked to this paper.

## Acknowledgments

The authors acknowledge the excellent technical staff at the Imaging Facility, Core Facility of Zhejiang University School of Medicine and Center of Cryo-Electron Microscopy of Zhejiang University for their assistance with confocal laser scanning microscopy and scanning electron microscopy. This study was supported by the Scientific and Technological Innovation 2030 Program of China - major projects (2021ZD0200408 to X.W.), the National Natural Science Foundation of China (82371374 to X.W.), the Zhejiang Provincial Natural Science Foundation of China for Distinguished Young Scholars (RG25H090001 to X.W.) and the Fundamental Research Funds for the Central Universities (226-2025-00226, 226-2024-00087 to X.W.), the grant from Nanhu Brain-computer Interface Institute (010904006 and 010904009 to X.W.).

## Author Contributions

B.G., H.Z., X.L, and H.J. contributed equally to this work. X.W., H.Z. and B.G. conceptualized and designed the study. H.Z., B.G., X.L., H.J., H.L., Y.W., A.M., H.W., and R.Z. conducted the experiments and collected the data. X.W., H.Z., B.G., R.Z., Y.W., A.M., H.W., Y.H., Z.C., Y.Y., Z.C., X.G. and B.Y. analyzed and interpreted the data. H.Z., B.G., X.W., N.C. drafted the paper. All the authors critically revised the manuscript and approved the final version for submission.

---

**Extended Data Fig. 1.**
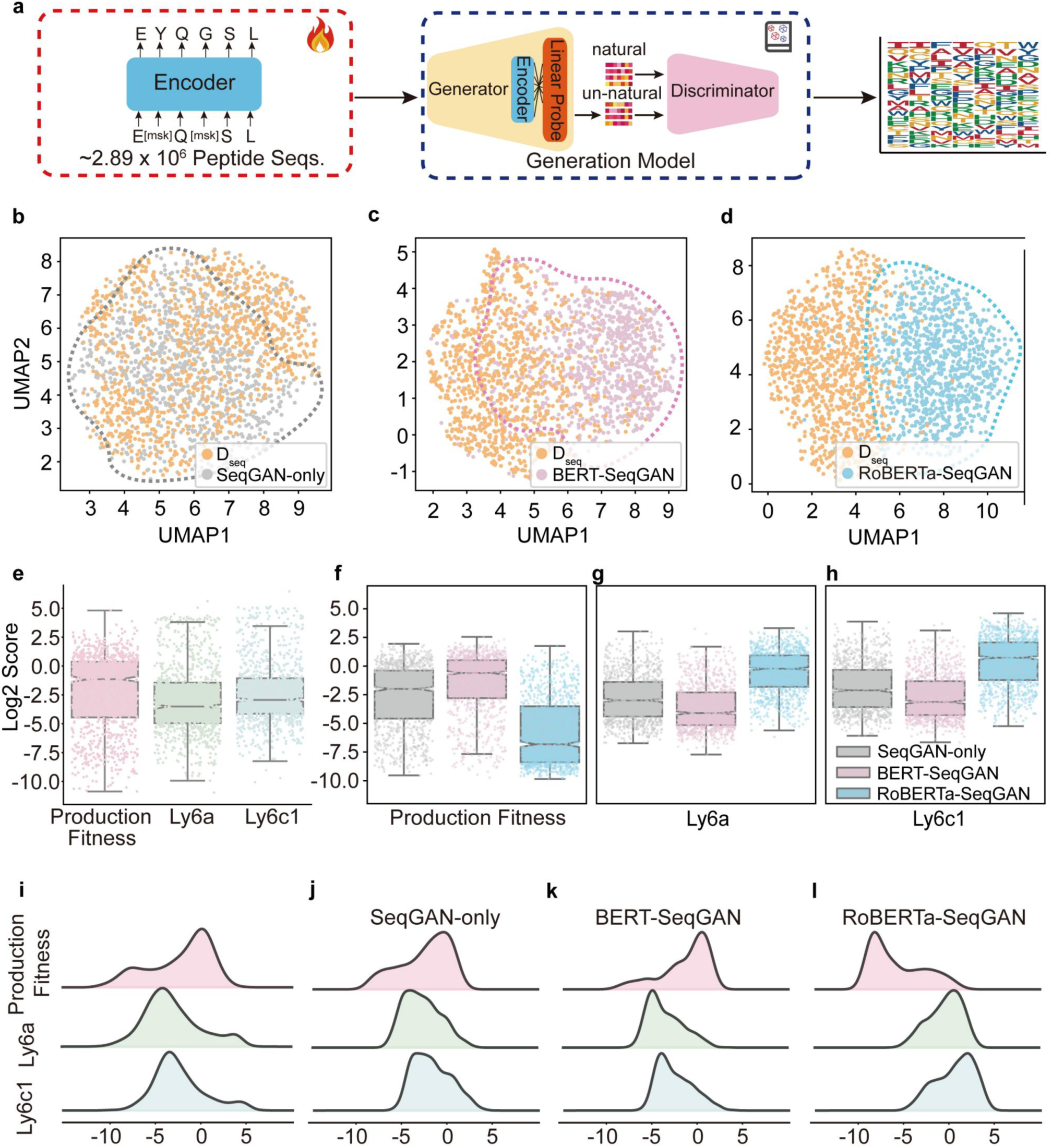
Functional outcomes of capsid sequences designed through pretraining and semantic tuning strategies. **a,** Schematic diagram of the pretraining and semantic tuning modules. **b-d,** Diagrams depict the feature distributions of sequences generated by the language models with different architectures, including SeqGAN-only (b), BERT-SeqGAN (c), and RoBERTa-SeqGAN (d), in comparison to those of the sequences in the semantic tuning training dataset ( ***D***_***seq***_). In each category, a random sample of 1000 sequences was extracted for analysis. **e-h,** Histograms showing the distributions of three functions (production fitness, Ly6a and Ly6c1 binding ability) of the sequences in the virtual evaluation dataset (***D***_***Production Fitness***_, ***D***_***Ly6a***_ and ***D***_***Ly6c1***_) (e) and the sequences generated by language models with different architectures, including SeqGAN-only (f), BERT-SeqGAN (g), and RoBERTa-SeqGAN (h). In each category, a random sample of 1000 sequences was extracted for analysis. Box plots are shown as the median, top and bottom quartiles. The color of the bar indicates the outcomes from different architectures, as indicated by the color bar provided in (h). **i‒l,** Ridgeline plots showing the distributions of three functions (production fitness, Ly6a and Ly6c1 binding ability) of the sequences in the virtual evaluation dataset (***D***_***Production Fitness***_ , ***D***_***Ly6a***_ and ***D***_***Ly6c1***_) (i) and the sequences generated by language models with different architectures, including SeqGAN-only (j), BERT-SeqGAN (k), and RoBERTa-SeqGAN (l). In each category, a random sample of 1000 sequences was extracted for analysis.

**Extended Data Fig. 2.**
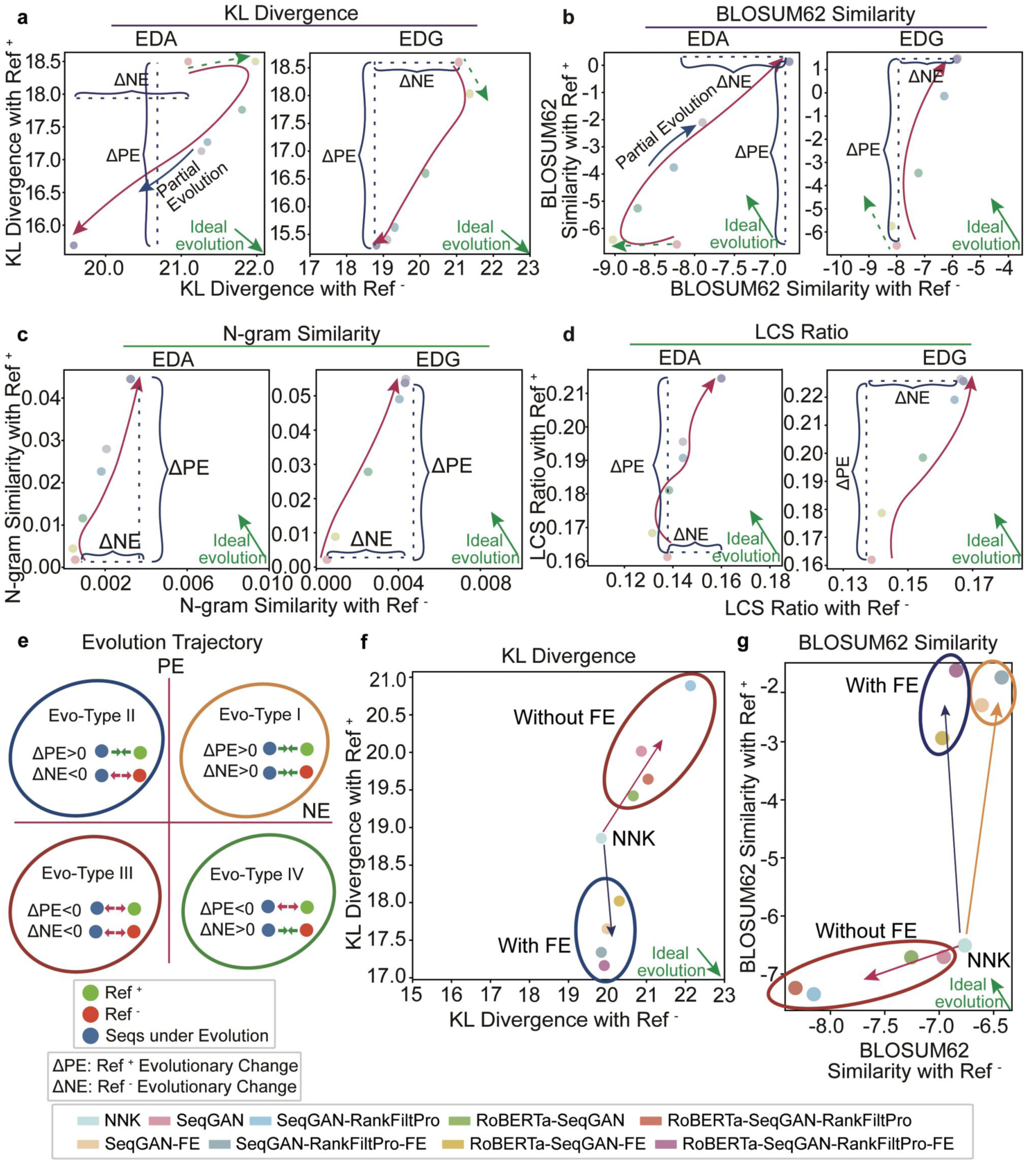
The ablation of each module significantly affects the evolutionary trajectory of sequences. **a-d,** The plots illustrate the evolutionary trajectories measured by KL divergence (a), BLOSUM62 similarity (b), N-gram similarity (c), and the LCS ratio (d) reveal the evolving functionality and semantic characteristics. The green arrow indicates the ideal evolution direction, whereas the green dashed arrow shows the direction of evolution within the initial 5 epochs. **e,** Definitions of the four evolutionary trajectory types. **f-g,** The narrative depicts the evolutionary changes in sequences engineered by architectures with different ablations, as measured by KL divergence (f) and BLOSUM62 similarity (g), in comparison to the sequences in the control NNK dataset. The colors of the dots represent the outcomes from architectures with distinct ablations, as indicated by the color bar provided at the bottom. The green arrow signifies the ideal evolution direction. The arrows colored accordingly show the direction of evolution of sequences engineered by architectures with different ablations.

**Extended Data Fig. 3.**
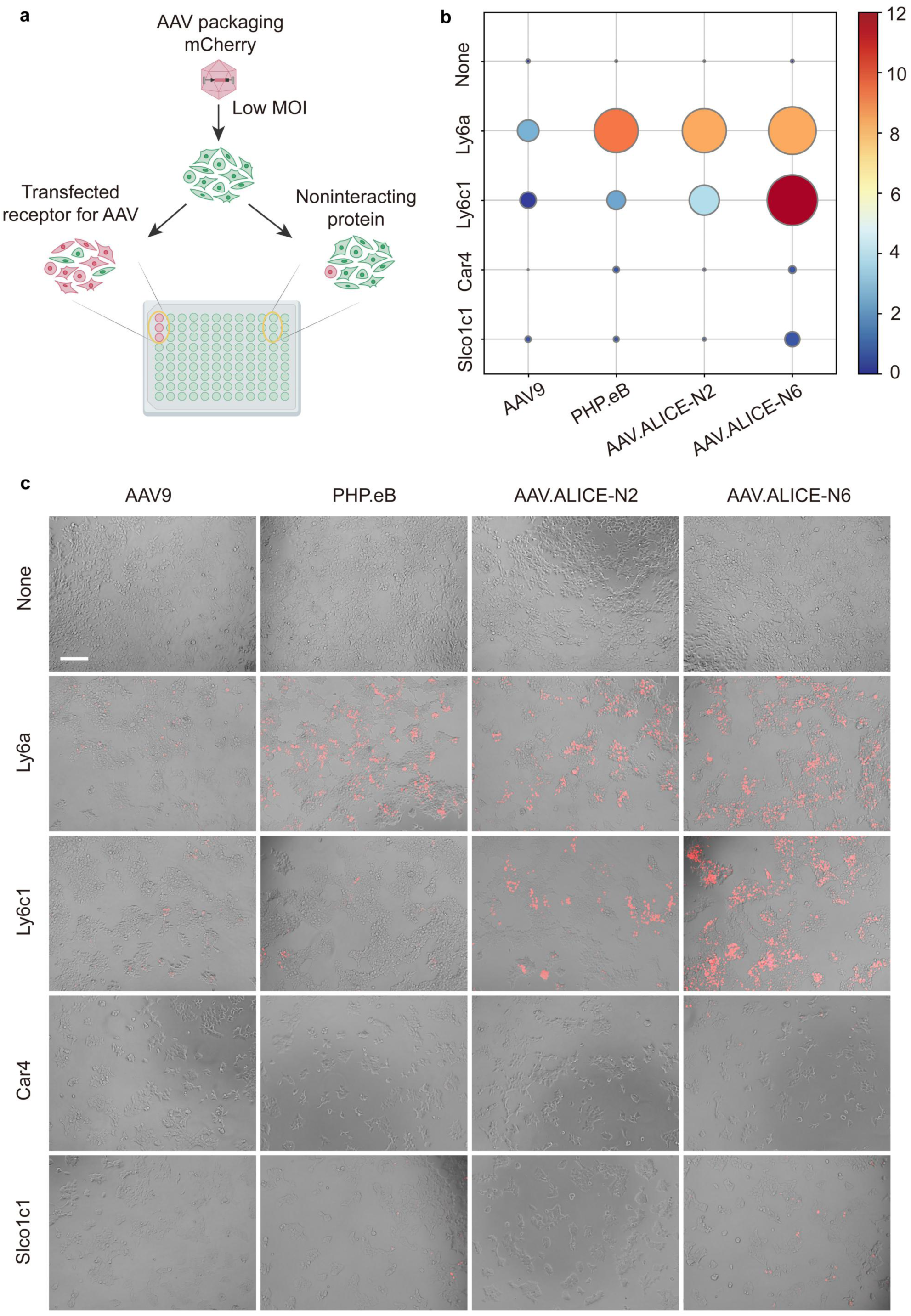
Validation of the mechanisms of action for AAV.ALICE-N2 and AAV.ALICE-N6 through in vitro experiments. **a,** The mechanisms of action of AAV variants were clarified by observing the AAV fluorescent protein transgene levels at a low multiplicity of infection (MOI) for different protein receptors. **b,** Infection efficacy of AAV9, PHP.eB, and AAV.ALICE-N2, and AAV.ALICE-N2 in HEK293T cells and HEK293T cells transfected with different protein receptors (Ly6a, Ly6c1, Car4 and Slco1c1). Extent of infection (max, 36.918; min, 0.031) and total brightness per signal area (max, 11.880; min, 0.035) were recorded. **c,** Representative images of HEK293T cells cultured in 96-well plates without and with receptor protein transfection, infected with 1 × 10^7^ v.g. AAV variant per well, and imaged 24 hours after transduction. An overlay of brightfield and fluorescence images is shown. Scale bar, 200 μm.

**Extended Data Fig. 4.**
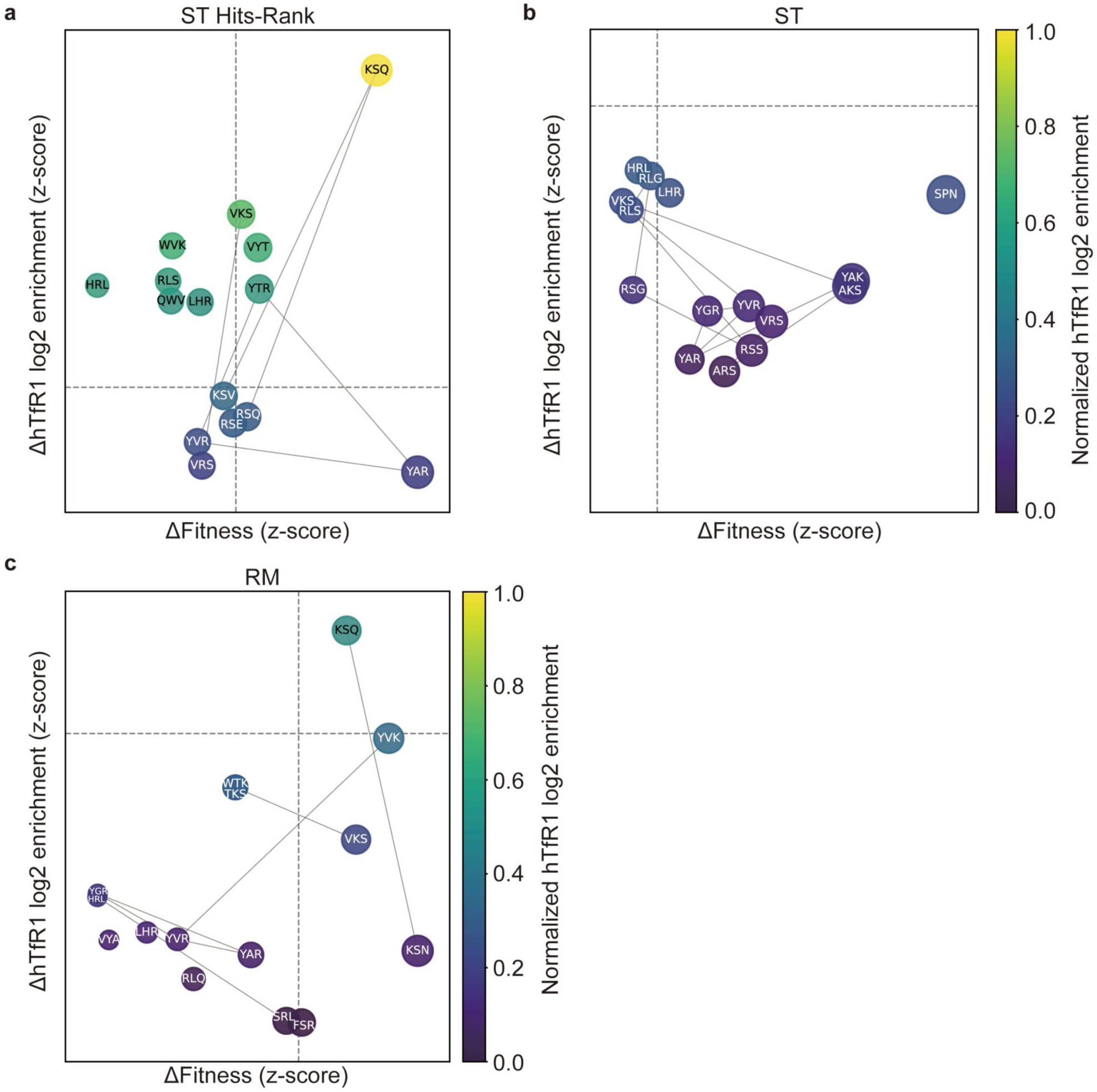
Motif-level contributions to hTfR1 log2 enrichment and production fitness in capsid sequences generated by different models. **a-c,** The performance of motifs generated by Semantic Tuning—Hits Ranking (ST Hits-Rank) model (a), Semantic Tuning—only (ST) model (b), and Random Mutation (RM) of sequences in ***Ref***\*(c). Scatter plots depict the effects of the top 15 motifs identified from each model on human transferrin receptor 1 (hTfR1) log2 enrichment and production fitness, calculated as the normalized z-score difference relative to the distribution of other sequences. Each circle represents a motif, with the x-axis indicating ΔFitness (z-score) and the y-axis indicating ΔhTfR1 log2 enrichment (z-score). Node color encodes normalized hTfR1 log2 enrichment, and edges connect motifs with shared sequence similarity.

**Extended Data Fig. 5.**
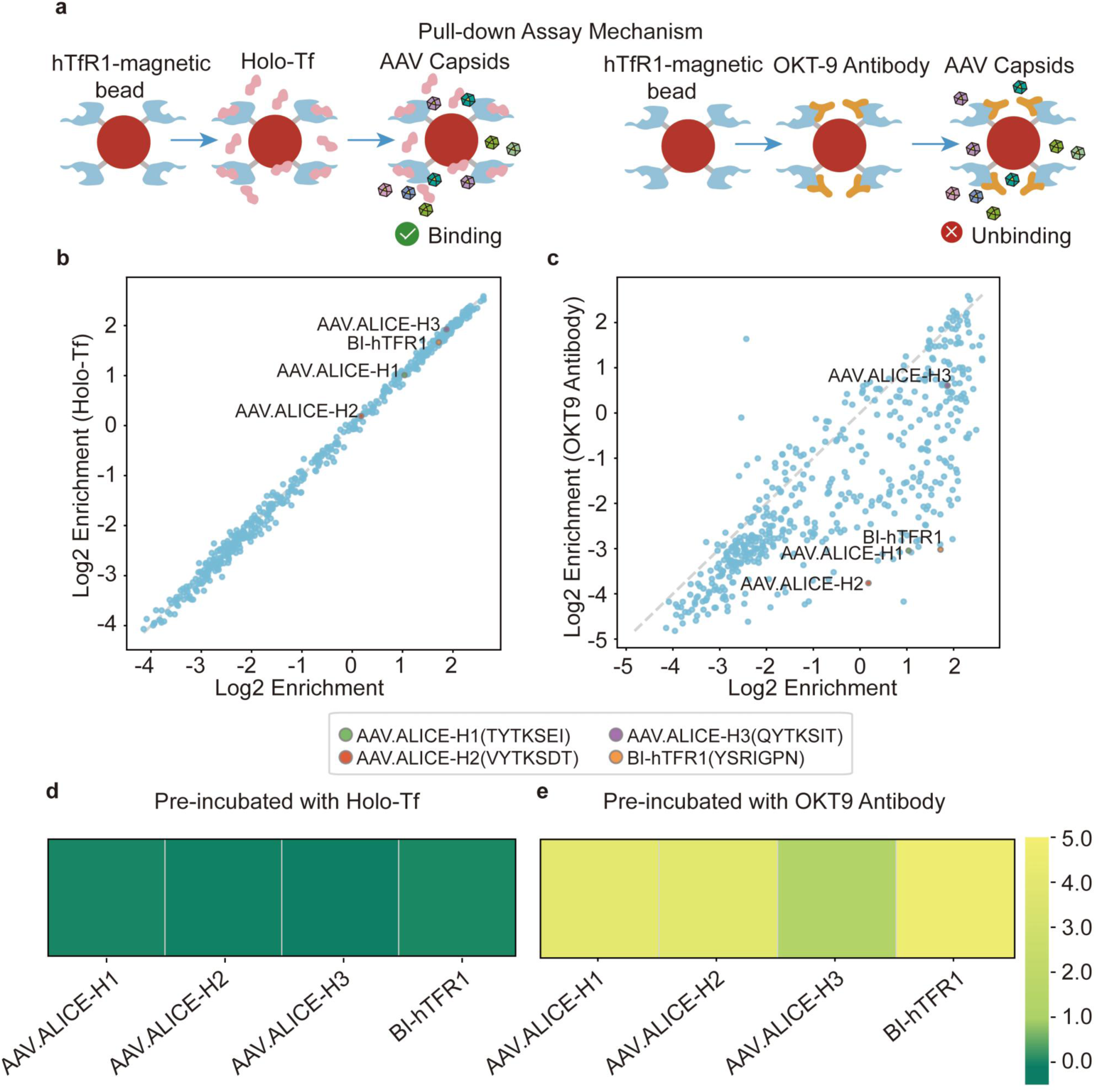
Mechanistic studies of engineered AAV variants binding to human TfR1. **a,** Schematic of the competitive pulldown assay. hTfR1-Fc-coated beads were pre-incubated with either the OKT9 antibody or human holo-transferrin (holo-Tf) prior to exposure to the AAV library, to assess competition for receptor binding. **b-c,** Comparison of variant enrichment (log2) on hTfR1 in the absence (x-axis) and presence (y-axis) of holo-Tf (b) and OKT9 antibody (c). **d-e,** Heatmap depicting the difference in log2 enrichment (Δlog2) for selected variants (AAV.ALICE-H1, H2, H3, and BI-hTFR1) between the hTfR1 and hTfR1 + holo-Tf (E) or hTfR1 + OKT9 antibody (e) conditions.

