## Supplemental Methods and Figures for "Mapping AAV capsid sequences to functions through function-guided *in silico* evolution"

### Key Resource Table

| REAGENT or RESOURCE | SOURCE | IDENTIFIER |
|---|---|---|
| Antibodies | | |
| rabbit anti-NeuN | Abcam | Cat# ab177487, RRID: AB_2532109 |
| chicken anti-GFAP | Abcam | Cat# ab4674, RRID: AB_304558 |
| rabbit anti-Olig2 | Millipore | Cat# AB9610, RRID: AB_570666 |
| goat anti-ChAT | Millipore | Cat# AB144P, RRID: AB_2079751 |
| rabbit Anti-SOX9 | Abcam | Cat#ab185966, RRID: AB_2728660 |
| goat anti-chicken Alexa Fluor 647 | Thermo Fisher | Cat# A32933, RRID: AB_2762845 |
| donkey anti-rabbit Alexa Fluor 647 | Thermo Fisher | Cat# A31573; RRID: AB_2536183 |
| donkey anti-goat Alexa Fluor 647 | Thermo Fisher | Cat# A21447, RRID: AB_2535864 |
| donkey anti-goat YSFluor 488 | Yeasen | 34306ES60 |
| Bacterial and virus strains | | |
| DH5α Chemically Competent Cell | Tsingke | TSC-C01 |
| AAV9 ss-EF1α-H2B-mCherry | This paper | N/A |
| AAV.ALICE-N1 ss-EF1α-H2B-mCherry | This paper | N/A |
| AAV.ALICE-N2 ss-EF1α-H2B-mCherry | This paper | N/A |
| AAV.ALICE-N3 ss-EF1α-H2B-mCherry | This paper | N/A |
| AAV.ALICE-N4 ss-EF1α-H2B-mCherry | This paper | N/A |
| AAV.ALICE-N5 ss-EF1α-H2B-mCherry | This paper | N/A |
| AAV.ALICE-N6 ss-EF1α-H2B-mCherry | This paper | N/A |
| AAV.ALICE-N7 ss-EF1α-H2B-mCherry | This paper | N/A |
| AAV.ALICE-N8 ss-EF1α-H2B-mCherry | This paper | N/A |
| AAV.ALICE-H1 ss-EF1α-H2B-mCherry | This paper | N/A |
| AAV.ALICE-H2 ss-EF1α-H2B-mCherry | This paper | N/A |
| AAV.ALICE-H3 ss-EF1α-H2B-mCherry | This paper | N/A |
| Chemicals, peptides, and recombinant proteins | | |
| Recombinant mouse Ly6C1-Fc Protein | Genscript | N/A |
| Recombinant mouse Ly6a-Fc Protein | Genscript | N/A |
| Recombinant human TfR1-Fc Protein | Genscript | N/A |
| Recombinant macaque TfR1-Fc Protein | Genscript | N/A |
| Recombinant Fc Protein | Genscript | N/A |
| Recombinant human IgG1-Fc Protein (C103S) | SinoBiological | 10702-HNAH |
| Human Holo Transferrin | Beyotime | ST1135 |
| OKT9 | Thermo Fisher | 14-0719-82 |
| Proteinase K | TIANGEN | RT403 |
| PEG8000 | BIOFROXX | 1363GR500 |
| DMEM | Gibco | C11995500 |
| 0.25% Trypsin-EDTA | Gibco | 25200-072 |
| 10X Penicillin-streptomycin | Beyotime | C0222 |
| iodixanol | Sigma | D1556-250ML |
| ampicillin | Beyotime | ST008 |

| | | |
|---|---|---|
| DPBS | Gibco | 14190144 |
| Tween-20 | Sigma-Aldrich | P7949 |
| BSA | Sigma-Aldrich | A1933 |
| 10% PLURONIC F-68 | gibco | 24040-032 |
| PEI | Biohub | 78PEI40000 |
| DNaseI | Sigma-Aldrich | AMPD1 |
| Critical commercial assays | | |
| Dynabeads protein A | Thermo Fisher | 10001D |
| cDNA First-Strand Synthesis Kit | ABclonal | RK20400 |
| SYBR Green Fast qPCR Mix | ABclonal | RK21203 |
| Ultrapure RNA Kit | Cowin | CW0581 |
| Endo-Free Plasmid Maxi Kit | Omega | D6926 |
| 2× Phanta Max Master Mix | Vazyme | P515 |
| XbaI | NEB | R0145V |
| AgeI | NEB | R3552L |
| Gibson Assembly Master Mix | NEB | E2611 |
| Gel DNA Recovery kit | Genstone | TD407 |
| Trypsin-EDTA | gibco | 25200-072 |
| Deposited Data | | |
| RNA sequencing data | This paper | Accessible upon publication |
| Experimental Models: Cell Lines | | |
| HEK-293T | ATCC | Cat# CRL-3216 |
| Experimental Models: Organisms/Strains | | |
| Mouse: BALB/c | Zhejiang Center of Laboratory Animal | N/A |
| Mouse: C57BL/6 | Zhejiang Center of Laboratory Animal | N/A |
| Mouse: FVB/NJF | Shanghai SLAC Laboratory Animal | N/A |
| Mouse: hTFRC knock in (B-hTFR1) | Biocytogen | 110861 |
| Oligonucleotides | | |
| Assembly-XbaI-F, see Supplementary Table9 | This paper | N/A |
| Assembly-AAV9-588i7NNK, see Supplementary Table9 | This paper | N/A |
| ITR-F, see Supplementary Table9 | This paper | N/A |
| ITR-R, see Supplementary Table9 | This paper | N/A |

| | | |
|---|---|---|
| NGS-1ST-F, see Supplementary Table9 | This paper | N/A |
| NGS-1ST-R, see Supplementary Table9 | This paper | N/A |
| Oligonucleotide library synthesis pool 200, see Source Data Supplementary Table22 | This paper | N/A |
| Oligonucleotide library synthesis pool 1, see Source Data Table Lib1 Source | This paper | N/A |
| Oligonucleotide library synthesis pool 2, see Source Data Table Lib2 Source | This paper | N/A |
| Recombinant DNA | | |
| pAAV2/9n | Addgene | 112865 |
| pUC-Cap9XA | This paper | N/A |
| pAAV-EF1α-H2B-mCherry | This paper | N/A |
| pAAV-P41-Rep78-Cap9_588iSTOP-WPRE-hGH | This paper | N/A |
| pRepAAP | This paper | N/A |
| pHelper | Agilent | 240071-12 |
| pUC19 | Sangon Biotech | B610005-0050 |
| pAAV-588-OligoPool-lib200 | This paper | N/A |
| pAAV-588-7NNK-lib | This paper | N/A |
| pAAV-588-OligoPool-lib1 | This paper | N/A |
| pAAV-588-OligoPool-lib2 | This paper | N/A |
| pAAV-P41-Rep78-Cap9-588i7mer-WPRE-hGH oligo library | This paper | N/A |
| pcDNA3.4 | GeneScript | N/A |
| pHAGE-P2A-eGFP | This paper | N/A |
| PMD2.G | Addgene | 12259 |
| psPAX2 | Addgene | 12260 |
| Software and Algorithms | | |
| Python | Open Source | https://www.python.org/ |
| RDKit | Open Source | https://github.com/rdkit |
| Pytorch | Open Source | https://pytorch.org/ |
| Biopython | Open Source | https://biopython.org/ |
| Adobe Illustrator 2023 | Adobe System | https://www.adobe.com/ |

### RESOURCE AVAILABILITY

### METHODS DETAILS

#### 1. Statistical analysis between nature language and biology language

The essence of deep learning lies in harnessing vast datasets to train neural networks, enabling them to autonomously learn features from the data. This learning process is grounded in the statistical properties inherent to the data itself, a principle known as statistical learning. To investigate the alignment of semantic information in biological languages with the statistical regularities found in human natural language, we embarked on a three-pronged approach:

i) We sourced 570,830 reviewed protein sequence entries from UniProt[1] (high-quality, Swiss-Prot database, https://www.uniprot.org/), incorporating 20 standard amino acids along with 5 nonstandard amino acids (denoted X, Z, B, U, O), to construct a corpus for biological language analysis, designated $C_{\alpha}$.

ii) We collected 72,753 high-quality AAV capsid sequences, each containing 7 amino acids, from open-source data[2]. These sequences were processed similarly to protein sequences to construct a corpus for AAV capsid sequence language analysis, designated $C_{\beta}$.

iii) We obtained raw data from the Book1 datasets (an open-sourced dataset)[3], and after a thorough data cleansing process (eliminating punctuation, spaces, digits, and special characters), we retained only the 52 alphabetic characters, thus forming a corpus for human natural language, designated $C_{\gamma}$.

From these corpora, we segment tokens on the basis of k-gram divisions, ranging from unigrams to pentagrams, guided by Zipf's Law, which postulates that the frequency $n_i$ of the $i^{th}$ most common word is as follows:

$$n_i = \frac{1}{i^{\alpha}}$$

namely,

$$\log n_i = -\alpha \log i + c$$

where $\alpha$ represents the distribution's exponent and where $c$ is a constant. Obviously, we observed that the frequency distribution of biological language adheres to:

$$\log n_i^{\alpha} = -\alpha^{\alpha} \log i + c^{\alpha}$$

in which the AAV capsid conforms to:

$$\log n_i^{\beta} = -\alpha^{\beta} \log i + c^{\beta}$$

in which human natural language conforms to the following:

$$\log n_i^{\gamma} = -\alpha^{\gamma} \log i + c^{\gamma}$$

where $\alpha^{\gamma} > \alpha^{\alpha} > \alpha^{\beta}$.

This finding is exciting for several reasons:

i) This finding demonstrates that not only does human natural language syntax adhere to Zipf's law but also biological languages (protein and AAV sequences) do so (Supplementary Fig. 1).

ii) The quantity of n-grams in the vocabularies of $C_{\alpha}$, $C_{\beta}$ and $C_{\gamma}$ is not excessively large, suggesting the presence of significant structural elements within languages.

These outcomes lay the groundwork for harnessing the capabilities of language models. At their core, language models are indispensable for the analysis of both natural and biological languages. In addition to Zipf's law, a previous study[4] has shown that natural amino acid sequences conform to a set of statistical patterns, thereby constituting a distinct biological lexicon. It is evident that the expertise and insights accrued over time in the field of natural language processing, especially those pertaining to language modeling and analytical methodologies[5], offer significant promise in this domain.

### 2. Sequence function evaluation model

According to the three target tasks (production fitness evaluation, ly6a target binding ability evaluation, and ly6c1 target binding ability evaluation), we divided the $\boldsymbol{D_{Production\ Fitness}}$, $\boldsymbol{D_{Ly6a}}$ and $\boldsymbol{D_{Ly6c1}}$ datasets into training and test sets at a ratio of 8:2, respectively, and adopted fivefold cross-validation. We established six evaluation indicators for these three tasks: RMSE, Pearson, Spearman, $R^2$, MedAE, and the C-index. Seven evaluation models were trained: support vector regression (SVR), eXtreme gradient boosting (XGBoost), k-nearest neighbors (KNN), random forest (RF), gradient boosting (GB), Bayesian ridge, and AdaBoost. The hyperparameters of these models can be found in Supplementary Table3. After the prediction performance of the seven candidate models on the three target tasks was analyzed, gradient boosting was selected as the optimal model because of its superior overall performance. The detailed performance data of each candidate model on the three tasks can be found in Tables S4--6. Each set of results was averaged, and the standard deviation was calculated after fivefold cross-validation.

### 3. Development of the ALICE system

#### 3.1 Pretraining Module

The pretraining module capitalizes on the encoder architecture of the transformer model[5], primarily because of the encoder's exceptional proficiency in encoding and compressing linguistic data, and encoder-only configurations (e.g., BERT and RoBERTa) have advantages in absorbing biosemantic and syntactic information in the pretraining stage[5], outperforming models based on the decoder-only or encoder-decoder paradigm. For example, the BERT framework adopts the bidirectional encoder of the transformer to demonstrate strong contextual understanding and higher data retention efficiency, as reflected in its proficiency in next sentence prediction and its ability to discern contextual relationships by strategically masking tokens. The effectiveness of these methods is particularly evident in settings with limited computing resources and restricted data access, which is attributed to their low training requirements.

##### 3.1.1 Illustration of the Pretraining Tasks

We consider two critical considerations: first, the ALICE system integrates an extensive array of complex data and models within its downstream processes, which necessitates the precise splicing and assembly of these components to ensure vocabulary uniformity; second, the range of amino acid sequences encountered during the upstream pretraining phase is highly variable. In contrast, the downstream fine-tuning phase is characterized by a fixed amino acid length of seven, coupled with a specific level of data complexity. To ensure vocabulary consistency across both the upstream and downstream tasks, a character-based tokenizer is used to segment each amino acid into individual characters, thereby creating an exhaustive vocabulary. This methodology promotes seamless integration and enhances the efficiency of managing the diverse datasets inherent to the system architecture.

We proceed to illustrate examples of pretraining tasks within the BERT[6] and RoBERTa[7] architecture training processes as follows:

- **Pretraining Process of the BERT Architecture**

Amino acid sequences can be analogously viewed as sentences within natural language. Adhering to the data construction protocol of BERT, the pretraining phase encompasses two pivotal tasks[6]: the masked language model (MLM) and next sentence prediction (NSP).

**Algorithm I: Pretraining** [BERT Model]

```
def BERT (input_ids, segment_ids, attention_mask, N_layers, head_size, num_heads):
    # Embedding Layers
    embeddings = WordEmbeddingLayer(input_ids) + PositionEmbeddingLayer((input_ids) + SegmentEmbeddingLayer(segment_ids)
    # Transformer Layers with Self-Attention
    for _ in range(N_layers):
        attention_output = MultiHeadAttention(embeddings, attention_mask, head_size, num_heads)
        embeddings = LayerNorm(embeddings + attention_output)
        ff_output = FeedForwardNetwork(embeddings)
        embeddings = LayerNorm(embeddings + ff_output)
return embeddings
```

**Algorithm II:** Pretraining [next sentence prediction (NSP) task]

```
def NextSentencePrediction(embeddings):
    # Linear Mapping hidden_size → 2
    mapping = Linear (embeddings)
    # Binary Classification of NSP
    return LogSoftmax(mapping)
```

**Algorithm III: Pretraining** [Predicting the origin token from the masked input sequence]

```
def MaskedLanguageModel(embeddings):
    # Linear Mapping hidden_size → vocab_size
    mapping = Linear (embeddings)
    # Predicting the origin token from the masked input sequence
    return LogSoftmax(mapping)
```

where $\mathbf{input}_{\mathbf{ids}}$ represents the sequence of word IDs derived from the input text, $\mathbf{segment}_{\mathbf{ids}}$ are utilized to distinguish between various segments of the sequence (e.g., Sentence A and Sentence B in sentence pair tasks), and $\mathbf{attention}_{\mathbf{mask}}$ provides the model with information regarding which tokens are actual words and which are padding, ensuring that the model processes only the relevant parts of the input sequence. $\mathbf{N_{layers}}$ refers to the number of transformer layers within the architecture, and $\mathbf{head_{size}}$ indicates the size of each attention head. Finally, $\mathbf{num_{heads}}$ denotes the number of attention heads used in the model, which is a key parameter that influences the model's capacity to capture different aspects of the input data through parallel processing.

**I. MLM and the masking procedure**

Consider an unlabeled amino acid sequence, "MRWQEMGYIFYPRKLR". During the random masking process, we select the i-th token (where $\mathrm{i} <$ is the length of the unlabeled sequence) and engage in a masking strategy where 15% of the amino acid "words" are subjected to masking, substitution, or retention. For illustrative purposes, the distribution of these modifications follows the ratio of mask:substitution: keep = 0.8:0.1:0.1, described as follows:

- 80% of the time: The word is replaced with a [MASK] token; for instance, "MRWQEMGYIFYPRKLR" becomes "MRW[MASK]EMGYIF[MASK]PR[MASK]LR".
- 10% of the time: The original word is substituted with a random amino acid, exemplifying that "MRWQEMGYIFYPRKLR" may be altered to "MRUQEQGYIFYLRKLR".
- 10% of the time: The original word is retained unchanged; hence, "MRWQEMGYIFYPRKLR" remains "MRWQEMGYIFYPRKLR". This approach biases the model toward learning authentic word representations.

The effectiveness of this approach is rooted in the transformer encoder's lack of prior knowledge regarding the specific words it will predict or those that are substituted randomly, which compels the model to develop a contextual representation for each input token. Moreover, since random substitutions affect only 1.5% of all tokens (10% of the 15% that are modified), this technique does not adversely affect the model's linguistic understanding.

**II. Next Sentence Prediction**

For the next sentence prediction task, each amino acid sequence is bisected, with each half either retaining its original context or being randomly paired. This task is exemplified as follows:

- Input = "MR[MASK]QEMG [SEP] YIFYPR[MASK]LR", Label = "IsNext".
- Input = "MR[MASK]QEMG [SEP] penguin Q[MASK]M[MASK]Y", Label = "NotNext".

In this context, the [SEP] token serves to delineate the boundary between distinct segments. This architectural choice is instrumental in enabling the model to discern contextual relationships among adjacent amino acid sequences, thereby bolstering its predictive precision and semantic comprehension within the realm of bioinformatics applications.

**III. Hyperparameter Optimization**

Following the determination of the model architectures, the subsequent step involves optimizing the hyperparameters for the two model architectures. Using the random search method, the learning rate was identified to be 1e-4, with warmup steps set at 10,000 and weight decay at 0.01. Additionally, $\beta_1$ and $\beta_2$ were set to 0.9 and 0.999, respectively. On the basis of these settings, a random search was conducted for parameters such as the number of layers, hidden size, dropout rate, and number of attention heads, as detailed in the table below. To ensure the robustness of our findings, each set of experimental results is repeated five times, allowing us to calculate the average and variance, thereby enhancing the reliability of our hyperparameter optimization process (Supplementary Table7).

- **Pretraining Process of the RoBERTa Architecture**

Building upon the BERT framework, the RoBERTa architecture introduces two significant technical advancements to enhance model performance[7]: dynamic masking and elimination of next-sentence prediction (NSP).

**Algorithm IV: Pretraining** [RoBERTa Model]

```
def RoBERTa (input_ids, segment_ids, attention_mask, N_layers, head_size, num_heads):
    # Embedding Layers
    embeddings = WordEmbeddingLayer(input_ids) + PositionEmbeddingLayer((input_ids) + SegmentEmbeddingLayer(segment_ids)
    # Transformer Layers with Self-Attention
    for _ in range(N_layers):
        attention_output = MultiHeadAttention(embeddings, attention_mask, head_size, num_heads)
        embeddings = LayerNorm(embeddings + attention_output)
        ff_output = FeedForwardNetwork(embeddings)
        embeddings = LayerNorm(embeddings + ff_output)
    return embeddings
```

**Algorithm V: Pretraining** [Predicting the origin token from the masked input sequence]

```
def MaskedLanguageModel(embeddings):
    # Linear Mapping hidden_size → vocab_size
    mapping = Linear (embeddings)
    # Predicting the origin token from the masked input sequence
    return LogSoftmax(mapping)
```

where $\mathbf{input}_{\mathbf{ids}}$ represents the sequence of word IDs for the input text, segment_ids are utilized to distinguish between different segments of the sequence (e.g., Sentence A and Sentence B in sentence pair tasks), and $\mathbf{attention}_{\mathbf{mask}}$ informs the model about which words are actual words and which ones are padding. $\mathbf{N}_{\mathbf{layers}}$ refers to the number of transformer layers, $\mathbf{head}_{\mathbf{size}}$ indicates the size of each attention head, and $\mathbf{num}_{\mathbf{heads}}$ represents the number of attention heads. RoBERTa and BERT are built upon a comparable foundational architecture and engage in analogous training tasks; however, they diverge in their pretraining methodologies. RoBERTa enhances training velocity and resource optimization by refining the selection of training data, modifying batch sizes, and implementing strategic learning rate schedules. These refinements contribute to a more efficient and effective pretraining phase, distinguishing RoBERTa from BERT in terms of performance and scalability.

**I. Dynamic Masking**

Diverging from BERT's static masking protocol, RoBERTa implements a dynamic masking strategy. This approach involves (1) decuplicating the original corpus, with each instance undergoing random masking of 15% of tokens, and (2) maintaining the predetermined distribution of masking (80%), replacement (10%), and retention (10%) activities. Analogous to cross-validation methodologies, this dynamic masking generates distinct masked variations within each corpus iteration, broadening the model's exposure to diverse information and fostering more comprehensive learning.

**II. Omission of NSP**

RoBERTa eliminates the NSP component, following insights from SpanBERT[8], which highlights potential drawbacks associated with NSP, including noise introduction into the masked language modeling (MLM) task. This decision addresses two primary concerns: (1) sentence concatenation potentially leads to truncation and diminishes the model's ability to learn from longer sequences crucial for predicting masked tokens; and (2) irrelevant negative samples in NSP detracting from accurate mask prediction within the current sentence. Consequently, removing NSP aims to enhance the model's focus on MLM, potentially improving performance by reducing irrelevant noise and aligning more closely with the objective of understanding complex linguistic and biological sequences.

**III. Hyperparameter Optimization**

RoBERTa's architecture closely resembles that of BERT, albeit with distinctions in training data processing and methods. Thus, we adhere to a consistent configuration: setting epochs to 4, learning rate to 1e-4, warming up with 10,000 steps, employing a weight decay of 0.01, and using $\boldsymbol{\beta_1}$=0.9 and $\boldsymbol{\beta_2}$=0.999. Furthermore, we employ the random search technique to explore hyperparameters and refine the model architecture holistically (Supplementary Table8). Specifically, RoBERTa is not a new pretraining model but rather an optimization method tailored for BERT, similar to the techniques outlined in the original paper. Therefore, at the model level, RoBERTa is essentially BERT, but it is a high-performance version achieved through a series of carefully crafted optimizations.

#### 5.2 Semantic Tuning module

In biology, the amino acid sequence distribution patterns of viruses differ from the discrete distributions typically found in most protein sequences. Instead, they resemble a dialect within the protein language family. While models in the pretraining phase learn semantic and syntactic information from the vast UniProt protein database, generating accurate and high-quality authentic viral capsid sequences of AAV requires a deeper understanding of the "language" and distribution rules of viruses through precise learning methods.

To optimize training data quality and process effectiveness, we refined our dataset by selecting viable AAV capsid sequences from a constructed library, carefully filtering out capsid sequences with weak Ly6a or Ly6c1 target binding and weak virus production ability. This process yielded 72,753 viable AAV capsid sequences for the generative model's semantic tuning stage. As in the pretraining stage, all sequences underwent char-level tokenization, with a consistent vocabulary established. To leverage the biological semantic and syntactic information embedded in the pretrained RoBERTa model, we implemented a semantic tuning strategy, enriching our comprehension of AAV semantics and programming aspects on the basis of previously acquired protein knowledge.

#### 3.2.1 Capsid generation module architecture

The ALICE framework employs a SeqGAN architecture for its generative model to produce high-quality, viable, and sufficiently diverse AAV sequences. During semantic tuning, pretrained BERT and RoBERTa models serve as the SeqGAN generator's core, with a bespoke output layer for sequence generation. Using AAV capsid sequences from the library as true samples, the model iteratively samples signals via Gaussian noise. The generator aims to produce sequences closely resembling true samples, whereas the discriminator strives to accurately differentiate between positive and negative samples. This adversarial process enables the generator to create samples that mimic real viral sequences, avoiding discriminator detection, whereas the discriminator refines its ability to identify increasingly authentic-looking fake samples.

Through this iterative process, the semantic and syntactic information within viral sequences is precisely recognized and compressed into the SeqGAN model architecture via adversarial learning. Notably, semantic and syntactic information of biological languages is crucial: training directly on a SeqGAN architecture with initialized hyperparameters without a pretraining phase generates sequences that bind weakerly to the Ly6a and Ly6c1 targets than sequences generated by a SeqGAN using a pretrained RoBERTa encoder architecture as the generator core.

#### 3.2.2 Architecture of the Semantic Tuning Module

**Algorithm VI: Semantic Tuning** [Generator with Masked Language Model]

```
def SemanticTuning_Generator(sequence):
    # Features learned from the pretraining stage
    mlm_ebed = RoBERTa (sequence)
    # Mapping
    embed = Linear(mlm_ebed)
    # LSTM Generate Sequences to Learn the Semantic of AAV Capsid
    output = LSTM(embed)
    hw = Linear (output)
    # Applied Highway Linear with Gate Control
    features = Dropout(Sigmoid(hw)×ReLU(hw)+(1-Sigmoid(hw))×output)
    return GumbelSoftmax(Linear(features))
```

**Algorithm VII: Semantic Tuning** [Discriminator]

```
def SemanticTuning_Discriminator(sequence):
    embed = Linear(seq)
    cnn_lst = []       # ReLU+CNN1d
    for _ in N_CNN_layers:
        cnn_lst.append(ReLU(Conv1d(embed)))
    # Splice pools at dim=1 - splice each batch together
    pred = concatenate(cnn_lst, dim=1)
    for _ in Res_block_layers:
        pred = ResBlock(pred)
        pred = LayerNorm(pred)
    hw = Linear(pred)
    # Applied Highway Linear with Gate Control
    pred = Sigmoid(hw)×ReLU(hw) + (1-Sigmoid(hw))×pred
    pred = Linear(pred)
    return Sigmoid(Linear(Dropout(pred)))
```

**Algorithm VIII: Semantic Tuning** [GumbelSoftmax]

```
def GumbelSoftmax (logits, temperature):
    # Step 1: Sample from Gumbel(0, 1)
    def sample_gumbel(shape):
        U = Uniform(0, 1, shape)
        return -Log(-Log(U))
    # Step 2: Add Gumbel noise to the logits
    gumbel_noise = sample_gumbel(logits.shape)
    noisy_logits = logits + gumbel_noise
    # Step 3: Apply Softmax with temperature
    Softmax_output = Softmax(noisy_logits/temperature)
    return Softmax_output
```

**Algorithm IX: Semantic Tuning [ResBlock]**

```
def ResBlock(input):
    x = Conv1d(ReLU(input))
    output = Conv1d(ReLU(x))
    return input + 0.3×output
```

In our model, $\mathbf{sequence}$ refers to authentic AAV sequences derived from the library. $\mathbf{mlm_{embed}}$ denotes embedding features obtained by pretraining input sequences via the RoBERTa model, which is employed for semantic adjustment within the generator. $\mathbf{cnn_{lst}}$ represents a list of embedding features processed through ReLU activation functions and 1D convolutional layers, which serve for sequence feature extraction within the discriminator.

Within the Gumbel–Softmax[9] sampling framework, $\mathbf{logits}$ refers to unnormalized probability values serving as inputs for generating pseudodiscrete samples. $\mathbf{Temperature}$ signifies the parameter regulating the smoothness of the generated distribution in the Gumbel-Softmax function. $\mathbf{Gumbel_{noise}}$ denotes the sampled noise from the Gumbel distribution, which perturbs the logits. $\mathbf{Noisy_{logits}}$ refers to logits augmented with Gumbel noise and utilized for pseudodiscrete sample generation.

The Softmax function, which is applied to noisy logits and scaled by the temperature parameter, controls the distribution's smoothness. Lower temperatures produce distributions closer to one-hot vectors. The Gumbel-Softmax function returns samples approximating one-hot vectors while remaining differentiable, making it suitable for neural network training involving discrete choices.

#### 3.3 Ranking Filtration Process

To quantitatively assess the quality of capsid sequences generated during the Semantic Tuning stage and to identify the top 1000 sequences with the most robust overall capabilities, we employed the GB model to predict and evaluate the production fitness and binding abilities to Ly6a and Ly6c1 of the generated capsid sequences.

For the selection process, we defined the RankFiltPro score to select the top 1000 candidates on the basis of multiple functions, including the binding capabilities to Ly6a and Ly6c1 and production fitness, via the following function:

$$\mathrm{RankFiltPro_{Score}} \propto Pred_{Ly6c1} + Pred_{Ly6a} + Pred_{ProductionFitness}$$

where $Pred_{Ly6c1}$ and $Pred_{Ly6a}$ represent the predicted binding abilities of the capsid sequences to these receptors and where $Pred_{ProductionFitness}$ represents the predicted production fitness, all of which are evaluated by the GB model. This score is used to rank and select the top 1000 candidates from the capsid sequences generated during the semantic tuning stage.

#### 3.4 Function-guided evolution

##### 3.4.1. MSA methods for constructing the FE function

We conceptualize the sequence optimization process in the function-guided evolution (FE) algorithm as a combinatorial optimization problem. The algorithm builds learning and evolution guidance around three functional optimization goals: production fitness, Ly6a binding affinity, and Ly6c1 binding affinity. It promotes iterative optimization through mutations within the AAV capsid "population", aiming to design sequences closer to more functional AAV variants ($\boldsymbol{Ref^{+}}$) and divergent from less functional variants ($\boldsymbol{Ref^{-}}$).

To simulate AAV evolution, we selected several heuristic algorithms: a genetic algorithm (GA)[10], simulated annealing (SA)[11], a distribution estimation algorithm (EDA)[12], and a hybrid of a distribution estimation algorithm and a genetic algorithm (EDG)[12].

We designed an FE function to quantitatively evaluate the differences between AAV sequences during optimization and between $\boldsymbol{Ref^{+}}$ and $\boldsymbol{Ref^{-}}$. To ensure a reasonable and objective evaluation of the degree of optimization, we incorporated the concept of multiple sequence alignment (MSA).

The FE function is defined as follows:

$$\mathbf{max}: \mathbf{Target}(\mathbf{Seq}, \mathbf{Ref^{+}}, \mathbf{Ref^{-}}) \propto \mathbf{MSA}(\mathbf{Seq}, \mathbf{Ref^{+}}) - \mathbf{MSA}(\mathbf{Seq}, \mathbf{Ref^{-}})$$

where $\mathrm{MSA}$ indicates the Smith–Waterman algorithm [13], which indicates each sequence to be evaluated with the optimization function.

$$\mathrm{MSA}_{i,j} = \max \begin{cases} H_{i-1,j-1} + s(a_i, b_j), \\ \max_{k\geq 1}\{H_{i-k,j} - W_k\} \\ \max_{l\geq 1}\{H_{i,j-l} - W_l\} \\ 0 \end{cases}$$

where $1 \leq i \leq n, 1 \leq j \leq m$. The detailed pseudocode is as follows:

**Algorithm X: Smith–Waterman Algorithm**

**def MSA** (sequence_a, sequence_b):

Initialization: $\text{Set } H_{k,0} = 0 \text{ for } 0 \leq k \leq n \text{ and } H_{k,1} = 0 \text{ for } 0 \leq l \leq m$

$\text{for } i = 1,2,\ldots,m \text{ do:}$

$$H_{i,j} = \max \begin{cases} 0 \\ H_{i-1,j-1} + s(a_i, b_j) \\ H_{i-k,j} - (G_s + kG_e) \\ H_{i,j-1} - (G_s + lG_e) \end{cases}$$

$\text{set backtrace } T(i,j) \text{to the maximizing pair } (i,j)$

$\text{set } (i,j) := \operatorname{argmax}\{H_{i,j} | i = 1,2,\ldots,n, j = 1,2,\ldots,m\}$

$\text{the best score is } \alpha := H_{i,j} \text{ repeat}$

$\text{if } T(i,j)(i-1.j-1):$

$\quad \text{print}(x_{i-1}, y_{j-1})$

$\text{elif } T(i,j)(i-1.j-1):$

$\quad \text{print}(x_{i-1}, -)$

```
        else:
            print(−, y_{i−1})
            set (i, j) ≔ T(i, j)
            until H_{i,j} = 0
    return score α
```

#### 3.4.2. Heuristic Algorithms for AAV Sequence Optimization

Unlike deep learning algorithms, which typically require large datasets, heuristic algorithms can yield meaningful results with only a few hundred data points, a feature particularly advantageous when dealing with scarce experimental data such as AAV. In this study, we established the top 1000 functional sequences filtered by the ranking filtration process module (Fig. 1c) as the initial population. We explored four heuristic algorithms to investigate their specific impacts on the functional evolution of AAV sequences:

- **Genetic algorithms (GAs)[10]:** GAs simulate natural selection, making them well suited for optimizing discrete data such as amino acid sequences. Through iterative selection, crossover, and mutation, GAs explore a broad search space, facilitating the discovery of highly optimized sequences.
- **Simulated Annealing (SA)[11]:** Inspired by metallurgical annealing, SA excels in escaping local optima by occasionally accepting suboptimal solutions. This approach is effective for continuous attribute optimization, enabling fine-tuning of continuous parameters in AAV sequences.
- **Estimation of distribution algorithms (EDAs)[12]:** EDAs construct probabilistic models of promising solutions to guide the search process. This approach is particularly advantageous for discrete data, where modeling the distribution of amino acid sequences can lead to effective optimization.
- **Estimation of the distribution generic algorithm (EDG)[12]:** EDG combines elements of GAs and EDAs, leveraging both natural selection and probabilistic modeling. This hybrid approach enhances the optimization process by balancing exploration and exploitation.

#### 3.4.3 Genetic Algorithm (GA) for Sequence Optimization

In the genetic algorithm (GA) for sequence optimization, the evolution process commences with crossover, mutation, and selection strategies to simulate evolutionary progress. By applying multiple sequence alignment (MSA) search in the contrastive purpose function, we align the optimized sequences' divergence distance after each evolution epoch between the sets of $\boldsymbol{Ref}^{+}$ and $\boldsymbol{Ref}^{-}$. This approach enables effective evolution with limited data and a clear optimization purpose.

**Algorithm XI: Genetic Algorithm**

```
def GA(candidate_sequences):
    # Top 1000 functional sequences filtered out via the ranking filtration process
    def purposeFunction(seq):
        # Calculate the MSA score between seq and Ref+ and the negative MSA score between
    seq andRef−
```

```
        return min(MSA(Ref, Ref^+) − MSA(Ref, Ref^−))
    while purposeFunction(candidate_sequences) > stopping_threshold:
        new_seqs = []
        evolve_score = [(seq, purposeFunction(seq)) for seq in candidate_sequences]
        # Descend rank through contrastive score
        selected_score = evolve_score.sort(key=lambda x: x[1])
        # Select the top n number sequences as the most adaptable sequences in the current iteration
        selected_seqs = [seq for seq, score in selected_score[: topn]]
        # Apply crossover on optimized sequences
        new_seqs.append(crossover(selected_seqs))
        # Apply mutation to offspring with a predefined mutation rate
        new_seqs.append(mutate(selected_seqs))
        n_evolve_score = [(seq, purposeFunction(seq)) for seq in new_seqs]
        # Descend rank through contrastive score
        n_selected_score = n_evolve_score.sort(key=lambda x: x[1])
        # Select the top n number sequences as the most adaptable sequences in the current iteration
        n_selected_seqs = [seq for seq, score in n_selected_score[: topn]]
        # Replace the least fit sequences in P with new offspring
        candidate_sequences = selected_seqs + n_selected_seqs[:P-topn]
    return the sequence with the highest purpose score as the optimized AAV variant
```

Note:

- Purpose Function: We define an $\boldsymbol{fitness\ function}$ that evaluates each sequence on the basis of its similarity to $\boldsymbol{Ref^{+}}$ and dissimilarity to $\boldsymbol{Ref^{-}}$. This involves calculating the MSA score.
- Selection: Sequences are selected on the basis of their fitness scores, with higher fitness conferring a greater probability of selection for the next generation, mimicking natural selection.
- Crossover: Selected sequences are paired and undergo crossover to create new offspring. This process involves exchanging sequence segments to combine desirable traits from parent sequences.
- Mutation: Random mutations are applied to the offspring at a predefined rate, introducing genetic diversity and facilitating exploration of the search space.

#### 3.4.4 Simulated Annealing Algorithm (SA) for Sequence Optimization

Next, in addition to the genetic algorithm (GA), we implement the simulated annealing (SA) algorithm as a fundamental component of AAV sequence optimization. SA, a versatile heuristic algorithm inspired by metallurgical annealing processes, has been explored in this context. Our methodological examination encompasses the underlying principles and techniques of SA, its unique advantages in AAV sequence optimization, and its contribution to enhancing AAV sequences for specific biomedical applications.

The optimization process begins with iterative annealing steps, where sequences undergo perturbation and evaluation on the basis of a $\boldsymbol{fitness\ function}$. By applying the MSA search within the contrastive purpose function, we align the optimized sequences' divergence distance after each annealing step between the sets of $\boldsymbol{Ref^+}$ and $\boldsymbol{Ref^-}$. This approach enables effective optimization with limited data and a clear optimization purpose.

**Algorithm XII: Simulation Annealing Algorithm**

```
def SA (candidate_sequences):
    # Top 1000 functional sequences filtered out via the ranking filtration process
    population = candidate_sequences
    T = initial_temperature
    alpha = cooling_rate
    def purposeFunction(seq):
        # Calculate the MSA score between seq and Ref+ and the negative MSA score between seq andRef−
        return min(MSA(Ref, Ref+) − MSA(Ref, Ref−))
    while T > stopping_threshold:
        new_population = []
        for seq in population:
            # Generate a new sequence seq' by perturbing seq
            new_seq = perturb(seq)
            # Calculate the fitness of seq and new_seq
            current_fitness = purposeFunction(seq)
            new_fitness = purposeFunction(new_seq)
            # Accept new_seq on the basis of the SA acceptance probability
            if new_fitness > current_fitness or random.uniform(0, 1) < math.exp((new_fitness - current_fitness)/T):
                new_population.append(new_seq)
            else:
                new_population.append(seq)
        # Reduce the temperature
        T *= alpha
        # Update the population with the new sequences
        population = new_population
        # Align optimized sequences via MSA to measure the divergence from Ref+ andRef−
    # Return the sequence with the highest fitness score as the optimized AAV variant
    best_sequence = max(population, key=purposeFunction)
    return best_sequence
```

Note:

The initial temperature $\mathrm{T}$ and the cooling rate $\alpha$ are defined.

Annealing process:

For each sequence $\mathrm{seq}$ in the population:

- Generate a new sequence $\mathrm{seq}'$ by making a small perturbation to $\mathrm{seq}$.
- The fitness of both $\mathrm{seq}$ and $\mathrm{seq}'$ is calculated via $\boldsymbol{fitness\ function}$.
- If $\mathrm{F(seq')}$ is greater than $\mathrm{F(seq)}$, accept $\mathrm{seq}'$ as the new sequence.

- If $F(seq')$ is less than or equal to $F(seq)$, accept a probability $e^{\frac{F(seq')-F(seq)}{T}}$.

Temperature reduction: Reduce the temperature $T$ by multiplying it by the cooling rate $\alpha$.

#### 3.4.5 Estimation of the Distribution Algorithm (EDA) for Sequence Optimization

The estimation of distribution algorithms (EDAs) represents a class of evolutionary algorithms that optimize a population $(P)$ of candidate solutions. The process begins with the generation and evaluation of an initial population $P$ via an objective function. The best-performing candidates are then selected to construct a probabilistic model of $P$, from which a new set of points is sampled. This iterative process continues until a termination criterion is met.

The key distinction between EDAs and conventional evolutionary algorithms lies in their method of generating new candidate solutions. While traditional evolutionary algorithms employ implicit distributions defined by variation operators, EDAs utilize explicit probability distributions encoded by models such as Bayesian networks or multivariate normal distributions. Like other evolutionary algorithms, EDAs can address optimization problems across various representations, from vectors to LISP-style S-expressions, with candidate solutions evaluated via one or more objective functions.

By leveraging explicit probabilistic models, EDAs effectively address optimization problems that challenge conventional evolutionary algorithms and traditional optimization techniques, particularly those with high levels of epistasis. Furthermore, EDAs provide valuable insights into the problem being solved through their probabilistic models. This information can be utilized to design problem-specific neighborhood operators for local search, bias future runs of EDAs on similar problems, or create efficient computational models of the problem.

The EDA approach is predicated on evolving a probabilistic model of the solution space. The potential solutions in $P$ are treated as realizations of multivariate random variables, whose joint probability distribution is estimated and updated iteratively. Under the assumption of independent variables, the product of their marginal distributions constitutes the joint distribution of all variables, which is particularly effective in handling limited data for evolutionary progress.

The general procedure of an EDA is as follows:

```
t := 0
initialize model M(0) to represent a uniform distribution over admissible solutions
while (termination criteria not met) do
    P := generate N>0 candidate solutions by sampling M(t)
    F := evaluate all candidate solutions in P
    M(t + 1) := adjust_model(P, F, M(t))
    t := t +
```

**Algorithm XIII: Estimation of the distribution algorithm**

**def EDA**(candidate_sequences):

    # Top 1000 functional sequences filtered out via the ranking filtration process
    population = candidate_sequences
    **def** purposeFunction(seq):
        # Calculate the MSA score between seq and $\boldsymbol{Ref}^{+}$ and the negative MSA score between seq and $\boldsymbol{Ref}^{-}$

```
        return min(MSA(Ref, Ref+) − MSA(Ref, Ref−))
    # Initialize model M(0) to represent uniform distribution over admissible solutions
    model = initialize_uniform_model(population)
    t = 0
    # Determine if the stopping criterion is met
    while not stopping_criterion_met(t):
        # Generate N > 0 candidate solutions by sampling M(t)
        sampled_population = sample_from_model(model, population_size=1000)
        # Evaluate all candidate solutions in P
        fitness_scores = [(seq, purposeFunction(seq)) for seq in sampled_population]
        # Adjust model M(t + 1) on the basis of population P and fitness scores F
        model = adjust_model(sampled_population, fitness_scores, model)
        t += 1
    # Return the sequence with the highest fitness score as the optimized AAV variant
    best_sequence = max(sampled_population, key=purposeFunction)
    return best_sequence
```

Note:

- Initialization: Initialize the model $\mathrm{M}(0)$ to represent a uniform distribution over the admissible solutions.
- $\boldsymbol{fitness\,function}$: Define a $\boldsymbol{fitness\,function}$ that evaluates each sequence on the basis of its similarity to $\boldsymbol{Ref}^{+}$ and dissimilarity to $\boldsymbol{Ref}^{-}$. This involves calculating the multiple sequence alignment (MSA) score.
- Sampling: Generate more than zero candidate solutions by sampling from the model $\mathrm{M(t)}$.
- Evaluation: Evaluate all candidate solutions using the$\boldsymbol{fitness\,function}$
- Model Adjustment: Adjust the probabilistic model $\mathrm{M(t+1)}$ on the basis of the evaluated population and their fitness scores.
- Alignment and Measurement: After each iteration, align the optimized sequences via MSA to measure their divergence distance from $\boldsymbol{Ref}^{+}$ and $\boldsymbol{Ref}^{-}$. This ensures that the optimization process is guided by clear objectives and maintains the semantic integrity of the sequences.

#### 3.4.6 Mixture of the estimation of distribution algorithm and the genetic algorithm (EDA-GA, also referred to here as the EDG) method

The mixture of the estimation of distribution algorithm and the genetic algorithm (EDG) method represents an innovative and powerful approach in AAV sequence optimization. EDG combines the strengths of both the genetic algorithm (GA) and estimation of distribution algorithm (EDA) to create a hybrid approach that excels in addressing the intricate challenges posed by AAV sequences.

This hybrid algorithm aims to achieve superior AAV function design performance by combining the rapid convergence characteristic of EDAs with the broad search capabilities of GAs. By leveraging the complementary strengths of these two algorithms, EDG offers a robust and efficient method for exploring the complex solution space of AAV sequences, potentially leading to more effective and targeted optimization outcomes.

**Algorithm XIV: Estimation of the distribution algorithm and genetic algorithm**

```
def EDA_GA(candidate_sequences, max_generations, population_size):
    # Top 1000 functional sequences filtered out via the ranking filtration process
    population = initialize_population(candidate_sequences, population_size)
    def fitness_function(seq):
        # Calculate the fitness of the sequence on the basis of the task scheduling criteria
        return evaluate_fitness(seq)
    # Initialize model M(0) to represent a uniform distribution over admissible solutions
    model = initialize_uniform_model(population)
    t = 0
    while t < max_generations:
        # EDA Part
        # Select top-performing individuals on the basis of fitness
        selected_for_eda = select_top_individuals(population, fitness_function, selection_size)
        # Build probabilistic model from selected individuals
        model = build_probabilistic_model(selected_for_eda)
        # Sample new candidates from the probabilistic model
        eda_offspring = sample_from_model(model, population_size//2)
        # GA Part
        # Select top-performing individuals on the basis of fitness
        selected_for_ga = select_top_individuals(population, fitness_function, selection_size)
        # Apply crossover and mutation to generate new candidates
        ga_offspring = []
        for _ in range(population_size//2):
            # Select two parents from the population for crossover
            parent1, parent2 = select_parents(selected_for_ga)
            child = crossover(parent1, parent2)
            mutated_child = mutate(child)
            ga_offspring.append(mutated_child)
        # Combine offspring from EDA and GA
        combined_offspring = eda_offspring + ga_offspring
        # Evaluate fitness of all offspring
        offspring_fitness = [(seq, fitness_function(seq)) for seq in combined_offspring]
        # Select the best individuals from the combined offspring and the original population
        population = select_best_individuals(population, combined_offspring, offspring_fitness, population_size)
        t += 1
    # Return the sequence with the highest fitness score as the optimized solution
    best_sequence = max(population, key=fitness_function)
    return best_sequence
```

**Algorithm XV: Estimation of** the distribution algorithm and genetic algorithm [select the top individuals]

```
def select_top_individuals(population, fitness_function, selection_size):
    # Select the top-performing individuals on the basis of fitness
    sorted_population = sorted(population, key=fitness_function, reverse=True)
    return sorted_population[:selection_size]
```

**Algorithm XVI: Estimation of** the distribution algorithm and genetic algorithm [select the best individuals]

```
def select_best_individuals(population, offspring, offspring_fitness, population_size):
    # Select the best individuals from the combined offspring and original population
    combined_population = population + offspring
    sorted_population = sorted(combined_population, key=lambda x: offspring_fitness[x],
reverse=True)
    return sorted_population[:population_size]
```

**Note:**

EDA Model Building and Sampling:

- The top-performing individuals are selected on the basis of fitness.
- A probabilistic model is built from these selected individuals.
- Sample new candidate solutions from this probabilistic model to create EDA-evolved offspring.

GA operations:

- The top-performing individuals are selected from the population on the basis of fitness.
- Crossover and mutation are applied to these selected individuals to generate GA-evolved offspring.

Combining offspring:

- The EDA-evolved and GA-evolved offspring are combined.
- The best individuals from the combined pool and the original population are selected to form the new population.

#### 3.5 Hits-Ranking Module

We use the Hits-Ranking (HR) module to evaluate the sequences optimized via function-guided evolution (FE), selecting the most functional sequences for final in vivo wet experiment verification. The implementation method is as follows:

First, we used the FE score to screen the top 500 sequences. Next, we narrowed this down to the top 100 sequences on the basis of their virus production capacity (production fitness). Finally, we considered both the HR score and multitarget binding ability (Ly6a and Ly6c1) to select the top 8 sequences for functional verification through wet experiments.

### 4. KL-based regularization for optimizing the objective function to promote diversification in capsid sequence design.

To implement novelty-guided regularization, we employed a KL-based regularization procedure within the original objective loss in the FE module. For each generated sequence $s$, a position-specific one-hot distribution $Q \in \mathbb{R}^{L\times 20}$ was constructed with additive smoothing $\varepsilon$. The positive reference distribution $Ref^+$ was obtained from position-wise amino acid frequencies of the training positive set, softened by a temperature parameter $\tau$:

$$Ref^+_{i,j} \propto (f_{i,j} + \varepsilon)^{1/\tau}, \quad \sum_{j=1}^{20} Ref^+_{i,j} = 1,$$

where $f_{i,j}$ denotes the observed frequency of residue $j$ at position $i$.

The novelty-control component was formulated as the KL divergence between the generated distribution and the reference:

$$\mathrm{KL}(Q \parallel Ref^+) = \sum_{i=1}^{L}\sum_{j=1}^{20} Q_{i,j} \log \frac{Q_{i,j}}{Ref^+_{i,j}}.$$

To ensure comparability across sequence lengths, the divergence was normalized by the theoretical maximum per-position divergence, $L \cdot \ln(20)$, and clipped to the interval $[0,1]$:

$$\mathrm{KL}_{\mathrm{norm}}(Q, Ref^+) = \min\left(1, \max\left(0, \frac{\mathrm{KL}(Q \parallel Ref^+)}{L\ln(20)}\right)\right).$$

The final loss function under scheme 1 was expressed as

$$\mathcal{L}(s) = \mathcal{L}_{\mathrm{basic}} - w_{\mathrm{novel}} \cdot \mathrm{KL}_{\mathrm{norm}}(Q, Ref^+),$$

where $\mathcal{L}_{\mathrm{basic}}$ denotes the original objective loss in the FE module, and $w_{\mathrm{novel}}$ is the weight controlling the contribution of the novelty term. This formulation allowed systematic modulation of the balance between functional fidelity and exploratory novelty, with weights set to 0.2, 1.1, and 2.0 in the experiments corresponding to Supplementary Fig. 23 a–f.

At lower weights, KL-regularized optimization allowed FE to achieve functional integration but generated sequences that remained close to the $Ref^+$ set. In contrast, at higher weights, KL regularization drove FE to produce sequences more distinct from $Ref^+$, thereby increasing diversity, though at the expense of multifunctional integration.

### 5. Data preprocessing protocol and development of ALICE-X system

#### 5.1 Data Preprocessing and Evaluation Model Training

From the AAV-588-OligoPool-lib1 dataset, rows containing non-missing values for both production fitness and hTfR1 $\log_2$ enrichment were retained after data cleaning and filtering. Following the workflow described previously, we employed a gradient boosting regressor (GBR) as the evaluation model to ensure consistency across analyses. The $D_{hTfR1}$ and $D_{fitness-x}$ datasets, built from cleaned data, were used to train and evaluate models for hTfR1 log2 enrichment and production fitness prediction.

The dataset was randomly partitioned into a training set (n = 6,131) and a test set (n = 256). Two independent GBR models were then trained: one to predict production fitness, and the other to predict hTfR1 log2 enrichment. These regressors serve as evaluation components in subsequent modules of the ALICE-X framework, providing predictive guidance for sequence prioritization and optimization.

#### 5.2 Pre-training Stage

The pre-training stage leveraged the existing RoBERTa model as implemented in the ALICE framework. The model's parameter set remained unchanged, aligning with the configurations detailed in the preceding sections to ensure experimental consistency.

#### 5.3 Semantic-tuning Stage

##### 5.3.1 Semantic-tuning dataset $D_{Seq-x}$

We curated capsid sequences from the oligo dataset by selecting variants with comparatively favorable receptor engagement and packaging profiles using two relative (relative high fitness and hTfR1) log2 enrichment proxies: hTfR1 log2 enrichment > −2 and production fitness > −1. This filter yielded 1,388 unique capsid sequences.

For semantic tuning, we utilized a SecGAN model based on the SeqGAN framework, employing a RoBERTa-based pretraining initializer. The model was trained with a learning rate of $2 \times 10^{-6}$ and a BERT hidden dimension of 2048. The loss function incorporated weighted components ($\lambda_{mle}$ = 1.0, λ_adv = 0.25, λ_ent = 0.01), and the Gumbel-Softmax temperature was annealed from 1.5 to 0.7 (γ = 0.97). The model's capacity was 860,054 parameters for the generator and 1,435,530 for the discriminator. The learning rates for both networks were maintained at $1.0 \times 10^{-6}$, with a final Gumbel-Softmax temperature (τ) of 0.841.

##### 5.3.2 Semantic-tuning Training Process

During the Semantic Tuning stage, we continued to adopt a SeqGAN-based architecture to guide sequence generation. The training procedure, optimization strategy, and hyperparameter settings were kept consistent with those applied in the ALICE framework, ensuring methodological continuity and fair comparability across modules.

#### 5.4 Data processing and development of FE-X module

##### 5.4.1 Reference datasets construction for FE-X

Reference sets of capsid sequences were derived from the AAV-588-OligoPool-lib1 dataset to provide positive and negative reference sequences for the FE-X architecture. Specifically, 27 high-quality sequences were selected as the positive reference set ($Ref^+$), defined by stringent thresholds of production fitness greater than 2 and hTfR1 log2 enrichment greater than 2. To construct the negative reference set ($Ref^-$), 48 sequences with production fitness below −2 and hTfR1 log2 enrichment below −2 were randomly sampled. Random sampling was intentionally applied to mitigate potential class imbalance and to ensure that the negative set did not overly concentrate on any specific motif family. Together, these curated $Ref^+$ and $Ref^-$ datasets serve as the foundation for contrastive learning in Function Integration Stage and as anchor points for distance-based regularization in Exploration Stage.

#### 5.4.2 Designing Exploration Rewards for Novel Landscapes in FE-X

Direct modification of the objective function in the original FE module proved insufficient to achieve our dual goals (Supplementary Fig. 23): the discovery of novel sequence motifs and the simultaneous integration of multiple functional traits. This limitation is unsurprising, since a single objective function often induces strong biases toward local optima, leading the evolutionary process to converge prematurely on familiar motifs rather than exploring new regions of sequence space. To address this challenge, we reasoned that more sophisticated machine learning strategies were required—methods capable of balancing *exploitation* (retaining functional characteristics of known beneficial motifs) with *exploration* (venturing into unexplored but potentially valuable areas of the landscape).

To this end, we drew inspiration from the paradigm of curriculum learning, in which training is structured in stages of gradually increasing difficulty or abstraction. Following this principle, we restructured the functional evolution (FE) pipeline into a two-stage sequential framework, which we designate the FE-X module. This upgraded design explicitly separates the tasks of functional integration and novelty exploration, thereby enabling a more balanced optimization strategy.

1) **Function Integration Stage: Multi-functional Fusion.**

The first stage focuses on ensuring that candidate capsid sequences integrate two essential properties: production fitness and hTfR1 binding capability. In the original FE module, this objective was pursued through contrastive evolution, using $Ref^+$ sequences as positive exemplars and $Ref^-$ sequences as negative controls. This contrastive design encourages candidates to resemble beneficial motifs while avoiding deleterious patterns. Function Integration Stage in FE-X preserves this principle, acting as a functional filter to ensure that early-generation sequences are viable and multifunctionally competent.

2) **Exploration Stage: Exploration and diversification.**

By contrast, the second stage is explicitly designed to escape the local attractor basin of the reference sequences and to promote exploration of novel sequence motifs. The central aim of Exploration Stage is not only to generalize beyond the training distribution but also to ensure that novel capsid variants continue to support robust multi-functional integration. In this sense, Exploration Stage extends the curriculum: candidates that have passed the basic functional criteria of Function Integration Stage are now subjected to more demanding constraints that reward novelty, diversity, and balance across objectives. This design specifically addresses the well-known limitations of KL-based regularization, which often fails to achieve an effective trade-off between functional preservation and exploration, particularly in high-dimensional and sparsely sampled sequence landscapes.

**I. Composite Loss in Exploration Stage:**

To operationalize this exploration phase, exploration stage introduces a composite scoring function that integrates multiple complementary terms:

1) Predictive quality signals, derived from pretrained machine-learning models that independently estimate production fitness and hTfR1 binding capacity.

2) Exploration-oriented novelty measures, designed to quantify the dissimilarity of candidates relative to known reference sequences ($Ref^{+}/Ref^{-}$) and to prior generations.

3) Constraint and penalty terms, which act as regularizers, ensuring that the optimization process does not collapse into trivial, imitative, or non-viable solutions.

The resulting scoring function $S(x)$ is explicitly decomposable into interpretable components (Fig. 4c-f, Supplementary Fig. 25), making it possible to analyze the balance between exploitation and exploration achieved during optimization.

**II. Predictive Quality Components**

Let each candidate sequence $x$ be evaluated by two evaluation models constructed above:

1) $\hat{f}(x) \in \mathbb{R}$: predicted production fitness.

2) $\hat{h}(x) \in \mathbb{R}$: predicted hTfR1 binding capability.

Because the two predictive heads may operate on different scales, and because their outputs may contain heterogeneous noise distributions, we apply a standardization and calibration step before integrating them into downstream objectives. Several normalization strategies are available:

**1) Score normalization (z-score or robust min–max):**

$$z_f(x) = \frac{\hat{f}(x) - \mu_f}{\sigma_f}, \qquad z_h(x) = \frac{\hat{h}(x) - \mu_h}{\sigma_h},$$

with $\mu_{\bullet}, \sigma_{\bullet}$ are computed either per generation or from a held-out calibration set.

**2) Robust alternatives:** Replace means and standard deviations with medians and median absolute deviations (MADs) to reduce the influence of outliers.

**3) Bounded normalization:** Apply quantile clipping, restricting predictions to between the $q$-th and $(1-q)$-th quantiles, followed by rescaling to a fixed range such as $[-1,1]$.

In our implementation, both predictors are further min–max normalized to $[0,1]$ across the candidate pool of each generation. This guarantees scale comparability, avoids domination of one predictor over the other, and stabilizes the optimization dynamics.

In conventional multi-objective optimization, the natural next step would be to analyze Pareto fronts.

However, in this application, such an approach is impractical: the dataset of high-quality references is small (only 27 $Ref^+$ and 48 $Ref^-$ sequences), and the multi-functional fusion problem in this novel sequence landscape is NP-hard. As a pragmatic alternative, we collapse the bi-objective signals into a single scalar reward using the sum and min strategies described later, thereby retaining interpretability while enabling tractable evolutionary optimization.

**III. Exploration and Novelty Components**

A central requirement for Exploration Stage is diversification. The evolutionary algorithm must not only refine and optimize within already known sequence families but also expand into previously uncharted regions of sequence space, thereby increasing the likelihood of discovering novel and functionally relevant motifs. To explicitly encourage this behavior, we introduce quantitative novelty measures that act as exploration-oriented regularizers within the composite loss function.

**(1) Distance from reference sets ($Ref^+/Ref^-$)**

For each candidate sequence $x$, we compute its minimum Hamming distance to the set of reference sequences, including both beneficial ($Ref^+$) and deleterious ($Ref^-$) controls. The Hamming distance between two sequences of equal length is defined as:

$$d_{Hamming}(x, y) = \sum_{i=1}^{L} 1\{x_i \neq y_i\}$$

where $L$ is the sequence length, $x_i$ and $y_i$ denote the residues at position $\boldsymbol{i}$, and $1\{\cdot\}$ is the indicator function. We then define the reference distance as:

$$d_{ref}(x) = \frac{1}{L} \min_{r \in \{Ref^+ \cup Ref^-\}} d_{Hamming}(x, r)$$

This normalized measure lies in the range $[0,1]$. A higher value of $d_{ref}(x)$ indicates that the candidate is more dissimilar to all known references, which we interpret as an indicator of novelty. By contrast, sequences with low $d_{ref}(x)$ are judged to be close to existing reference motifs and therefore less exploratory.

**(2) Diversity relative to the previous generation**

To avoid premature convergence and over-exploitation of recent high-scoring solutions, we additionally encourage diversity relative to the immediately preceding generation $G_{t-1}$. For candidate $x$, we compute:

$$d_{prev}(x) = \frac{1}{L} \min_{y \in G_{t-1}} d_{Hamming}(x, y)$$

Here, $d_{prev}(x)$ represents the normalized distance to the nearest neighbor from the prior generation. Larger values imply that $\boldsymbol{x}$ contributes novel information to the evolving population, while smaller values suggest redundancy or duplication of recent candidates.

Both $d_{ref}(x)$ and $d_{prev}(x)$ serve as exploration-promoting regularizers, shifting the optimization away from pure exploitation of predictive peaks (i.e., maximizing only fitness and binding predictions). Incorporating these terms helps ensure that the evolutionary search does not stagnate in narrow basins of attraction, but instead explores a wider range of the sequence landscape. Conceptually, this balances **quality exploitation** with **novelty exploration**, a trade-off central to efficient evolutionary search in high-dimensional sequence spaces.

**IV. Penalty and Constraint Terms**

While novelty measures encourage exploration, unconstrained diversification can lead to degenerate solutions that sacrifice functional viability in favor of mere dissimilarity. To safeguard against such outcomes, we introduce **explicit penalties and constraints** that ensure candidates retain a baseline level of performance. These penalties operate as *soft regularizers*, reducing the score of sequences that are either too close to known references or that fall below functional baselines.

**1) Proximity penalty**

The first constraint addresses the risk of generating candidates that imitate known reference motifs. Even if such sequences appear viable, they provide little value in terms of novelty or exploration. To prevent this, we impose a penalty that scales with proximity to $Ref^{+}$ and $Ref^{-}$ sequences:

$$P_{ref}(x) = \lambda_{ref\_pen} \cdot (1 - d_{ref}(x))$$

where $d_{ref}(x)$ is the normalized distance from the reference sets (as defined above), and $\lambda_{ref\_pen}$ is a tunable hyperparameter controlling penalty severity. This formulation ensures that sequences with small $d_{ref}(x)$ (i.e., very close to references) are heavily penalized, while sufficiently novel sequences experience little to no penalty.

**2) Baseline threshold penalties**

In addition to penalizing proximity, we enforce functional viability baselines for both production fitness and hTfR1 binding capability. Let $f(x)$ and $h(x)$ denote the normalized predictions for fitness and binding capability, respectively. From the terminal population of Function Integration Stage, we define baseline thresholds $f_{base}$ and $h_{base}$ as the median or chosen quantile of the respective predictive distributions. These values serve as soft viability cutoffs.

Any candidate whose predicted scores fall below these baselines is linearly penalized:

$$P_{fit}(x) = \lambda_{fit\_pen} \cdot \max\{0, f_{base} - f(x)\}$$

$$P_{ht}(x) = \lambda_{ht\_pen} \cdot \max\{0, h_{base} - h(x)\}$$

Here, $\lambda_{fit\_pen}$ and $\lambda_{ht\_pen}$ are tunable weights. By design, these penalties do not suppress sequences performing above baseline; instead, they only penalize those below the threshold, thereby enforcing minimum viability guarantees while preserving freedom for exploration above baseline.

Together, the penalties $P_{ref}(x)$, $P_{fit}(x)$, $P_{ht}(x)$ serve to regularize optimization, preventing collapse into trivial imitation or into non-functional sequence space.

#### 5.4.3 Reward Strategies: Sum vs. Min

After quality normalization and baseline enforcement, the central question remains how to combine fitness and binding predictions into a single reward. To this end, we explored two complementary aggregation strategies:

**1) Sum strategy**

The **sum reward** aggregates both predictive dimensions linearly:

$$R_{sum}(x) = f(x) + h(x)$$

This formulation reflects *aggregate performance*. It rewards candidates that are strong in either fitness or binding, while also favoring improvements in both. Because it tolerates asymmetric trade-offs, the sum strategy tends to produce a more diverse set of viable candidates, albeit with the possibility of admitting partially unbalanced solutions (e.g., very high fitness but moderate binding).

**2) Min strategy**

The **min reward** adopts a "weakest-link" principle:

$$R_{min}(x) = \min\{f(x), h(x)\}$$

This formulation enforces *balanced multifunctional integration*. Candidates are only rewarded if both predicted functions are simultaneously strong; a deficiency in either dimension sharply limits the reward.

##### 5.4.4 Definition of Exploration Stage Score

The Exploration Stage reward function integrates predictive quality, exploration incentives, novelty regularization, and viability constraints into a single scalar score. This design provides a transparent and interpretable means of balancing competing objectives in the evolutionary search process.

Formally, for a candidate sequence $x$, the final Exploration Stage score is defined as:

$$S(x) = \underbrace{R_{\{\text{sum/min}\}}(x)}_{\text{quality}} + \underbrace{\alpha \cdot d_{\text{prev}}(x)}_{\text{exploration}} + \underbrace{\gamma \cdot d_{\text{ref}}(x)}_{\text{novelty}} - (P_{\text{ref}}(x) + P_{\text{fit}}(x) + P_{\text{ht}}(x)).$$

Here, the score is decomposed into four interpretable categories:

1. **Quality term:** $R_{\{\text{sum/min}\}}(x)$ aggregates the outputs of two pretrained predictors, fitness $f(x)$ and hTfR1 binding $h(x)$.
   - In the **sum strategy**, $R_{\text{sum}}(x) = f(x) + h(x)$, representing aggregate functional performance.
   - In the **min strategy**, $R_{\text{min}}(x) = \min\{f(x), h(x)\}$, enforcing balanced multifunctionality through a weakest-link principle.
2. **Exploration term:** $d_{\text{prev}}(x)$ is the normalized minimum Hamming distance to sequences in the previous generation, encouraging exploration of new variants rather than duplication of recently selected motifs. Its contribution is scaled by hyperparameter $\alpha$.
3. **Novelty term:** $d_{\text{ref}}(x)$ is the normalized minimum distance to reference sets ($Ref^{+}/Ref^{-}$). A larger value indicates greater novelty relative to known motifs. Its contribution is scaled by hyperparameter $\gamma$.
4. **Penalty terms:** The sum of $P_{\text{ref}}(x), P_{\text{fit}}(x), P_{\text{ht}}(x)$ enforces proximity constraints and minimum viability guarantees. These are weighted by independent penalty parameters $\lambda$, ensuring that candidates too close to references or falling below functional baselines are systematically discouraged.

##### 5.4.5 Pseudocode for Exploration Stage Evaluation

**Architecture of the FE-X Module**

**Algorithm XVII: FE-X Module** [Architecture]

```
def FE_X(sequence_population, epochs_stage1, epochs_stage2):
    # ----- Function Integration Stage: Multi-functional Fusion -----
    population = Function_Integration_Stage(sequence_population, epochs_stage1)
    # ----- Exploration Stage: Exploration and Diversification -----
    population = Exploration_Stage(population, epochs_stage2)
    return population
```

**Algorithm XVIII: Function Integration Stage** [Multi-functional Fusion]

```
def Function_Integration_Stage(population, epochs):
    for t in range(epochs):
        # Contrastive scoring against Ref+ / Ref- sequences
        scores = []
        for seq in population:
            score = Contrastive_Evaluate(seq, Ref_plus, Ref_minus)
            scores.append(score)
        # Selection + Variation
        population = EvolutionaryStep(population, scores)
    return population
```

**Algorithm XIX: Exploration Stage** [Exploration & Diversification]

```
def Exploration_Stage (population, epochs):
    # Baseline thresholds from Function Integration Stage
    f_base = median(FitnessPredictor(population))
    h_base = median(BindingPredictor(population))
    for t in range(epochs):
        next_population = []
        scores = []
        for seq in population:
            # Predictive quality
            f_pred = Normalize(FitnessPredictor(seq))
            h_pred = Normalize(BindingPredictor(seq))
            # Novelty measures
            d_ref = Distance_to_Refs(seq, Ref_plus, Ref_minus)
            d_prev = Distance_to_Prev(seq, population)
            # Compute reward
            reward = RewardStrategy(seq, f_pred, h_pred, d_ref, d_prev,
                                    f_base, h_base, mode="sum_or_min")
            scores.append(reward)
            next_population.append(seq)
        # Evolutionary update
        population = EvolutionaryStep(next_population, scores)
    return population
```

**Algorithm XX: Contrastive Evaluation** [Function Integration Stage Scoring]

```
def Contrastive_Evaluate(seq, Ref_plus, Ref_minus):
    # Alignment-based similarity scoring
```

```
    sim_pos = mean(SmithWaterman(seq, r) for r in Ref_plus)
    sim_neg = mean(SmithWaterman(seq, r) for r in Ref_minus)

    # Normalize and combine
    base_score = (sim_pos - sim_neg) / 2 + 0.5
    # Optional novelty penalty relative to Ref+
    novelty_penalty = KL_divergence(seq, Ref_plus)
    return Clamp(base_score - novelty_penalty, 0, 1)
```

**Algorithm XXI: Exploration Stage Reward Function** [Reward Function]

```
def RewardStrategy(seq, f_pred, h_pred, d_ref, d_prev, f_base, h_base, mode):
    # Quality term
    if mode == "sum":
        R = f_pred + h_pred
    elif mode == "min":
        R = min(f_pred, h_pred)
    else:
        R = f_pred + h_pred
    # Exploration bonuses
    R = R + alpha * d_prev + gamma * d_ref
    # Penalties
    P_ref = lambda_ref * (1 - d_ref)
    P_fit = lambda_fit * max(0, f_base - f_pred)
    P_ht = lambda_ht * max(0, h_base - h_pred)
    score = R - (P_ref + P_fit + P_ht)
    return Clamp (score, 0, 1)
```

**Algorithm XXII: Evolutionary Step** [Evolution for Exploration and Exploitation]

```
def EvolutionaryStep(population, scores):
    # Select top sequences based on scores
    selected = Selection(population, scores)
    # Generate offspring by mutation / crossover
    offspring = Variation(selected)
    # Return next generation
    return offspring
```

**Notes**

- FitnessPredictor() and BindingPredictor() → evalution models from Function Integration Stage training.
- Clamp() → enforce score range [0,1].
- Hyperparameters: $\alpha$, $\gamma$, $\lambda_{ref}$, $\lambda_{fit}$, $\lambda_{ht}$.
- mode can be "sum" or "min" depending on reward strategy.

This unified formulation allows flexible tuning of evolutionary dynamics through hyperparameters $\alpha$, $\gamma$ and $\lambda$. Increasing $\alpha$ or $\gamma$ biases the search toward exploration and novelty, while higher penalty weights impose stricter viability filters. In practice, these parameters can be adapted depending on the specific experimental priorities—whether the goal is broad exploration of new sequence space or focused refinement of multifunctionally balanced variants.

Overall, this design offers a principled and interpretable scoring system that integrates predictive modeling with explicit evolutionary heuristics, thereby extending the FE module into the FE-X module capable of discovering innovative and multifunctionally competent AAV capsid sequences.

#### 5.5 Hits-Ranking Module——Post-Evolution Ranking and Candidate Selection

Following multiple rounds of FE-X–driven exploratory evolution and functional integration, the generated capsid sequences were initially ranked using the composite scoring function defined in the Exploration Stage:

$$S(x) = \underbrace{R_{\{\text{sum/min}\}}(x)}_{\text{quality}} + \underbrace{\alpha \cdot d_{\text{prev}}(x)}_{\text{exploration}} + \underbrace{\gamma \cdot d_{\text{ref}}(x)}_{\text{novelty}} - (P_{\text{ref}}(x) + P_{\text{fit}}(x) + P_{\text{ht}}(x)).$$

This unified scoring function integrates predictive quality, exploration incentives, novelty contributions, and explicit penalty terms. Based on these scores, the top 1,000 sequences were retained as preliminary candidates.

To further prioritize sequences with balanced performance, we applied pretrained evaluation models to obtain independent predictions of production fitness ($f(x)$) and hTfR1 log2 enrichment $h(x)$). These two predictive readouts were then combined into a comprehensive prediction score (CP Score) defined as:

$$CP\ Score(x) = \frac{f(x) + h(x)}{2}$$

This averaging scheme gives equal weight to production fitness and receptor-targeting capability, thereby enforcing balanced multifunctional integration. Sequences were re-ranked according to this comprehensive prediction score, and the top 500 sequences were selected for oligonucleotide library synthesis.

### 6.Training Generative Models from Scratch for the Design of Multifunctional Capsid Sequences

To investigate whether traditional AI architecture models can directly generate capsid sequences with integrated multifunctionality, we trained models based on SeqGAN-only, VAE, and Diffusion architectures from scratch. The training was performed on a dataset $\boldsymbol{D_{Seq}}$ (n = 72,753; details provided in Supplementary Methods) using an 80:20 split for the training and test sets, respectively. Hyperparameters were optimized via a randomized parameter search. The parameters for SeqGAN remained consistent with those described in the main text, while the architectural parameters for the VAE and Diffusion models are specified in the TableS11.

After training, each generative model was used to design 1,000 novel capsid sequences. These sequences were subsequently evaluated using an evaluation model to assess key properties: production fitness, Ly6a log2 enrichment score, and Ly6c1 log2 enrichment score.

The results indicate that sequences generated by all three architectures largely recapitulate the distribution of the training set. However, none of the models succeeded in achieving integrated multifunctionality. This highlights the inherent challenges conventional deep learning models face when confronted with highly complex tasks that demand multifunctional integration under limited data conditions. In particular, although these generative models are adept at reproducing the statistical properties of the training distribution, they remain limited in their ability to explore the broader functional landscape required for effective multi-feature fusion.

### 7. Minimum Spanning Tree (MST) Construction on 7-mer Hamming Space

We visualize the relational "backbone" of capsid variants in sequence space by constructing a **minimum spanning tree (MST)** over AAV capsid sequences. Two panels are produced: (i) $\boldsymbol{Ref^+}$ ∪ **ALICE-X**, and (ii) $\boldsymbol{Ref^+}$ ∪ **ALICE**. Nodes encode hTfR1 binding capability (measured by log2 enrichment) by color; node shape distinguishes **AI** (circle) from $\boldsymbol{Ref^+}$ (square).

#### 7.1 Data preparation

When a cap on total nodes is imposed, all $\boldsymbol{Ref^+}$ sequences are preserved in full, while the AI set is proportionally reduced so that the combined dataset does not exceed the specified node limit. A fixed ordering is applied to ensure reproducibility across runs. The hTfR1 metric is coerced to floating point.

#### 7.2 Sequence distance calculation

Let a sequence be $s_i = (a_{i1}, \dots, a_{i7})$ with $a_{ik} \in \mathcal{A}$ (20 canonical amino acids). The **H**amming distance between two sequences is

$$d_H(s_i, s_j) \;=\; \sum_{k=1}^{7} \mathbf{1}\{a_{ik} \neq a_{jk}\},$$

an integer in $[0,7]$ equal to the number of positions at which the 7-mers differ. This yields a symmetric distance matrix $D = [d_H(s_i, s_j)]$ on the combined node set.

#### 7.3 Graph and MST construction

We build the complete weighted graph $G = (V, E)$ with $V$ the unique sequences and edge weights $w_{ij} = d_H(s_i, s_j)$. The minimum spanning tree $T^*$

$$T^* = arg \underset{T \text{ is a spanning tree on } V}{min} \sum_{(i,j)\in T} w_{ij}.$$

By favoring the shortest available mutational steps (small $d_H$) while minimizing total cost globally, the MST provides a sparse, comprehensible backbone of nearest-neighbor relationships.

##### 7.3.1 Layout and axes

The tree is embedded in 2D using a force-directed (spring) layout. Let $x_i \in \mathbb{R}^2$ be the coordinates returned by the layout algorithm that approximately preserve graph-theoretic proximities. The x- and y-axes are unitless and do not correspond to biophysical variables; they simply provide a visually stable arrangement in 2D. We therefore suppress ticks and labels and emphasize connectivity, edge weights, and group envelopes rather than absolute positions.

#### 7.3.2 Visual encodings

**Node color (hTfR1 log2 enrichment).**

Let $y_i$ be hTfR1 $\log_2$ enrichment for node $i$. Colors are mapped using a linear normalization

$$c_i = \frac{y_i - y_{\min}}{y_{\max} - y_{\min}},$$

where $y_{\min}, y_{\max}$ are either (a) panel-specific or (b) shared across both panels to support cross-panel comparability.

**Node shape and size.**

Nodes from ALICE-X or ALICE are drawn as circles; $\boldsymbol{Ref^{+}}$ nodes as squares.

**Edge width (sequence similarity).**

For each edge $(i,j)$ in the MST, let $d_{ij} = d_H(s_i, s_j)$ denote the Hamming distance and $s_{ij} = 7 - d_H \in [0,7]$ the corresponding sequence similarity (larger values indicate more similar sequences). The visual line width assigned to that edge is

$$\text{lw}_{ij} = a + b\log(1 + s_{ij}),$$

where $\text{lw}_{ij}$ denotes the plotted edge thickness in the figure (in points), $s_{ij}$ is the similarity score, and $\boldsymbol{a}, \boldsymbol{b} > \boldsymbol{0}$ are small constants chosen for display scaling. Thus, edges between highly similar sequences (small Hamming distance, larger $s_{ij}$) are drawn thicker, facilitating visual discrimination of short mutational steps.

### 8. Deep Motif Relationship Analysis

To characterize recurrent sequence motifs across multiple architecture in ablation study, we constructed motif-level maps integrating global normalization, motif aggregation, and sequence-similarity edges. For each sequence $s$, fitness $x(s)$ and hTfR1 $\log_2$ enrichment $y(s)$ were pooled across all datasets and normalized using min–max scaling:

$$x^*(s) = \frac{x(s) - \min(x)}{\max(x) - \min(x)}, \quad y^*(s) = \frac{y(s) - \min(y)}{\max(y) - \min(y)}.$$

The global means $\mu_x$ and $\mu_y$ of normalized values served as baselines for effect-aware coordinate construction.

Motifs were extracted as overlapping $k$-mers ($k = 3$) from each sequence, and duplicates within a sequence were collapsed. For each motif $m$, we computed mean normalized values

$$\overline{x}_m = \frac{1}{|S_m|}\sum_{s \in S_m} x^*(s), \quad \overline{y}_m = \frac{1}{|S_m|}\sum_{s \in S_m} y^*(s),$$

with $S_m$ denoting the set of sequences containing $m$. Motifs were ranked by relative frequency $freq_m = |S_m|/|S_f|$, and the top 15 motifs generated by each architecture in the ablation study were retained.

Each motif was positioned on effect-aware axes as standardized deviations from the global means:

$$X_m = \frac{\overline{x}_m - \mu_x}{\sigma_x}, \quad Y_m = \frac{\overline{y}_m - \mu_y}{\sigma_y},$$

where $\sigma_x$ and $\sigma_y$ are standard deviations across motifs. Thus, $X = 0$ and $Y = 0$ mark the global reference, with quadrants reflecting relative gains or losses in fitness and enrichment. Nodes were encoded by size (proportional to $\overline{x}_m$) and color (proportional to $\overline{y}_m$).

To reveal sequence neighborhoods, motifs were connected by edges when their Hamming distance satisfied:

$$d_H(m_i, m_j) = \sum_{k=1}^{K} \mathbf{1}\{m_{i,k} \neq m_{j,k}\} \leq 1.$$

Edge thickness scaled inversely with distance, emphasizing single-substitution neighbors. The resulting graphs provide a dual representation: **axes** capture functional deviation from global means, while **edges** reveal local mutational proximity.

The resulting graphs illustrate both the relative effect profile of motifs (via axes) and their local sequence relatedness (via Hamming edges). Motifs clustering near the upper-right quadrant (X>0, Y>0) exhibit above-average performance in both production fitness and hTfR1 binding capability, while edges reveal minimal substitutions that connect them to neighboring motifs of differing functional profiles.

### 9. Trajectory Relationship Analysis

This analysis illustrates the evolutionary trajectory of sequence populations across epochs by simultaneously tracking their mean proximity to positive ($Ref^+$) and negative ($Ref^-$) references. The resulting bivariate path embeds each epoch as a point $(x_t, y_t)$ and connects points in chronological order to visualize directional movement in reference space.

#### 9.1 Data sources and preprocessing

To maintain computational consistency, a fixed set of 500 sequences was retained from each evolutionary epoch in FE-X. Each row contains an amino-acid sequence $s$ (uppercase, 20-letter alphabet). Two external reference sets were supplied: a set of **positives** $Ref^+ = \{r_1^+, \dots, r_{n_+}^+\}$ and a set of **negatives** $Ref^- = \{r_1^-, \dots, r_{n_-}^-\}$.

#### 9.2 Pairwise metric families

For any two sequences $\boldsymbol{a}, \boldsymbol{b}$, a scalar score $\psi(a, b)$ is computed using one of the following families; a small pseudocount $\varepsilon$ is employed where needed to avoid zero divisions.

**a) Position-wise KL divergence.**

For a sequence set $\mathcal{S}$ of equal-length strings, define the per-position empirical distributions $P_i(a)$ over the 20 amino acids $a$ at position $i$ (rows sum to 1). Given two distributions $P = \{P_i\}$ and $Q = \{Q_i\}$, the Kullback–Leibler divergence is

$$D_{\mathrm{KL}}(P\|Q) = \sum_{i=1}^{L}\sum_{a} P_i\,(a)\log\frac{P_i(a)}{Q_i(a)},$$

with $Q_i(a) \leftarrow Q_i(a) + \varepsilon$ and re-normalization. In the minimal setting where $P$ and $Q$ arise from single sequences, the per-position distributions are one-hot (smoothed by $\varepsilon$).

**b) $n$-gram overlap (contiguous $n$-mers).**

Let $\mathcal{N}_n(a)$ be the multiset of $n$-grams in $a$. The overlap ratio is

$$\psi_{n\text{-gram}}(a,b) = \frac{|\mathcal{N}_n(a) \cap \mathcal{N}_n(b)|}{\max\{|\mathcal{N}_n(a)|, |\mathcal{N}_n(b)|, 1\}}.$$

**c) LCS ratio (longest common subsequence).**

$$\psi_{\mathrm{LCS}}(a,b) = \frac{|\mathrm{LCS}(a,b)|}{\max\{|a|, |b|\}}.$$

These metrics deliberately span position-aware (KL), substring (n-gram), and global composition perspectives, with LCS providing order-aware complements.

#### 9.3 Epoch-level aggregation to references

For epoch $t$, let $S_t$ denote the set of sequences retained. The epoch-level mean score to the positive and negative reference sets is defined as a two-way average:

$$\mu_t^+ = \frac{1}{|S_t|\,|R^+|}\sum_{s\in S_t}\sum_{r^+\in R^+}\psi(s,r^+), \qquad \mu_t^- = \frac{1}{|S_t|\,|R^-|}\sum_{s\in S_t}\sum_{r^-\in R^-}\psi\,(s,r^-).$$

Here, $\psi(s,r)$ serves as a generic sequence–sequence comparison function.

#### 9.4 Trajectory embedding and rendering

Each epoch $t$ is embedded as the point

$$(x_t, y_t) = (\mu_t^-, \mu_t^+),$$

and epochs are connected in ascending order to form a polyline. Points are color-mapped by epoch index (continuous colormap), with the first epoch (start) highlighted by an open circle and the last epoch (end) by a filled star.

#### 9.5 Axis semantics and quadrant interpretation

Axis labels explicitly denote the metric and target set:

- **X-axis:** "$\psi$ to $Ref^-$" $\equiv \mu_t^-$.
- **Y-axis:** "$\psi$ to $Ref^+$" $\equiv \mu_t^+$.

Because metrics differ in polarity, we separately consider similarity-oriented measures (where higher values denote stronger similarity, such as LCS or n-gram overlap) and distance-oriented measures (where higher values denote greater dissimilarity, such as KL divergence).

Ultimately, the resulting trajectory graphs provide a compact, metric-aware visualization of how evolving sequence cohorts move in $Ref^{+}/Ref^{-}$ space over epochs, revealing convergence toward desirable regions or trade-offs that depend on the choice of similarity/distance criterion.

### 10. Efficacy Analysis for Reward, Reference Distance, and Diversity

To systematically assess how different evolutionary drivers contribute to functional optimization, we established a unified framework that places reward signals, reference distances, and inter-generational diversity under the same analytical lens. The goal was to quantify whether sequences stratified by these variables at epoch $t$ are predictive of subsequent improvements in production fitness or hTfR1 enrichment at epoch $t + 1$.

This approach extends the classical notion of "reward efficacy" into a generalizable paradigm, allowing us to treat reward design, deviation from reference sequences (measured by reference distance), and evolution diversity as parallel forces in shaping evolutionary trajectories. By applying a unified median-shift procedure across all three variables, we enabled direct comparison of each factor's contribution to hTfR1 $\log_2$ enrichment and production fitness throughout the Exploration Stage of evolution.

#### 10.1 Population and associated metrics

At every epoch $t$, the evolving sequence population is denoted as:

$$P_t = s_1, s_2, \ldots, s_{n_t}.$$

Each sequence $s_i \in P_t$ carries a set of associated metrics. First, a reward score $r_i^{(t)}$ is assigned, reflecting the optimization objective under the chosen reward mode (e.g., sum, min, or extended forms that integrate diversity and reference distance).

Second, two predictive outcomes are recorded: the predicted production fitness $\hat{f}_i^{(t)}$ and the predicted hTfR1 enrichment $\hat{h}_i^{(t)}$. Both are obtained from supervised predictors and rescaled to the unit interval via min–max normalization to ensure comparability across evaluation batches.

Beyond these predictions, two distance-based signals are central to Stage-2 evolution. The first is the reference distance $\rho_i^{(t)}$, which measures the minimal Hamming distance between sequence $s_i$ and any sequence in the positive or negative reference sets $(\mathrm{Ref}^{+}, \mathrm{Ref}^{-})$, normalized by sequence length $L$:

$$\rho_i^{(t)} = \frac{1}{L} \min_{r \in \mathrm{Ref}^{+} \cup \mathrm{Ref}^{-}} \mathrm{Ham}(s_i, r).$$

Larger values indicate greater divergence from known sequences and are thus interpreted as favorable for innovation.

The second is the diversity score $\delta_i^{(t)}$, which measures the minimal Hamming distance between $s_i$ and sequences from the preceding epoch $P_{t-1}$:

$$\delta_i^{(t)} = \frac{1}{L} \min_{p \in P_{t-1}} \mathrm{Ham}(s_i, p).$$

This term captures the extent to which a candidate represents genuine exploration rather than rediscovery of prior solutions.

Finally, soft penalties are incorporated to prevent the evolutionary process from collapsing onto trivial but non-functional solutions. Baseline values for production fitness and enrichment $(f_{\text{base}}, h_{\text{base}})$ are defined as the median predictions within the starting population, and sequences falling below these thresholds accrue linear penalties. Together, these metrics provide a multi-dimensional characterization of each sequence, balancing predicted performance with novelty and exploration.

#### 10.2 Stratification and Quartiles

To analyze causal influence, we stratified sequences at epoch $t$ according to quartiles of a chosen variable $X_i^{(t)} \in r_i^{(t)}, \rho_i^{(t)}, \delta_i^{(t)}$.

Within each epoch, sequences were ranked by their $X$-values and partitioned into quartile subsets:

$$Q_X^{(t)} = Q_{\text{Top25}}^{(t)}, Q_{50-75}^{(t)}, Q_{25-50}^{(t)}, Q_{\text{Bottom25}}^{(t)}.$$

This stratification allows us to disentangle whether the highest-scoring quartiles (e.g., the most diverse or most reference-distant sequences) exhibit distinct evolutionary consequences relative to their lower-scoring counterparts.

#### 10.3 Stratification and Quartiles

For each quartile $q$, we calculated the median performance at epoch $t$ for a given outcome metric $M$ (fitness or enrichment):

$$\widetilde{M}_{(q,X)}^{(t)} = \mathrm{median} M_i^{(t)} \mid s_i \in Q_X^{(t)}.$$

At the subsequent epoch, we then computed the population-level median:

$$\widetilde{M}^{(t+1)} = \mathrm{median} M_j^{(t+1)} \mid s_j \in P_{t+1}.$$

The efficacy score of quartile $q$ under stratification variable $X$ is defined as the shift:

$$\Delta M_{(q,X)}^{(t \to t+1)} = \widetilde{M}^{(t+1)} - \widetilde{M}_{(q,X)}^{(t)}.$$

A positive value indicates that sequences belonging to quartile $q$ at epoch $t$ were predictive of improvement in the next generation; a negative value indicates that these sequences were disadvantageous.

#### 10.4 Stratification and Quartiles

This unified procedure was applied in three distinct contexts.

When $X = r$, the analysis captures **reward efficacy**, testing whether sequences exposed to different levels of reward at epoch $t$ genuinely translate into improvements at epoch $t + 1$.

When $X = \rho$, it captures reference **distance efficacy**, quantifying whether divergence from canonical sequences/motifs promotes functional gains.

When $X = \delta$, it captures **diversity efficacy**, revealing whether inter-generational dissimilarity effectively broadens the search space and enhances outcomes.

In all cases, the quartile-based design enables nuanced interpretation: strong positive efficacy in the top quartile suggests that maximizing the corresponding driver is beneficial, whereas flat or negative shifts suggest diminishing returns or counterproductive effects.

The efficacy scores were visualized as heatmaps spanning training epochs. Rows correspond to quartiles, columns to epochs, and color encodes the magnitude of $\Delta M$. These visualizations provide an intuitive, time-resolved picture of how reward strength, reference divergence, and evolution diversity shape the functional landscape.

By embedding reward, distance, and diversity signals in a single formalism, this framework moves beyond ad hoc comparisons and delivers a systematic, causally interpretable analysis of evolutionary pressures. Crucially, it allows us to pinpoint not only which strategies are beneficial, but also when during training their influence is most pronounced—an essential insight for the design of adaptive evolutionary objectives.

### 11. Pareto Front–based Ranking and Selection of Sequences

To identify sequences that simultaneously achieve high production fitness and improved receptor targeting capability, we employed a Pareto front–based multi-objective ranking framework. This approach avoids reducing the problem to a single weighted objective, which may bias selection toward one function at the expense of the other, and instead preserves solutions that represent optimal trade-offs between competing criteria.

#### 11.1 Objective Definition

Each candidate sequence $i$ was jointly evaluated on two maximization objectives:

**Production fitness** ($f_i$): Defined as the experimentally measured or model-predicted production fitness value of sequence $i$.

**Targeting enrichment gain relative to wild-type (WT)** ($\Delta_i$): Defined as the relative increase in transduction efficiency observed in TFRC knock-in (KI) mice compared to WT controls:

$$\Delta_i = \text{Log2\_Enrichment}_{\text{TFRC KI},\, i} - \text{Log2\_Enrichment}_{\text{WT},\, i}.$$

Thus, each sequence can be represented as a two-dimensional objective vector:

$$\mathbf{v}_i = (f_i, \Delta_i).$$

#### 11.2 Pareto Dominance

Pareto dominance was defined following standard multi-objective optimization theory. Specifically, sequence $p$ is said to dominate sequence $q$ (denoted $p \prec q$) if:

$$f_p \geq f_q \quad \text{and} \quad \Delta_p \geq \Delta_q,$$

with at least one strict inequality.

This definition ensures that no dominated sequence can be strictly better in both objectives simultaneously, thereby filtering out candidates that are inferior across the board.

#### 11.3 Non-Dominated Sorting and Ranking

Using this dominance relation, we applied **non-dominated sorting** to partition the population into Pareto fronts:

- **First Pareto front (Rank = 1):** Contains all sequences that are not dominated by any other sequence in the dataset.
- **Second Pareto front (Rank = 2):** Contains sequences that are dominated only by members of the first front, and so forth.

This stratification yields a hierarchy of trade-off solutions, with lower-ranked fronts corresponding to better compromise sets.

#### 11.4 Within-Front Prioritization

To refine candidate selection within the top-ranked front, we implemented a lexicographic ordering scheme:

1. Sequences were first ordered by **descending fitness** ($f_i$), ensuring that production viability remains a primary determinant.
2. In cases of ties, sequences were further ranked by **descending enrichment gain** ($\Delta_i$), prioritizing improved targeting performance.

This two-level ordering ensures that the selected set contains sequences that are not only Pareto-optimal but also practically viable in terms of production.

#### 11.5 Candidate Selection

We select sequences from the top 6 Pareto-ranked sets(~ 100 capsid sequences). This ensures that all chosen candidates represent non-dominated solutions, i.e., no selected sequence is simultaneously outperformed in both objectives by another sequence.

By design, this Pareto-based selection process guarantees balanced optimization of fitness and targeting gain, avoids collapsing into one-dimensional rankings, and ensures that the final candidate set maintains both biological relevance and diversity in performance trade-offs.

### 12. Ablation Study for Multifunctional Integration and Novel Sequence Exploration Assessed by Pull-down Assays

The ablation analysis was performed to clarify the functional role of individual ALICE-X modules in achieving multifunctional integration of capsid properties. In particular, we aimed to assess whether combinations of semantic tuning, ranking, and FE/FE-X modules could balance production fitness with hTfR1 binding capability, while still exploring novel sequence motifs.

For each architecture under comparison—ALICE-X, ALICE, ST Hits-Rank, ST, and RM—500 sequences were generated and jointly synthesized into a single AAV variant library—pAAV-588-OligoPool-lib2. This design ensured that all variants were evaluated in parallel under identical pulldown assay conditions, minimizing experimental batch effects.

To examine the trade-off between fitness and targeting, we applied a Top@k analysis, whereby sequences are first ranked by production fitness $f(s)$ and the top-$k$ subset retained:

$$\boldsymbol{Top@k} = \{\boldsymbol{s} \in \boldsymbol{S} \mid \boldsymbol{rank}(\boldsymbol{f}(\boldsymbol{s})) \leq \boldsymbol{k}\}.$$

For each group, the Top@100 sequences were analyzed for their hTfR1 log2 enrichment values, providing a focused view of targeting capability among the most viable variants. This approach allowed us to directly assess the extent to which functional fidelity (fitness) and exploratory novelty (binding to hTfR1) could be simultaneously achieved.

The comparative results indicated that ALICE-X and ALICE architectures successfully realized such functional fusion, retaining high production fitness while maintaining strong hTfR1 binding. In contrast, architectures lacking the FE or FE-X modules produced sequences with reduced binding capacity, highlighting the indispensability of these modules for multifunctional optimization.

### 13. UMAP visualization of sequence embeddings

We employed uniform manifold approximation and projection (UMAP) to visualize the high-dimensional sequence data generated by our ALICE system in two-dimensional space. UMAP is a dimension reduction technique that preserves both the local and global structure of the data, making it particularly suitable for visualizing complex biological sequences[14].

Our analysis pipeline begins with the conversion of amino acid sequences into numerical representations. Each amino acid is assigned a unique integer value according to a predefined dictionary, allowing for the transformation of sequences into vectors of equal length.

The numerical sequence data are then standardized via scikit-learn (https://scikit-learn.org/stable/)'s StandardScaler to ensure that all the features contribute equally to the analysis. These standardized data serve as inputs to the UMAP algorithm.

UMAP constructs a topological representation of high-dimensional data on the basis of fuzzy simplicial sets. For each point $x_I$, UMAP computes a local fuzzy simplicial set representation:

$$\sigma_i = \sum_{x_i, x_j < r_i}^{j} e^{\frac{-d(x_i - x_j) - r_i}{\sigma_i}}$$

where $d(x_i - x_j)$ is the distance between points $x_i$ and $x_j$, $r_i$ is the distance to the nearest neighbor of $x_i$, and $\sigma_i$ is a normalization factor.

The global fuzzy simplicial set is then obtained by combining these local representations:

$$\mu(X) = \cup_{i=1}^{n} \sigma_i$$

UMAP then finds a low-dimensional representation $Y$ that minimizes the cross-entropy between the high-dimensional and low-dimensional fuzzy simplicial sets:

$$CE(\mu(X), v(Y)) = \sum_{i,j} \mu_{i,j}(X) \log\left(\frac{\mu_{i,j}(X)}{v_{i,j}(Y)}\right) + (1 - \mu_{i,j}(X)) \log\left(\frac{(1 - \mu_{i,j}(X))}{1 - v_{i,j}(Y)}\right)$$

We utilized the UMAP implementation available in the umap-learn Python library (https://github.com/lmcinnes/umap). The UMAP algorithm constructs a high-dimensional graph representation of the data and then finds a low-dimensional embedding that preserves the topological structure of this graph. This process results in a two-dimensional representation of our sequence data, where similar sequences are positioned closer together.

To enhance the interpretability of the visualization, we incorporated a kernel density estimation (KDE) via the Gaussian_kde function. This allows for the visualization of the density of points in the two-dimensional UMAP space, highlighting areas of high sequence concentration.

The resulting visualization combines scatter plots of individual sequences, color-coded by their source (e.g., positive references, negative references, EDA-generated sequences), with a contour plot representing the density estimation. This approach provides a comprehensive view of the distribution and relationships among different sequence sets in our dataset.

### 14. WebLogo Computations

We appreciate the reviewer’s attention to the interpretation of the WebLogo plots. By default, WebLogo computes the stack height at each position as the information content (IC), defined as:

$$IC(i) = \log_2(K) - H(i), H(i) = -\sum_{a} p_i(a) \log_2 p_i(a)$$

where K is the alphabet size (20 for amino acids) and $p_i(a)$ is the frequency of amino acid $a$ at position i. Thus, positions with strong conservation show tall stacks, and positions with diverse residues show shorter stacks.

In the WebLogo visualizations used in our manuscript (including Fig. 5a and Fig. 6b), we intentionally applied the “unit height” (frequency-based) normalization mode, in which the total stack height is fixed to 1 for all positions and only the relative frequencies of amino acids are displayed. This mode was selected to highlight positional preferences and residue composition rather than global conservation differences across sites. Because of this normalization, all positions display stacks of equal height even though their underlying Shannon entropy values differ.

Importantly, the conservation values are fully computed during WebLogo generation; we simply chose the frequency-normalized visualization for interpretability and comparability across peptide positions. When using the default information-content mode, the logos show variable stack heights consistent with positional entropy calculations.

**Supplementary Table 1. Dataset distribution.**

The table outlines the distribution and characteristics of the datasets used in the study, providing data sources and insights into the diversity and scope of the data analyzed.

| Datasets | Usage Stage | Total Amount |
|---|---|---|
| Book1 datasets | Statistics for Zipf's Law and Sentence Structure in NLP | Containing 11,038 books (approximately 1,816,414 sentences) of 16 different subgenres |
| Swiss-Prot database | Statistics for Zipf’s law and Sentence Structure in biology | 570,830 reviewed protein sequence entries |
| UniProt database | Pretraining | 2,896,202 protein sequence entries with sequence lengths shorter than 50 amino acids (AA) |
| AAV Capsid Library | Semantic Tuning | 72,753 capsid sequences collected for Semantic Tuning to learn the semantic language of AAV |
| AAV Capsids reported and verified in the literature | Function-guided Evolution (FE) | 129 capsid sequences including $\boldsymbol{Ref^{+}}$ and$\boldsymbol{Ref^{-}}$ |
| Features Evaluation datasets | Ranking Filtration Process | A total of 119,851 capsid sequences, designated as $\boldsymbol{D_{Production\ Fitness}}$, were collected for training the model to predict production fitness. Additionally, 89,169 capsid sequences, labeled as $\boldsymbol{D_{Ly6a}}$, were gathered for training the model to predict Ly6a binding ability. Finally, 89,040 capsid sequences, referred to as $\boldsymbol{D_{Ly6c1}}$, were assembled for training the model to predict Ly6c1 binding ability. |

**Supplementary Table 2. Datasets constructed with $\boldsymbol{Ref^{+}}$ and $\boldsymbol{Ref^{-}}$ for functional evolutionary analysis.**

This table presents the $\boldsymbol{Ref^{+}}$ and $\boldsymbol{Ref^{-}}$ datasets utilized for the functional evolutionary analysis. The $\boldsymbol{Ref^{+}}$ dataset comprises sequences that exhibit the following: 1) High production fitness: These sequences demonstrate superior replication and infectivity capabilities. 2) Binding affinity to Ly6a and Ly6c1: These sequences have the ability to bind to Ly6a or Ly6c1, which are key cellular receptors involved in viral entry and immune regulation. 3) Documented BBB permeability, high viability, and in vivo validation: These sequences have been reported in the literature to traverse the blood–brain barrier (BBB), exhibit high viability, and have been validated in vivo across multiple mouse strains (C57BL/6 and BALB/c). The $\boldsymbol{Ref^{-}}$ dataset, in contrast, includes sequences that 1) Exhibit insufficient production fitness: These sequences demonstrate subpar replication and infectivity capabilities. 2) Lack of binding affinity to Ly6a and Ly6c1: These sequences are unable to bind to either of the key cellular receptors Ly6a and Ly6c1, thereby hindering viral entry and immune regulation. The analysis of these $\boldsymbol{Ref^{+}}$ and $\boldsymbol{Ref^{-}}$ datasets aims to elucidate the underlying features that contribute to the superior characteristics of $\boldsymbol{Ref^{+}}$ sequences and the detrimental characteristics of $\boldsymbol{Ref^{-}}$ sequences in the context of functional evolutionary processes.

| $\boldsymbol{Ref^{+}}$ | | | | | | | |
|---|---|---|---|---|---|---|---|
| No. | Sequence | No. | Sequence | No. | Sequence | No. | Sequence |
| 1 | AKAGWSS | 17 | IKAGYSS | 33 | RFAGDAS | 49 | VGVAPPR |
| 2 | APGSAVW | 18 | KYLGDLS | 34 | RLAGASV | 50 | VHQGYSS |
| 3 | ARVGYAQ | 19 | LGVRLGT | 35 | RSVGTIY | 51 | VRTGYAQ |
| 4 | ATVSAVV | 20 | LKPGWAQ | 36 | RWTGESQ | 52 | VRTGYST |
| 5 | DGTLSRA | 21 | LKSGYSQ | 37 | RYAGDST | 53 | VRVGLAQ |
| 6 | DQSSARW | 22 | LRAGFSS | 38 | RYAGDSV | 54 | VSRGYST |
| 7 | ENRGFST | 23 | LRAGYSM | 39 | RYSGDAA | 55 | VTRGYSS |
| 8 | ERAGYSS | 24 | LRAGYSS | 40 | RYTGEAQ | 56 | VTYGGSQ |
| 9 | ETRGANS | 25 | LRVGTVY | 41 | SAPFWTE | 57 | WKTMGAS |
| 10 | FRLQGSA | 26 | LSRGMAQ | 42 | SHNFASI | 58 | WRDMGSQ |
| 11 | GGTKESL | 27 | LTLTTSK | 43 | TRAGMSQ | 59 | WRNLGSA |
| 12 | GGYSERF | 28 | LYVGLSS | 44 | TRAGYSS | 60 | YGSLNAL |
| 13 | GNDPGRW | 29 | PAPGWSS | 45 | TRNGYST | 61 | YLVGFKP |
| 14 | GNETGRW | 30 | PMQRSFG | 46 | TTRGYSV | | |
| 15 | IARGYSV | 31 | PYVGMAS | 47 | VAVGASG | | |
| 16 | IIKGYSS | 32 | REGEARW | 48 | VDRGMVI | | |
| Reported Capsid Sequences | | | | | | | |
| No. | Sequence(Name, Method)[ref] | No. | Sequence(Name, Method)[ref] | No. | Sequence(Name, Method)[ref] | No. | Sequence(Name, Method)[ref] |
| 1 | ERVGFAQ(PHP.C3, | 5 | SIERPFK(PHP.B7, | 9 | TLQIPFK(PHP.B4, | 13 | TTLKPFL(PHP.V2, |

| | | | | | | | |
|---|---|---|---|---|---|---|---|
| | CREATE selection methods)[15] | | M-CREATE selection methods)[15] | | M-CREATE selection methods)[15] | | M-CREATE selection methods, )[15] |
| 2 | FTLTTPK(PHP.B3, M-CREATE selection methods)[16] | 6 | SVSKPFL(PHP.B2, CREATE selection methods, Ly6a)[16] | 10 | TLQLPFK(PHP.B5, M-CREATE selection methods)[15] | 14 | TVSALFK(AAV.CPP.21, rational design)[17] |
| 3 | QAVRTSL(PHP.S, CREATE selection methods, unknown)[18] | 7 | TALKPFL(PHP.V1, M-CREATE selection methods)[15] | 11 | TLQQPFK(PHP.B6, M-CREATE selection methods)[15] | 15 | WSTNAGY(PHP.C2, M-CREATE selection methods)[15] |
| 4 | RYQGDSV(PHP.C1, M-CREATE selection methods)[15] | 8 | TLAVPFK(PHP.B, M-CREATE selection methods)[16] | 12 | TMQKPFI(PHP.B8, M-CREATE selection methods)[15] | | |
| $\boldsymbol{Ref^{-}}$ | | | | | | | |
| No. | Sequence | No. | Sequence | No. | Sequence | No. | Sequence |
| 1 | AGTALGR | 15 | GSNRGGG | 29 | QRLAAGQ | 43 | THLKPFI |
| 2 | ALGSGSW | 16 | GTSKPFL | 30 | RFSPDAA | 44 | TSIGFAA |
| 3 | ARAGENV | 17 | IRGATGP | 31 | RNDATLT | 45 | TSRLPFI |
| 4 | ASHGPSP | 18 | KGTGGYV | 32 | RPASTSW | 46 | TTGRPFV |
| 5 | DATRTYA | 19 | LGRALGY | 33 | RYVGSSS | 47 | VANRNTA |
| 6 | DSAGTRN | 20 | LMAGYSA | 34 | SGKPPDD | 48 | VGTRLGL |
| 7 | EHSGGRW | 21 | LNDHRSQ | 35 | SGRPLGY | 49 | VVAGSVV |
| 8 | FDRNGPK | 22 | LVEKPFR | 36 | SPSRTSW | 50 | VVPPLSN |
| 9 | GDRDGRT | 23 | NGGVGMG | 37 | SRSSETR | 51 | VYGEKER |
| 10 | GDSMHLA | 24 | NGRPVAG | 38 | TAGWGAP | 52 | WDHSGDK |
| 11 | GGTASYS | 25 | NNFHEAR | 39 | TEKTAQI | 53 | YASAERN |
| 12 | GHADNRT | 26 | NRALGSS | 40 | TGRIDTR | | |
| 13 | GSFMPPP | 27 | PKTGSIL | 41 | TGSGHSV | | |
| 14 | GSLAAQD | 28 | PSHAIGV | 42 | TGVAPAA | | |

**Supplementary Table 3. Hyperparameters of the sequence function evaluation model.**
We used various regression models as candidate models for sequential function evaluation, including support vector regression (SVR), eXtreme gradient boosting (XGBoost), K-nearest neighbors (KNN), random forest (RF), gradient boosting (GB), Bayesian ridge, and AdaBoost. Each model's hyperparameters were individually adjusted and experimented with, resulting in the determination of the best candidate model for this module.

| Model Name | Parameter Name | Value |
|---|---|---|
| KNN | n_neighbors | 5 |
| | weights | "uniform" |
| | algorithm | "auto" |
| | leaf_size | 30 |
| | p | 2 |
| | metric | "minkowski" |
| | metric_params | None |
| | n_jobs | None |
| SVR | kernel | "rbf" |
| | degree | 3 |
| | gamma | "scale" |
| | coef0 | 0 |
| | tol | 1.00E-03 |
| | C | 1 |
| | epsilon | 0.1 |
| | shrinking | TRUE |
| | cache_size | 200 |
| | verbose | FALSE |
| | max_iter | -1 |
| Bayesian Ridge | compute_score | TRUE |
| | n_iter | 300 |
| | tol | 1.00E-03 |
| | alpha_1 | 1.00E-06 |
| | alpha_2 | 1.00E-06 |
| | lambda_1 | 1.00E-06 |
| | lambda_2 | 1.00E-06 |
| | alpha_init | None |
| | lambda_init | None |
| | fit_intercept | TRUE |
| | normalize | "deprecated" |
| | copy_X | TRUE |
| | verbose | FALSE |
| RF | n_estimators | 500 |
| | n_jobs | -1 |
| | oob_score | TRUE |

| | | |
|---|---|---|
| | max_features | "sqrt" |
| | criterion | "squared_error" |
| | max_depth | None |
| | min_samples_split | 2 |
| | min_samples_leaf | 1 |
| | min_weight_fraction_leaf | 0 |
| GB | n_estimators | 500 |
| | learning_rate | 0.1 |
| | max_depth | 15 |
| | max_features | "sqrt" |
| | min_samples_leaf | 10 |
| | min_samples_split | 10 |
| | loss | "ls" |
| | random_state | 42 |
| AdaBoost | n_estimators | 50 |
| | learning_rate | 1 |
| | loss | "linear" |
| | random_state | None |
| XGBoost | max_depth | 7 |
| | n_estimators | 200 |
| | learning_rate | 0.1 |
| | use_label_encoder | FALSE |
| | objective | "rank:pairwise" |

**Supplementary Table 4. Evaluation of Production Fitness Prediction Models.**
This analysis compares the performance of seven candidate machine learning models for predicting production fitness. The models evaluated include support vector regression (SVR), eXtreme gradient boosting (XGBoost), K-nearest neighbors (KNN), random forest (RF), gradient boosting (GB), Bayesian ridge, and AdaBoost.

| Production Fitness Prediction Performances | | | | | | | |
|---|---|---|---|---|---|---|---|
| Model | Fold | Pearson | Spearman | RMSE | $R^2$ | MedAE | C-index |
| SVR | 1 | 0.67186 | 0.64604 | 2.71153 | 0.39203 | 1.60187 | 0.73111 |
| | 2 | 0.66729 | 0.63975 | 2.70299 | 0.39346 | 1.59531 | 0.72890 |
| | 3 | 0.67724 | 0.65131 | 2.67003 | 0.41219 | 1.56678 | 0.73361 |
| | 4 | 0.66785 | 0.64055 | 2.71167 | 0.39097 | 1.61444 | 0.72878 |
| | 5 | 0.73609 | 0.71266 | 2.40424 | 0.52374 | 1.44232 | 0.75855 |
| KNN | 1 | 0.87509 | 0.84453 | 1.68340 | 0.76567 | 0.88892 | 0.82939 |
| | 2 | 0.87339 | 0.84330 | 1.69060 | 0.76273 | 0.89713 | 0.82842 |
| | 3 | 0.87842 | 0.84985 | 1.66475 | 0.77149 | 0.90161 | 0.83185 |
| | 4 | 0.87555 | 0.84572 | 1.67959 | 0.76635 | 0.90202 | 0.82929 |
| | 5 | 0.87447 | 0.84438 | 1.69028 | 0.76460 | 0.90754 | 0.82894 |
| Bayesian Ridge | 1 | 0.89077 | 0.86262 | 1.58052 | 0.79344 | 0.97228 | 0.83907 |
| | 2 | 0.88932 | 0.85955 | 1.58708 | 0.79089 | 0.98896 | 0.83686 |
| | 3 | 0.89262 | 0.86495 | 1.57012 | 0.79673 | 0.97297 | 0.84002 |
| | 4 | 0.89014 | 0.86076 | 1.58335 | 0.79236 | 0.97604 | 0.83803 |
| | 5 | 0.87452 | 0.85987 | 1.69392 | 0.76359 | 0.97905 | 0.83764 |
| AdaBoost | 1 | 0.88652 | 0.83480 | 2.17592 | 0.60849 | 1.79396 | 0.82220 |
| | 2 | 0.89245 | 0.82910 | 2.11035 | 0.63028 | 1.72841 | 0.81917 |
| | 3 | 0.89343 | 0.83689 | 2.09087 | 0.63954 | 1.71635 | 0.82308 |
| | 4 | 0.88568 | 0.82391 | 2.21855 | 0.59234 | 1.85756 | 0.81684 |
| | 5 | 0.89208 | 0.83244 | 2.06523 | 0.64858 | 1.67295 | 0.82176 |
| XGBoost | 1 | 0.94041 | 0.92047 | 2.97121 | 0.27000 | 2.26539 | 0.87966 |
| | 2 | 0.94070 | 0.92044 | 2.96489 | 0.27023 | 2.25937 | 0.87956 |
| | 3 | 0.94234 | 0.92281 | 2.97217 | 0.27163 | 2.25363 | 0.88105 |
| | 4 | 0.94046 | 0.91961 | 2.98362 | 0.26269 | 2.26622 | 0.87888 |
| | 5 | 0.93968 | 0.91933 | 2.97493 | 0.27081 | 2.28058 | 0.87873 |
| RF | 1 | 0.95142 | 0.92953 | 1.08827 | 0.90207 | 0.61561 | 0.88655 |
| | 2 | 0.95074 | 0.92850 | 1.09450 | 0.90055 | 0.62214 | 0.88572 |
| | 3 | 0.95191 | 0.93161 | 1.08680 | 0.90261 | 0.62526 | 0.88782 |

| | | | | | | | |
|---|---|---|---|---|---|---|---|
| | 4 | 0.95037 | 0.92816 | 1.09921 | 0.89993 | 0.63011 | 0.88523 |
| | 5 | 0.95004 | 0.92807 | 1.10567 | 0.89928 | 0.63268 | 0.88520 |
| GB | 1 | 0.95622 | 0.93581 | 1.01909 | 0.91412 | 0.58017 | 0.89177 |
| | 2 | 0.95550 | 0.93503 | 1.02546 | 0.91270 | 0.58800 | 0.89082 |
| | 3 | 0.95597 | 0.93658 | 1.02367 | 0.91360 | 0.58804 | 0.89199 |
| | 4 | 0.95585 | 0.93467 | 1.02238 | 0.91343 | 0.58842 | 0.89075 |
| | 5 | 0.95503 | 0.93484 | 1.03444 | 0.91183 | 0.59716 | 0.89070 |

**Supplementary Table 5. Evaluation of the Ly6a Binding Ability Prediction Model.**
This table summarizes the performance of seven candidate machine learning models for predicting Ly6a binding ability. The models evaluated include support vector regression (SVR), eXtreme gradient boosting (XGBoost), K-nearest neighbors (KNN), random forest (RF), gradient boosting (GB), Bayesian ridge, and AdaBoost.

| Ly6a Binding Ability Performances | | | | | | | |
|---|---|---|---|---|---|---|---|
| Model | Fold | Pearson | Spearman | RMSE | $R^2$ | MedAE | C-index |
| SVR | 1 | 0.61000 | 0.57704 | 2.48906 | 0.32567 | 1.42542 | 0.70013 |
| | 2 | 0.60559 | 0.57190 | 2.52726 | 0.31996 | 1.46323 | 0.69706 |
| | 3 | 0.60990 | 0.57689 | 2.50378 | 0.32363 | 1.45488 | 0.69900 |
| | 4 | 0.61355 | 0.57755 | 2.49584 | 0.33460 | 1.47437 | 0.69951 |
| | 5 | 0.63104 | 0.59733 | 2.44681 | 0.37505 | 1.44878 | 0.70754 |
| KNN | 1 | 0.71241 | 0.64265 | 2.14332 | 0.50000 | 1.28291 | 0.73059 |
| | 2 | 0.70882 | 0.64362 | 2.17502 | 0.49631 | 1.30410 | 0.73074 |
| | 3 | 0.70771 | 0.63719 | 2.16756 | 0.49309 | 1.30565 | 0.72797 |
| | 4 | 0.71264 | 0.64138 | 2.15803 | 0.50253 | 1.32544 | 0.72941 |
| | 5 | 0.70910 | 0.63805 | 2.19140 | 0.49871 | 1.35586 | 0.72799 |
| Bayesian Ridge | 1 | 0.72646 | 0.68555 | 2.08346 | 0.52753 | 1.27467 | 0.74653 |
| | 2 | 0.72925 | 0.68860 | 2.09705 | 0.53178 | 1.28845 | 0.74706 |
| | 3 | 0.72994 | 0.68876 | 2.08110 | 0.53272 | 1.28553 | 0.74725 |
| | 4 | 0.72703 | 0.68701 | 2.10087 | 0.52854 | 1.30095 | 0.74694 |
| | 5 | 0.70613 | 0.68343 | 2.19404 | 0.49750 | 1.32261 | 0.74513 |
| AdaBoost | 1 | 0.67863 | 0.65321 | 2.28953 | 0.42945 | 1.41373 | 0.72990 |
| | 2 | 0.68530 | 0.65129 | 2.30795 | 0.43287 | 1.40435 | 0.72841 |
| | 3 | 0.68672 | 0.65401 | 2.28196 | 0.43817 | 1.40889 | 0.72962 |
| | 4 | 0.68906 | 0.65139 | 2.28594 | 0.44181 | 1.44817 | 0.72914 |
| | 5 | 0.67283 | 0.65009 | 2.40062 | 0.39842 | 1.50741 | 0.72834 |
| XGBoost | 1 | 0.78857 | 0.73401 | 4.06534 | -0.79884 | 3.66384 | 0.77020 |
| | 2 | 0.78756 | 0.73516 | 4.08751 | -0.77890 | 3.65703 | 0.76997 |
| | 3 | 0.78996 | 0.73491 | 4.06488 | -0.78274 | 3.64276 | 0.77025 |
| | 4 | 0.79522 | 0.73972 | 4.10141 | -0.79686 | 3.68867 | 0.77290 |
| | 5 | 0.79077 | 0.73482 | 4.10900 | -0.76246 | 3.66750 | 0.77014 |
| RF | 1 | 0.81052 | 0.73526 | 1.78235 | 0.65423 | 1.08343 | 0.77282 |
| | 2 | 0.80869 | 0.73787 | 1.81131 | 0.65068 | 1.09923 | 0.77342 |
| | 3 | 0.80857 | 0.73315 | 1.79758 | 0.65136 | 1.10994 | 0.77149 |

| | | | | | | | |
|---|---|---|---|---|---|---|---|
| | 4 | 0.81478 | 0.74358 | 1.78631 | 0.65915 | 1.10718 | 0.77626 |
| | 5 | 0.81017 | 0.73678 | 1.82756 | 0.65135 | 1.13053 | 0.77301 |
| GB | 1 | 0.81061 | 0.72531 | 1.77580 | 0.65677 | 1.08904 | 0.76883 |
| | 2 | 0.80956 | 0.73072 | 1.79962 | 0.65518 | 1.08384 | 0.77106 |
| | 3 | 0.80695 | 0.72434 | 1.79937 | 0.65067 | 1.10404 | 0.76804 |
| | 4 | 0.81498 | 0.73328 | 1.77314 | 0.66416 | 1.09377 | 0.77224 |
| | 5 | 0.81053 | 0.72926 | 1.81284 | 0.65694 | 1.10737 | 0.77055 |

**Supplementary Table 6. Evaluation of Ly6c1 binding ability prediction models.** This table summarizes the performance of seven candidate machine learning models for predicting Ly6c1 binding ability. The models evaluated include support vector regression (SVR), eXtreme gradient boosting (XGBoost), K-nearest neighbors (KNN), random forest (RF), gradient boosting (GB), Bayesian ridge, and AdaBoost.

| Ly6c1 Binding Ability Performances | | | | | | | |
|---|---|---|---|---|---|---|---|
| Model | Fold | Pearson | Spearman | RMSE | $R^2$ | MedAE | C-index |
| SVR | 1 | 0.51954 | 0.50290 | 2.51427 | 0.17964 | 1.34660 | 0.67447 |
| | 2 | 0.50901 | 0.49666 | 2.54234 | 0.16551 | 1.34009 | 0.67224 |
| | 3 | 0.50220 | 0.48699 | 2.52967 | 0.16639 | 1.32983 | 0.66829 |
| | 4 | 0.52442 | 0.51223 | 2.55160 | 0.17291 | 1.35581 | 0.67796 |
| | 5 | 0.54642 | 0.53192 | 2.40825 | 0.25499 | 1.29684 | 0.68559 |
| KNN | 1 | 0.67399 | 0.61477 | 2.06400 | 0.44716 | 1.17711 | 0.71945 |
| | 2 | 0.67330 | 0.61144 | 2.06843 | 0.44763 | 1.15910 | 0.71823 |
| | 3 | 0.67274 | 0.60291 | 2.06309 | 0.44554 | 1.18628 | 0.71508 |
| | 4 | 0.67159 | 0.60882 | 2.09031 | 0.44493 | 1.19533 | 0.71773 |
| | 5 | 0.67114 | 0.60392 | 2.08219 | 0.44307 | 1.20054 | 0.71539 |
| Bayesian Ridge | 1 | 0.70603 | 0.67899 | 1.96606 | 0.49838 | 1.15721 | 0.74455 |
| | 2 | 0.70225 | 0.67035 | 1.98146 | 0.49310 | 1.16119 | 0.74080 |
| | 3 | 0.68670 | 0.66079 | 2.01494 | 0.47112 | 1.17947 | 0.73658 |
| | 4 | 0.70394 | 0.67597 | 1.99341 | 0.49520 | 1.18193 | 0.74298 |
| | 5 | 0.66907 | 0.66410 | 2.08038 | 0.44404 | 1.17512 | 0.73836 |
| AdaBoost | 1 | 0.63476 | 0.58316 | 2.26185 | 0.33609 | 1.42728 | 0.70496 |
| | 2 | 0.64676 | 0.59303 | 2.22821 | 0.35899 | 1.41354 | 0.70888 |
| | 3 | 0.62443 | 0.57921 | 2.25591 | 0.33705 | 1.40829 | 0.70326 |
| | 4 | 0.64302 | 0.59288 | 2.27384 | 0.34318 | 1.44472 | 0.70857 |
| | 5 | 0.63729 | 0.58655 | 2.26604 | 0.34038 | 1.43468 | 0.70602 |
| XGBoost | 1 | 0.77535 | 0.73524 | 3.41292 | -0.51159 | 2.99702 | 0.77092 |
| | 2 | 0.77622 | 0.73242 | 3.42236 | -0.51218 | 3.02954 | 0.76983 |
| | 3 | 0.76850 | 0.72347 | 3.41827 | -0.52212 | 3.02442 | 0.76560 |
| | 4 | 0.77750 | 0.73508 | 3.44997 | -0.51202 | 3.02345 | 0.77114 |
| | 5 | 0.77363 | 0.72733 | 3.44295 | -0.52272 | 3.04000 | 0.76747 |
| RF | 1 | 0.80000 | 0.73565 | 1.68294 | 0.63245 | 0.97301 | 0.77266 |
| | 2 | 0.80145 | 0.73002 | 1.68364 | 0.63403 | 0.97792 | 0.77030 |
| | 3 | 0.79629 | 0.72497 | 1.68984 | 0.62802 | 0.96935 | 0.76781 |

| | | | | | | | |
|---|---|---|---|---|---|---|---|
| | 4 | 0.80103 | 0.73471 | 1.69802 | 0.63372 | 0.98654 | 0.77251 |
| | 5 | 0.79904 | 0.72546 | 1.69431 | 0.63124 | 0.99744 | 0.76807 |
| GB | 1 | 0.80750 | 0.73279 | 1.63770 | 0.65194 | 0.96471 | 0.77186 |
| | 2 | 0.80720 | 0.72793 | 1.64297 | 0.65149 | 0.95963 | 0.76994 |
| | 3 | 0.80681 | 0.72737 | 1.63704 | 0.65090 | 0.95443 | 0.76965 |
| | 4 | 0.80714 | 0.73273 | 1.65658 | 0.65138 | 0.96228 | 0.77235 |
| | 5 | 0.80607 | 0.72304 | 1.65132 | 0.64971 | 0.97990 | 0.76793 |

**Supplementary Table 7. Hyperparameter Optimization of the BERT Architecture.** Optimization of the BERT encoder. Each set of optimization experiment results was repeated 5 times, and the mean and variance were calculated.

| Bert-based Pretraining Module | | | | |
|---|---|---|---|---|
| Hidden Size | Drop out | Attention Head | Layers | Accuracy |
| 256 | 0.1 | 16 | 1 | 8.520±1.300 |
| 256 | 0.1 | 16 | 2 | 8.533±1.651 |
| 256 | 0.1 | 16 | 3 | 8.151±1.433 |
| 256 | 0.1 | 16 | 4 | 10.479±1.399 |
| 256 | 0.3 | 16 | 1 | 7.963±1.460 |
| 256 | 0.4 | 16 | 1 | 7.552±1.412 |
| 256 | 0.5 | 16 | 1 | 7.471±1.316 |
| 256 | 0.6 | 16 | 1 | 6.857±1.238 |
| 256 | 0.7 | 16 | 1 | 6.356±0.978 |
| 256 | 0.8 | 16 | 1 | 5.867±0.769 |
| 256 | 0.1 | 4 | 1 | 8.092±1.702 |
| 256 | 0.1 | 8 | 1 | 8.428±1.479 |
| 256 | 0.1 | 32 | 1 | 8.153±1.699 |
| 256 | 0.1 | 64 | 1 | 8.340±1.573 |
| 512 | 0.1 | 16 | 1 | 9.370±1.201 |
| 1024 | 0.1 | 16 | 1 | 10.659±1.072 |
| 2048 | 0.1 | 16 | 1 | 11.308±0.732 |
| 4096 | 0.1 | 16 | 1 | 11.534±0.461 |

**Supplementary Table 8. Hyperparameter Optimization of the RoBERTa Architecture.** Optimization of the RoBERTa encoder. Each set of optimization experiment results was repeated 5 times, and the mean and variance were calculated.

| RoBERTa-based Pretraining Module | | | | |
|---|---|---|---|---|
| **Hidden Size** | **Drop out** | **Attention Head** | **Layers** | **Accuracy** |
| 256 | 0.1 | 16 | 1 | 10.282±1.791 |
| 256 | 0.1 | 16 | 2 | 10.593±1.304 |
| 256 | 0.1 | 16 | 3 | 10.518±1.691 |
| 256 | 0.1 | 16 | 4 | 10.443±1.010 |
| 256 | 0.3 | 16 | 1 | 9.301±2.271 |
| 256 | 0.4 | 16 | 1 | 8.870±2.396 |
| 256 | 0.5 | 16 | 1 | 8.950±2.225 |
| 256 | 0.6 | 16 | 1 | 8.860±2.442 |
| 256 | 0.7 | 16 | 1 | 8.375±2.377 |
| 256 | 0.8 | 16 | 1 | 7.913±2.385 |
| 256 | 0.1 | 4 | 1 | 9.656±2.313 |
| 256 | 0.1 | 8 | 1 | 9.582±2.383 |
| 256 | 0.1 | 32 | 1 | 9.503±2.521 |
| 256 | 0.1 | 64 | 1 | 9.622±2.337 |
| 512 | 0.1 | 16 | 1 | 10.804±1.615 |
| 1024 | 0.1 | 16 | 1 | 11.074±0.797 |
| 2048 | 0.1 | 16 | 1 | 11.241±0.491 |
| 4096 | 0.1 | 16 | 1 | 11.356±0.301 |

**Supplementary Table 9. Primer List**

| Name | Sequence 5’ to 3’ |
|---|---|
| Assembly-XbaI-F | CACTCATCGACCAATACTTGTACTATCTCT |
| Assembly-NNK-AAV9-588 | CCCGGAAGTATTCCTTGGTTTTGAACCCAACCGGTCTGCGCCTGTGCMNNMNNMNNMNNMNNMNNMNNTTGGGCACTCTGGTGGTT |
| ITR-F | CGGCCTCAGTGAGCGAGC |
| ITR-R | AGGAACCCCTAGTGATGG |
| NGS-1ST-F | ACACTCTTTCCCTACACGACGCTCTTCCGATCTACTAACCCGGTAGCAACGGAGT |
| NGS-1ST-R | GTGACTGGAGTTCAGACGTGTGCTCTTCCGATCTCTGCCAAACCATACCCGGAAGTATTCC |

**Supplementary Table 10. Model Architectures and Hyperparameters for Generative Training of Multifunctional Capsid Sequences**

This table summarizes the architectural and training parameters used for generative model development from scratch. SeqGAN was trained with the same parameters described in the main text, while the specific architectural configurations for the VAE and Diffusion models are listed here.

| **VAE Model Parameters** | |
|---|---|
| **Parameter** | **Value** |
| Vocab Size | 20 |
| hidden size | 1024 |
| z dim | 16 |
| num layers | 7 |
| learning rate | 5e-4 |
| beta | 0.1 |
| Optimizer | Adam |
| **Diffusion Model Parameters** | |
| **Parameter** | **Value** |
| Vocab Size | 20 |
| hidden size | 512 |
| num layers | 16 |
| num heads | 32 |
| dropout | 0.1 |
| diffusion steps | 500 |
| beta start | 1e-4 |
| beta end | 0.02 |

**Supplementary Table 11. Ranking Model Construction for hTfR1 Target Design.**

Considering the need for better convergence on the training set we constructed, we have partially optimized the original gradient boosting model by adopting LightGBM. The convergence results are presented in the table above.

| Task Name | RMSE | Pearsonr | Spearmanr | R2 | MedAE | MAE |
|---|---|---|---|---|---|---|
| Fitness Prediction | 1.387 | 0.878 | 0.876 | 0.768 | 0.861 | 1.077 |
| hTfR1 Prediction | 1.351 | 0.660 | 0.593 | 0.432 | 0.782 | 1.026 |

**Supplementary Table 12. Datasets construction of $Ref^+$ and $Ref^-$ for FE-X.**

To establish a clear benchmark for the FE-X architecture, we defined reference sets of capsid sequences from the AAV-588-OligoPool-lib1 dataset. The positive set ($Ref^+$) was established by applying stringent thresholds (production fitness > 2, hTfR1 $log_2$ enrichment > 2), yielding 27 high-quality sequences. In contrast, the negative set ($Ref^-$) consists of 48 sequences randomly sampled from those meeting opposite criteria (production fitness < −2, hTfR1 log2 enrichment < −2). This randomization ensures the negative set is representative and mitigates class imbalance by preventing overrepresentation of any single motif family. These curated reference sets underpin the contrastive learning in the Function Integration Stage and provide anchors for distance-based regularization in the Exploration Stage.

| $Ref^+$ | | | | | | | |
|---|---|---|---|---|---|---|---|
| No. | Sequence | No. | Sequence | No. | Sequence | No. | Sequence |
| 1 | KYHAMSG | 8 | LSRISIN | 15 | YLKSQTF | 22 | YSRGGPN |
| 2 | TYHKSTV | 9 | VYTRSQP | 16 | YAKSSPN | 23 | LHRLSSN |
| 3 | IYSKSVI | 10 | VYTRTMS | 17 | YARSGTN | 24 | LHRLQLG |
| 4 | LHLRGVN | 11 | YHRLSNN | 18 | YVKSNTQ | 25 | LHRSLPD |
| 5 | LHKMGLN | 12 | YSRIGPN | 19 | YSRNAPN | 26 | LARSGTN |
| 6 | LHKLAYS | 13 | YSRLNKD | 20 | YSRLNMN | 27 | LSRTGVN |
| 7 | LHRLSEH | 14 | YHRLGDD | 21 | YSRLNLD | | |
| $Ref^-$ | | | | | | | |
| No. | Sequence | No. | Sequence | No. | Sequence | No. | Sequence |
| 1 | TSTGPWP | 13 | SLSPLWP | 25 | TKAYLAF | 37 | RPLLVRV |
| 2 | HVALRLH | 14 | EWLHKTQ | 26 | FLELARR | 38 | RSKLVLP |
| 3 | EHVYQTY | 15 | PAVNAYG | 27 | RSKIEAW | 39 | TVFRVGF |
| 4 | VPPRNAK | 16 | LSLVRRL | 28 | SSVRRPV | 40 | ASMKTWQ |
| 5 | PSVWPMD | 17 | WPSELPY | 29 | RGSFTML | 41 | ASLKSFF |
| 6 | TLEGKVT | 18 | KLGLKTT | 30 | RKTLSMP | 42 | VPMTGRK |
| 7 | KLGSVWI | 19 | PSPYVIV | 31 | LFIVAKG | 43 | IELWGNR |
| 8 | KVFVHIP | 20 | MSVFHPV | 32 | ARLDFFP | 44 | LFNSNRF |
| 9 | SLPMMFR | 21 | SKFAVQR | 33 | CEHGAMA | 45 | RGSFSLI |
| 10 | GTWPAIR | 22 | QSEQVGQ | 34 | TTSPWTA | 46 | LVDPLSC |
| 11 | SLSFSYM | 23 | PVDTPFE | 35 | DVIRTYR | 47 | CNDRPSG |
| 12 | LVRAYLP | 24 | ENTPYDI | 36 | LGMIRKT | 48 | ERGRTAL |

**Supplementary Table 13. Top 200 Sequences designed by ALICE-X and positive control ranked by Pareto Rank.**

| Sequence | hTFRC KI mice Brain log2 Enrichment | WT mice Brain log2 Enrichment | Production Fitness | Pareto Rank |
|---|---|---|---|---|
| VYTRSAG | -0.272 | -1.823 | 2.194 | 1 |
| QYIKSNT | 0.615 | -3.157 | 1.746 | 1 |
| VYTRAAT | 0.943 | -3.314 | 1.257 | 1 |
| VYTKVQG | 1.009 | -3.963 | 1.205 | 1 |
| VWVKSQP | 1.700 | -3.494 | 1.089 | 1 |
| QYIKSVP | 2.565 | -2.869 | 0.942 | 1 |
| QYTKSIT | 2.346 | -3.294 | 0.623 | 1 |
| VYTKSDT | 3.724 | -2.734 | 0.340 | 1 |
| TYTKSEI | 6.184 | -3.506 | 0.267 | 1 |
| QWVKSPT | 5.411 | -5.876 | -0.691 | 1 |
| TYTKSSI | 3.908 | 0.563 | 1.571 | 2 |
| IYTKSQP | 1.454 | -2.263 | 1.369 | 2 |
| TYMKSQI | 1.133 | -2.933 | 1.256 | 2 |
| VYTKSLS | 1.399 | -3.058 | 1.101 | 2 |
| QYTKSSS | 2.164 | -2.391 | 1.077 | 2 |
| TYTRSVA | 1.898 | -3.285 | 1.070 | 2 |
| VYTRSTP | 2.751 | -2.515 | 0.404 | 2 |
| QWAKSVP | 3.296 | -3.005 | 0.231 | 2 |
| QWTKSLA | 1.915 | -5.292 | 0.209 | 2 |
| AYIKSVS | 3.628 | -4.435 | 0.194 | 2 |
| VYTKTIS | 1.828 | -6.693 | 0.177 | 2 |
| VYTRSLA | 2.564 | -6.040 | -0.074 | 2 |
| QYTKSID | 5.475 | -3.203 | -0.360 | 2 |
| NWVKSST | 5.232 | -3.636 | -0.461 | 2 |
| VYTRVMT | 3.655 | -5.742 | -0.499 | 2 |
| QYTRSMQ | 4.269 | -5.501 | -1.646 | 2 |
| QYTKSIQ | 0.117 | 0.171 | 1.399 | 3 |
| TYTRSQP | 0.444 | -1.385 | 1.312 | 3 |
| MYTRSES | -0.265 | -2.318 | 1.301 | 3 |
| VYTRTEN | 1.231 | -2.644 | 1.199 | 3 |
| VYTKSVV | 0.679 | -3.505 | 1.079 | 3 |
| MYTKSIS | 1.945 | -2.728 | 1.063 | 3 |
| VYTKSAT | 1.666 | -3.410 | 1.021 | 3 |
| QYTKSIV | 2.501 | -2.679 | 0.772 | 3 |
| PWVKSNP | 1.795 | -3.455 | 0.346 | 3 |
| AWVKSVP | 2.982 | -2.832 | 0.220 | 3 |
| VYTRNIS | 1.641 | -4.450 | 0.194 | 3 |
| VYTRSLE | 2.427 | -4.368 | 0.091 | 3 |
| VYTKMVA | 2.069 | -4.995 | 0.072 | 3 |

| | | | | |
|---|---|---|---|---|
| VYTKSYQ | 3.712 | -3.421 | -0.066 | 3 |
| VYTRSDL | 3.812 | -3.762 | -0.136 | 3 |
| SFVKSEI | 4.043 | -3.597 | -0.190 | 3 |
| VWTKSLT | 2.871 | -5.483 | -0.287 | 3 |
| VYMRSED | 4.403 | -4.483 | -0.715 | 3 |
| VYTRSDV | -4.794 | 0.111 | 1.256 | 4 |
| QYTKSNN | 0.478 | 0.572 | 1.235 | 4 |
| TYTRSIA | 3.184 | 0.001 | 1.151 | 4 |
| TFVKSQI | 0.373 | -3.212 | 1.113 | 4 |
| QYTKSQA | 2.388 | -1.980 | 1.050 | 4 |
| VYQRSQT | 1.365 | -3.232 | 1.024 | 4 |
| VYTRSTD | 1.001 | -3.993 | 0.945 | 4 |
| QYTRSAP | 2.436 | -2.726 | 0.604 | 4 |
| VYTRTMA | 1.317 | -3.865 | 0.198 | 4 |
| VYTKGES | 2.804 | -2.540 | 0.167 | 4 |
| YVLKSQA | 4.228 | -1.551 | 0.098 | 4 |
| VYTKTDL | 3.033 | -2.854 | 0.010 | 4 |
| NWVKSIN | 2.230 | -3.841 | 0.003 | 4 |
| AWVKSGV | 3.461 | -3.347 | -0.007 | 4 |
| MVYKSQV | 4.135 | -2.796 | -0.269 | 4 |
| VYTRAMS | 3.353 | -3.808 | -0.347 | 4 |
| VYTRSQM | 5.531 | -2.376 | -0.386 | 4 |
| VWTKSIP | 1.825 | -6.253 | -0.395 | 4 |
| VYTRVVT | 2.995 | -5.272 | -0.747 | 4 |
| VYTRSLL | 2.842 | -5.790 | -1.457 | 4 |
| VYTKGMS | -2.988 | -2.736 | 1.174 | 5 |
| TYTRSSE | 1.725 | -2.205 | 1.039 | 5 |
| VYTKSML | 0.727 | -3.593 | 0.960 | 5 |
| VYTKGIM | 1.769 | -2.726 | 0.946 | 5 |
| SYTRTNV | 0.891 | -3.665 | 0.645 | 5 |
| VYTKTHM | 1.680 | -3.198 | 0.542 | 5 |
| VYTKTMM | 2.280 | -2.863 | 0.526 | 5 |
| QWVKSNA | 1.404 | -3.749 | 0.151 | 5 |
| VYMKSVT | 1.313 | -4.109 | 0.052 | 5 |
| VYTKTMQ | 2.278 | -3.168 | 0.029 | 5 |
| QYTKSLN | 2.764 | -2.714 | -0.021 | 5 |
| VYIKSDP | 3.248 | -3.440 | -0.023 | 5 |
| VYTKMMT | 2.205 | -4.515 | -0.168 | 5 |
| VYIRTNS | 1.988 | -5.614 | -0.423 | 5 |
| TYTRSIV | 2.228 | -5.855 | -0.884 | 5 |
| IYTRSEV | -5.237 | -2.127 | 1.123 | 6 |
| QYTKSIP | 0.434 | -2.073 | 1.023 | 6 |
| QYIKSLG | -0.220 | -2.908 | 0.986 | 6 |

| | | | | |
|---|---|---|---|---|
| VYPRSEA | 0.656 | -2.769 | 0.980 | 6 |
| VYTRVSA | 1.920 | -2.478 | 0.920 | 6 |
| VYTKSAP | 1.824 | -2.866 | 0.536 | 6 |
| IYTKSLS | 1.633 | -3.205 | 0.502 | 6 |
| AFVKSQA | 1.562 | -3.377 | 0.496 | 6 |
| IYTRSEP | 1.460 | -3.532 | 0.433 | 6 |
| QYTRSQG | 2.522 | -2.557 | 0.396 | 6 |
| IYTKSVN | 2.213 | -2.907 | 0.373 | 6 |
| VYTRSID | 2.839 | -3.583 | -0.045 | 6 |
| QWVKSTQ | 0.020 | -6.664 | -0.233 | 6 |
| QWVKSHP | 3.725 | -2.976 | -0.345 | 6 |
| VYTKSDE | 4.990 | -1.720 | -0.462 | 6 |
| IYTRSVS | 3.740 | -3.579 | -0.497 | 6 |
| SFVKSQF | 4.379 | -3.106 | -0.630 | 6 |
| VYTRSSM | 4.639 | -3.060 | -0.923 | 6 |
| MYTRSQI | 3.535 | -4.448 | -1.676 | 6 |
| PYTKSQI | -0.507 | -3.427 | 0.974 | 7 |
| QLVKSQI | 0.576 | -2.350 | 0.970 | 7 |
| VYTKGLS | 0.841 | -2.429 | 0.965 | 7 |
| VYTRGVA | 1.611 | -1.688 | 0.924 | 7 |
| TYTKSPV | 1.334 | -3.043 | 0.763 | 7 |
| VYTKSLM | 1.236 | -3.631 | 0.492 | 7 |
| VYMKSQP | 2.483 | -2.578 | 0.283 | 7 |
| AFVKSQV | 2.753 | -2.779 | -0.079 | 7 |
| VYTRTLD | 1.879 | -4.410 | -0.124 | 7 |
| SYTRSLN | 4.108 | -2.492 | -0.242 | 7 |
| VYTRIQE | 4.756 | -1.851 | -0.363 | 7 |
| QYTKSII | 2.894 | -3.805 | -0.494 | 7 |
| VYTRSIL | 3.272 | -3.574 | -0.533 | 7 |
| SFVKSEL | 1.389 | -6.075 | -0.724 | 7 |
| VYTRIQV | 5.186 | -2.347 | -1.170 | 7 |
| QYTRNQS | 4.630 | -2.914 | -1.425 | 7 |
| QYTRTSS | 4.483 | -3.329 | -1.886 | 7 |
| YVHKSQV | 0.041 | -2.737 | 0.918 | 8 |
| VYNRSVD | 0.233 | -2.655 | 0.778 | 8 |
| VYTRAVP | 0.451 | -2.522 | 0.763 | 8 |
| QFVKSQT | 1.551 | -2.055 | 0.734 | 8 |
| VYTRSEI | 1.533 | -2.218 | 0.695 | 8 |
| SYTRSQG | 2.725 | -1.293 | 0.686 | 8 |
| VYTKGIT | 0.926 | -3.172 | 0.620 | 8 |
| TYQRSET | 1.608 | -2.505 | 0.557 | 8 |
| VYTRTTS | 2.378 | -2.396 | 0.480 | 8 |
| VYTKSFN | 0.411 | -4.440 | 0.412 | 8 |

| | | | | |
|---|---|---|---|---|
| QYTRSAA | 0.699 | -4.269 | 0.207 | 8 |
| QLVKSQY | 1.753 | -3.218 | -0.118 | 8 |
| QLIKSQV | 2.789 | -2.618 | -0.155 | 8 |
| TFVKSTI | 3.805 | -2.191 | -0.160 | 8 |
| VFTRSQS | 3.708 | -2.308 | -0.182 | 8 |
| VYTRIDN | 0.663 | -5.376 | -0.184 | 8 |
| TYTRSLN | 1.767 | -4.343 | -0.244 | 8 |
| QWVKSLP | 2.695 | -3.721 | -0.284 | 8 |
| QWVKSQG | 3.654 | -2.938 | -0.541 | 8 |
| TWVKSLE | 3.244 | -3.596 | -0.705 | 8 |
| VYTRSMM | 2.532 | -4.627 | -0.835 | 8 |
| QYTRSVP | 1.908 | 0.139 | 0.915 | 9 |
| VYTRVQG | 0.052 | -2.352 | 0.893 | 9 |
| VYTRTVV | -0.028 | -2.788 | 0.850 | 9 |
| QYIKSLP | 0.137 | -3.331 | 0.715 | 9 |
| VYTKTNV | 1.298 | -2.188 | 0.685 | 9 |
| TYTKSTI | 0.796 | -3.090 | 0.673 | 9 |
| TYTRSVG | 1.103 | -2.791 | 0.672 | 9 |
| QYTKSVI | 1.713 | -2.195 | 0.627 | 9 |
| VYTRADS | 0.232 | -3.766 | 0.595 | 9 |
| QVLKSQY | 1.037 | -2.964 | 0.571 | 9 |
| VYTRGNN | 2.580 | -1.762 | 0.434 | 9 |
| VYTKSPV | 0.625 | -3.910 | 0.360 | 9 |
| NWVKSQN | 1.636 | -3.152 | 0.336 | 9 |
| VYTRAVS | 2.077 | -2.859 | -0.023 | 9 |
| VYTKSVY | 1.381 | -4.006 | -0.172 | 9 |
| VYTRGIV | 3.128 | -2.263 | -0.199 | 9 |
| VYTKTIH | 0.649 | -5.381 | -0.241 | 9 |
| VYTKLVA | 0.683 | -5.656 | -0.321 | 9 |
| YSRIGPN | 3.053 | -3.357 | -0.468 | 9 |
| VYTRSHD | 2.029 | -4.387 | -0.733 | 9 |
| QYTRSAN | 2.887 | -3.731 | -0.837 | 9 |
| VWTKSLQ | 3.362 | -3.788 | -0.865 | 9 |
| SYTKSQI | -6.193 | -2.840 | 0.882 | 10 |
| QYTKSYS | -5.597 | -6.445 | 0.845 | 10 |
| QYMKSQV | -0.505 | -2.256 | 0.823 | 10 |
| SYTRSLP | -0.407 | -3.351 | 0.698 | 10 |
| VYTKSTI | 0.806 | -2.496 | 0.674 | 10 |
| IYTRSVQ | 0.518 | -3.077 | 0.591 | 10 |
| QYTRSQE | 1.545 | -2.239 | 0.580 | 10 |
| TYTKSQL | 1.703 | -2.104 | 0.575 | 10 |
| QYTRSVV | 0.443 | -3.487 | 0.571 | 10 |
| SFVKSSV | 1.752 | -2.376 | 0.378 | 10 |

| | | | | |
|---|---|---|---|---|
| VYTRAQP | 4.493 | 0.295 | 0.361 | 10 |
| VYTKSMS | 0.470 | -3.759 | 0.349 | 10 |
| TYTKSAQ | 2.280 | -1.972 | 0.339 | 10 |
| QYTRLQP | 1.825 | -2.598 | 0.308 | 10 |
| QYTRSEV | 2.777 | -1.925 | 0.193 | 10 |
| MYTRSNV | 1.639 | -3.129 | 0.094 | 10 |
| QWTKSID | -0.129 | -5.140 | -0.184 | 10 |
| VYTKSVM | 3.103 | -1.997 | -0.198 | 10 |
| VYTRGAP | 2.963 | -2.633 | -0.296 | 10 |
| VYTRVEN | 3.540 | -2.081 | -0.307 | 10 |
| VYTKSIT | 0.437 | -5.321 | -0.330 | 10 |
| VYTKTLV | 3.104 | -2.666 | -0.369 | 10 |
| TYTRSLT | 2.118 | -3.741 | -0.378 | 10 |
| VYTKTLM | 2.345 | -3.798 | -0.515 | 10 |
| VWVKSST | 1.541 | -4.751 | -0.615 | 10 |
| IYTRSQI | 2.671 | -3.656 | -0.818 | 10 |
| VYTRVLS | 2.422 | -4.389 | -0.953 | 10 |
| VYTKSHN | 2.643 | -4.390 | -1.414 | 10 |
| QWTKSTV | -2.556 | -2.911 | 0.822 | 11 |
| QYTRLSE | -1.200 | -2.478 | 0.708 | 11 |
| VYTKSMV | -0.893 | -2.592 | 0.684 | 11 |
| VYTRTNN | -0.174 | -2.779 | 0.570 | 11 |
| QWTKSQA | 1.029 | -2.239 | 0.554 | 11 |
| VYTRTTP | 0.870 | -2.528 | 0.507 | 11 |
| VYTRSEN | 1.616 | -1.836 | 0.457 | 11 |
| VYTRIAP | 1.268 | -2.515 | 0.402 | 11 |
| TFVKSQM | 1.143 | -2.652 | 0.401 | 11 |
| QWVKSIP | 1.774 | -2.309 | 0.311 | 11 |
| NWVKSNP | 1.600 | -2.664 | 0.182 | 11 |
| VYTKVSQ | 1.506 | -2.824 | 0.139 | 11 |
| VYIKSTN | 2.454 | -2.222 | 0.130 | 11 |
| VYTRTSM | 2.681 | -2.041 | 0.063 | 11 |

**Supplementary Table 14. Viral Package Prediction Results and Viral Titer.**

| Capsids | Sequence | Predicted Production Fitness | Average Experimental Titer (vg/ml) | High/Normal/Low |
|---|---|---|---|---|
| AAV9 | SAQAQAQ | / | $2.41 \times 10^{13}$ vg/ml | Normal |
| AAV.ALICE-N1 | LRSGYSS | 0.93 | $9.59 \times 10^{10}$ vg/ml | Low |
| AAV.ALICE-N2 | MMRGYSS | -0.49 | $5.55 \times 10^{13}$ vg/ml | High |
| AAV.ALICE-N3 | MRPGYSS | 1.85 | $6.00 \times 10^{12}$ vg/ml | Normal |
| AAV.ALICE-N4 | MRSGYSS | 0.94 | $6.02 \times 10^{12}$ vg/ml | Normal |
| AAV.ALICE-N5 | DRYGYSS | -0.07 | $9.59 \times 10^{10}$ vg/ml | Low |
| AAV.ALICE-N6 | TGFGYSS | -0.28 | $4.95 \times 10^{13}$ vg/ml | High |
| AAV.ALICE-N7 | TNYGYSS | 0.10 | $8.30 \times 10^{12}$ vg/ml | Normal |
| AAV.ALICE-N8 | WERGYSS | 0.24 | $3.86 \times 10^{13}$ vg/ml | High |
| AAV.ALICE-N9 | IRQGYSQ | 0.67 | $5.92 \times 10^{13}$vg/ml | High |
| AAV.ALICE-H1 | TYTKSEI | 0.95 | $4.05 \times 10^{12}$ vg/ml | Normal |
| AAV.ALICE-H2 | VYTKSDT | 1.08 | $5.58 \times 10^{12}$ vg/ml | Normal |
| AAV.ALICE-H3 | QYTKSIT | 2.26 | $3.33 \times 10^{13}$ vg/ml | High |

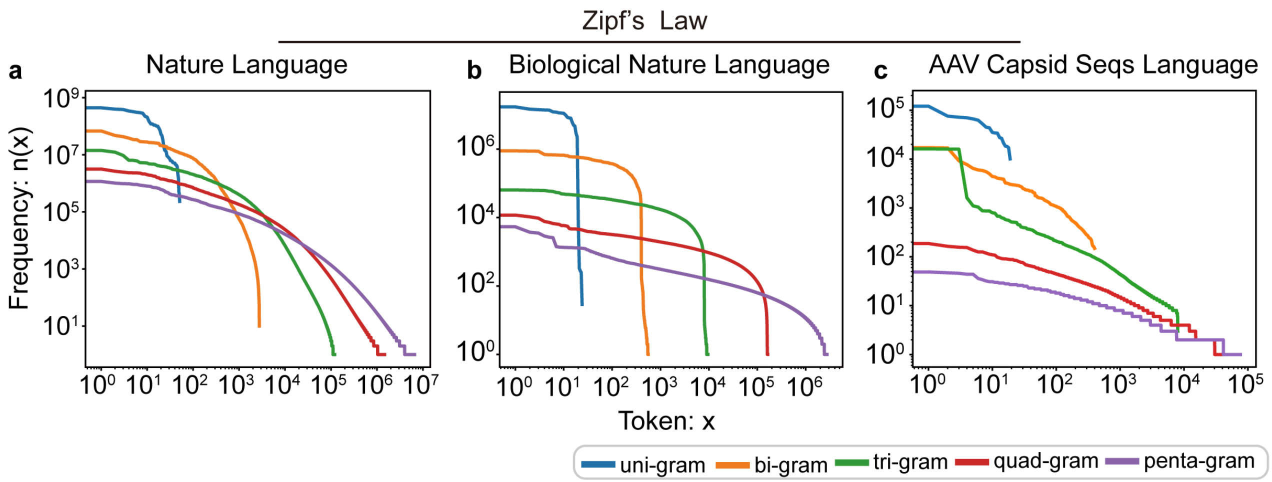

**Supplementary Fig. 1 | Distribution of Zipf's law across different language domains.**

**a-c,** Illustration of Zipf's law in human natural language (**a**), amino acid-based biological language (**b**), and AAV capsid sequences (**c**). The AAV capsid data used here were sampled from the AAV Capsid Library dataset recorded in Supplementary Table 1.

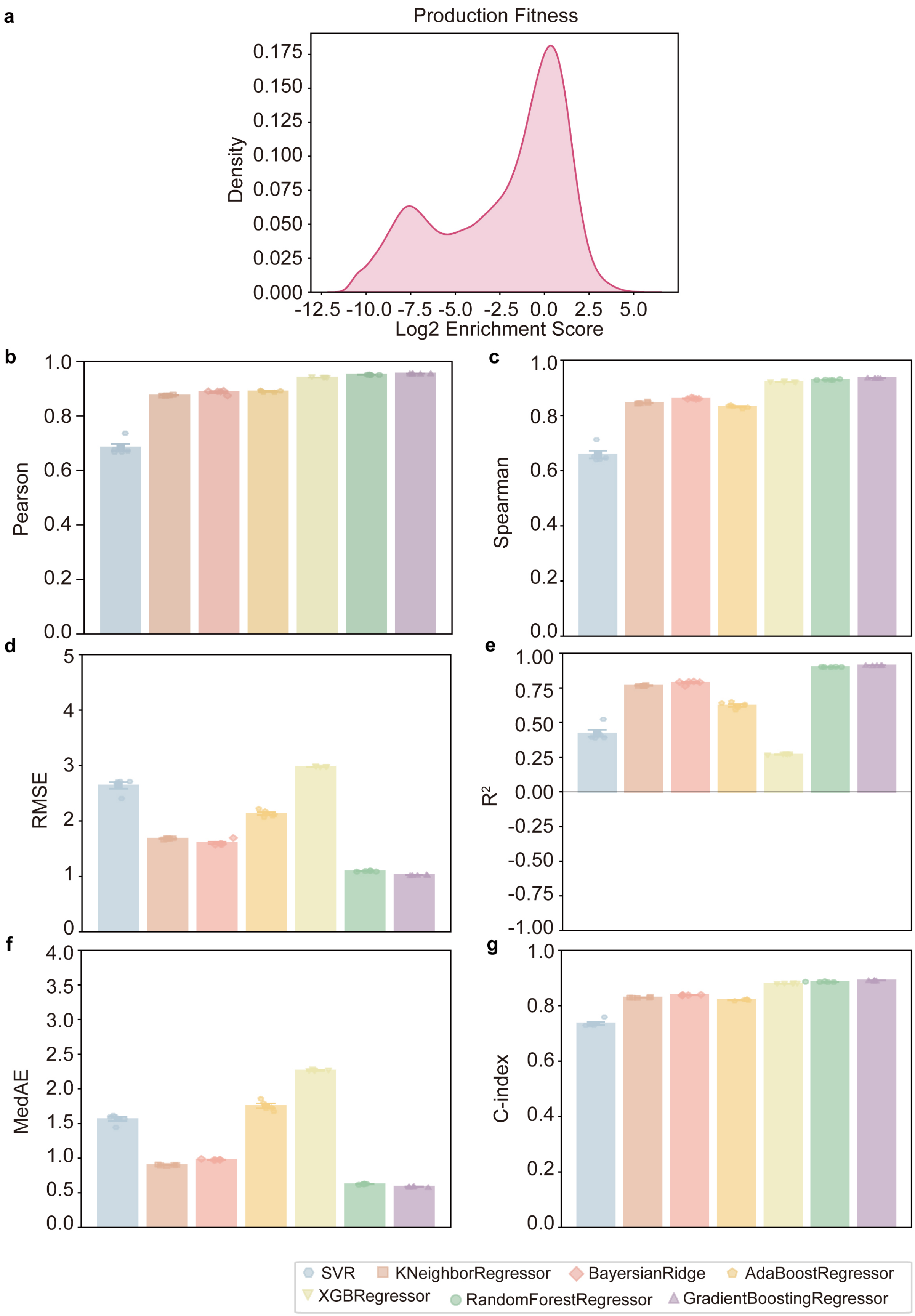

1572

1573 **Supplementary Fig. 2 | The performance of the models in evaluating AAV production fitness.**

**a,** Distribution of training data for the AAV production fitness assessment models ($\boldsymbol{D_{Production\ Fitness}}$). **b-g,** Evaluation results of seven candidate machine learning models for predicting production fitness via six evaluation indicators: Pearson correlation (b), Spearman's rank correlation (c), root mean square error (RMSE) (d), coefficient of determination ($R^2$) (e), median absolute error (MedAE) (f), and concordance index (g).

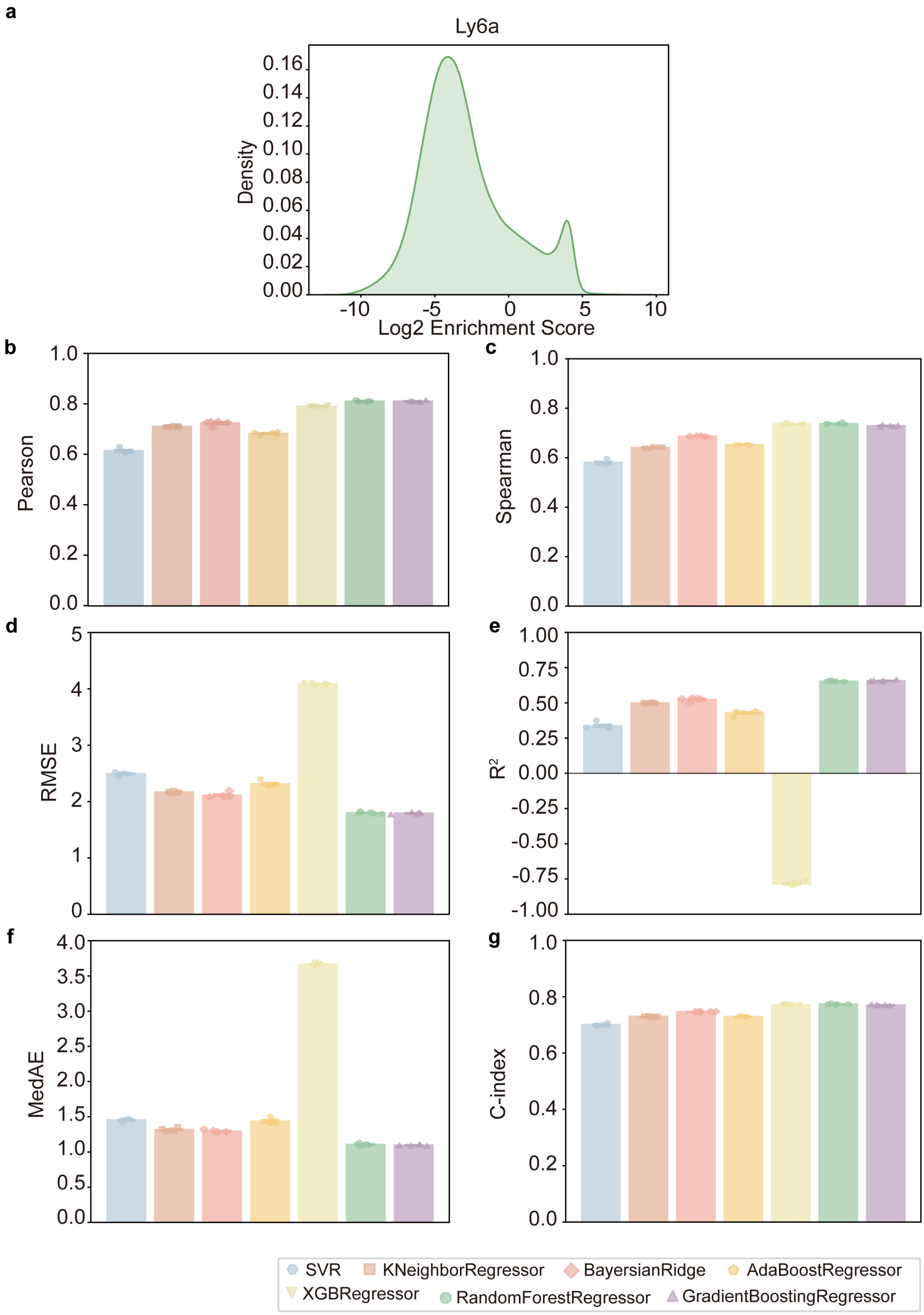

**Supplementary Fig. 3 | The performance of the models in evaluating the binding capacity of AAV variants and Ly6a target proteins.**

**a,** Distribution of training data for the evaluation model of AAV and Ly6a target protein binding capacity

($D_{Ly6a}$).

**b-g,** Evaluation results of seven candidate machine learning models for predicting binding capacity via six evaluation indicators: Pearson correlation (b), Spearman's rank correlation (c), root mean square error (RMSE) (d), coefficient of determination ($R^2$) (e), median absolute error (MedAE) (f), and concordance index (g).

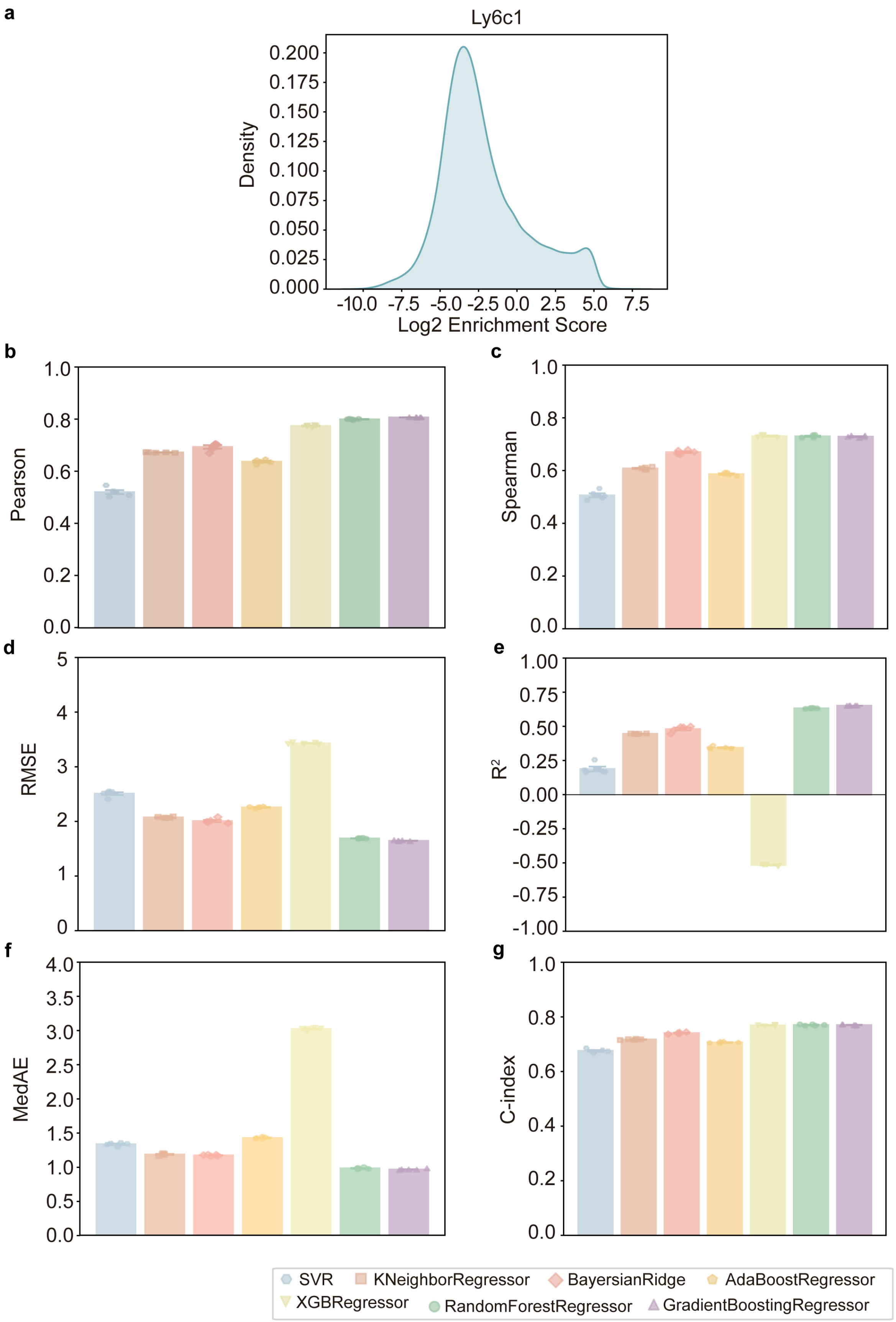

**Supplementary Fig. 4 | The performance of the models in evaluating the binding capacity of AAV variants and Ly6c1 target proteins.**

**a,** Distribution of training data for the evaluation model of AAV and Ly6c1 target protein binding capacity ($\boldsymbol{D_{Ly6c1}}$).

**b-g,** Evaluation results of seven candidate machine learning models for predicting binding capacity via six evaluation indicators: Pearson correlation (b), Spearman's rank correlation (c), root mean square error (RMSE) (d), coefficient of determination ($R^2$) (e), median absolute error (MedAE) (f), and concordance index (g).

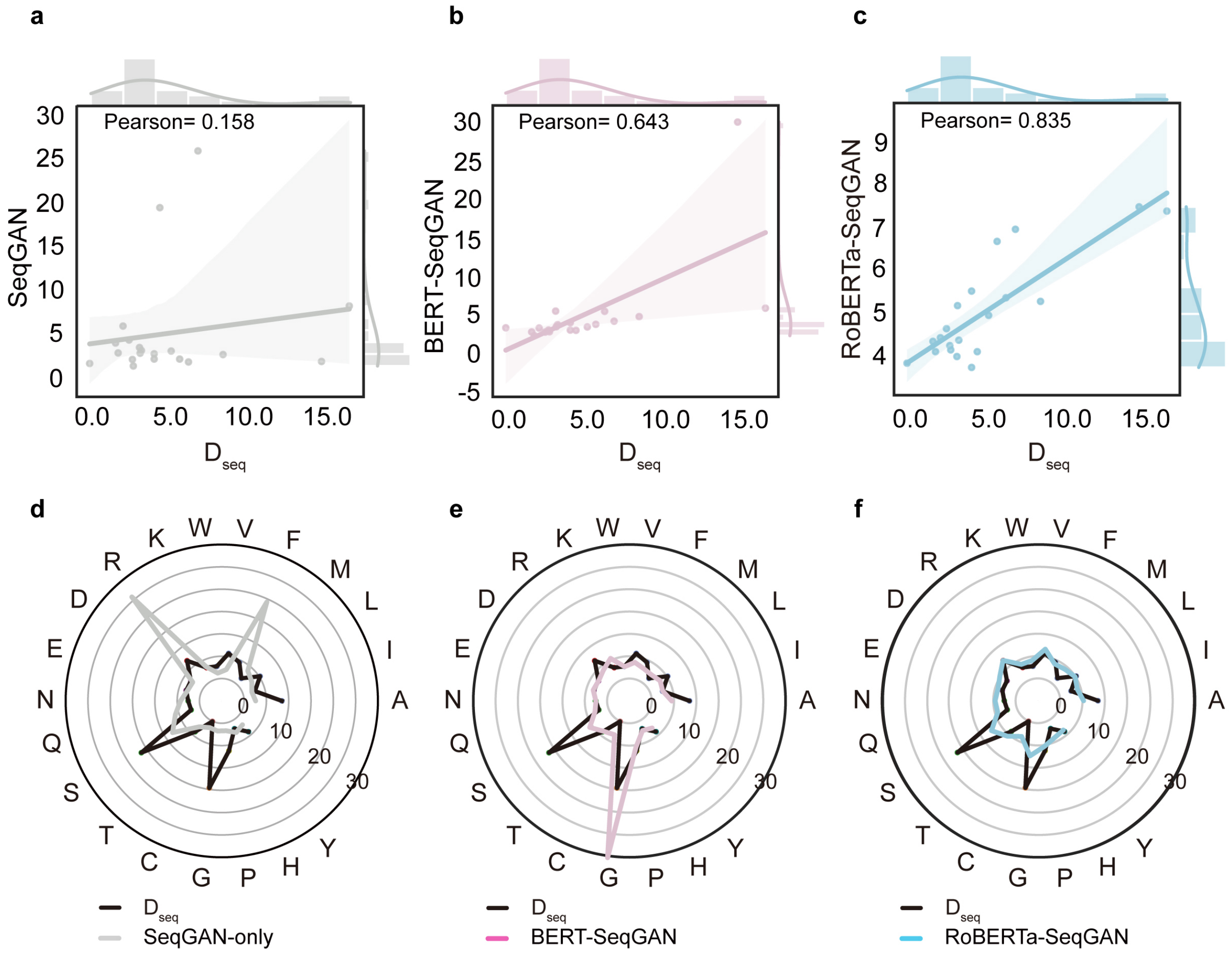

**Supplementary Fig. 5 | Amino acid grammar semantic analysis of sequences designed through pretraining and semantic tuning strategies**

**a-c,** The plots show the Pearson correlation of the amino acid propensities of capsid sequences designed by three different generative architectures with those of the sequences in the semantic tuning dataset ($\boldsymbol{D_{seq}}$). The correlations for SeqGAN-only (a), BERT-SeqGAN (b), and RoBERTa-SeqGAN (c) are 0.158, 0.643, and 0.835, respectively. In the categories of SeqGAN-only, BERT-SeqGAN, and RoBERTa-SeqGAN, a random sample of 1000 sequences was extracted for analysis**.**

**d-f,** The diagrams show the amino acid frequency distribution of capsid sequences designed by three different generative architectures, including SeqGAN-only (d), BERT-SeqGAN (e), and RoBERTa-SeqGAN (f), in comparison to the sequences in the semantic tuning dataset ($\boldsymbol{D_{seq}}$). **In the categories of** SeqGAN-only, BERT-SeqGAN, and RoBERTa-SeqGAN, a random sample of 1000 sequences was extracted for analysis**.**

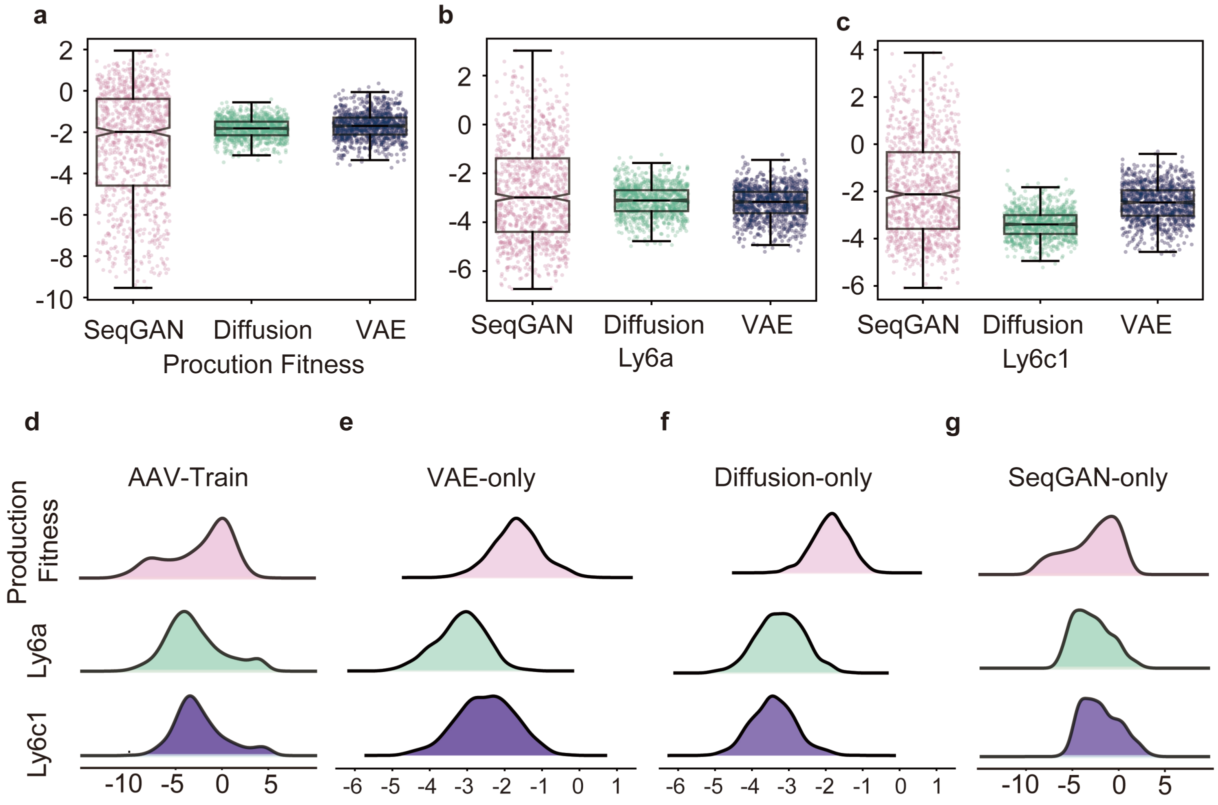

**Supplementary Fig. 6 | Comparative performance of generative models for capsid sequence design.**

**a~c,** Performance comparison of VAE-only, Diffusion-only, and SeqGAN-only models in capsid generation. For each model, 1,000 generated capsids were evaluated for production fitness (a), Ly6a binding capability (b), and Ly6c1 binding capability (c) using the evaluation model.

**d,** Distributions of fitness, Ly6a enrichment, and Ly6c1 enrichment in the training set.

**e~g,** Distributions of the predicted values of fitness, Ly6a enrichment, and Ly6c1 enrichment for capsids generated by the VAE-only (e), Diffusion-only (f), and SeqGAN-only (g) models.

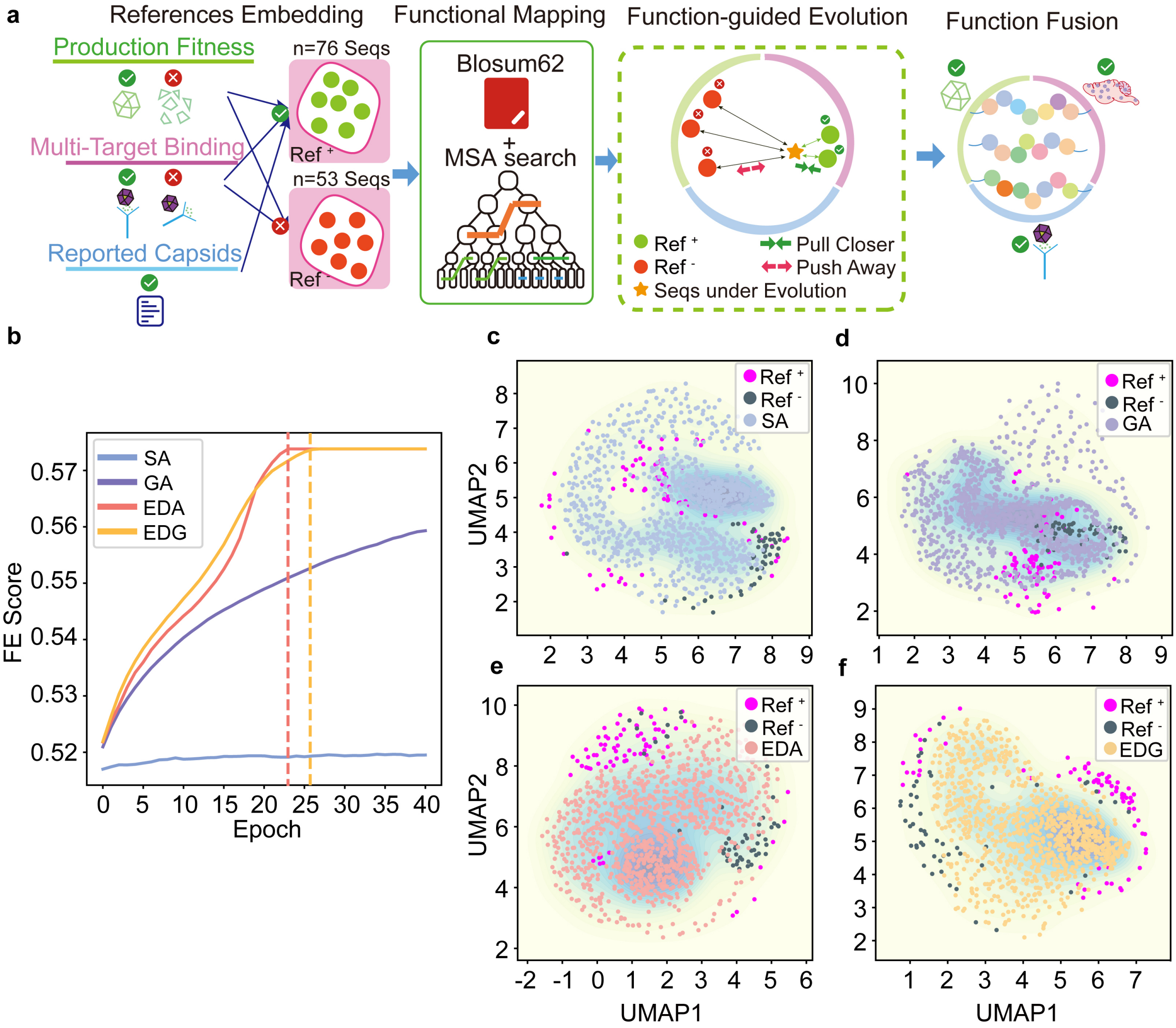

**Supplementary Fig. 7 | Operating mechanism and functionality of the function-guided evolution (FE) module.**

**a,** Positive and negative samples ($\boldsymbol{Ref^+}$ and $\boldsymbol{Ref^-}$) for the FE module were chosen on the basis of viability, multitargeting, and reported capsid (76 $\boldsymbol{Ref^+}$ sequences, 53 $\boldsymbol{Ref^-}$ sequences). The sequences filtered from RankFiltPro evolve toward the features of positive samples and away from those of negative samples in a model employing a heuristic algorithm coupled with contrastive learning (Multiple Sequence Alignment, MSA) and reinforcement learning. This enables the fusing of multiple functions into a single sequence.

**b,** The plot illustrates the FE scores of sequences engineered by the SA, GA, EDA, or EDG model at different epochs on their progress toward the $\boldsymbol{Ref^+}/\boldsymbol{Ref^-}$ sample.

**c-f,** The plots show the functional distributions of sequences that have been engineered via SA (c), GA (d), EDA (e), and EDG (f) compared with those of the sequences from the $\boldsymbol{Ref^+}$ and $\boldsymbol{Ref^-}$ datasets within a multidimensional space.

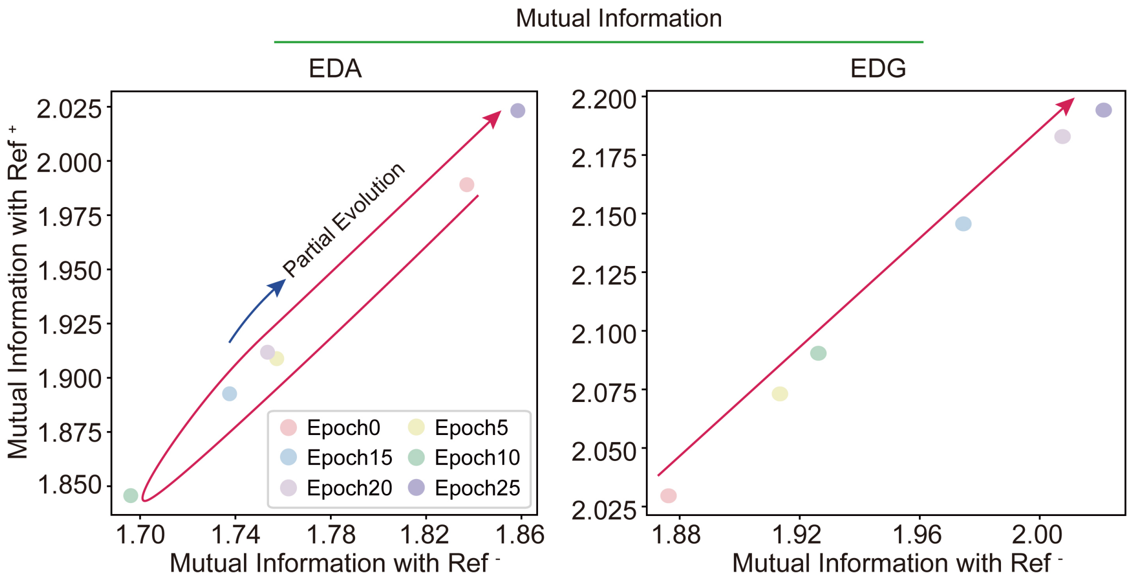

**Supplementary Fig. 8 | The evolutionary dynamics of the mutual information of the sequences designed by the FE module with the EDA or EDG model.**

The plots illustrate the changes in mutual information during the evolutionary process within the FE module when the EDA or EDG model is employed.

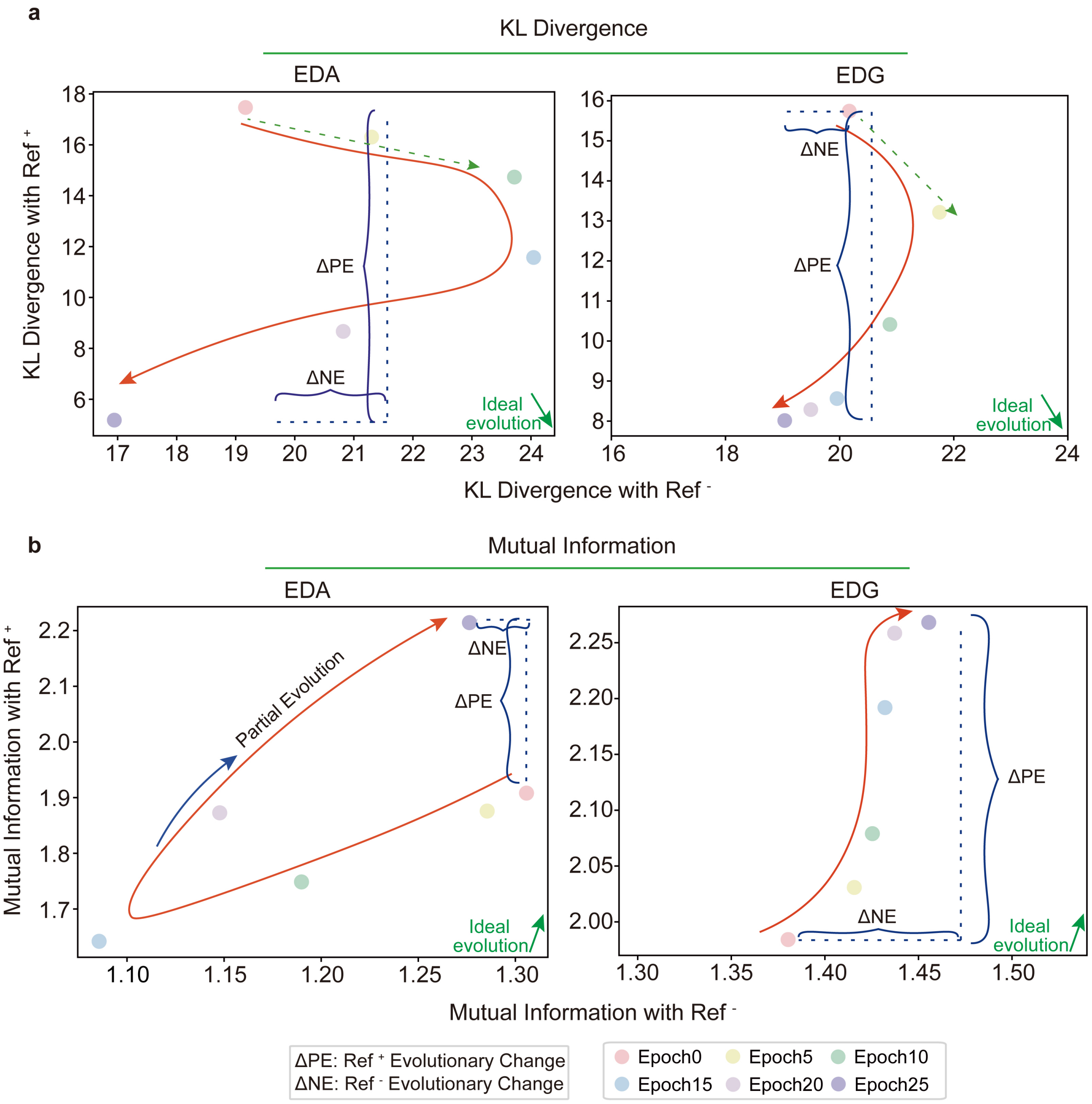

**Supplementary Fig. 9 | The evolutionary trajectories with a single reference are consistent with those with multiple references in the FE module.**

**a,** This panel shows the evolutionary trajectory of sequences designed by the EDA and EDG-based FE modules, measured by KL divergence. The analysis is conducted with a single sequence in the dataset of $\boldsymbol{Ref^{+}}$ and $\boldsymbol{Ref^{-}}$ as the reference point. The colors of the dots represent the outcomes after different epochs, as indicated by the color bar provided at the bottom.

**b,** This panel shows the evolutionary trajectory using mutual information as the measurement method, with a single sequence in the dataset of $\boldsymbol{Ref^{+}}$ and $\boldsymbol{Ref^{-}}$ as the reference point. The colors of the dots represent the outcomes after different epochs, as indicated by the color bar provided at the bottom.

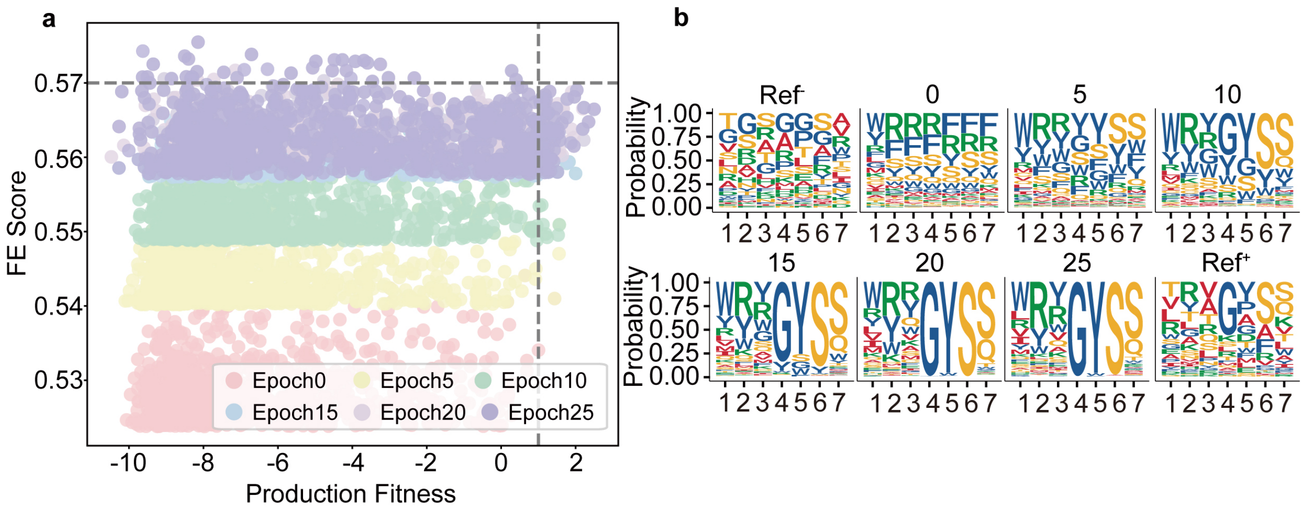

**Supplementary Fig. 10 | The evolutionary process of the functional and structural semantics of the sequence designed by the FE modules.**

**a,** The distribution of the FE score and production fitness of the sequences in different epochs of evolution in the FE module. The colors of the dots represent the outcomes after different epochs, as indicated by the color bar provided at the bottom.

**b,** The logo diagram shows the amino acid frequencies of the sequences in different epochs of evolution in the FE module and $\boldsymbol{Ref^{+}}$ , $\boldsymbol{Ref^{-}}$.

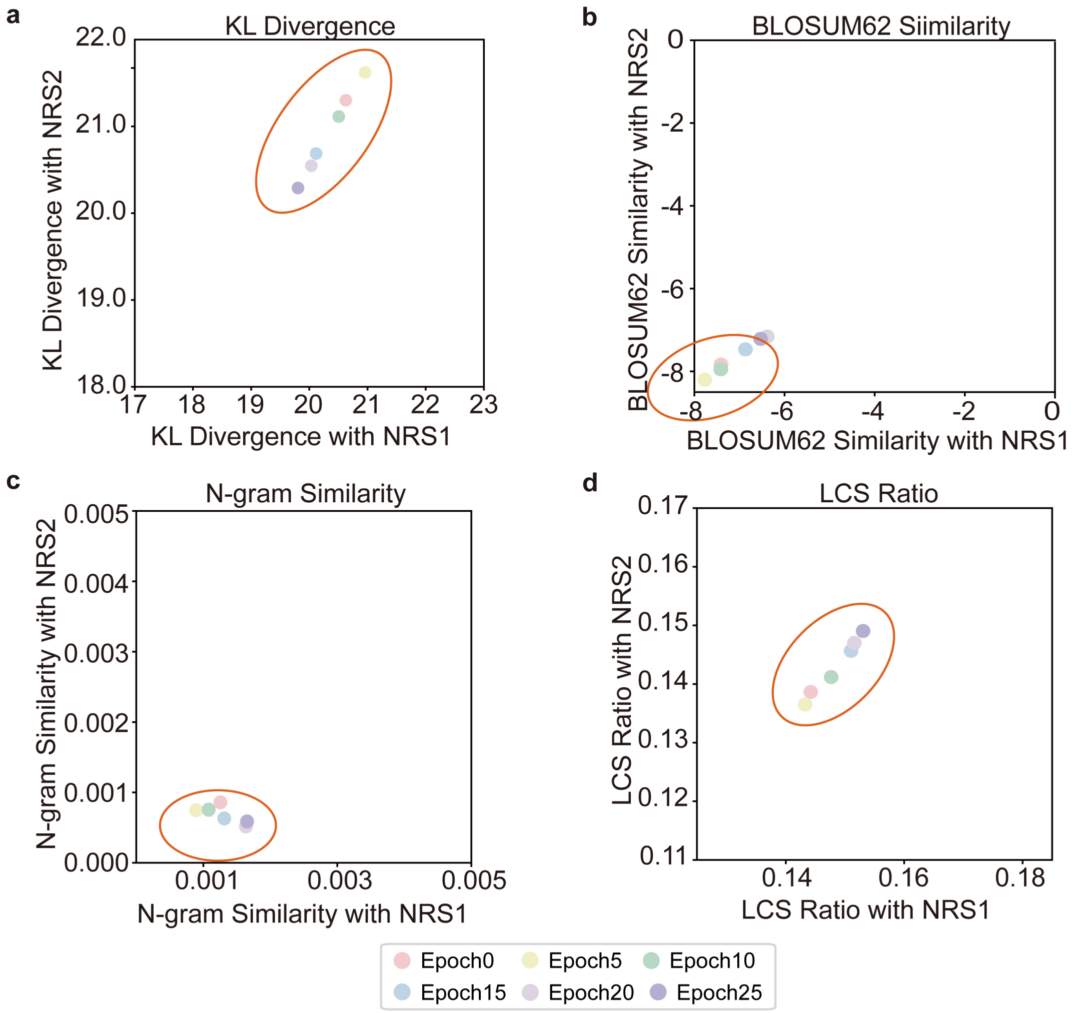

**Supplementary Fig. 11 | Robustness and controllability assessment of the FE module.**

The capsid sequences designed by the EDG-based FE module were contrastively assessed by two datasets of capsid sequences that were not utilized during the guided evolution process. These sequences are also not related to the sequences (NRS) that appear in the datasets of $\boldsymbol{Ref^{+}}$ or $\boldsymbol{Ref^{-}}$.

**a-d,** Plots showing the perturbations of the sequences in terms of evolution progress (a), functional similarity (b), local semantic information (c), and global semantic information (d). NRS1 and NRS2 indicate two datasets of sequences that are not related to the sequences that appeared in the dataset of $\boldsymbol{Ref^{+}}$ or $\boldsymbol{Ref^{-}}$.

The colors of the dots represent the outcomes after different epochs, as indicated by the color bar provided at the bottom.

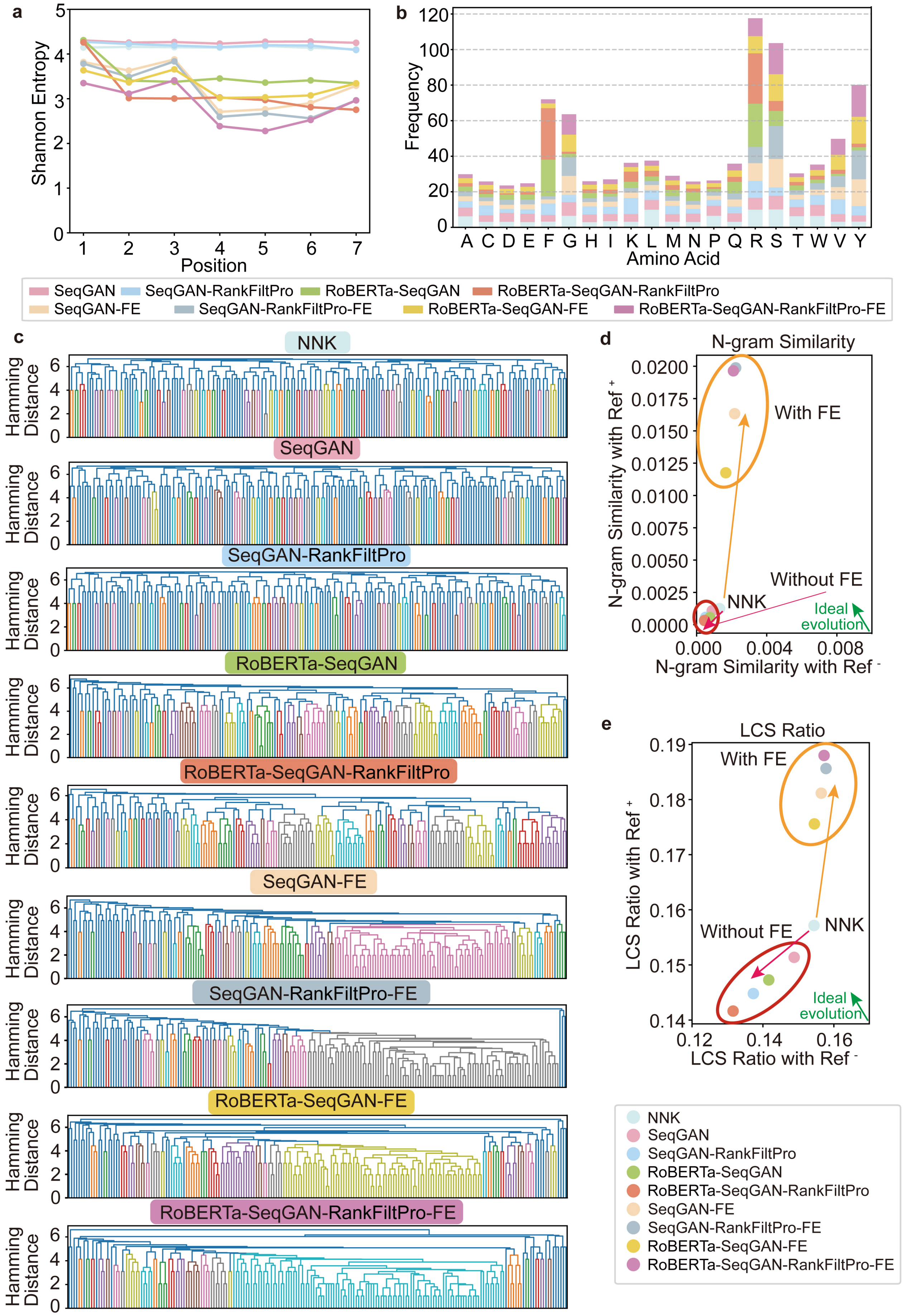

a
Shannon Entropy
Position
b
Frequency
Amino Acid
A C D E F G H I K L M N P Q R S T W V Y
SeqGAN
SeqGAN-RankFiltPro
RoBERTa-SeqGAN
RoBERTa-SeqGAN-RankFiltPro
SeqGAN-FE
SeqGAN-RankFiltPro-FE
RoBERTa-SeqGAN-FE
RoBERTa-SeqGAN-RankFiltPro-FE
c
Hamming Distance
NNK
SeqGAN
SeqGAN-RankFiltPro
RoBERTa-SeqGAN
RoBERTa-SeqGAN-RankFiltPro
SeqGAN-FE
SeqGAN-RankFiltPro-FE
RoBERTa-SeqGAN-FE
RoBERTa-SeqGAN-RankFiltPro-FE
d
N-gram Similarity
N-gram Similarity with Ref +
N-gram Similarity with Ref -
With FE
Without FE
NNK
Ideal evolution
e
LCS Ratio
LCS Ratio with Ref +
LCS Ratio with Ref -
With FE
Without FE
NNK
Ideal evolution
NNK
SeqGAN
SeqGAN-RankFiltPro
RoBERTa-SeqGAN
RoBERTa-SeqGAN-RankFiltPro
SeqGAN-FE
SeqGAN-RankFiltPro-FE
RoBERTa-SeqGAN-FE
RoBERTa-SeqGAN-RankFiltPro-FE

**Supplementary Fig. 12 | Semantic analysis of the sequences designed by the architectures with the ablation of distinct modules.**

**a,** The plot shows the sequence diversity distribution of capsid sequences generated by nine combined architectures in the ablation study at seven sequence positions measured via Shannon entropy.

**b,** The diagram shows the amino acid frequency distribution of capsid sequences designed by nine combined architectures in the ablation study.

**c,** Hamming distance display of capsid sequences designed by nine combined architectures in the ablation study.

**d-e,** Using the capsid sequences in the dataset of NNK as the control group, the diagram shows the evolutionary trajectory distribution of the capsid sequences designed by nine combined architectures in the ablation study from the aspects of local (d) and global (e) semantic information. The colors of the dots represent the outcomes from architectures with distinct ablations, as indicated by the legend provided at the bottom.

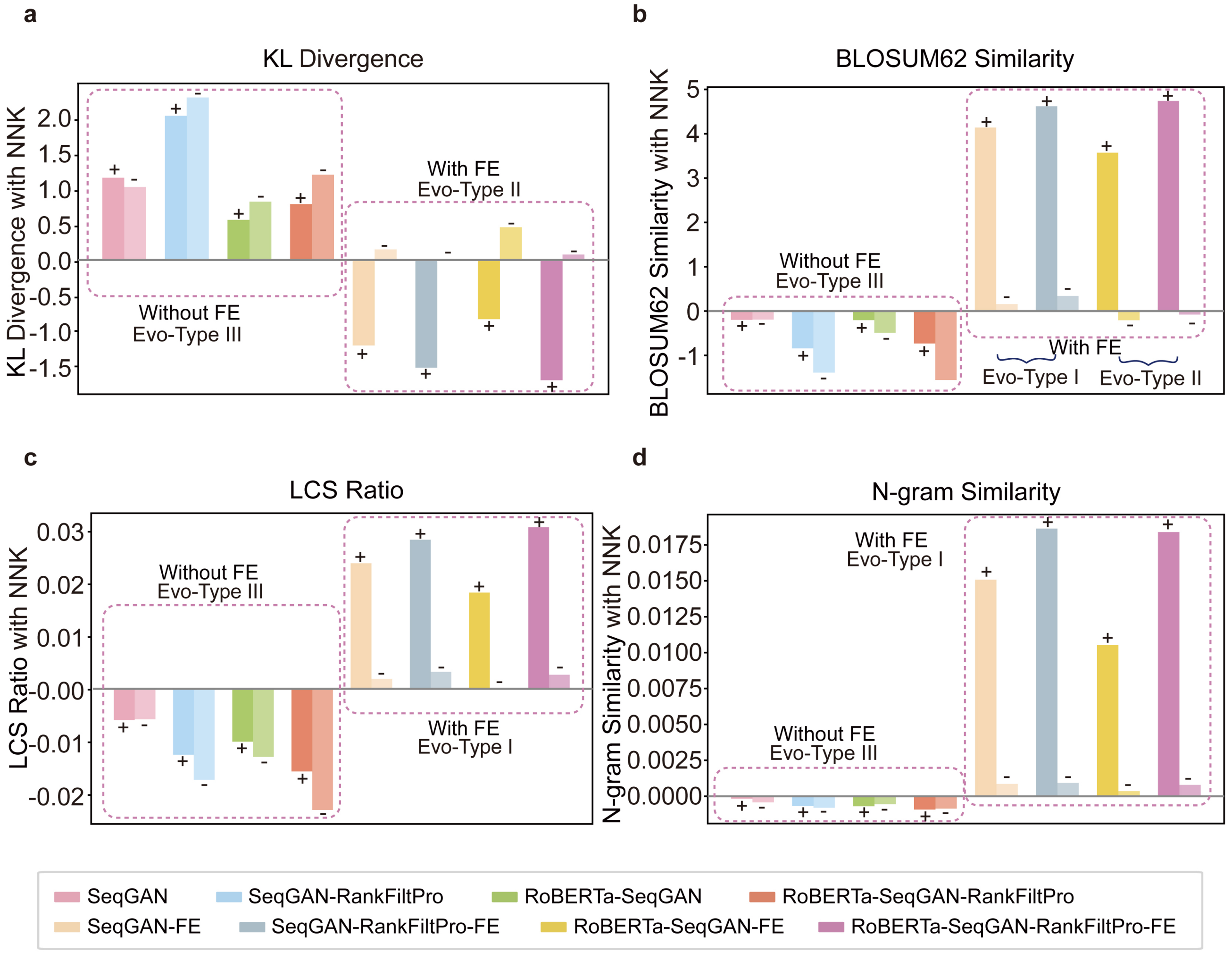

**Supplementary Fig. 13 | Quantification of the evolutionary changes in sequences designed by the architectures with the ablation of distinct modules.**

**a-b,** Histograms showing the evolutionary and functional changes toward $\boldsymbol{Ref}^{+}$ and $\boldsymbol{Ref}^{-}$ measured by KL divergence (a) and BLOSUM62 similarity (b), respectively. The colors of the histograms represent the outcomes from architectures with distinct ablations, as indicated by the color bar provided at the bottom. **c-d,** Histograms illustrating the evolutionary changes in the global and local semantic information of the sequences measured by the LCS ratio (c) and N-gram similarity (d), respectively. The sequences in the NNK dataset serve as controls. + and - indicate changes compared with $\boldsymbol{Ref}^{+}$ and $\boldsymbol{Ref}^{-}$, respectively. The colors of the histograms represent the outcomes from architectures with distinct ablations, as indicated by the color bar provided at the bottom.

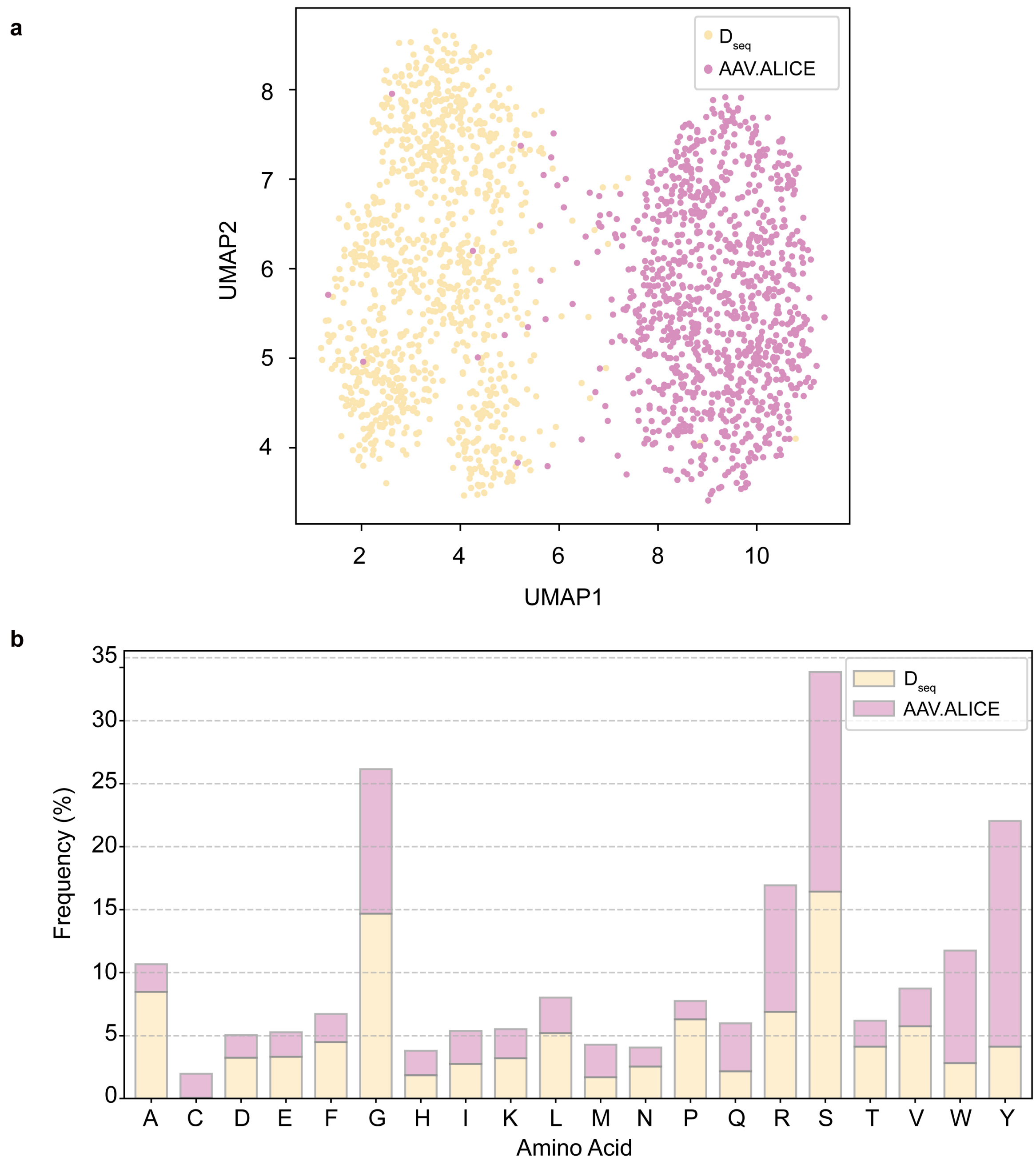

**Supplementary Fig. 14 | Comparative analysis of the capsid sequences designed by ALICE and those in the semantic tuning training dataset.**

**a,** Sequence feature analysis of capsid sequences designed by ALICE (AAV.ALCIE) and those in the semantic tuning training dataset ( $\boldsymbol{D_{seq}}$). In each category, a random sample of 1000 sequences was extracted for analysis**.**

**b,** Amino acid frequency distribution of capsid sequences designed by ALICE (AAV.ALCIE) and those in the semantic tuning training dataset ( $\boldsymbol{D_{seq}}$).

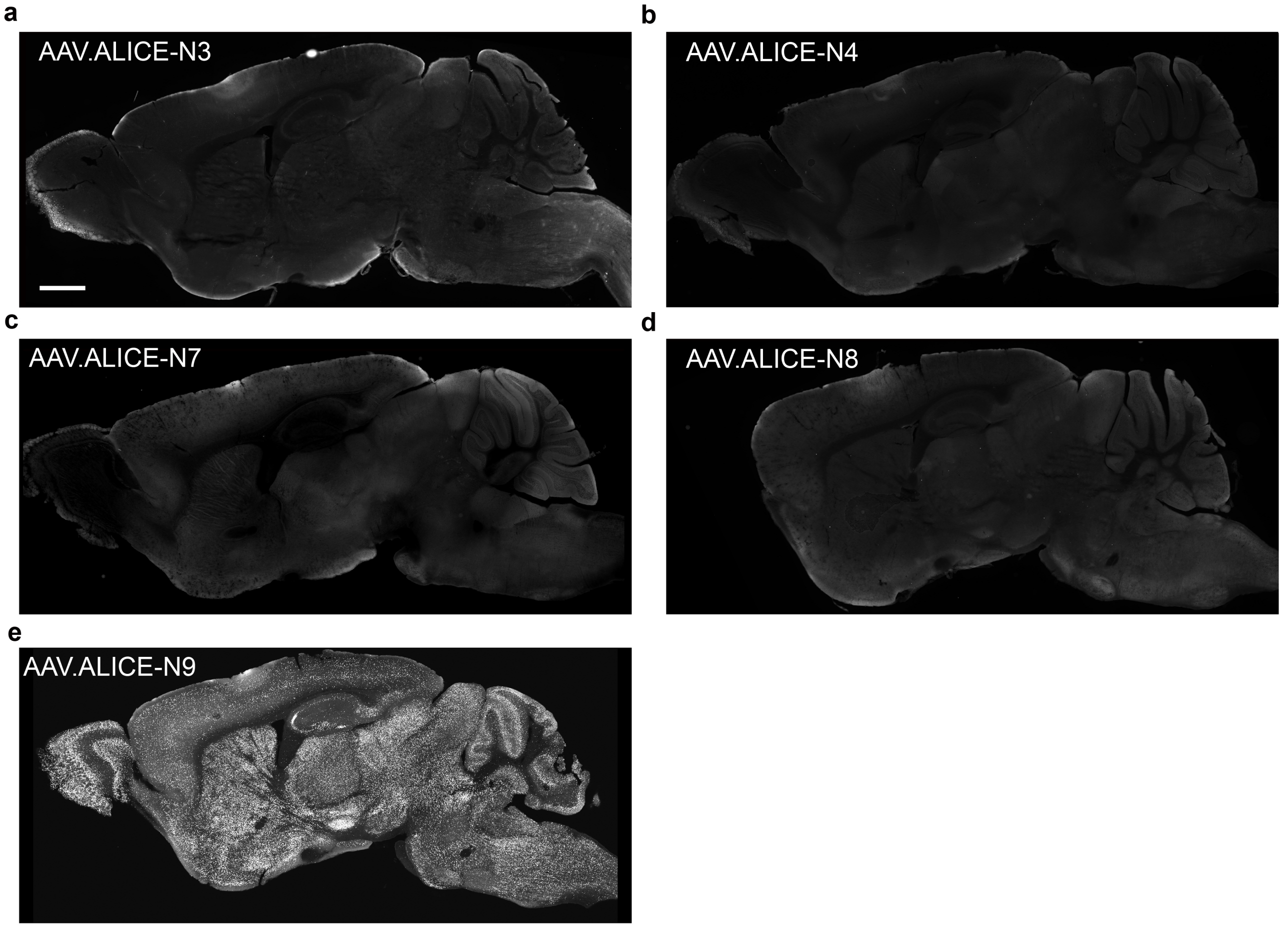

**Supplementary Fig. 15 | The engineered AAV variants did not efficiently infect CNS cells.**
**a-e,** These mice were administered intravenous injections of $5 \times 10^{11}$ vg of AAV.ALICE-N3 (a), AAV.ALICE-N4 (b), AAV.ALICE-N7 (c), AAV.ALICE-N8 (d) and AAV.ALICE-N9 (e). Scale bar,1 mm. After a 21-day recovery period post injection, the brains of mice were harvested. Representative images of excised brains showed only minimal detectable fluorescence.

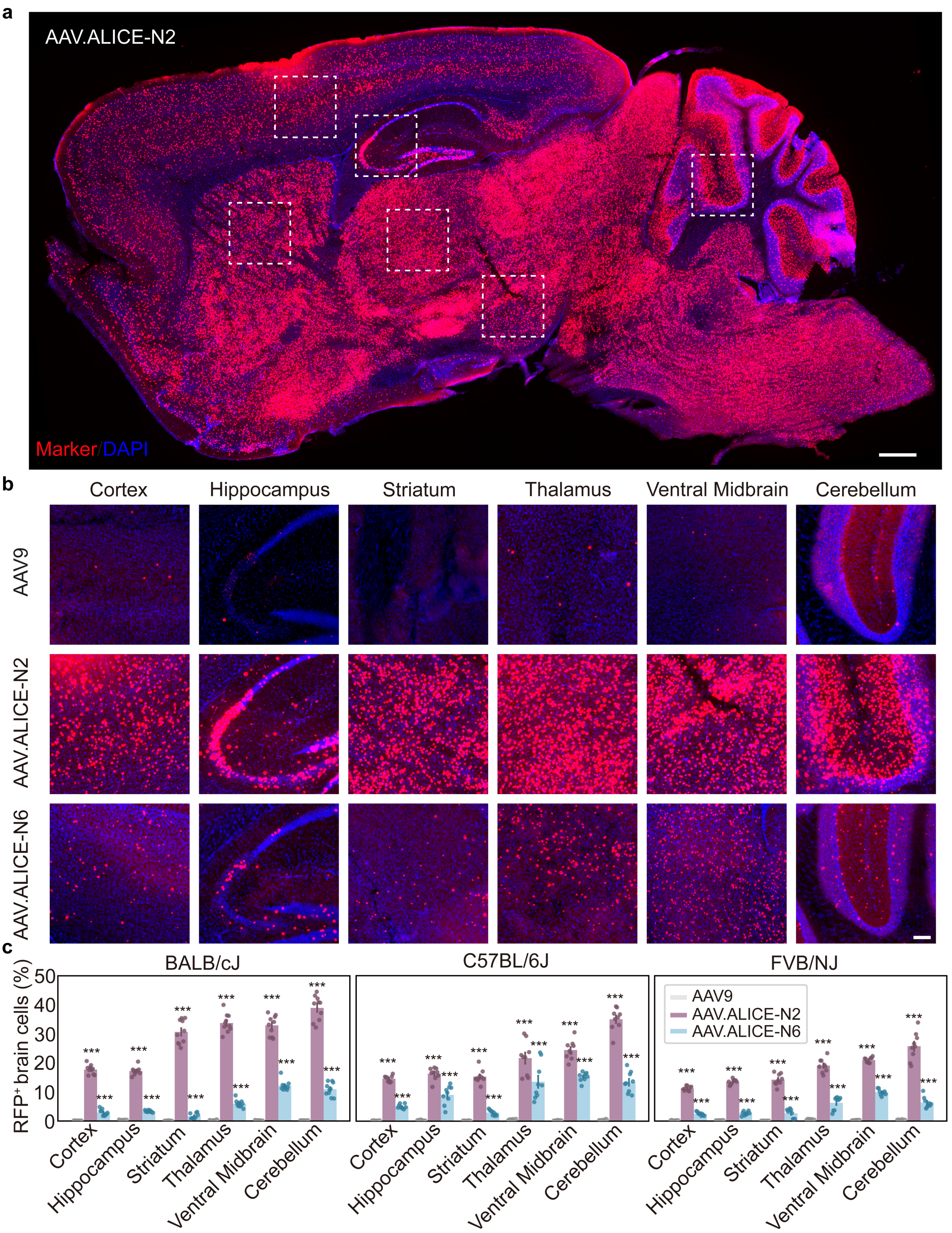

**Supplementary Fig.16 | Broad brain biodistribution of AAV9, AAV.ALICE-N2 and AAV.ALICE-N6 across multiple mouse strains.**

**a,** Representative fluorescence images are presented, depicting brain sections from BALB/cJ mice 21 days following intravenous injection with AAV.ALICE-N2 at a dosage of 5 × $10^{11}$ vg per animal. Scale bar,500 μm.

**b,** Detailed images of various brain regions within BALB/cJ mice, corresponding to the framed areas in (a), are provided to illustrate the distribution of the reporter gene. Scale bar,100 μm.

**c,** Depicted are quantitative analysis of the proportion of RFP (red fluorescent protein) positive cells in different brain regions of mice after injection of AAV9, AAV.ALICE-N2 and AAV.ALICE-N6 ($5 \times 10^{11}$ vg per animal) in BALB/cJ, C57BL/6J, and FVB/NJ mice. Data are presented as the mean ± s.e.m., n = 3-4 mice per group. Statistical significance was determined using one-way ANOVA with Tukey's posttest (*$P < 0.05$, **$P < 0.01$，***$P < 0.001$ versus AAV9).

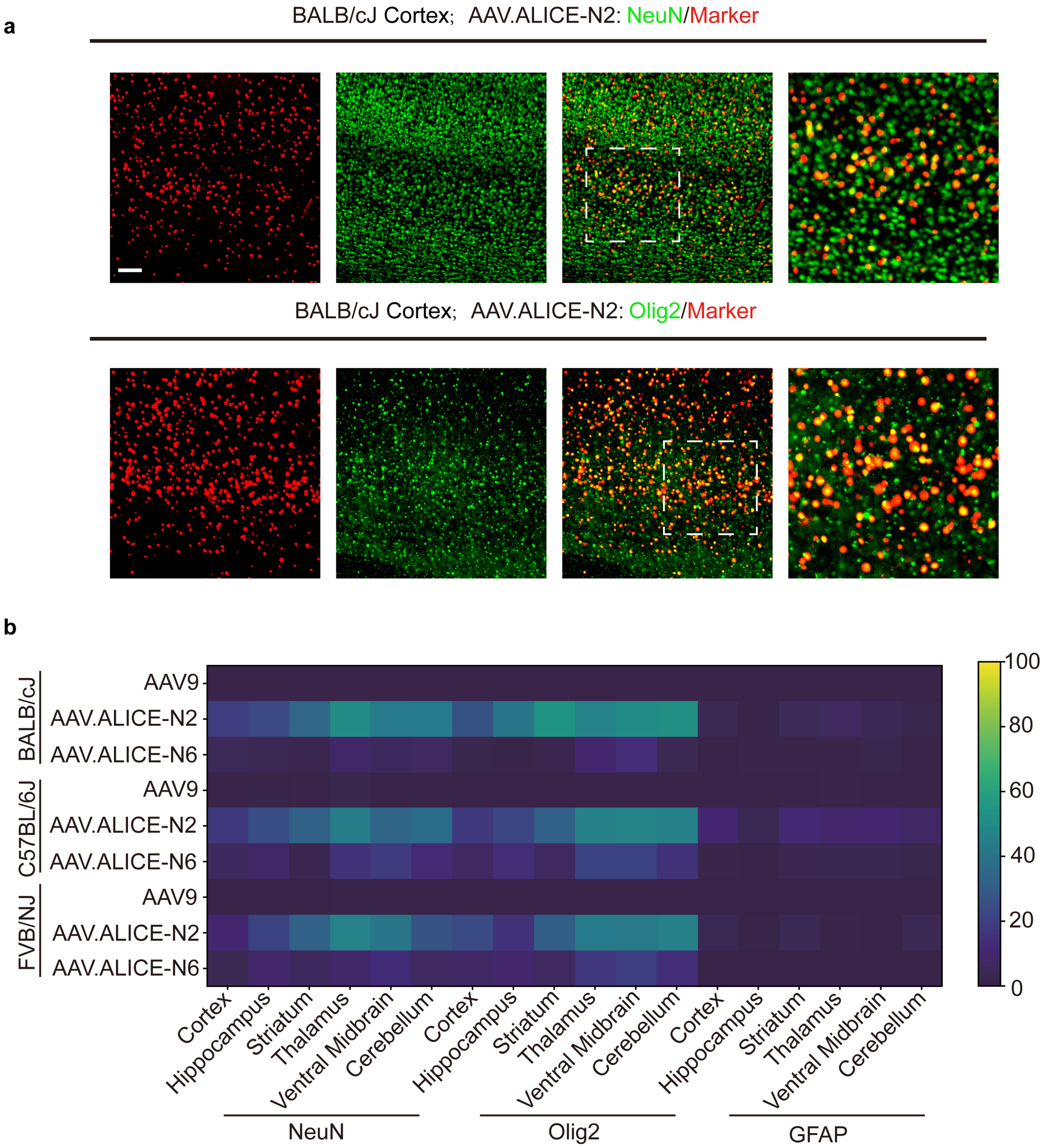

**Supplementary Fig. 17 | Types of CNS cells transduced by AAV9, AAV.ALICE-N2 and AAV.ALICE-N6.**

**a,** Representative images showing transduced neurons ($NeuN^+$) and oligodendrocytes ($Olig2^+$) in the brain. Panels display representative cortical regions with NeuN and Olig2 nuclear colabeling. Scale bars, 100 µm.
**b,** Heatmap of the enrichment of AAV9, AAV.ALICE-N2 and AAV.ALICE-N6 for NeuN, Olig2, GFAP cell colabeling in six brain regions of BALB/cJ, C57BL/6J and FVB/NJ mice (n = 3 mice per group, mean is plotted).

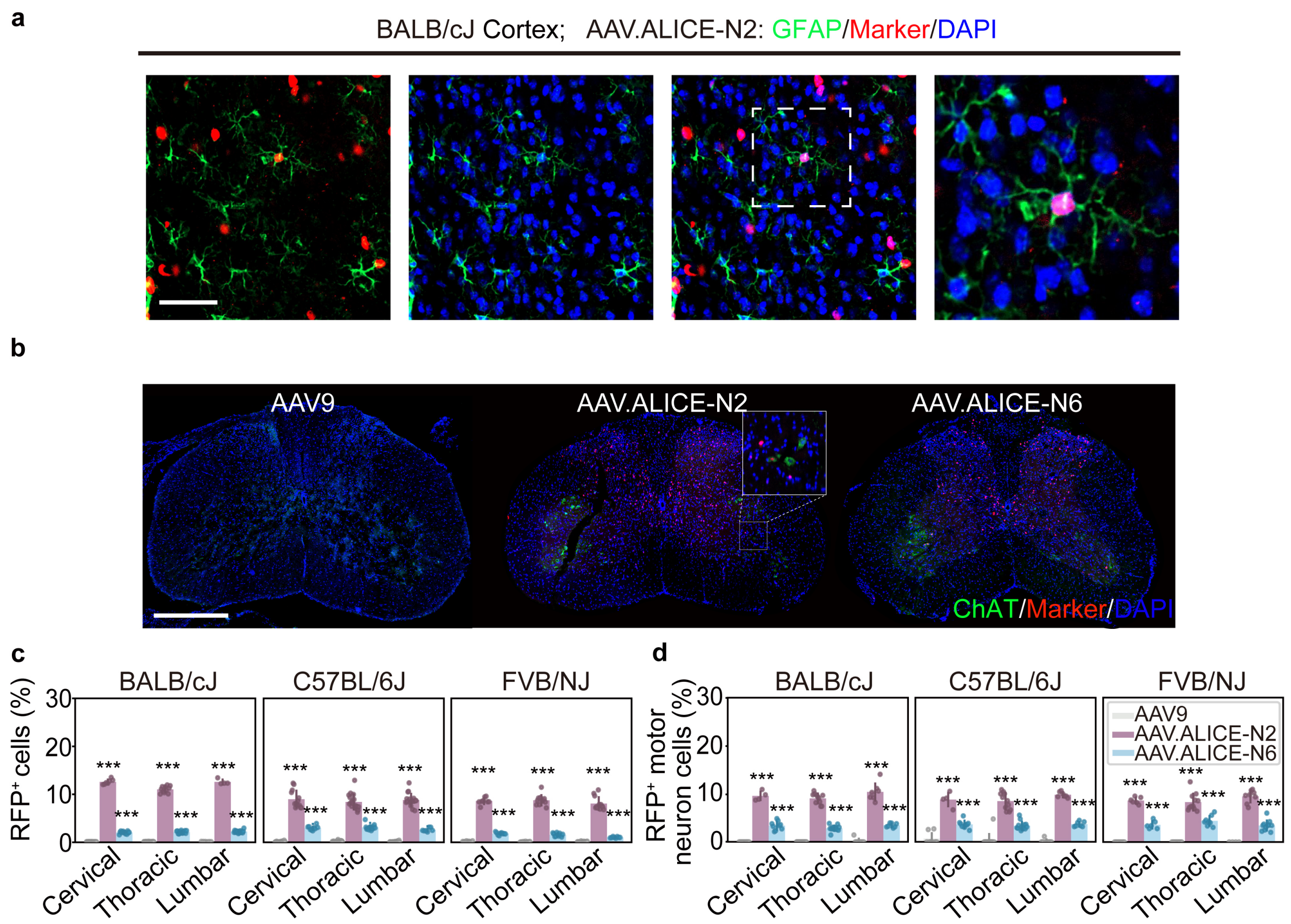

**Supplementary Fig. 18 | Transduction of the cells in the CNS with AAV9, AAV.ALICE-N2, and AAV.ALICE-N6 in different mouse strains.**

**a,** The representative pictures show the transduced astrocytes ($GFAP^+$) in the brain. Panels display representative cortical regions with GFAP and DAPI nuclear colabeling. Scale bars, 100 μm.

**b,** The representative pictures show transduced motor neurons ($ChAT^+$) in the spinal cord. Scale bars, 500 μm. Panels display representative spinal cord with ChAT and DAPI nuclear colabeling.

**c,** Quantitative analysis of neuronal labeling by AAV9, AAV.ALICE-N2, and AAV.ALICE-N6 across different segments of the spinal cord (cervical, thoracic, lumbar) is presented for various mouse strains.

**d,** A quantitative comparison of motor neuron labeling by AAV9, AAV.ALICE-N2, and AAV.ALICE-N6 in the cervical, thoracic, and lumbar segments of the spinal cord is shown across various mouse strains. Data are mean ± s.e.m. with n = 3-4 mice per group. Statistical significance was determined using one-way ANOVA with Tukey's posttest (*P < 0.05, **P < 0.01, ***P < 0.001 versus AAV9).

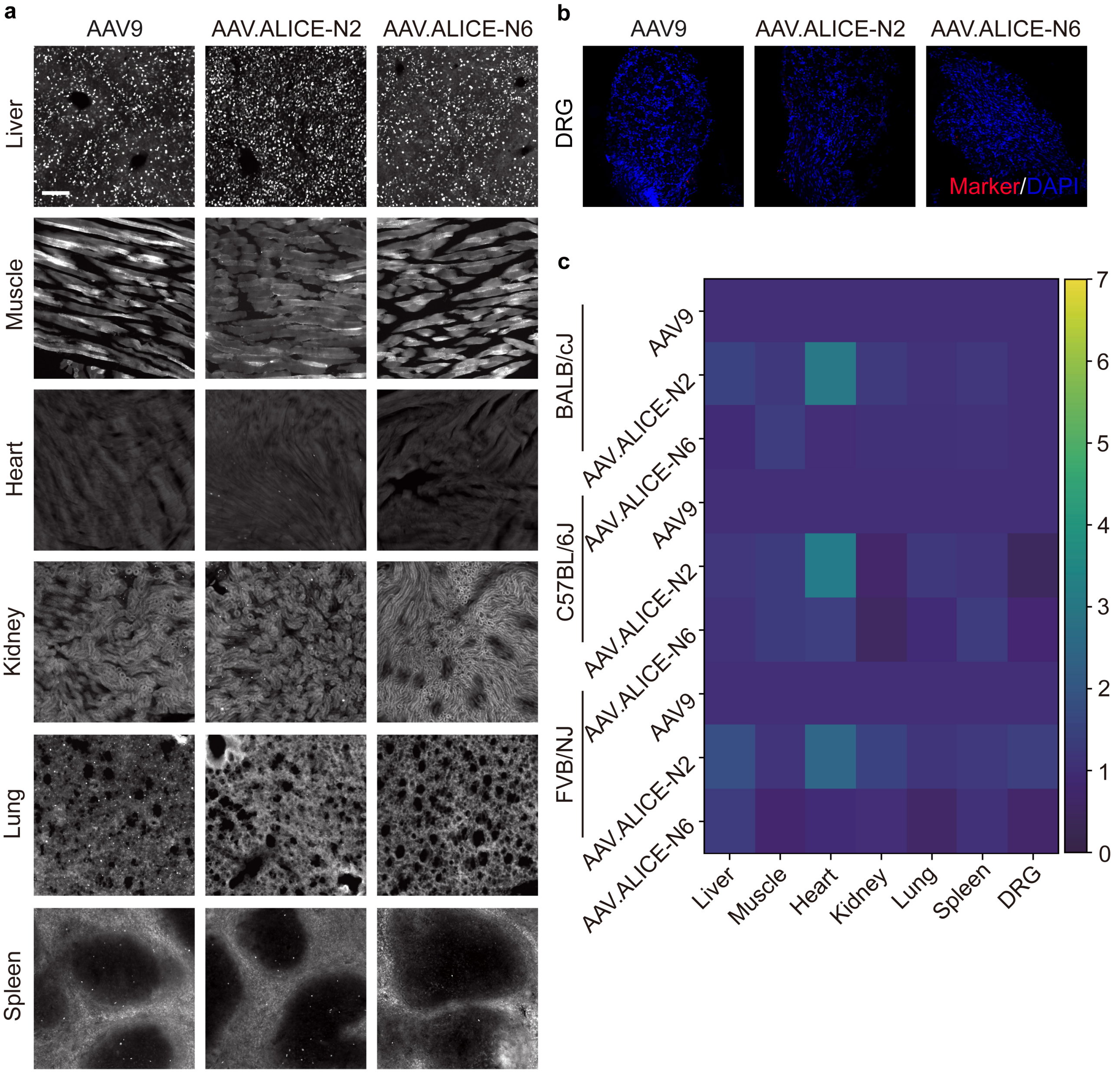

**Supplementary Fig. 19 | Transduction of peripheral tissues by AAV9, AAV.ALICE-N2, and AAV.ALICE-N6 in various mouse strains.**

**a,** Representative images of peripheral tissues (liver, muscle, heart, kidney, lung, and spleen) from mice transduced with AAV9, AAV.ALICE-N2, and AAV.ALICE-N6. Scale bars, 100 μm.

**b,** The distribution of AAV9, AAV.ALICE-N2, and AAV.ALICE-N6 in dorsal root ganglia (DRG) is shown. Scale bars, 100 μm.

**c,** A heatmap representation of the transduction efficiency of AAV9, AAV.ALICE-N2, and AAV.ALICE-N6 in the above peripheral tissues in different mouse strains, expressed as fold-change relative to AAV9 (n = 3 mice per group, mean is plotted). Enrichment ratios (dimensionless) compare ALICE-designed variants to AAV9, with values near 1 in the liver indicating comparable tropism.

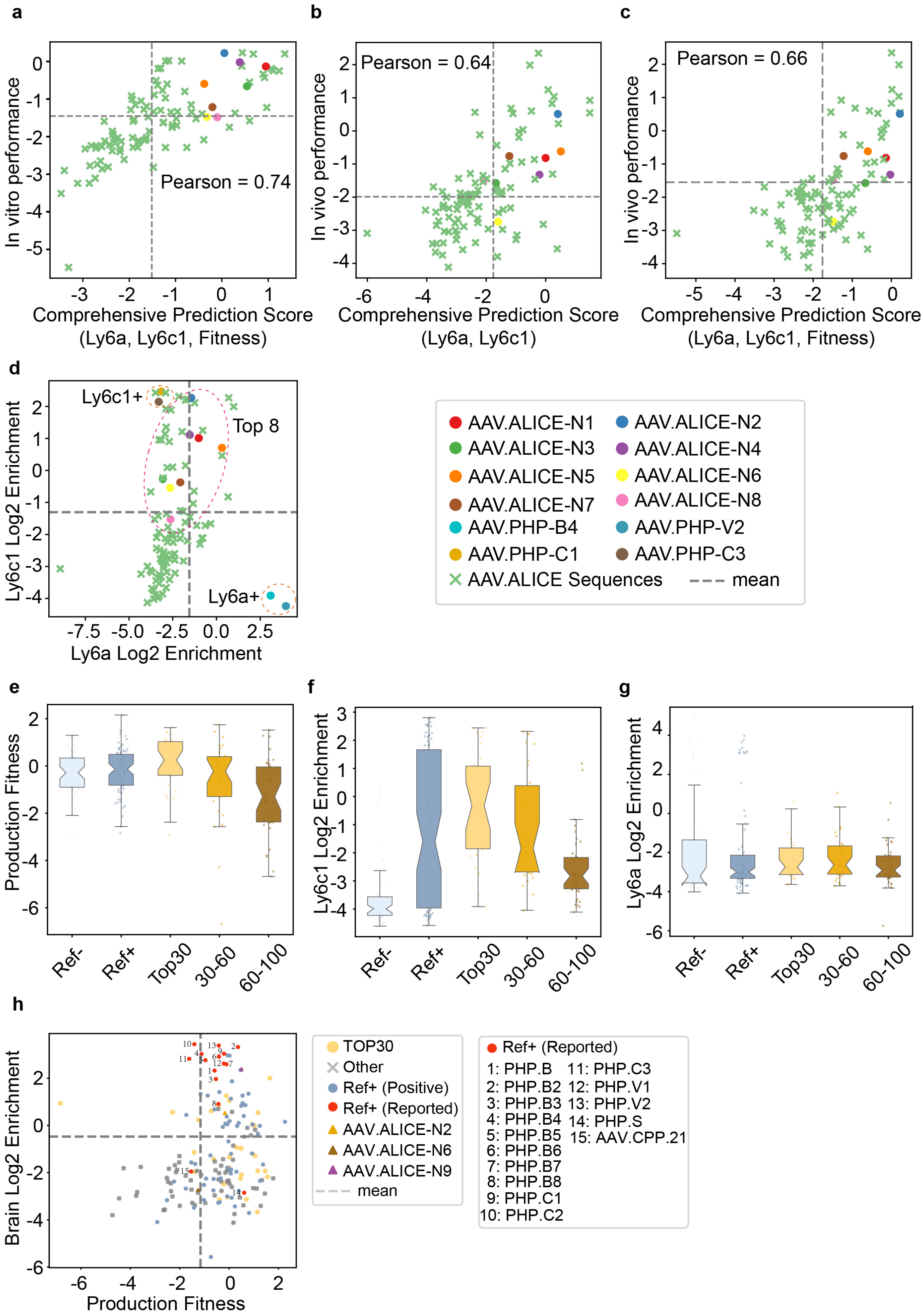

a
b
c
Pearson = 0.74
Pearson = 0.64
Pearson = 0.66
In vitro performance
In vivo performance
Comprehensive Prediction Score (Ly6a, Ly6c1, Fitness)
Comprehensive Prediction Score (Ly6a, Ly6c1)
d
Ly6c1+
Top 8
Ly6a+
Ly6c1 Log2 Enrichment
Ly6a Log2 Enrichment
AAV.ALICE-N1
AAV.ALICE-N2
AAV.ALICE-N3
AAV.ALICE-N4
AAV.ALICE-N5
AAV.ALICE-N6
AAV.ALICE-N7
AAV.ALICE-N8
AAV.PHP-B4
AAV.PHP-V2
AAV.PHP-C1
AAV.PHP-C3
AAV.ALICE Sequences
mean
e
f
g
Production Fitness
Ly6c1 Log2 Enrichment
Ly6a Log2 Enrichment
Ref-
Ref+
Top30
30-60
60-100
h
Brain Log2 Enrichment
Production Fitness
TOP30
Other
Ref+ (Positive)
Ref+ (Reported)
AAV.ALICE-N2
AAV.ALICE-N6
AAV.ALICE-N9
mean
Ref+ (Reported)
1: PHP.B
2: PHP.B2
3: PHP.B3
4: PHP.B4
5: PHP.B5
6: PHP.B6
7: PHP.B7
8: PHP.B8
9: PHP.C1
10: PHP.C2
11: PHP.C3
12: PHP.V1
13: PHP.V2
14: PHP.S
15: AAV.CPP.21

1767

**Supplementary Fig. 20 | Validation of the predictive power of ALICE.**

**a,** The correlation between the ALICE comprehensive prediction score and experimentally measured capsid performance is shown. Capsid performance was quantified as the mean of production fitness and Log2 enrichment of Ly6a and Ly6c1 from in vitro assays. The comprehensive prediction score was defined as the average of predicted production fitness, Ly6c1 log2 enrichment and Ly6a log2 enrichment values (detailed in Supplementary Methods).

**b,** The correlation between the ALICE predicted target binding capability (calculated by the average of Ly6c1 log2 enrichment and Ly6a log2 enrichment values) and in vivo performance is shown. In vivo performance is measured by brain Log2 enrichment.

**c,** The correlation between the comprehensive prediction score of ALICE and in vivo performance is shown. In vivo performance is measured by brain Log2 enrichment. The comprehensive prediction score was defined as the average of predicted production fitness, Ly6c1 log2 enrichment and Ly6a log2 enrichment values (detailed in Supplementary Methods).

**d,** The distribution of experimentally measured Ly6a and Ly6c1 Log2 enrichment for capsids generated by ALICE is shown. Ly6a-positive and Ly6c1-positive capsids are included as positive controls, and the top eight capsids were highlighted.

**e-g,** Box plots of the performance of capsids generated by ALICE, including in vitro measures: production fitness (e), Ly6c1 log2 enrichment (f), and Ly6a log2 enrichment (g). $\boldsymbol{Ref}^{+}$ and $\boldsymbol{Ref}^{-}$ are shown as controls.

**h,** Distribution of experimentally measured production fitness and brain log2 enrichment of capsids generated by ALICE, with $\boldsymbol{Ref}^{+}$ and $\boldsymbol{Ref}^{-}$ shown as controls. $\boldsymbol{Ref}^{+}$ (Positive) includes sequences selected for their high production fitness—reflecting superior replication efficiency and infectivity—as well as their strong binding affinity to Ly6a and Ly6c1, two key cellular receptors involved in viral entry and immune regulation. $\boldsymbol{Ref}^{+}$ (Reported) refers to sequences previously documented to exhibit robust blood–brain barrier (BBB) permeability, high viability, and consistent in vivo performance across multiple mouse strains, including C57BL/6 and BALB/c. Additional details are provided in Supplementary Table 2.

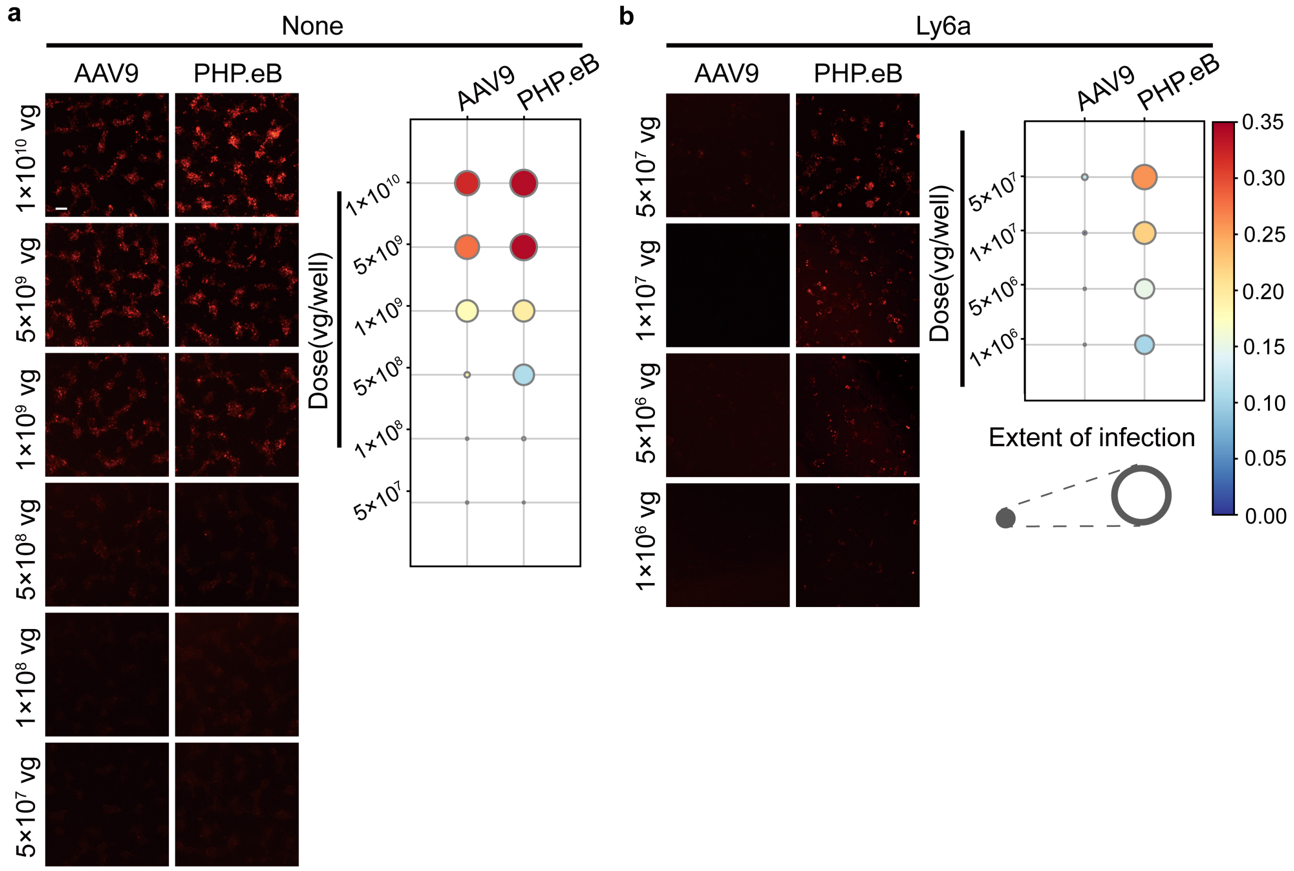

**Supplementary Fig. 21 | Gradient exploration of capsid titers.**

**a-b,** Fluorescence images showing the dose dependence of AAV9 and PHP.eB in HEK293T cells in 96-well plates. At 1 x $10^7$ v.g. per well, PHP.eB has markedly higher potency than AAV9 in Ly6a-transfected HEK293T cells. Representative images are shown. Scale bars = 50 μm.

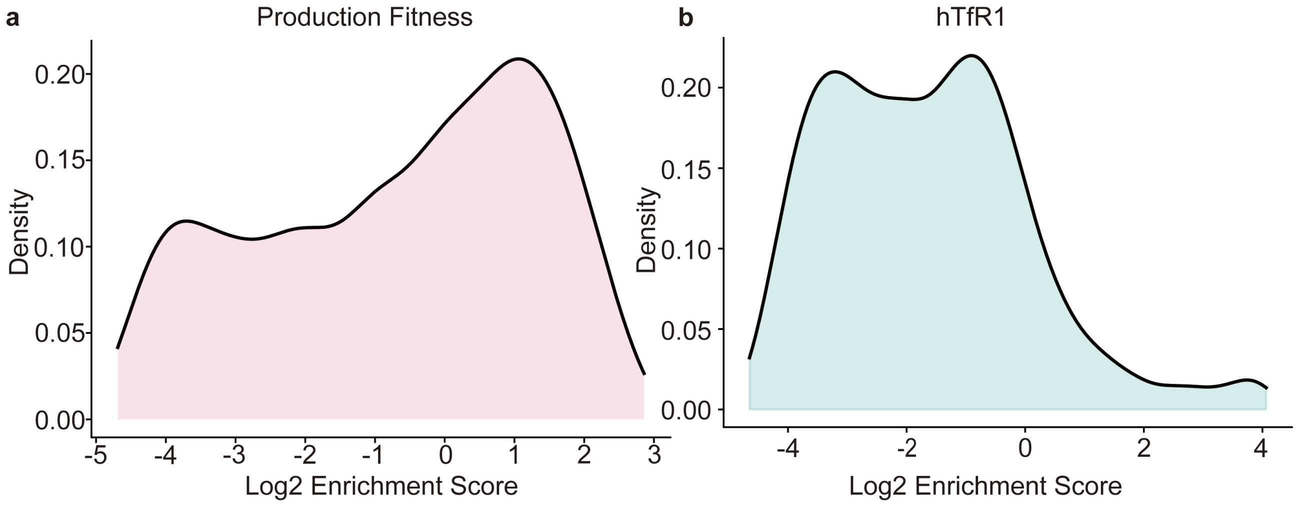

**Supplementary Fig. 22 | Distribution of training data for the evaluation model in ALICE-X.**

**a-b,** Distribution of training data ($\boldsymbol{D_{fitness-x}}$, $\boldsymbol{D_{hTfR1}}$). for the evaluation model of production fitness (a) and hTfR1 log2 enrichment score (b) (see Supplementary Methods for details).

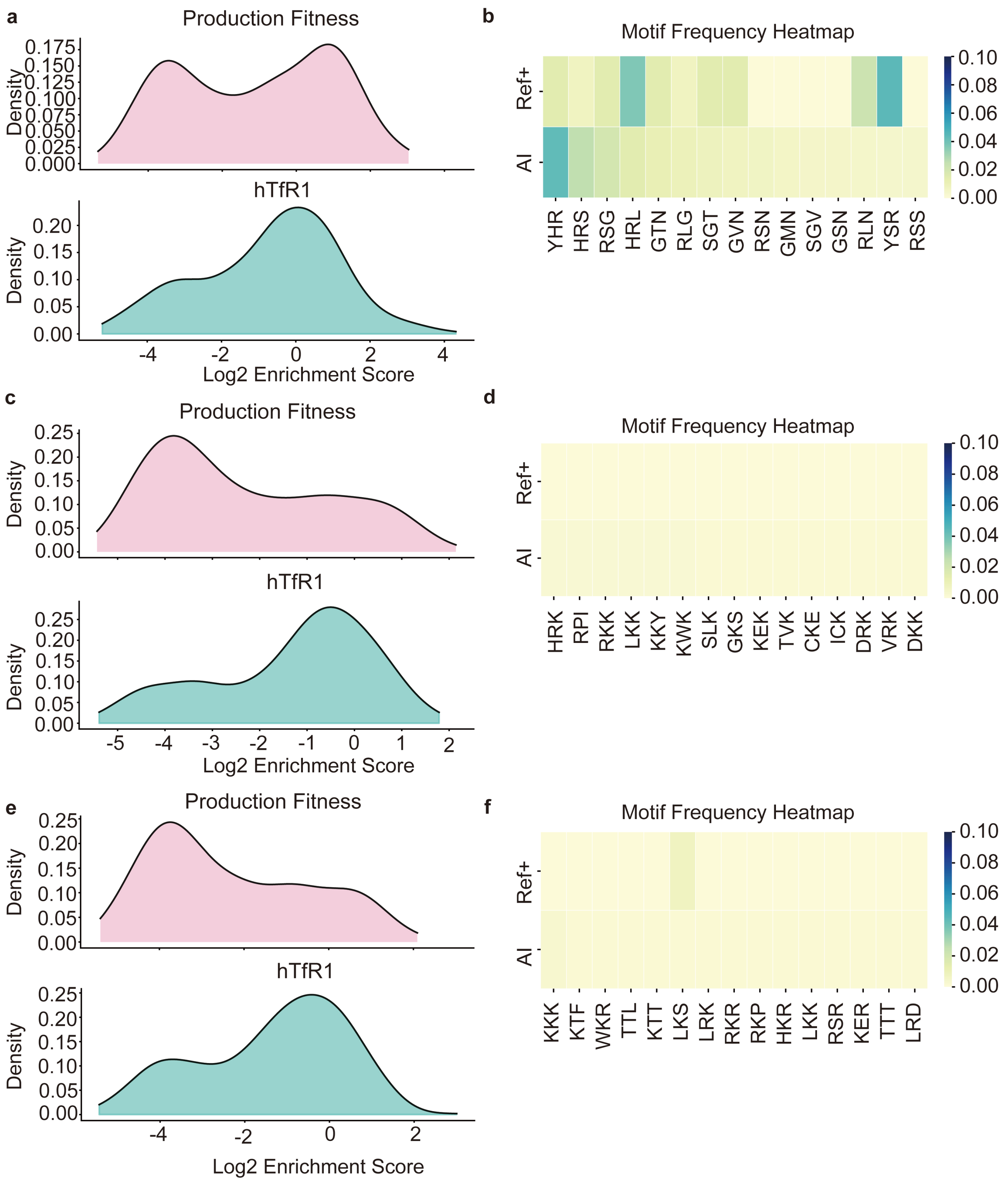

**Supplementary Fig. 23 | Trade-off between multi-function fusion and novelty exploration during sequence design.**

**a-b,** With a low novelty normalization weight (0.2), the generated sequences successfully fused both fitness and hTfR1 binding capability (a), but the explored motifs remained closely aligned with those in $\boldsymbol{Ref}^{+}$(b).

**c-d,** At a moderate weight (1.1), the explored motifs diverged from those in $\boldsymbol{Ref}^{+}$(d), but functional fusion was disrupted (c), resulting in reduced fitness.

**e-f,** A higher novelty weight (2.0) further disrupted functional integration, resulting in even lower fitness (e).

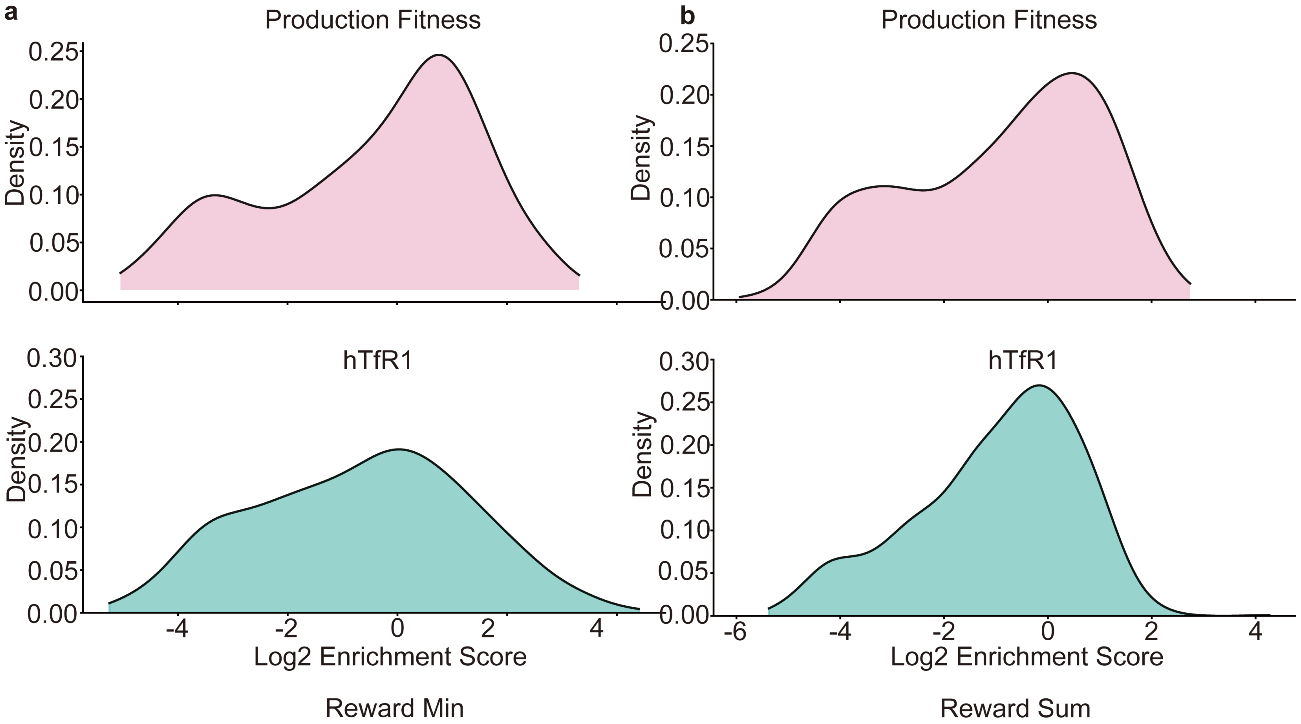

**Supplementary Fig. 24 | Performance comparison of exploration strategies using Min vs. Sum reward functions.**

**a,** Sequence distributions of production fitness and hTfR1 log2 enrichment after 20 evolutionary rounds using the Min reward function.

**b,** Sequence distributions of production fitness and hTfR1 log2 enrichment after 20 evolutionary rounds using the Sum reward function, showing improved hTfR1 log2 enrichment while preserving high fitness.

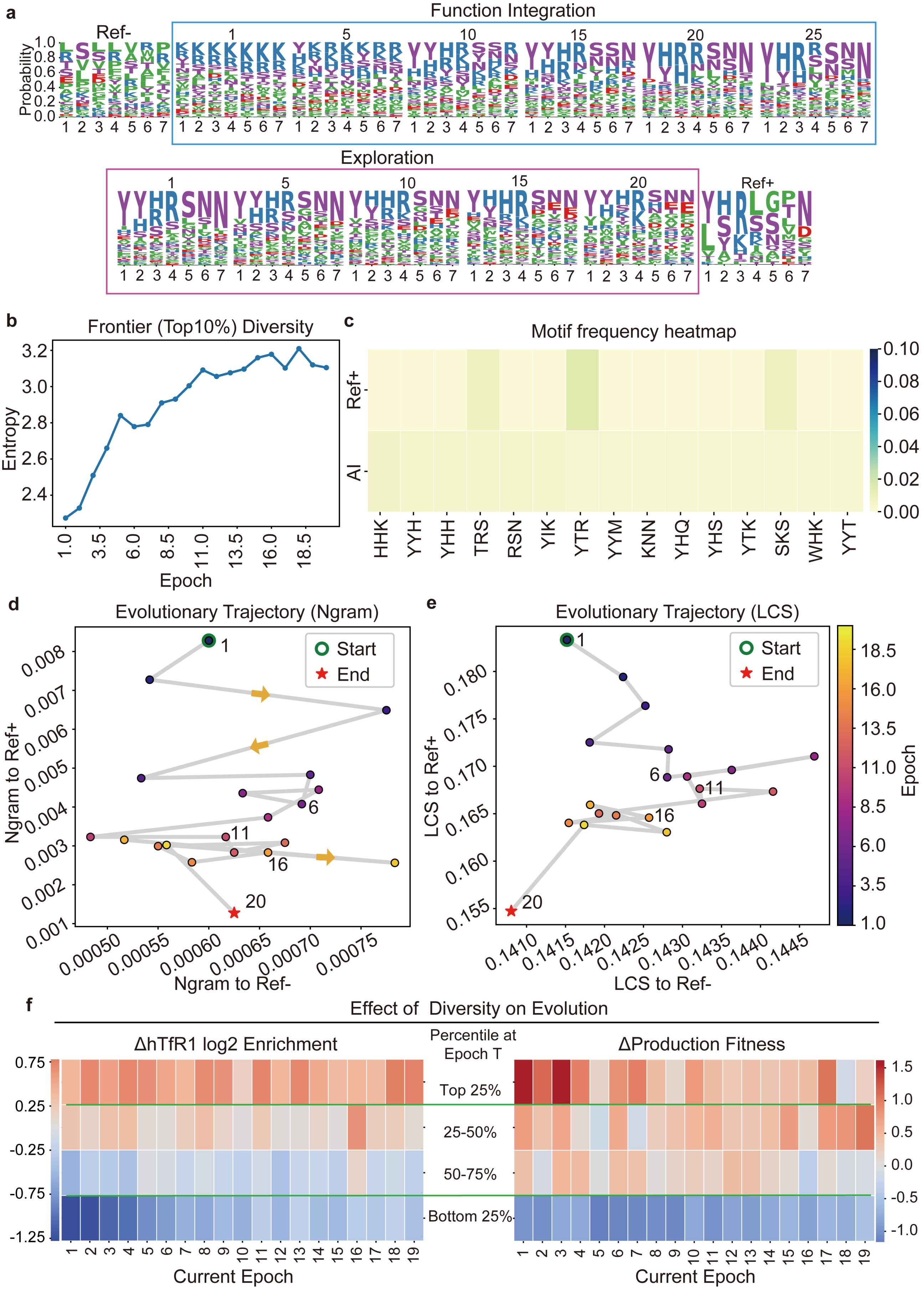
a
Ref-
Function Integration
Probability
Exploration
Ref+
b
Frontier (Top10%) Diversity
Entropy
Epoch
c
Motif frequency heatmap
Ref+
AI
HHK
YYH
YHH
TRS
RSN
YIK
YTR
YYM
KNN
YHQ
YHS
YTK
SKS
WHK
YYT
d
Evolutionary Trajectory (Ngram)
Start
End
Ngram to Ref+
Ngram to Ref-
e
Evolutionary Trajectory (LCS)
LCS to Ref+
LCS to Ref-
Epoch
f
Effect of Diversity on Evolution
ΔhTfR1 log2 Enrichment
Percentile at Epoch T
ΔProduction Fitness
Top 25%
25-50%
50-75%
Bottom 25%
Current Epoch
Current Epoch

**Supplementary Fig. 25 | Evolutionary trajectories of ALICE-X demonstrating novel sequence space exploration.**

**a,** The diagram illustrates the amino acid frequencies in ALICE-X's sequences during the Function Integration and Exploration stages, showing their progression across different evolutionary epochs of the FE-X module.

**b,** Increase in sequence diversity among the top 10% of variants generated during the exploration phase.

**c,** Motif distribution of ALICE-X–generated sequences compared with those in $\boldsymbol{Ref}^{+}$..

**d,** Local semantic evolutionary trajectories of ALICE-X during the exploration stage.

**e,** Global semantic evolutionary trajectories of ALICE-X during the exploration stage.

**f,** Effect of diversity on the median change in hTfR1 log2 enrichment and production fitness in the subsequent epoch during evolution.

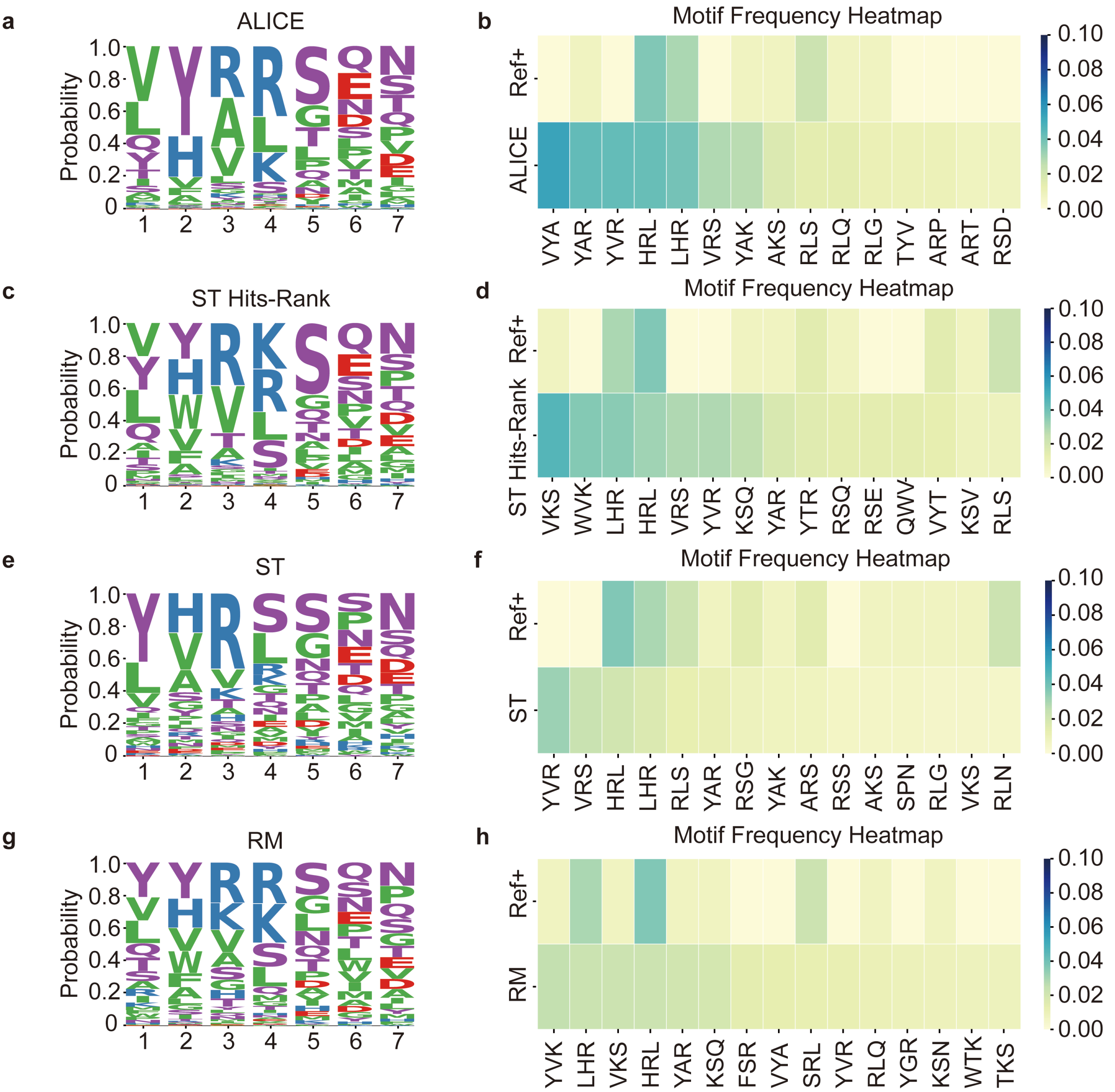

**Supplementary Fig. 26 | Comparative performance evaluation of ablation methods for spatial exploration and motif discovery.**

**a,** Amino acid frequency logo plot of the top 500 sequences generated by the ALICE model, showing the positional distribution of enriched residues.

**b,** Motif frequency heatmap of the top 15 motifs in ALICE-generated sequences compared with those in $\boldsymbol{Ref}^{+}$.

**c,** Amino acid frequency logo plot of the top 500 sequences generated by the Semantic Tuning + Hits Ranking (ST Hits-Rank) model.

**d,** Motif frequency heatmap of the top 15 motifs in ST Hits-Rank–generated sequences compared with those in $\boldsymbol{Ref}^{+}$.

**e,** Amino acid frequency logo plot of the top 500 sequences generated by the Semantic Tuning (ST)–only model.

**f,** Motif frequency heatmap of the top 15 motifs in ST-only–generated sequences compared with those in $\boldsymbol{Ref^{+}}$.

**g,** Amino acid frequency logo plot of the top 500 sequences generated by the Random Mutation (RM) model.

**h,** Motif frequency heatmap of the top 15 motifs in RM-generated sequences compared with those in $\boldsymbol{Ref^{+}}$.

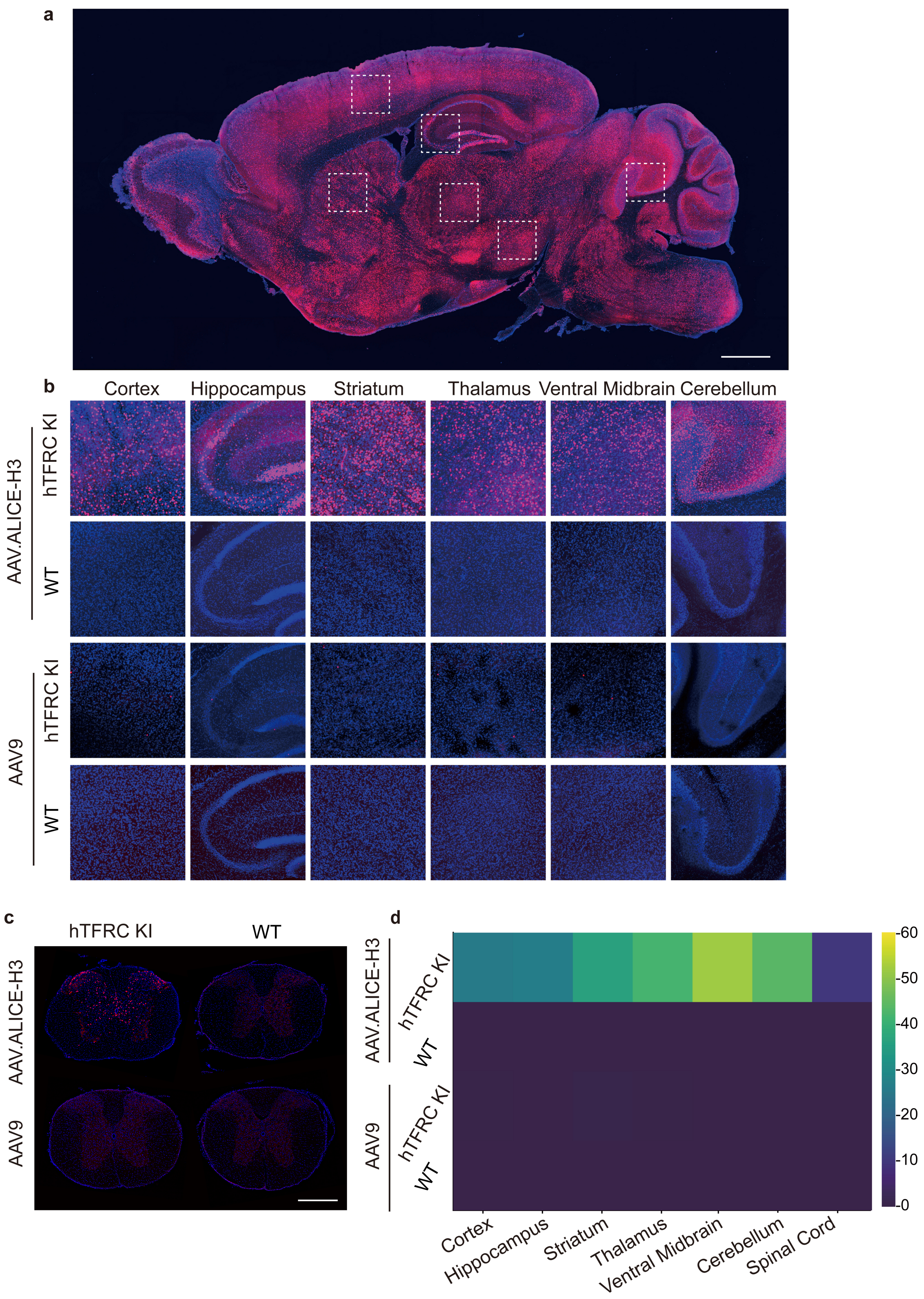
a
b
Cortex
Hippocampus
Striatum
Thalamus
Ventral Midbrain
Cerebellum
AAV.ALICE-H3
hTFRC KI
WT
AAV9
hTFRC KI
WT
c
hTFRC KI
WT
AAV.ALICE-H3
AAV9
d
AAV.ALICE-H3
hTFRC KI
WT
AAV9
hTFRC KI
WT
Cortex
Hippocampus
Striatum
Thalamus
Ventral Midbrain
Cerebellum
Spinal Cord
60
50
40
30
20
10
0

**Supplementary Fig. 27 | H2A-RFP expression profiles in CNS regions transduced by AAV9 versus AAV.ALICE-H3.**

**a,** Representative fluorescence images are presented, depicting brain sections from hTFR1 KI mice 21 days following intravenous injection with AAV.ALICE-H3 at a dosage of 5 × 10^11 vg per animal. Scale bar,1 mm.

**b,** Detailed images of various brain regions within hTFR1 KI mice and wild type mice (C57BL/6J), corresponding to the framed areas in (a), are provided to illustrate the distribution of the reporter gene.

**c,** The representative pictures show AAV9 and AAV.ALICE-H3 transduction in the spinal cord. Scale bars, 500 μm.

**d,** Heatmap representation of CNS cells transduction efficiency (H2A-RFP$^+$/DAPI$^+$ cells) transduced with AAV9 or AAV.ALICE-H3 across six brain regions and spinal cord of *hTFR1*-KI versus wild-type mice (n = 3 mice per group, mean is plotted).

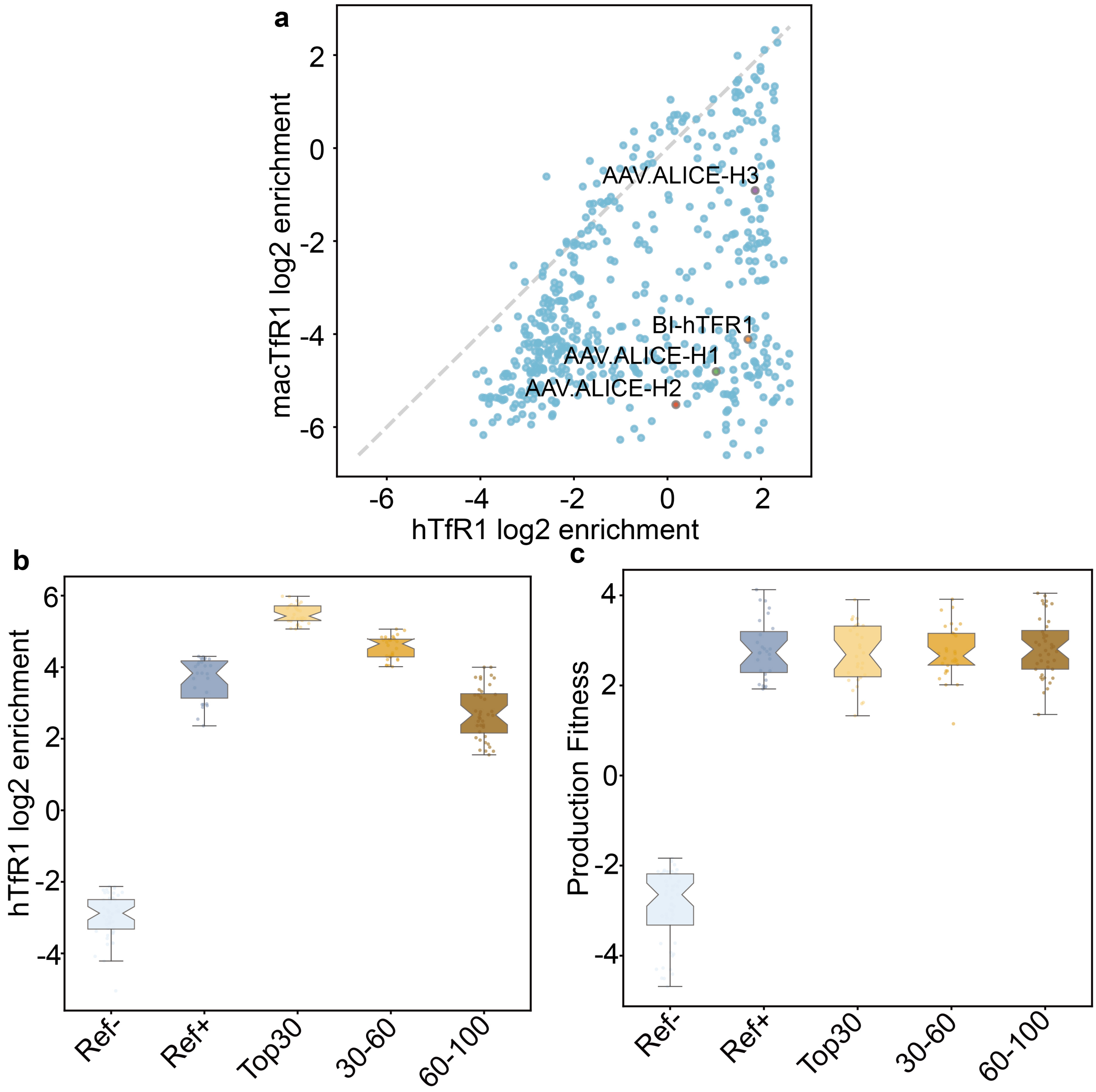

**Supplementary Fig. 28 | Features of Engineered AAV Variants and Comparison with Ref+.**

**a,** Correlation of variant enrichment (log2) between human and macaque TfR1 in a pulldown assay using the AAV-588-OligoPool-lib2 library.

**b,** The hTfR1 log2 enrichment of top sequences designed by ALICE-X, compared with $\boldsymbol{Ref}^{+}$ and $\boldsymbol{Ref}^{-}$.

**c,** The production fitness of top sequences designed by ALICE-X, compared with $\boldsymbol{Ref}^{+}$ and $\boldsymbol{Ref}^{-}$.

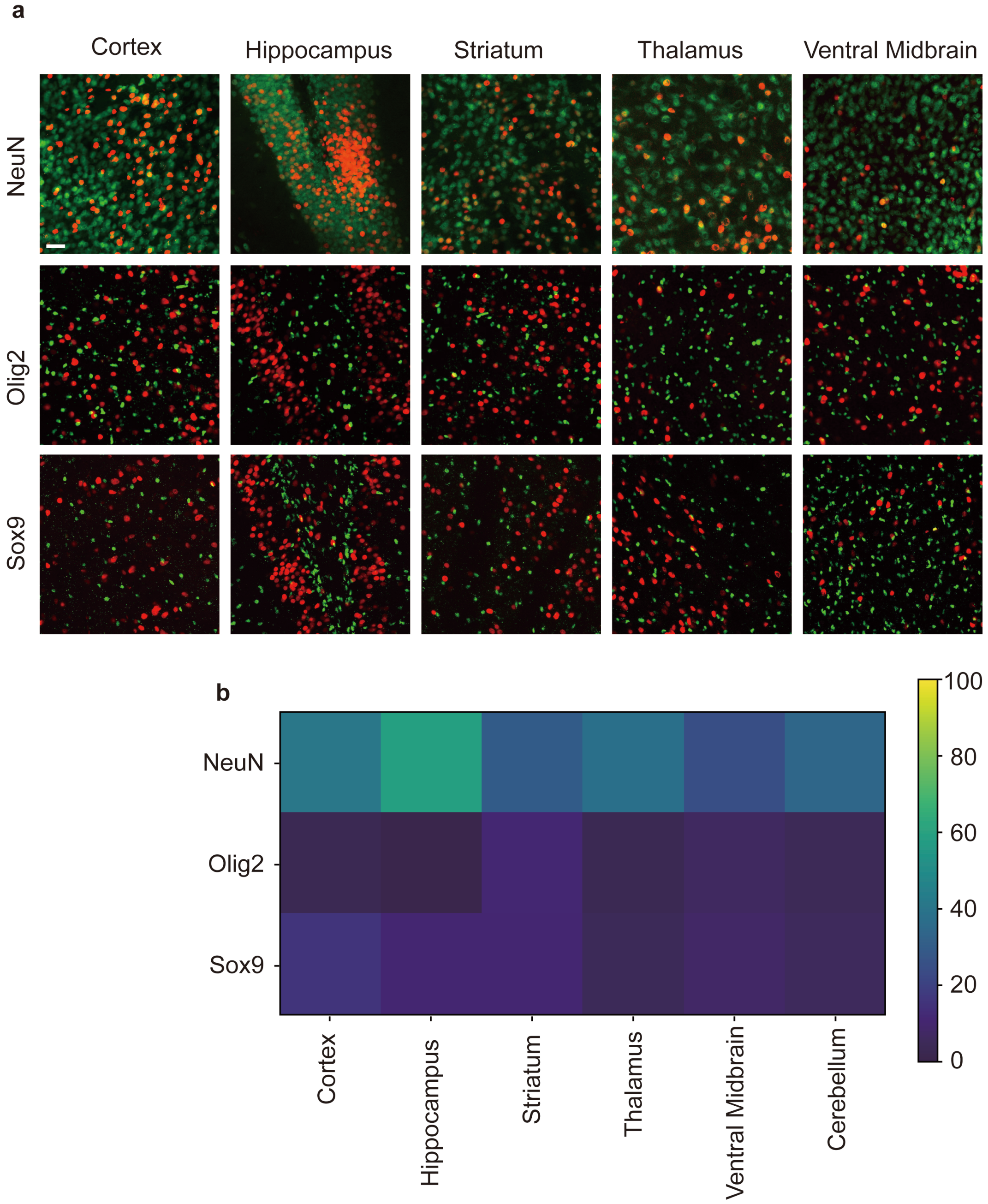

**Supplementary Fig. 29 | Cell-type-specific transduction efficiency of AAV.ALICE-H3 across brain regions.**

**a,** Representative images showing AAV.ALICE-H3-transduced cells (RFP+, red) and neurons (NeuN+, green), oligodendrocytes (Olig2+, green) and astrocytes (Sox9+, green) in the brain. Scale bars, 100 μm.

**b,** Heatmap representation of $NeuN^{+}$, $Olig2^{+}$, and $Sox9^{+}$ cell proportions among AAV.ALICE-H3-transduced (H2A-$RFP^{+}$) cells across six brain regions in hTFR1-KI mice (n = 3 mice per group, mean is plotted).

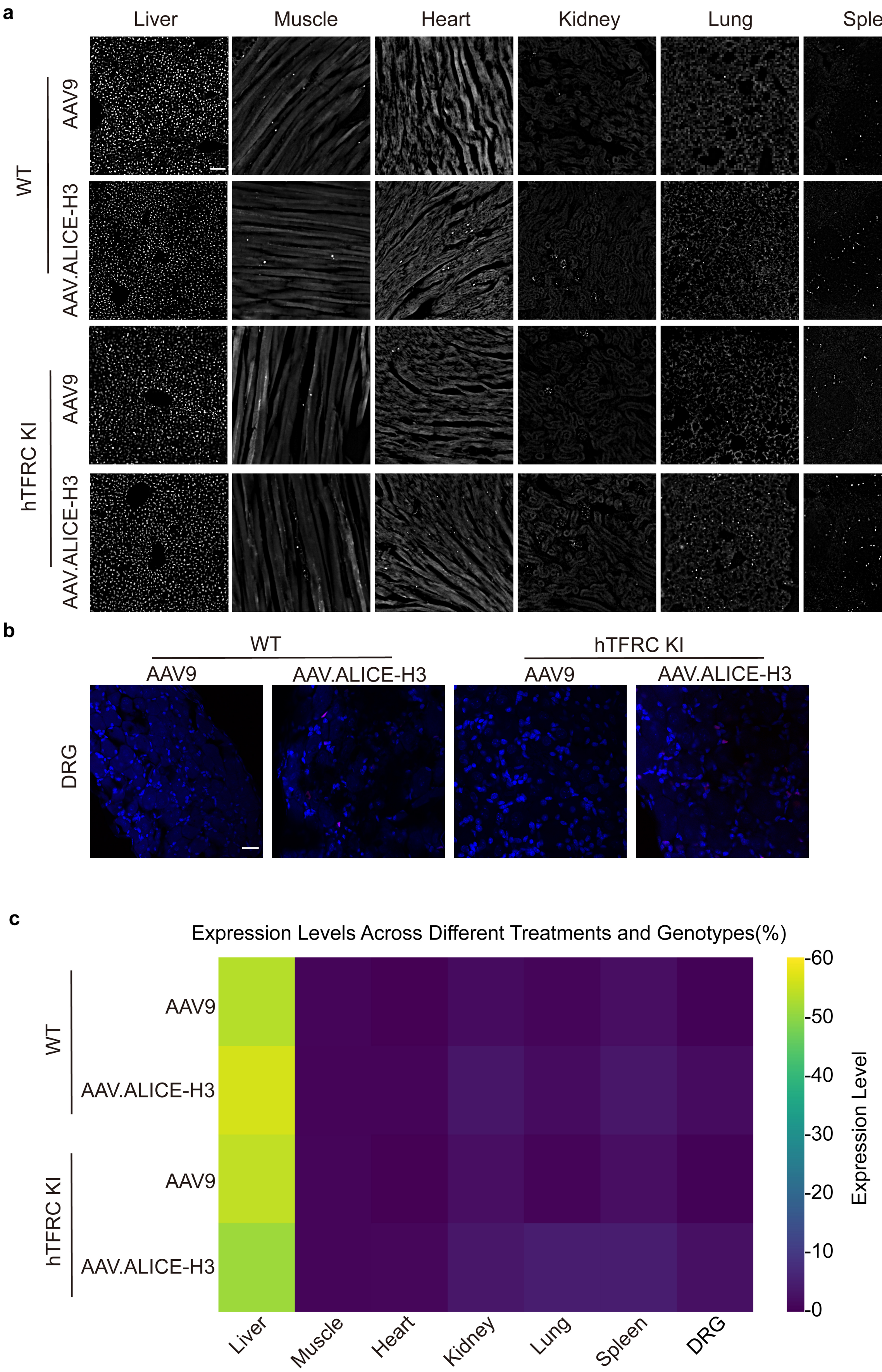
a
Liver
Muscle
Heart
Kidney
Lung
Spleen
WT
AAV9
AAV.ALICE-H3
hTFRC KI
AAV9
AAV.ALICE-H3
b
WT
hTFRC KI
AAV9
AAV.ALICE-H3
AAV9
AAV.ALICE-H3
DRG
c
Expression Levels Across Different Treatments and Genotypes(%)
WT
AAV9
AAV.ALICE-H3
hTFRC KI
AAV9
AAV.ALICE-H3
Liver
Muscle
Heart
Kidney
Lung
Spleen
DRG
-60
-50
-40
-30
-20
-10
-0
Expression Level

1890

**Supplementary Fig. 30 | Transduction efficiency of peripheral tissues by AAV9 and AAV.ALICE-H3.**

**a,** Representative images showing AAV.ALICE-H3 and AAV9-transduced cells in peripheral tissues, including liver, muscle, heart, kidney, lung, and spleen, are depicted in the illustrated figures. Scale bars, 100 μm.

**b,** Representative images showing AAV.ALICE-H3 and AAV9-transduced cells in dorsal root ganglia (DRG). Scale bars, 20 μm.

**c,** A heatmap representation of the transduction efficiency (H2A-RFP$^{+}$/DAPI$^{+}$ cells) of AAV9 and AAV.ALICE-H3 in the above peripheral tissues within hTFR1 KI mice and wild type mice. (n = 3 mice per group, mean is plotted).
